# Functional Genomic Complexity Defines Intratumor Heterogeneity and Tumor Aggressiveness in Liver Cancer

**DOI:** 10.1101/614057

**Authors:** So Mee Kwon, Anuradha Budhu, Hyun Goo Woo, Jittiporn Chaisaingmongkol, Hien Dang, Marshonna Forgues, Curtis C. Harris, Gao Zhang, Noam Auslander, Eytan Ruppin, Chulabhorn Mahidol, Mathuros Ruchirawat, Xin Wei Wang

**Affiliations:** Laboratory of Human Carcinogenesis and Liver Cancer Program, Center for Cancer Research, National Cancer Institute, Bethesda, Maryland 20892, USA; Department of Physiology, Ajou University School of Medicine, Suwon, 16499, Republic of Korea; Department of Biomedical Science, Graduate School, Ajou University, Suwon, 16499, Republic of Korea; Laboratory of Chemical Carcinogenesis, Chulabhorn Research Institute, Bangkok 10210, Thailand; Center of Excellence on Environmental Health and Toxicology, Office of the Higher Education Commission, Ministry of Education, Bangkok 10400, Thailand; Molecular and Cellular Oncogenesis Program and Melanoma Research Center, The Wistar Institute, Philadelphia, PA 19104, USA; Cancer Data Science Lab, National Cancer Institute, National Institute of health, MD 20892, USA

**Author notes:** Corresponding author, Contact information: Xin Wei Wang; Laboratory of Human Carcinogenesis and Liver Cancer Program, Center for Cancer Research, National Cancer Institute, Bethesda, Maryland 20892, USA.

**Keywords:** Hepatocellular carcinoma, intrahepatic cholangiocarcinoma, immune surveillance, functional aneuploidy, chromosomal instability

## Abstract

**Background:** Chronic inflammation and chromosome aneuploidy are major traits of primary liver cancer (PLC), which represent the second most common cause of cancer related death worldwide. Increased cancer fitness and aggressiveness of PLC may be achieved by enhancing tumoral genomic complexity that alters tumor biology.

**Method:** Here, we developed a scoring method, namely functional genomic complexity (FGC), to determine the degree of molecular heterogeneity among 580 liver tumors with diverse ethnicities and etiologies by assessing integrated genomic and transcriptomic data.

**Results:** We found that tumors with higher FGC scores are associated with chromosome instability and *TP53* mutations, and a worse prognosis, while tumors with lower FGC scores have elevated infiltrating lymphocytes and a better prognosis. These results indicate that FGC scores may serve as a surrogate to define genomic heterogeneity of PLC linked to chromosomal instability and evasion of immune surveillance.

**Conclusion:** Our findings demonstrate an ability to define genomic heterogeneity and corresponding tumor biology of liver cancer based only on bulk genomic and transcriptomic data. Our data also provide a rationale for applying this approach to survey liver tumor immunity and to stratify patients for immune-based therapy.

**STATEMENT OF SIGNIFICANCE:** Genomic heterogeneity contributes to therapeutic failure and poor outcome in patients with liver cancer and poses a challenge in defining targeted therapy. Our findings demonstrate an ability to define genomic heterogeneity and corresponding tumor biology of liver cancer based only on bulk genomic and transcriptomic data. Our data also provide a rationale for applying this approach to survey liver tumor immunity and to stratify patients for immune-based therapy. \body

## INTRODUCTION

PLC is the second leading cause of cancer-related mortality in the world ^1,2^. Hepatocellular carcinoma (HCC) and intrahepatic cholangiocarcinoma (iCCA) are two main histological subtypes of PLC, with molecular subtypes that differ in tumor biology and prognosis ^3–8^. Like other solid malignant tumors, HCC and iCCA are genomically, molecularly and biologically heterogeneous among individual tumors (inter-tumor) or within tumor lesions (intra-tumor) ^3,4,6,9–13^. Genomic instability and chromosome aneuploidy, two main cancer hallmarks found in human solid tumors ^14^, may be primary sources of cancer genomic diversity, which enable cancer cells to acquire mutations required for tumor fitness during carcinogenesis. Consequently, diverse tumor cell subpopulations are generated, resulting in both inter-tumor and intra-tumor heterogeneities (ITH) ^15^. We hypothesized that the ability of a premalignant cell to acquire genomic instability and chromosomal aneuploidy, giving rise to a more advanced tumor lesion, may determine the extent of ITH. Thus, a solid tumor mass with increased genomic complexity enriched with known hallmarks of cancer may determine its aggressiveness. Furthermore, recent genomic analysis at the single cell resolution indicates that aneuploidy occurs early in tumor evolution and may lead to extensive clonal diversity ^16^. However, a majority of genes in the loci with aneuploidy may be passengers ^17^. In contrast, chromosomal instability signature, inferred from the transcriptome, has been shown to associate with tumor metastasis and poor patient prognosis in diverse cancer types, indicating that its associated genes are more likely to be functional ^18^. These acquired traits collectively derived from the altered genome and functional networks, which we refer to as cancer functional genomic complexity (FGC), analogous to proposed functional variomics ^17^, may reflect the degree of ITH linked to tumor aggressiveness. In this study, we derived FGC in individual tumor based on the newly defined method and determined whether it is correlated with the degree of tumor aggressiveness in HCC and iCCA.

## RESULTS

### Patient correlation coefficient (PCC) defines functional genomic complexity (FGC)

Since it is generally accepted that somatic copy number alteration (SCNA)-dependent transcriptomic deregulation plays functional driver roles in cancer progression, we hypothesized that the estimate of the SCNA-dependency of transcripts for each patient may reflect genomic complexity with functional influence. We postulated that the SCNA-dependency of transcripts in individual sample could be estimated based on the correlation between SCNAs and their corresponding transcriptome among the globally correlated features. For this, we first performed global correlation analysis between transcriptome and DNA copy number in the TIGER-LC cohort ^4^ and selected significantly correlated features using a conservative approach (Methods and Fig 1A). In the overall distribution of global correlation estimates, we found a shift towards positive correlation of the tumor (T) (shaded curves) samples of Thai HCC and iCCA, compared to their corresponding non-tumor (NT) specimens (dotted areas) (Figure S1A-B, left panels). A positive correlation was more apparent in T samples than NT samples when we compared correlation coefficient value among the selected features (Methods and Figure S1A-B, right panels), indicating that molecular features associated with SCNA-dependent transcripts are tumor-specific. Using these features, we computed PCC per sample (Methods and Table S1a-S1c). To determine the biological relevance of the PCC with genomic complexity, we examined the PCC associated genes (Fig 1B). Briefly, we performed correlation analysis between PCC and all the transcriptome features and selected positively or negatively associated genes based on the correlation estimate and p-value (above top 5% or below bottom 5% of correlation estimate and p-value < 0.05). Gene Ontology (GO) analysis revealed that DNA repair and cell cycle related biological processes are enriched by PCC positively-associated genes, while immune response pathways are enriched by PCC negatively-associated genes (Fig 1C). Consistent results were also found when we analyzed Thai HCC and iCCA separately as well as TCGA HCC (Figure S2A-F). Since most of over-represented processes were closely related with aneuploidy and chromosomal instability, such as cell cycle and DNA damage response (DDR), we determined the association of PCC with chromosomal instability (CIN) score, which was calculated based on the copy number data only (Supplementary Materials and Methods and Table S1a-S1b). As expected, we observed a strong association between CIN and PCC (Fig 1D and Figure S3A-C). Consistent results were also obtained when we compared the genomic instability (GIN) scores, the length of chromosomes with SCNA (Supplementary Materials and Methods and Table S1a), as another estimate of genomic instability. We found a strong correlation between overall GIN scores and PCC values (Figure S3D-E). In addition, we found no evidence of preferential allelic gain or loss linked to PCC, when we compared the association of PCC with the amplified (CIN_ampl_) and deleted (CIN_del_) scores for individual samples (Supplementary Materials and Methods and Figure S4A-B). Interestingly, we found a strong association between CIN_ampl_ and CIN_del_ in both Thai HCC and iCCA (Figure S4C-D). Consistent trends were also observed in the broad chromosomal arm-level SCNA analyses in HCC and iCCA when we used the GISTIC 2.0 algorithm (Figure S4E-F). These results suggest that gains or losses of specific chromosomal regions may be a consequence of tumor cells acquiring genomic instability and then being selected during tumor evolution. However, it is likely that a majority of genes in the loci with aneuploidy may be passengers. Therefore, we determined whether the PCC also reflected the aneuploidy of the genes with functional role. For this, we compared the PCC with the estimate of total functional aneuploidy (tFA) (Table S1a-1b) ^18^, which was inferred from transcriptome data only. Briefly, we calculated tFA in each sample based on the coordinated aberrations in the expression of genes, which were localized to each chromosomal region, by adapting the method in Carter’s paper ^18^. Since tFA was the estimate of the aneuploidy reflected transcriptome level and more likely to be functional, we examined the functional relevance between PCC and tFA. In the comparison of GO analysis performed with significantly tFA associated genes, we found functional similarity of PCC (Figure S5A-F). Also, we found a strong association of PCC with tFA in TIGER-LC cohort (Fig 1E). A strong association maintained when Thai HCC and iCCA were examined separately as well as TCGA HCC (Figure S5G-I). Moreover, we found a high concordance among CIN, tFA and PCC scores (Fig 1F and Figure S6). Since immune response pathways were enriched by PCC negatively-associated genes, we examined the relevance of PCC with cancer immunity. Many studies have been reported the association of anti-tumor activities of effector T cells at the tumor sites with patient prognosis in solid tumor ^19,20^, we focused on the immune effector activity of the local immune infiltrates. For this, we calculated immune cytolytic activity (ICYT) based on the transcript levels of two key cytolytic effectors, granzyme A (*GZMA*) and perforin (*PRF1*)^21^. Consistent with the GO analysis with negatively PCC associated genes, we found a strong inverse correlation between ICYT and PCC in the tumor tissues. However, there was no association in adjacent non-tumor tissues in Thai HCC and iCCA (Fig 1G-H), indicating that these associations were only linked to local immune infiltrates in tumor and PCC may reflect the immune surveillance in tumor tissue. Taken together, these results indicate that PCC represents functional genomic complexity (FGC) conversantly reflecting both chromosome level and transcriptome level and selection of drivers during tumorigenesis. Therefore, we named PCC as “FGC” to refer the newly defined genomic complexity estimate in individual sample.

**Figure 1.**
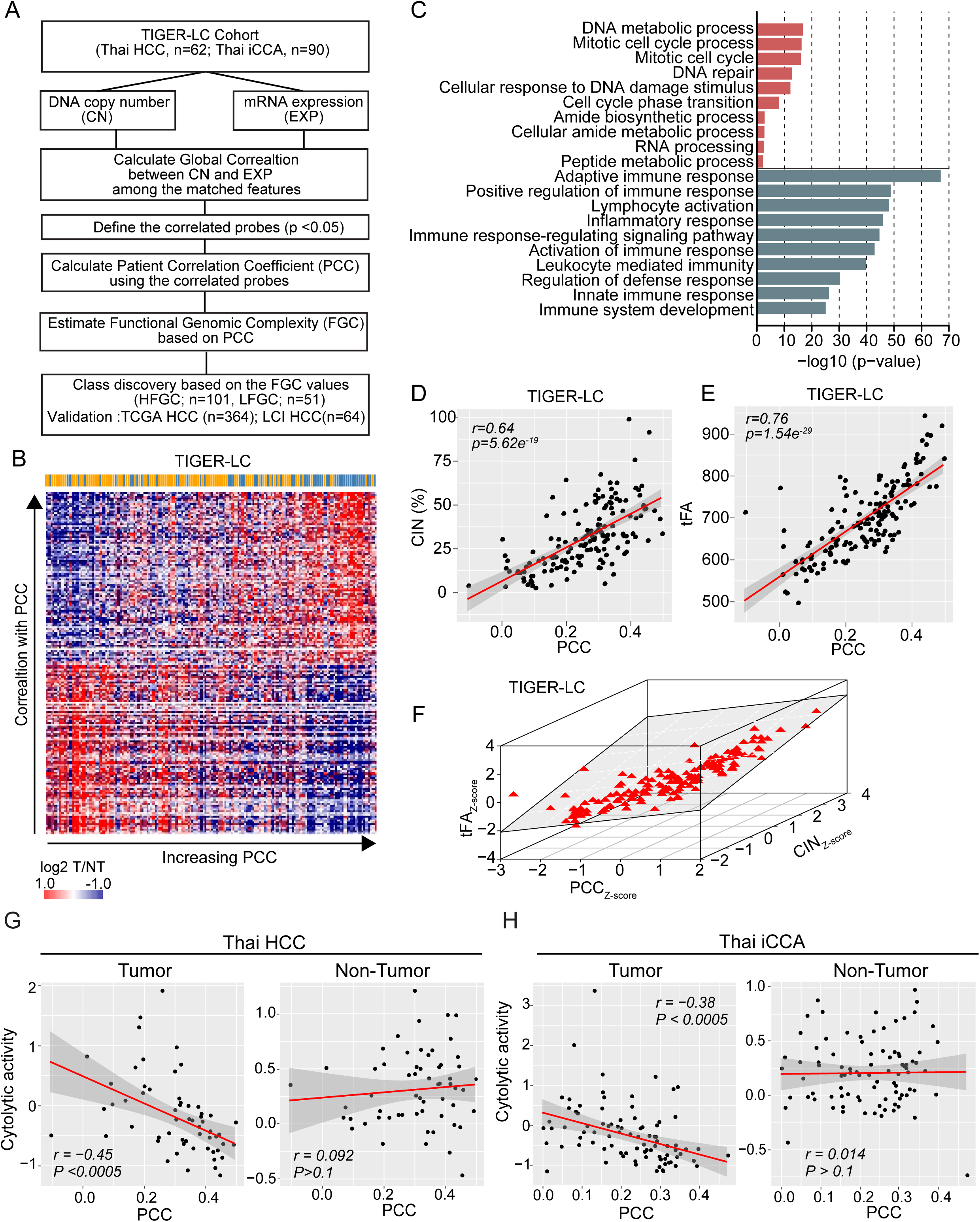
Patient correlation coefficient (PCC) defines functional genomic complexity (FGC). (A) Schematic overview of the study design. (B) Heatmap shows the expression level of positively or negatively PCC associated genes (> 95% or < 5% of correlation estimate and p-value <0.01) in TIGER-LC. Samples are represented according to the PCC increasing order in columns and genes were represented according to the decreasing of correlation order in coefficient in the row. Color bar indicates HCC and iCCA patients as in blue and orange color, respectively. (C) GO Enrichment Analysis of selected genes. Top10 ranked GO based on the precision rank were shown. Orange and green color indicate positively and negatively correlated gene sets, respectively. (D-E) PCC shows strong correlation with CIN (D) and tFA (E). Coefficient estimates and p-value based on Pearson’s correlation were depicted. (F) Collective association among PCC (x axis), CIN (y axis), and tFA (z axis) are shown. (G-H) The association between PCC and Immune Cytolytic activity (ICYT), defined as log-average of GZMA and PRF1 expression, derived from tumor (left panels of each) or non-tumor (right panels of each) tissue of Thai HCC and Thai iCCA are shown, respectively.

### Stratification PLC patients based on the FGC shows the clinical significance of FGC

To evaluate the clinical relevance of FGC, we divided 152 cases into a high (HFGC) or low group (LFGC) based on the FGC value (≥0.2) (left panel of Fig 2A and Table S1a) and found a significant difference in overall survival between HFGC and LFGC, with HFGC being more aggressive than LFGC (Fig 2B). It was noted that more HCC cases were found in HFGC whereas more iCCA cases were found in LFGC (Fig 2A, right panel) and the difference was statistically significant (p-value of 2.42e^-6^, Wilcoxon rank sum test). We observed similar results when Thai HCC and iCCA cohorts were analyzed separately (Figure S7A-D and Table S1a) and in the Cancer Genome Atlas (TCGA) HCC cohort and LCI Chinese HCC cohort (Figure S7E-H and Table S1b-S1c), indicating that FGC scores are robust indicators of PLC prognosis. Furthermore, HFGC had a significantly higher level of tFA than LFGC (Fig 2C and Figure S8A-C). Evidence for the difference in genomic instability between HFGC and LFGC was also found among SCNA of individual samples with a higher magnitude of SCNA values observed in the correlated segments in HFGC than LFGC (Fig 2D-E and Figure S9A-D). When we compared the GIN length between HFGC and LFGC, we found that HFGC had much higher GIN score, showing significant difference in both gain and loss compared to LFGC (Figure S9E-F). Upon examining allelic imbalance, only HFGC samples contained SCNA regions with loss of heterozygosity (LOH) in Thai HCC and iCCA (Fig 2F). Interestingly, the total number of peak segments was noticeably higher in HCC HFGC than iCCA HFGC (Figure S9G). We found more deleted segments with LOH (DEL W/LOH) than amplified segments with LOH (AMP W/LOH). Furthermore, the regions for copy neutral LOH (CN LOH), i.e., segments with LOH without SCNA, were only found in HFGC in HCC and iCCA (Figure S9H, upper panel). Moreover, CN LOH was distributed throughout all genomic regions of iCCA HFGC, while focal enrichment of CN LOH in specific chromosomal arms was found in HCC HFGC (Figure S9H, lower panel). Considering the likelihood of CN LOH to activate oncogenes and to unmask tumor suppressers ^22^, the high incidence of CN LOH in the HFGC subtype implies a strong contribution of genomic instability to carcinogenesis.

**Figure 2.**
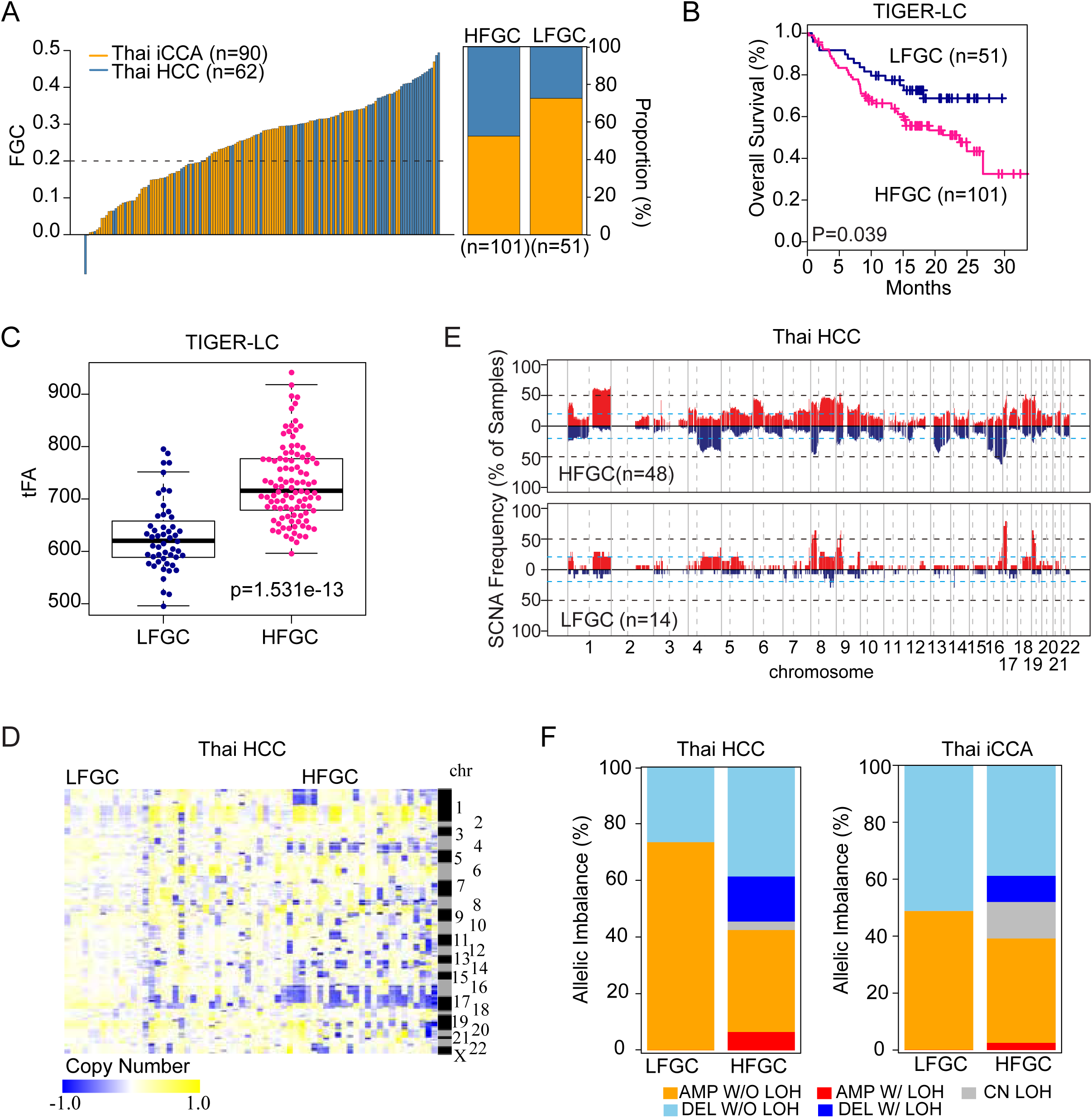
Stratification of PLC patients based on the FGC. (A) FGC values among the TIGER-LC cohort (n=152) are plotted in a rank order (left). By applying cut-off (dotted line, FGC=0.2), TIGER-LC cohort were separated into a FGC high (HFGC; n=101) and FGC low (LFGC; n=51) group. Relative proportion of HCC and iCCA among HFGC and LFGC are shown (right). HCC and iCCA patients are shown in blue and orange color, respectively. (B) Kaplan-Meier (KM) survival analysis based on the HFGC and LFGC subtype show significant difference of overall survival time. The statistical P value by the Cox-Mantel log-rank test were depicted. (C) HFGC shows higher tFA values compared to the LFGC. P-value based on Welch two sample t-test was depicted. (D) Heatmap shows copy number value of individual sample of Thai HCC corresponding to the correlated segments regions, respectively. Samples are grouped by the HFGC and LFGC in columns and segment regions are represented in rows according to the chromosomal location. (E) The frequency of SCNA among HFGC and LFGC subtype of Thai HCC are plotted corresponding to the correlated segments region, respectively. The sample frequencies with copy number gain and loss (log2 (copy number) >0.2 or log2 (copy number) < −0.2) are shown in red and blue, respectively. Chromosome boundaries and centromere position are indicated by vertical solid and dashed lines, respectively. Horizontal dashed blue lines indicate frequency of 50%. Horizontal dotted black lines indicate frequency of 20%. (F) Comparison of allelic imbalance frequency between HFGC and LFGC. Proportion of allelic imbalance of HFGC and LFGC of Thai HCC and Thai iCCA are plotted.

### Determination of candidate drivers based on the differentially expressed genes (DEGs) between HFGC and LFGC

To determine additional candidate drivers of HCC and iCCA, we selected DEGs between HFGC and LFGC (permutation t-test p< 0.005 and fold change > 0.5) among the globally correlated genes (Fig 3A-B). We found that the upregulated genes in HFGC were significantly enriched by cell cycle and DNA replication process-related genes, while immune response-related genes were down-regulated (Fig 3C-E and Table S2), with similar results found in TCGA HCC (Figure S10A-B and Table S3). Consistently, the gene set enrichment analysis (GSEA) demonstrated considerable number of overlapped pathways between Thai iCCA and HCC among the top-ranked 40 pathways of biological process, KEGG pathway, and oncogenic pathway (15, 29, and 27 pathways, respectively) (Fig 3F, Figure S11, and Table S4-5). Also, consistent results were found in TCGA HCC (Figure S10C-D and Table S6). These results suggest that tumor biology may differ between HFGC and LFGC regardless of PLC tumor type (Table S7).

**Figure 3.**
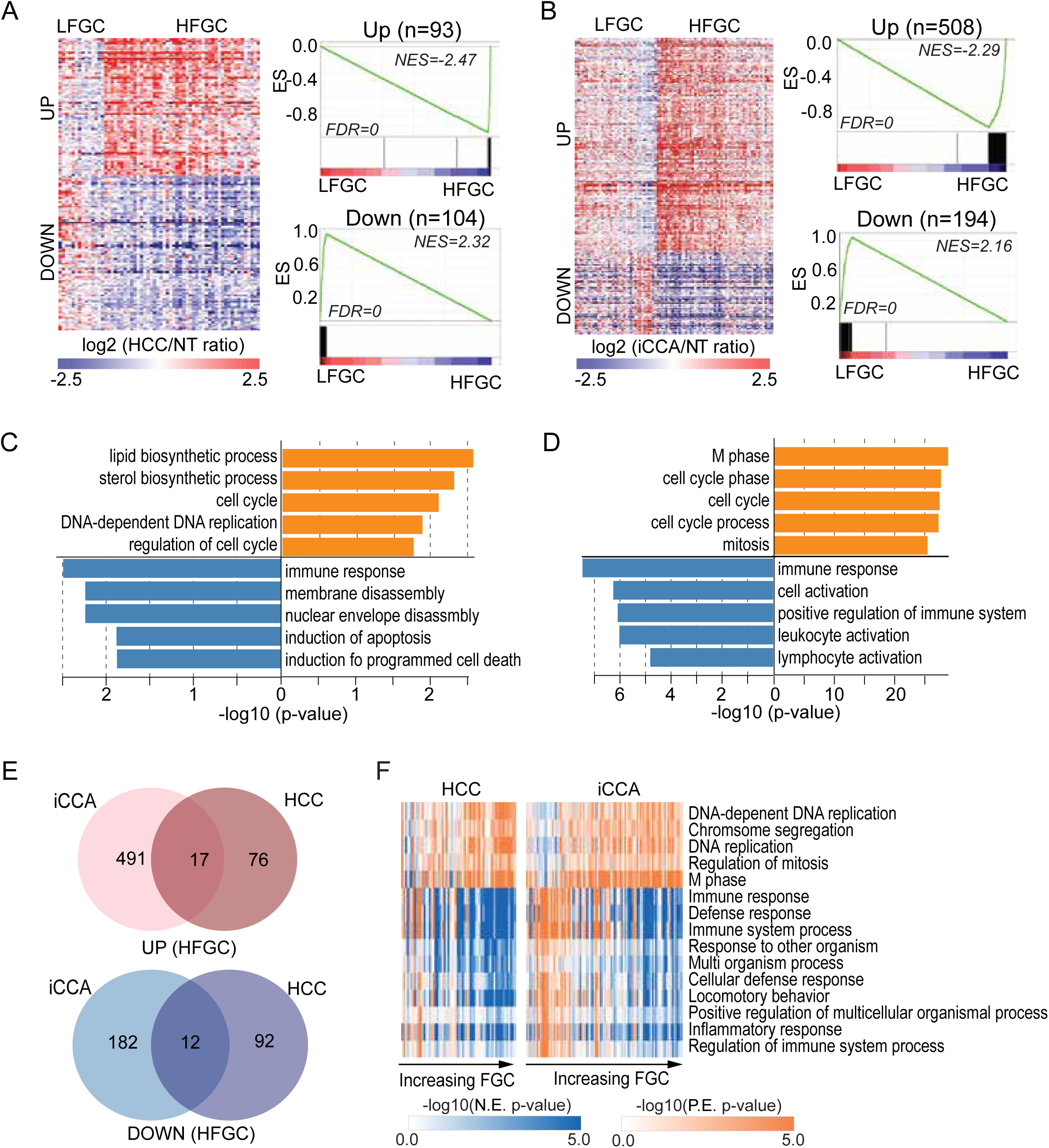
Differentially expressed genes (DEG) between HFGC and LFGC. (A-B) (Left panel of each) A heatmap shows the expression of DEG between HFGC and LFGC of Thai HCC (A) and Thai iCCA (B) Samples are represented in columns, grouped by HFGC and LFGC, and up-or down-regulated genes are represented in rows. (Right panel of each) Enrichment plots based on the 93 upregulated genes and 104 down-regulated genes are shown in the upper and lower panel, respectively. Normalized Enrichment scores (NES) and FDR for each gene set were noted. (C-D) GO analysis performed based on the DEGs of Thai HCC and Thai iCCA, respectively. The –log10 (p-value) is shown in orange and blue bars for up-and down-regulated genes, respectively. (E) Venn diagrams show the overlapping genes between DEG of Thai HCC and of Thai iCCA. Up-and down-regulated genes are analyzed separately. (F) Heatmaps show the ssGSEA of Thai HCC (left) and Thai iCCA (right) based on the gene sets derived from the biological process (BP) gene sets in Molecular Signatures Database (MSigDB database v5.2). The overlapping gene sets significantly enriched in both Thai HCC and Thai iCCA are shown. The P-value from the Kolmogorov-Smirnov (ks) test was transformed in -log scale and used in the plot. Samples were in columns according to the increasing order of FGC value and the log transformed p-value for each gene set is represented in rows.

### TP53 mutations are associated with increased FGC in PLC

Next, we examined whether specific gene mutations are associated with FGC. A total of 100 mutated genes were commonly found in HCC and iCCA (Table S8-10). Noticeably, patients with higher FGC scores exhibited more frequent *TP53* mutation, particularly in nonsynonymous mutations (Fig 4A and Figure S12A). Most *TP53* mutations were missense, located within its DNA binding domain (DBD) region (74.1%), while other mutations outside of the DBD were frameshift (18.52%) or nonsense mutations (Fig 4B, Figure S12B, and Table S11). Interestingly, *TP53* mutations at the DBD regions, including the most frequent R249S mutation in HCC, were more frequent in HFGC than LFGC tumors (Fig 4B, Figure S12B and S12E). Moreover, Thai HCC and iCCA cases with *TP53* mutations had a much worse survival than those with wild type p53 (Fig 4C-D) with comparable results found in TCGA HCC cases (Fig 4E). Since *TP53* DBD mutations are considered as a gain-of-function mutant p53 protein (mutp53) involved in the maintenance of genome integrity ^23–26^, it is plausible that the *TP53* mutations may result in increased genomic instability, leading to elevated FGC scores. Indeed, cases with *TP53* mutation were significantly enriched in HFGC (Fig 4F and Table S12). Also, patients with *TP53* mutation showed much higher CIN and FGC than wild type p53 (Fig 4G-H, Figure S12C-D, and S12F-G). Taken together, we suggest that tumors with higher FGC scores may originate from genomic instability due to p53 mutations, leading to carcinogenesis.

**Figure 4.**
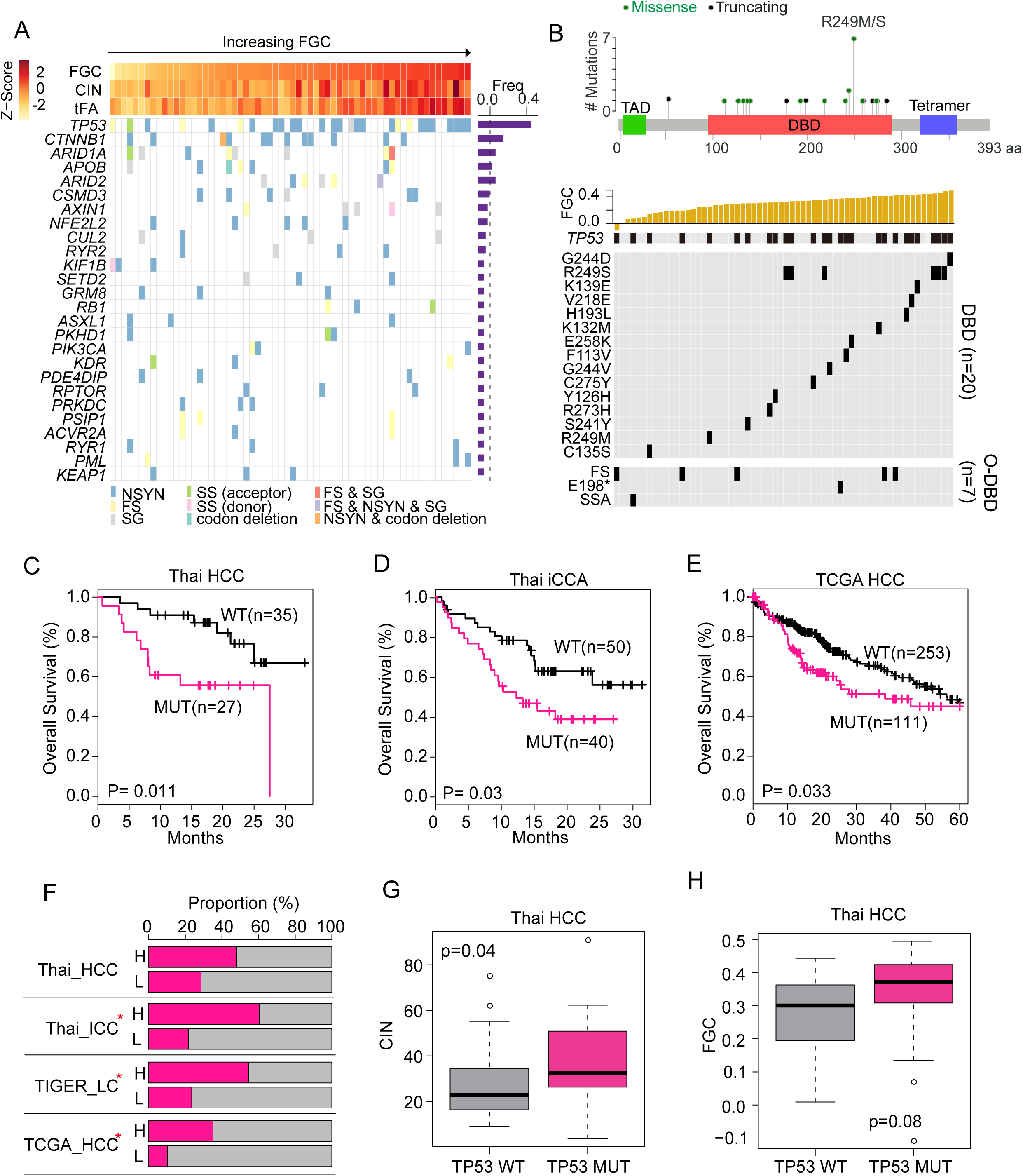
*TP53* functions as a cancer functional genomic complexity (FGC) driver. (A) (Top panel) Z-scores for FGC, CIN, and tFA in each HCC sample are plotted in each barplot in the FGC ranked order. (Bottom panel) The mutation frequency for 26 genes, mutated in more than 3 samples of Thai HCC, was shown (right panel). The occurrence of mutation of regarding gene in each sample and mutation types were indicated in different colors. Samples were represented in columns in the same order of top panel. (B) (Top panel) *TP53* mutations sites among Thai HCC were shown. Transactivation motif (TAD; 6-29), DNA binding motif (DBD; 95-288), and tetramerisation motif (Tetramer; 318-358) were depicted in different colored box; green, orange, and navy, respectively. Green or black dots indicates missense or truncating mutation, respectively. (Bottom panel) Top plot indicates the FGC score of each sample in the rank order. *TP53* mutation incidence in each sample were plotted in black according to the mutation sites. Mutation sites of *TP53* depending on DNA binding domain (DBD) and out of DBD (O-DBD) were plotted separately. (C-E) KM survival analysis based on *TP53* mutation status in the Thai HCC (C), Thai iCCA (D), and TCGA HCC (E) patients. (F) Proportion of occurrence of *TP53* mutation in HFGC and LFGC of TIGER-LC, Thai HCC, iCCA, TCGA HCC were shown. H or L indicates HFGC and LFGC, respectively. Significant enrichment of *TP53* mutation were marked with red star based on the Fisher’s exact test (p-value <0.05). (G-H) The CIN (G) and FGC (H) level between *TP53* WT and *TP53* mutation among Thai HCC. P-values based on the Welch two-sample t-test were depicted. NSYN, non-synonymous mutation; FS, frame shift mutation; SS, splice site mutation; NS, non-sense mutation; SSA, splice site acceptor.

### Association between FGC and Tumor infiltrating lymphocytes (TILs) in PLC

To analyze the extent of tumor infiltrating lymphocytes (TILs) in the TIGER-LC cohort, we applied the CIBERSORT (Supplementary Materials and Methods) ^27^ and calculated the immune score as the summation of transformed value of 22 TILs based on the CIBERSORT output (Supplementary Materials and Methods). Consistent with the close association between FGC and ICYT, we found a strong inverse correlation between immune scores (Supplementary Materials and Methods) and FGC in the tumor tissues. However, there was no association in adjacent non-tumor tissues in Thai HCC and iCCA (Figure S13A-B), indicating that the increase in TILs is tumor-specific. We found that TILs in LFCG were more actively associated each other compared to HFGC in both iCCA and HCC (Figure S13C-D). To determine FGC-associated TILs in either HCC or iCCA, we divided TILs into favorable or adverse TILs regarding to the enrichment of LFCG or HFGC. Specifically, a subset of favorable TILs is linked to LFGC, while a subset of adverse TILs is linked to HFGC cases (Supplementary Materials and Methods and Fig 5A and 5C). Interestingly, we found that immune scores with adverse TILs were positively associated with FGC, while immune score with favorable TILs showed inverse association with FGC (Fig 5B and 5D). This relationship is robust regardless of PLC subtype, indicating that increases in specific TILs may be a consequence of tumor cell-related FGC. Among adverse TILs, Tregs, NK cells and DC cells were consistently and significantly elevated in HFGC from TIGER-LC and TCGA HCC, suggesting that these immune cells may functionally contribute to tumors with an increased FGC (Fig 5E-F and Figure S13E). Currently, a number of immunomodulatory molecules are under investigation for immunotherapy in various cancer ^28^. To search for promising candidates of immunotherapy for PLC, we examined the expression profiles of the 54 genes with immunostimulatory or immunoinhibitory function. Most genes showed remarkable features, which were highly upregulated in LFGC but down-regulated in HFGC (Figure S14A-B, top panels), and showed negative association with FGC (Figure S14A-B, bottom panels) in TIGER-LC. Furthermore, several genes expressed much higher in LFGC than those in HFGC (Figure S14C-D) have been reported to be involved in the development and progression of liver cancer ^29–32^. Also, we found that *PDCD1LG2* (PD-L2), ligand for PD-1, was up-regulated in LFGC compared to HFGC indicative of the sensitivity to the immune-checkpoint inhibitor (ICI) _33_.

**Figure 5.**
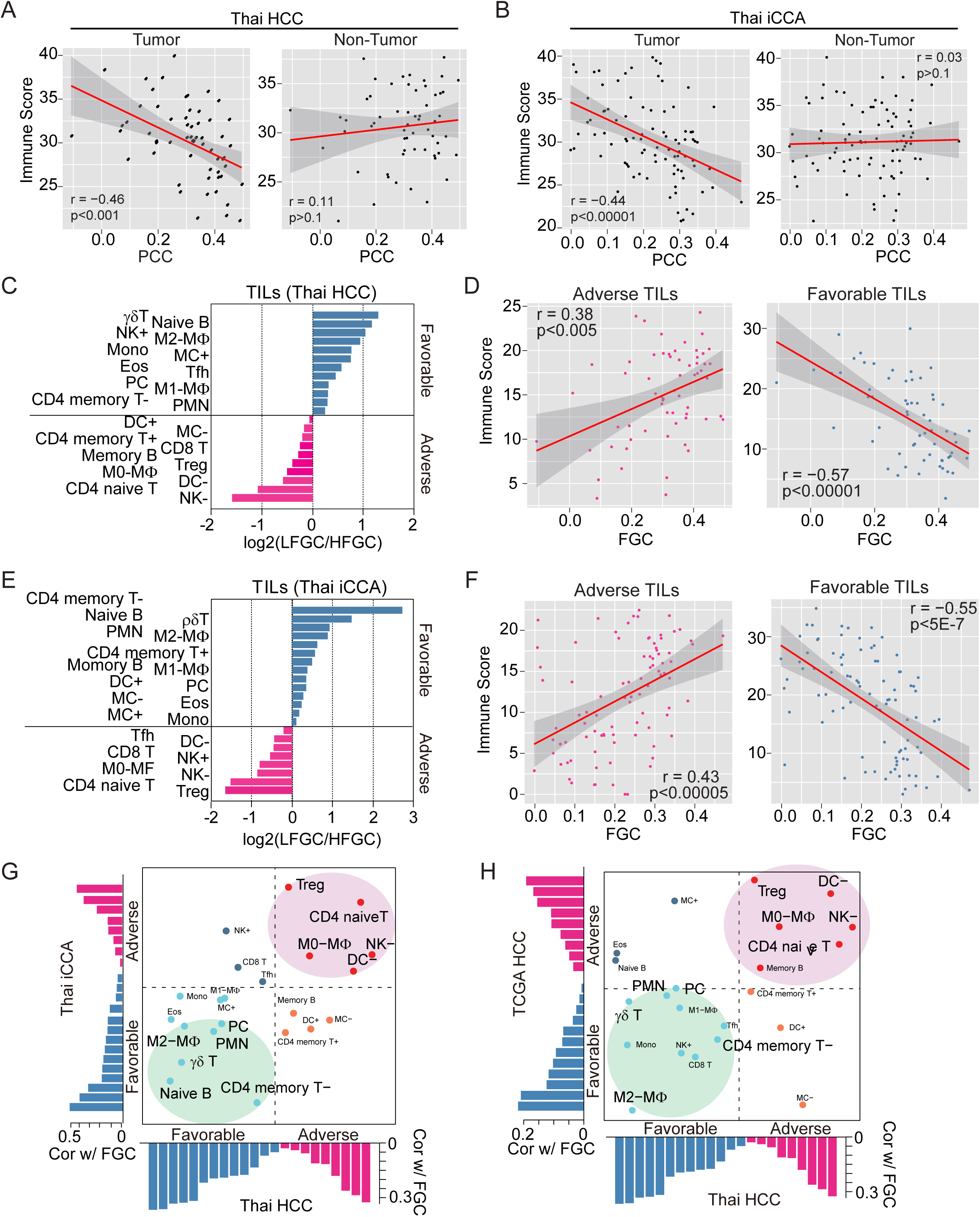
Tumor infiltrating lymphocytes (TILs) are related to FGC. (A-B) The association between PCC and immune score derived from tumor (left panel of each) or non-tumor (right panel of each) tissue of Thai HCC and Thai iCCA are shown, respectively. (C and E) Log2 ratios of the mean of 22 types of TILs in LFGC to HFGC of Thai HCC (C) and Thai iCCA (E) are shown, respectively. TILs enriched in the LFGC or HFGC group are assigned as favorable or adverse and plotted as either blue or pink bar, respectively. (D and F) The association between FGC and the immune score of adverse (left panel) or favorable TILs (right panel) among Thai HCC (D) and Thai iCCA (F), derived from Fig. 5C and 5E, are shown, respectively. (G-H) Concordance and difference of the associations of FGC with 22 types of TILs between Thai HCC and Thai iCCA (G) or Thai HCC and TCGA HCC (H) are shown, respectively. The x axis and y axis indicate the coefficient estimates from Pearson’s correlation between FGC and the relative proportion of TIL subpopulations in Thai HCC and Y-axis represents for coefficient estimates from Pearson’s correlation between FGC and the relative proportion of TIL subpopulations in Thai iCCA or TCGA HCC. Red and blue bars indicate adverse and favorable associations, predicted by the association with FGC, respectively. Pink or green circled areas indicate TILs commonly associated with FGC among Thai PLC and TCGA HCC.

### FGC may be useful to predict patients’ responses to ICI-based immunotherapy

Also, to examine whether increased FGC may be associated with ICI response, we analyzed the TCGA melanoma dataset as liver cancer-related ICI dataset is not available ^34^. To infer the FGC level in each patient, we calculated tFA as a surrogate for FGC. Consistently, we found that high tFA was correlated with poor survival in melanoma cohort (Figure S14E and Table S13a). Since patients who were treated with anti-CTLA-4 therapy were included in the TCGA melanoma dataset, we performed the Kaplan-Meier (KM) survival analysis by excluding them separately (Supplementary Materials and Methods) and found similar result (Figure S14F). Besides, the tFA level of the non-responders to the anti-CTLA-4 therapy was much higher compared to responders (Figure S14G), suggesting that the tFA level was closely associated with the efficacy of anti-CLTA-4 therapy. Similar trends were found in the analysis performed in the cohort of patients with metastatic melanoma who were treated with anti-PD-1 immunotherapy^35^ (Figure S14H and Table S13b). Overall, these results indicate that FGC could be useful to predict patients’ responses to ICI-based immunotherapy.

## DISCUSSION

PLC is clinically and molecularly heterogeneous and is highly refractory to treatment. The extensive heterogeneity of PLC may be due to the presence of complex etiological factors that causes chronic liver diseases. A common denominator at its origin is a perpetual wound-healing response triggered by parenchymal cell death, with an ensuing inflammatory cascade and concomitant fibrosis progression ^36^. In addition, inflammation-induced replication stress promotes DNA damage, which subsequently induces DDR, genomic instability and finally tumorigenesis ^37^. Thus, we hypothesized that chronic inflammation may be the main initiator of hepatocarcinogenesis through induction of genomic instability and aneuploidy. Consistently, *TP53* mutations may be a key driver for inflammatory-mediated hepatocarcinogenesis since it is the most frequently mutated gene in PLC ^38^. We found PLC cases with increased FGC enrich for TP53 mutations, consistent with the hypothesis that p53 inactivation may be a trigger to drive chromosome instability and consequently increased ITH. Interestingly, we found that tumors with different FGC scores have different immune cell infiltrates. It is conceivable that the immune surveillance program may help to remove cells with aneuploidy, perhaps by recognizing specific cell surface antigens such as calreticulin, to suppress the growth of cancer cells ^39^. However, increased ITH due to extended aneuploidy could overwhelm the immune system, possibly leading to T cell exhaustion ^40^. Chronic inflammation may activate several pathways to evade immune surveillance, providing an environment that is inhibitory to productive anti-tumor immune responses. Several types of immune suppressive cells, such as Tregs, associated with higher grade and poorly differentiated HCC with unfavorable outcome, may significantly undermine sustained cytotoxicity mediated by T and NK cells, allowing tumor cells to escape immune surveillance ^41,42^. Our results indicate that a unique landscape of immune cells is associated with FGC-high tumors, consistent with the central role of immune surveillance in hepatocarcinogenesis.

A recent study showed that aneuploidy of tumor cells adversely affected the immune cell reaction against tumors, suggesting an association of SCNA with immune evasion and the relevance of immunotherapy ^43^. In addition, checkpoint inhibitor-based immunotherapies targeting the regulatory pathways of T cells, cytotoxic T-lymphocyte associated antigen 4 (CTLA-4) (ipilimumab) and programmed cell death protein 1 (PD-1) (e.g., nivolumab or pembrolizumab), have enhanced anti-tumor activity with significant clinical benefit in patients with various cancers ^44^ including HCC ^45^. While it is unclear why most HCC patients do not benefit from immunotherapy, an ineffective immune surveillance program due to immunosuppressive mechanisms that are functional in tumor cells or the tumor-educated liver microenvironment has been suggested. Moreover, it has been difficult to identify predictive biomarkers of response to these agents. Our results show that the FGC score can distinguish TIL subpopulations associated with genomic instability. Therefore, we suggest that an FGC score may be a simple and reliable predictive indicator to stratify patients for immune therapy using only bulk genomic and transcriptomic analysis.

## METHODS

### Data sets

A previously described cohort of a set of 398 surgical paired tumor and non-tumor specimens derived from 199 patients of the TIGER-LC cohort (130 iCCA patients and 69 HCC patients) with publicly available Affymetrix Human Transcriptome Array 2.0 data and Affymetrix Genome-Wide Human SNP Nsp/Sty 6.0 data (NCBI GEO accession number GEO: GSE76297 and GSE76213, respectively) were used in this study ^4^. Somatic single nucleotide variants and small insertions and deletions among the TIGER-LC cohort were identified based on NCI OncoVar V4, an Agilent SureSelect Custom DNA kit (Agilent Technologies) targeting 2.93 Mb of sequence in 562 genes and publicly accessible (dbGAP (https://ww.ncbi.nlm.nih.gov/gap) Accession No. phs001199.v1.p1). For validation cohort, HCC cohort of 247 Chinese patients from LCI ^46^ and TCGA LIHC cohort with 377 HCC patients were used. To validate the association between FGC and immunotherapy with immune checkpoint blockade (ICB), we used transcriptome data from skin cutaneous melanoma datasets derived from TCGA_SKCM^34^ study (n=472) and metastatic melanoma from Hugo^35^ study (n=28). Data preprocessing of validation sets were described in the Supplementary Material and Methods.

### Calculation patient correlation coefficient (PCC)

We calculated patient correlation coefficient based on the pre-selected significantly associated features. To select significantly correlated global features, we calculated the global correlation coefficients and global correlation p-value based on the total transcriptome probes and corresponding genomic segments. Also, MAD of copy number values among the 64,597 transcript probes were calculated. Based on the global correlation p-value and MAD, significantly correlated features of transcriptome probes and corresponding segmented regions were selected by following cutoff by optimizing stratify patients. For Thai HCC and iCCA, p-value < 0.05 & MAD > 20% of overall distribution; for LCI HCC, p-value <0.0005 & MAD >10% for LCI cohort; for TCGA HCC, p-value <0.01 for TCGA HCC were applied to select for further analysis. Using only features that met p-level and MAD cutoffs, patient level correlation was calculated.

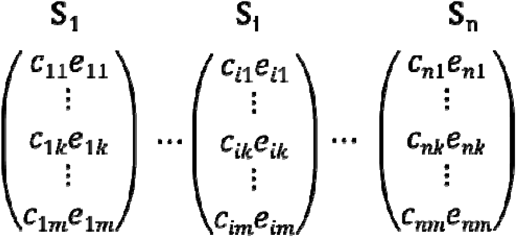

Where c is SCNA and e is expression for feature and n is the n^th^ sample and m is m^th^ feature Patient level correlation coefficient PCC was calculated based on the selected paired transcriptome probe and copy number value of segments for individual patient

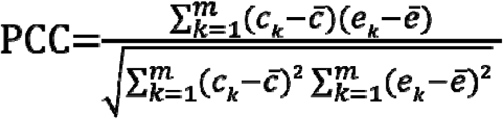

Where m is the number of selected features and c is the SCNA value and e is expression value for selected feature and is the mean of c and is the mean of e PCC is the measure of enrichment of the numbers and types of molecular features from genomic instability in individual sample.

Full methods and any associated references are provided in the Supplementary Information.

## Abbreviations

PLC: primary liver cancer;
FGC: functional genomic complexity;
HCC: Hepatocellular carcinoma;
iCCA: intrahepatic cholangiocarcinoma;
ITH: intra-tumor heterogeneities;
SCNA: somatic copy number alteration;
T: tumor;
NT: non-tumor;
PCC: patient correlation coefficient;
GO: Gene Ontology;
DDR: DNA damage response;
CIN: chromosomal instability;
tFA: total functional aneuploidy;
TILs: tumor infiltrating lymphocytes;
GIN: genomic instability;
HFGC: high functional genomic complexity group;
LFGC: low functional genomic complexity group;
TCGA: the Cancer Genome Atlas;
LOH: loss of heterozygosity;
DEL W/LOH: deleted segments with LOH;
AMP W/LOH: amplified segments with LOH;
CN LOH: copy neutral LOH;
GSEA: gene set enrichment analysis;
DBD: DNA binding domain;
Tregs: regulatory T cell;
NK: natural killer cell;
DC: dendritic cell;
KM: Kaplan-Meier (KM);
CTLA-4: cytotoxic T-lymphocyte associated antigen 4;
PD-1: programmed cell death protein 1.

## Acknowledgements

We thank Drs. Edward Gertz, Alejandro Schaffer and Snorri Thorgeirsson for their invaluable comments and critical reading of the manuscript.

## Author’s contributions

SMK and XWW conceived the idea. SMK carried out most of the data analysis. AB, HGW, JC, HD, MF, GZ, NA, ER performed data processing and advised experimental approaches. AB, JC, MR, XWW performed data management for the TIGER-LC consortium. SMK and XWW interpreted data and prepared the manuscript. All authors read, advised and approved the manuscript.

## Consent for publication

Not applicable.

## REFERENCES

1 Torre, L. A. et al. Global cancer statistics, 2012. CA Cancer J Clin 65, 87–108, doi:10.3322/caac.21262 (2015).

2 Global Burden of Disease Cancer, C. et al. Global, Regional, and National Cancer Incidence, Mortality, Years of Life Lost, Years Lived With Disability, and Disability-Adjusted Life-years for 32 Cancer Groups, 1990 to 2015: A Systematic Analysis for the Global Burden of Disease Study. JAMA Oncol 3, 524–548, doi:10.1001/jamaoncol.2016.5688 (2017).

3 The Cancer Genome Atlas Research Network. Comprehensive and Integrative Genomic Characterization of Hepatocellular Carcinoma. Cell 169, 1327–1341 e1323, doi:10.1016/j.cell.2017.05.046 (2017).

4 Chaisaingmongkol, J. et al. Common Molecular Subtypes Among Asian Hepatocellular Carcinoma and Cholangiocarcinoma. Cancer Cell 32, 57–70 e53, doi:10.1016/j.ccell.2017.05.009 (2017).

5 Jusakul, A. et al. Whole-Genome and Epigenomic Landscapes of Etiologically Distinct Subtypes of Cholangiocarcinoma. Cancer Discov 7, 1116–1135, doi:10.1158/2159-8290.CD-17-0368 (2017).

6 Hoshida, Y. et al. Integrative transcriptome analysis reveals common molecular subclasses of human hepatocellular carcinoma. Cancer Res 69, 7385–7392 (2009).

7 Lee, J. S. et al. A novel prognostic subtype of human hepatocellular carcinoma derived from hepatic progenitor cells. Nat.Med. 12, 410–416 (2006).

8 Yamashita, T. et al. EpCAM-positive hepatocellular carcinoma cells are tumor-initiating cells with stem/progenitor cell features. Gastroenterology 136, 1012–1024 (2009).

9 Nakamura, H. et al. Genomic spectra of biliary tract cancer. Nat Genet 47, 1003–1010, doi:10.1038/ng.3375 (2015).

10 Totoki, Y. et al. Trans-ancestry mutational landscape of hepatocellular carcinoma genomes. Nat Genet 46, 1267–1273, doi:10.1038/ng.3126 (2014).

11 Guichard, C. et al. Integrated analysis of somatic mutations and focal copy-number changes identifies key genes and pathways in hepatocellular carcinoma. Nat.Genet. 44, 694–698 (2012).

12 Zheng, H. et al. Single cell analysis reveals cancer stem cell heterogeneity in hepatocellular carcinoma. Hepatology, doi:10.1002/hep.29778 (2018).

13 Ling, S. et al. Extremely high genetic diversity in a single tumor points to prevalence of non-Darwinian cell evolution. Proc Natl Acad Sci U S A 112, E6496–6505, doi:10.1073/pnas.1519556112 (2015).

14 Beroukhim, R. et al. The landscape of somatic copy-number alteration across human cancers. Nature 463, 899–905 (2010).

15 Burrell, R. A., McGranahan, N., Bartek, J. & Swanton, C. The causes and consequences of genetic heterogeneity in cancer evolution. Nature 501, 338–345, doi:10.1038/nature12625 (2013).

16 Wang, Y. et al. Clonal evolution in breast cancer revealed by single nucleus genome sequencing. Nature 512, 155–160, doi:10.1038/nature13600 (2014).

17 Yi, S. et al. Functional variomics and network perturbation: connecting genotype to phenotype in cancer. Nat Rev Genet 18, 395–410, doi:10.1038/nrg.2017.8 (2017).

18 Carter, S. L., Eklund, A. C., Kohane, I. S., Harris, L. N. & Szallasi, Z. A signature of chromosomal instability inferred from gene expression profiles predicts clinical outcome in multiple human cancers. Nat Genet 38, 1043–1048, doi:10.1038/ng1861 (2006).

19 Pages, F. et al. Effector memory T cells, early metastasis, and survival in colorectal cancer. N Engl J Med 353, 2654–2666, doi:10.1056/NEJMoa051424 (2005).

20 Sato, E. et al. Intraepithelial CD8+ tumor-infiltrating lymphocytes and a high CD8+/regulatory T cell ratio are associated with favorable prognosis in ovarian cancer. Proc Natl Acad Sci U S A 102, 18538–18543, doi:10.1073/pnas.0509182102 (2005).

21 Rooney, M. S., Shukla, S. A., Wu, C. J., Getz, G. & Hacohen, N. Molecular and genetic properties of tumors associated with local immune cytolytic activity. Cell 160, 48–61, doi:10.1016/j.cell.2014.12.033 (2015).

22 Lapunzina, P. & Monk, D. The consequences of uniparental disomy and copy number neutral loss-of-heterozygosity during human development and cancer. Biol Cell 103, 303– 317, doi:10.1042/BC20110013 (2011).

23 Oren, M. & Rotter, V. Mutant p53 gain-of-function in cancer. Cold Spring Harb.Perspect.Biol. 2, a001107 (2010).

24 Gualberto, A., Aldape, K., Kozakiewicz, K. & Tlsty, T. D. An oncogenic form of p53 confers a dominant, gain-of-function phenotype that disrupts spindle checkpoint control. Proc Natl Acad Sci U S A 95, 5166–5171 (1998).

25 Restle, A. et al. Dissecting the role of p53 phosphorylation in homologous recombination provides new clues for gain-of-function mutants. Nucleic Acids Res 36, 5362–5375, doi:10.1093/nar/gkn503 (2008).

26 Olivier, M., Hollstein, M. & Hainaut, P. TP53 mutations in human cancers: origins, consequences, and clinical use. Cold Spring Harb Perspect Biol 2, a001008, doi:10.1101/cshperspect.a001008 (2010).

27 Newman, A. M. et al. Robust enumeration of cell subsets from tissue expression profiles. Nat Methods 12, 453–457, doi:10.1038/nmeth.3337 (2015).

28 Chen, L. & Flies, D. B. Molecular mechanisms of T cell co-stimulation and co-inhibition. Nat Rev Immunol 13, 227–242, doi:10.1038/nri3405 (2013).

29 Zhang, Q. F. et al. Liver-infiltrating CD11b(-)CD27(-) NK subsets account for NK-cell dysfunction in patients with hepatocellular carcinoma and are associated with tumor progression. Cell Mol Immunol 14, 819–829, doi:10.1038/cmi.2016.28 (2017).

30 Zekri, A. N. et al. Role of relevant immune-modulators and cytokines in hepatocellular carcinoma and premalignant hepatic lesions. World J Gastroenterol 24, 1228–1238, doi:10.3748/wjg.v24.i11.1228 (2018).

31 Circelli, L. et al. Immunological effects of a novel RNA-based adjuvant in liver cancer patients. Cancer Immunol Immunother 66, 103–112, doi:10.1007/s00262-016-1923-5 (2017).

32 Bu, Y., Liu, F., Jia, Q. A. & Yu, S. N. Decreased Expression of TMEM173 Predicts Poor Prognosis in Patients with Hepatocellular Carcinoma. PLoS One 11, e0165681, doi:10.1371/journal.pone.0165681 (2016).

33 Ahmad, S. M. et al. The inhibitory checkpoint, PD-L2, is a target for effector T cells: Novel possibilities for immune therapy. Oncoimmunology 7, e1390641, doi:10.1080/2162402X.2017.1390641 (2018).

34 Cancer Genome Atlas, N. Genomic Classification of Cutaneous Melanoma. Cell 161, 1681–1696, doi:10.1016/j.cell.2015.05.044 (2015).

35 Hugo, W. et al. Genomic and Transcriptomic Features of Response to Anti-PD-1 Therapy in Metastatic Melanoma. Cell 168, 542, doi:10.1016/j.cell.2017.01.010 (2017).

36 Berasain, C. et al. Inflammation and liver cancer: new molecular links. Ann N Y Acad Sci 1155, 206–221, doi:10.1111/j.1749-6632.2009.03704.x (2009).

37 Coussens, L. M. & Werb, Z. Inflammation and cancer. Nature 420, 860–867, doi:10.1038/nature01322 (2002).

38 Hussain, S. P., Hofseth, L. J. & Harris, C. C. Radical causes of cancer. Nat Rev Cancer 3, 276–285 (2003).

39 Senovilla, L., Galluzzi, L., Castedo, M. & Kroemer, G. Immunological control of cell cycle aberrations for the avoidance of oncogenesis: the case of tetraploidy. Ann N Y Acad Sci 1284, 57–61, doi:10.1111/nyas.12072 (2013).

40 Wherry, E. J. & Kurachi, M. Molecular and cellular insights into T cell exhaustion. Nat Rev Immunol 15, 486–499, doi:10.1038/nri3862 (2015).

41 Topalian, S. L., Weiner, G. J. & Pardoll, D. M. Cancer immunotherapy comes of age. J Clin Oncol 29, 4828–4836, doi:10.1200/JCO.2011.38.0899 (2011).

42 Mathai, A. M. et al. Role of Foxp3-positive tumor-infiltrating lymphocytes in the histologic features and clinical outcomes of hepatocellular carcinoma. Am J Surg Pathol 36, 980–986, doi:10.1097/PAS.0b013e31824e9b7c (2012).

43 Davoli, T., Uno, H., Wooten, E. C. & Elledge, S. J. Tumor aneuploidy correlates with markers of immune evasion and with reduced response to immunotherapy. Science 355, doi:10.1126/science.aaf8399 (2017).

44 Sharma, P. & Allison, J. P. The future of immune checkpoint therapy. Science 348, 56–61, doi:10.1126/science.aaa8172 (2015).

45 Greten, T. F. & Sangro, B. Targets for immunotherapy of liver cancer. J Hepatol, doi:10.1016/j.jhep.2017.09.007 (2017).

46 Roessler, S. et al. A unique metastasis gene signature enables prediction of tumor relapse in early-stage hepatocellular carcinoma patients. Cancer Res 70, 10202–10212, doi:10.1158/0008-5472.CAN-10-2607 (2010).

