## Supplementary Figures for "Functional Genomic Complexity Defines Intratumor Heterogeneity and Tumor Aggressiveness in Liver Cancer"

**This PDF file includes:**

Supplementary Materials and Methods
Figure S1 to S14
Figure S1. Distribution of global correlation coefficient in PLC.
Figure S2. Gene Ontology (GO) of PCC associated genes.
Figure S3. Association of PCC with CIN and GIN.

Figure S4. Association of amplified or deleted CIN (CINampl or CINdel) with PCC.
Figure S5. Gene Ontology (GO) of tFA associated genes.
Figure S6. Collective association among PCC, CIN, and tFA.
Figure S7. Validation of FGC in independent cohorts.
Figure S8. Comparison of tFA between HFGC and LFGC.
Figure S9. Comparison of SCNA in HFGC and LFGC.
Figure S10. Differentially expressed genes (DEG) between HFGC and LFGC of TCGA.
Figure S11. Gene Set Enrichment Analysis of HFGC and LFGC of Thai PLC.
Figure S12. Integrative analysis based on PCC showed TP53 as a Cancer functional genomic complexity (FGCs) driver.
Figure S13. Comparison of TIL subpopulations between HFGC and LFGC.
Figure S14. Association of FGC with immunomodulators

References

### Supplementary Materials and Methods

Data preprocessing

Transcriptome data and copy number data from TIGER-LC cohort was processed as follows. For Affymetrix Human Transcriptome Array 2.0 data, expression level of individual 914,585 exons were extracted and normalized based on the Robust Multi-array Average (RMA) method and sketch-quantile normalization method using the Transcriptome Analysis Console (TAC) Software 4.0. For transcripts with more than one exon probe sets, the mean expression was calculated and total 64,597 transcripts were used further analysis. For profiling of copy number for tumors and paired non-tumor tissues generated based on Affymetrix Genome-Wide Human SNP Nsp/Sty 6.0, we applied the crlmm R package into the raw CEL files to estimate copy number based on the CRLMM algorithm (1). Briefly, the ***crlmm*** package adapts the robust multichip average (RMA) to genotyping platforms based on the SNP-RMA algorithm (2). For probes for polymorphic loci, the raw intensities for each allele are quantile normalized (3) to a target reference distribution obtained from the HapMap phase 2 samples. The Affymetrix 6.0 platform contains 3 or 4 identical probes for each allele. The normalized intensities for a set of identical probes are summarized by the median. For nonpolymorphic loci, only one probe per loci is available and the intensities are quantile normalized without a subsequent summarization step. Additional details regarding the preprocessing of Affymetrix CEL files are described elsewhere(2). Somatic copy number variations were inferred by CBS (Circular binary segmentation) algorithm (4). The genomic locations of segmented regions were converted from hg19 to hg38 by applying the UCSC *liftOver* R package. Copy number value of segmented regions was merged or separated for corresponding transcriptome probes, resulting in allocation of copy number value for each segment corresponding 64,597 transcripts. For validation cohort, HCC cohort of 247 Chinese patients from LCI (5) and TCGA LIHC cohort with 377 HCC patients were used. Transcriptome data and aCGH data for LCI cohort was processed as described previously (6). Copy number value for each segmented region was allocated into each corresponding gene probe of the transcriptome data located in the segmented region resulting in 10,127 features. The level 3 RNA-seq v2.0 data and Affymetrix SNP 6.0 data were downloaded from TCGA Research Network (http://cancergenome.nih.gov/ ; release 1.0). Gene-level annotated transcriptome data segmented data were used for further analysis. All processing was conducted using R packages of Bioconductor 3.5 (https://cran.r-project.org/doc/FAQ/R-FAQ.html).

Calculation of global correlation

The global correlation coefficients and global correlation p-value based on the total transcriptome probes and corresponding genomic segments were calculated. For this, SCNA value for genomic segments corresponding to the 64,597 transcript probes was assigned using the GenomicRanges R packages. Hereafter, CN denotes copy number value and EXP denotes mRNA expression value.

$$\begin{matrix} \text{CN} & & \text{EXP} \\ \begin{pmatrix} c_{11} & \cdots & c_{1m} \\ \vdots & \ddots & \vdots \\ c_{n1} & \cdots & c_{nm} \end{pmatrix} & \text{x} & \begin{pmatrix} e_{11} & \cdots & e_{1m} \\ \vdots & \ddots & \vdots \\ e_{n1} & \cdots & e_{nm} \end{pmatrix} \end{matrix}$$

Where $c_{nm}$ represent the SCNA value of $n^{th}$ sample corresponding $m^{th}$ feature. Matched features are expressed like below.

$$\begin{matrix} & \mathbf{F_1} & & \mathbf{F_K} & & \mathbf{F_M} \\ \mathbf{S_1} & c_{11}e_{11} & \cdots & c_{1k}e_{1k} & \cdots & c_{1m}e_{1m} \\ & \vdots & & \vdots & & \vdots \\ \mathbf{S_i} & c_{i1}e_{i1} & & c_{ik}e_{ik} & & c_{im}e_{im} \\ & \vdots & & \vdots & & \vdots \\ \mathbf{S_n} & c_{n1}e_{n1} & \cdots & c_{nk}e_{nk} & \cdots & c_{nm}e_{nm} \end{matrix}$$

Where $F_K$ indicates $K^{th}$ feature and $S_i$ indicates for $i^{th}$ sample. Global correlation of M number of features from n number of tumor (T) and non-tumor (NT) of HCC and iCCA sample was calculated. Permutated correlation coefficient and p-value was used to compare between T and NT. Significantly correlated features of transcriptome probes and corresponding segmented regions were selected ($p$-value < 0.05 & median absolute deviation (MAD) >20% of overall distribution) for further analysis.

<u>Calculation of SCNA frequency and Inference of Arm-level SCNA</u>

To define amplified or deleted region, we applied threshold, 0.2 or -0.2, respectively, to the log2 transformed copy number value for individual 64,597 features. The fraction of patients who showed amplification or deletion was calculated for each feature. We calculated the frequency of arm-level amplifications and deletions based on the GISTIC (Genomic Identification of Significant Targets in Cancer) algorithm(7) in GISTIC_2.0 module of GenePattern (8). The segments with log2 ratio > 0.2 and < -0.2 were defined as chromosomal amplifications and deletions following the default value of the algorithm, respectively.

<u>LOH and allelic specific copy number</u>

LOH (Loss of Heterozygosity) for each sample was inferred using Genotyping Console 4.0 and output CHP file was used as an input file to calculate allele specific copy number using the Partek Genomics Suite 7.5. By merging copy number of segmented region defined by algorithm in Partek and LOH data, the allele specific copy number was estimated. For further analysis, we calculated the proportion of sample with allele specific copy number change in each segmented region.

<u>Biological relevance of PCC or tFA associated genes</u>

To examine if PCC or tFA was associated with the biological process, we performed correlation analysis between PCC and all the transcriptome features. Positively or negatively associated genes were selected based on the correlation estimate and p-value (above top 5% or below bottom 5% of correlation estimate and p-value < 0.05). Gene ontology enrichment analysis was performed using R package gProfileR based on GO: BP.

Gene Set Enrichment Analysis (GSEA) and Single-sample GSEA (ssGSEA)

GSEA was implemented in GenePattern (8) based on the C5 GO gene set of biological process, C2 curated gene sets of KEGG pathway, and C6 Oncogenic gene sets in Molecular Signatures Database (MSigDB database v5.2). Expression data of individual sample was transformed into gene set enrichment score P-value from the Kolmogorov-Smirnov (ks) test was used single sample enrichment score.

Measurement of chromosomal instability

To infer chromosomal instability, we devised two indicators, one is based on the SCNA proportion of individual sample level and the other is based on summation of length of segments with SCNA. For comparison of chromosomal instability of individual sample level, we defined the segments with log2 ratio > 0.2 and < -0.2 as chromosomal amplifications and deletions by applying noise cutoff of 0.2, respectively and proportion of amplified ($CIN_{ampl}$) or deleted features ($CIN_{del}$) over total features was calculated. $CIN_{ampl}$ and $CIN_{del}$ for individual patient was calculated based on the copy number value for 64,597 features. Summation of $CIN_{ampl}$ and $CIN_{del}$ was used as CIN score for further analysis. As another aspect of chromosome instability indicator, we also calculated the average length for total amplified ($GIN_{gain}$) or deleted regions ($GIN_{loss}$) and used the summation of the $GIN_{gain}$ and $GIN_{loss}$ for total SCNA length as a genomic instability (GIN) score.

Total functional aneuploidy (tFA)

We calculated total functional aneuploidy (tFA) in each sample based on coordinated aberrations in expression of genes localized to each chromosomal region using the adapted computational method from previously published paper (9). Briefly, it is a computational method to characterize aneuploidy in tumor samples based on coordinated aberrations in expression of genes localized to each chromosomal region. For a given data set, all of the normalized gene expression measurements present on the microarray and mapping to a given chromosomal cytoband region were grouped into a set designated 'B' (short for band). The rest of the genes, localized elsewhere on the genome, were grouped into a set 'G' (short for genome). The functional aneuploidy measure for the given cytoband is the value of student's t statistic comparing sets B and G. Sum of all functional aneuploidy magnitudes (the absolute t statistics) in a given tumor sample. Therefore, the tFA is total summarized level of chromosomal aberration in a given tumor in a univariate measure.

Differentially Expressed Genes (DEG) and Gene Ontology analysis

By comparing the expression between HFGC and LFGC of each tumor type, we selected differentially expressed genes in each subtype based on the fold change and permutation p-value from permutation t-test with 1,000 resampling (FC >0.5 or -0.5 & perm p-value <0.005). Gene ontology enrichment analysis was performed based on the DAVID 6.7(10).

Immune score

An estimation of the relative fractions of immune/inflammatory cell subsets from tissue expression profiles of Thai HCC, iCCA or TCGA HCC was conducted using CIBERSORT (11). The gene expression data was converted by quantile normalization of the log2 scaled expression matrix and relative fractions of leukocytes were quantified according to the website (https://cibersort.stanford.edu/index.php) with implemented analyses using the built-in LM22 signature matrix (LM22). Immune score of individual tumor or non-tumor tissue was calculated as summation of the 22 types of tumor infiltrating lymphocytes (TILs) fraction based on CIBERSORT output. Since the output value was ranged from 0 to 1, for calculation convenience, we transformed the output value by multiplying by 100 and added one before log2 transformation. The summation of transformed value for each TILs was used as the estimate of immune score. Considering the difference of clinical outcome between LFGC and HFGC, TILs enriched in LFGC than HFGC were defined as favorable or adverse, vice versa. The summation of adverse or favorable TILs fractions were used as immune score of adverse or favorable TILs.

Mutation Map

MutationMapper (version 1.0) in the cBioPortal (http://www.cbioportal.org/tools.jsp) was used to plot lollipop mutation diagram view with genomic coordinates to annotate TP53 variants (12, 13).

Validation with melanoma dataset

We used transcriptome data from skin cutaneous melanoma datasets derived from TCGA_SKCM(14) study (n=472) and metastatic melanoma from Hugo(15) study (n=28) to validate the association between FGC and immunotherapy with immune checkpoint blockade (ICB). We calculated tFAs in the individual sample and used them as a surrogate of PCC on the assumption that tFA were strongly associated with PCC based on our findings on liver cancer. Among TCGA_SKCM, 13 samples, which were pre-treated with anti-CTLA-4 therapy, were included or excluded to perform KM survival analysis to examine whether tFA level predict responsiveness to ICB treatment. To compare high and low group, patients were stratified based on the tFA level into high (above 3$^{rd}$ quartile) and low group (below 1$^{st}$ quartile) in the TCGA_SKCM. Among the patients with anti-CTLA-4 pretreatment, tFA levels were compared between responders and non-responders based on the Welch's two sample t-test. As another independent cohort, metastatic melanoma samples from Hugo study(15), where 28 patients were pre-treated with anti-PD-1 therapy, were used to validate the association between tFA and ICB responsiveness. To perform KM survival analysis, we divided patients into high and low group based on the median level of tFA. We classified patients into "Responder" and "Non-Responder" as followed; "Responder" indicates those who marked as "Complete

Response" or "Partial Response", while "Non-Responder" indicates those who marked as "Clinical Progressive Disease" or "Stable Disease" according to the response column of the clinical data.

Statistical Analyses

Kaplan-Meier (KM) Survival Analysis was performed based on the survival R package and p-value from log-rank test based on the Cox Proportional-Hazards Regression model was used to compare overall survival. Permutation t-test was calculated based on the perm R package by 1,000 resampling. Correlation coefficient and p-value was calculated based on the Pearson's product moment correlation. After filtering based on the global correlation p-value (p-value<0.05) and MAD of copy number value (MAD > the value of 20% of MAD percentile), correlation coefficient was calculated in the individual subject using the corresponding correlated segment and transcriptome sets. All statistical tests were performed using R.

**Supplementary Figures**

Figure S1

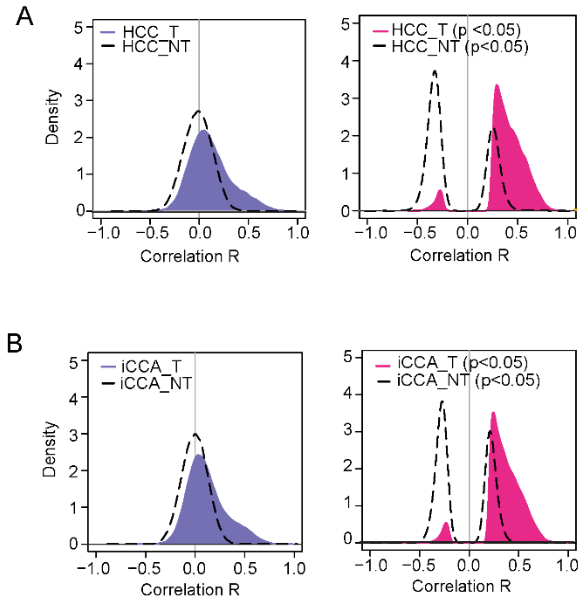


**Figure S1. Distribution of global correlation coefficient in PLC** (A-B) The density histogram shows the distribution of global correlation coefficient based on the permutated Pearson's correlation of DNA copy number (CN) and mRNA expression (EXP) from Tumor tissues and corresponding non-tumor tissue (A: HCC; n=64, HCC_NT; n=59, B: iCCA ; n=90, iCCA_NT; n=90). The distribution of correlation R is shown before (left panel) and after applying cut-off based on p-value (p-value <0.05) (right panel).

Figure S2

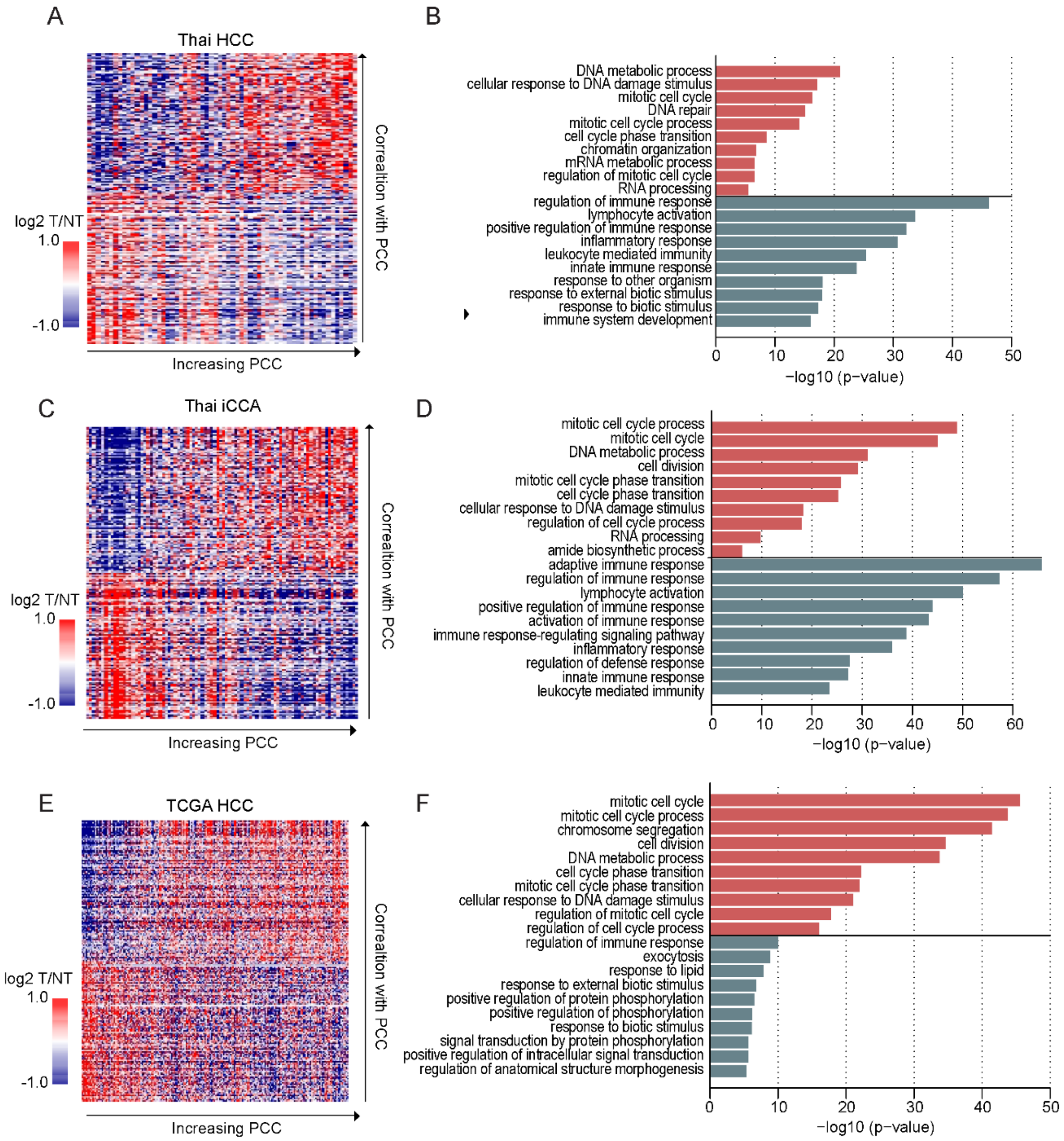


**Figure S2. Gene Ontology (GO) of PCC associated genes** (A, C, and E) Positively or negatively PCC associated genes were selected based on the correlation coefficient and p-value (more than 95% or less than 5% of estimate and p-value <0.01). Heatmap shows the expression level of selected genes in Thai HCC, iCCA, and TCGA HCC cohorts (A, C, and E, respectively). Samples are represented in columns according to the PCC increasing order. Selected genes were represented in the row according to the decreasing of correlation coefficient with PCC. (B, D, and F) GO Enrichment Analysis of selected genes in Thai HCC, iCCA, and TCGA HCC cohorts was performed (B, D, and F, respectively). Top10 ranked process based on the precision rank were shown. Orange and green color indicate positively and negatively correlated gene sets, respectively.

Figure S3

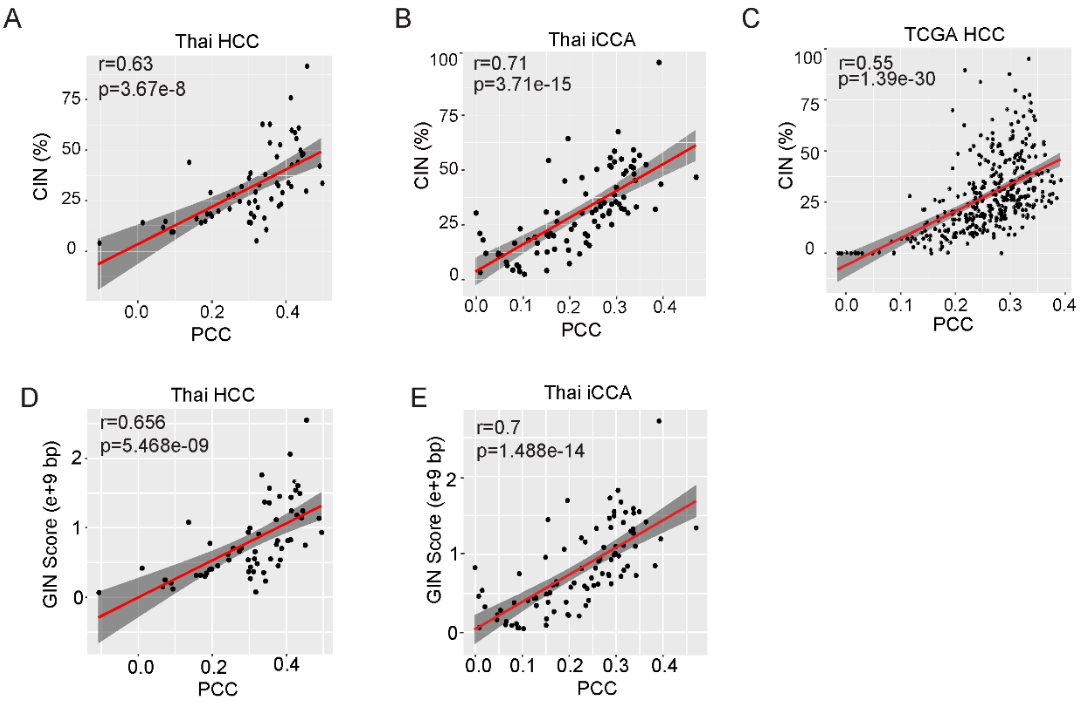


**Figure S3. Association of PCC with CIN and GIN** (A-C) PCC shows strong association with CIN in Thai HCC, Thai iCCA and TCGA HCC, respectively. (D-E) Genomic instability (GIN) length regarding to the copy number gain or copy number loss (Methods) was calculated in the individual sample and the summation of the total SCNA length was calculated as GIN score. PCC shows strong association with GIN score of individual Thai HCC (D) and iCCA (E) samples

Figure S4

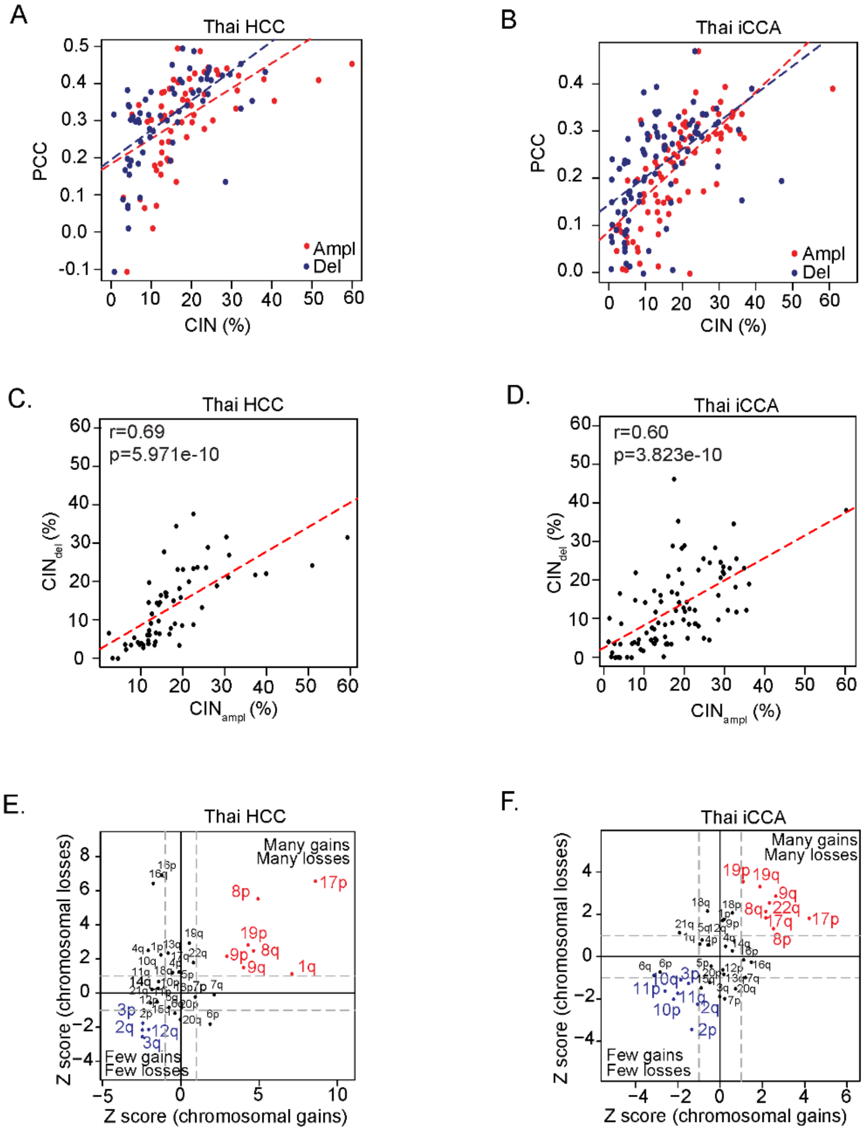


**Figure S4. Association of amplified or deleted CIN ($CIN_{ampl}$ or $CIN_{del}$) with PCC** (A-B) Strong associations of $CIN_{ampl}$ or $CIN_{del}$ with PCC in Thai HCC (A) and Thai iCCA (B) are shown. Red or blue dots indicate $CIN_{ampl}$ or $CIN_{del}$, respectively. Red or blue dots indicate $CIN_{ampl}$ or $CIN_{del}$, respectively. (C-D) Strong linear association between $CIN_{ampl}$ and $CIN_{del}$ were shown in Thai HCC (C) and Thai iCCA (D). Coefficient estimates and p-value based on Pearson's correlation were depicted. (E-F) Frequency of recurrent arm-level SCNA of Thai HCC (E) and Thai iCCA (F) are shown. Chromosomal arms are shown with respect to the frequency of arm-level gain (x axis) and loss (y axis), respectively. As a frequency measure, Z score from GISTIC output was used. Vertical dotted blue lines indicate Z score of the arm-level gain frequency is 1 and horizontal dotted blue lines indicate Z score of the arm-level loss frequency is 1. The arms with many gains and many losses or with few gains or few losses were high-lighted in red or blue colors, respectively.

Figure S5

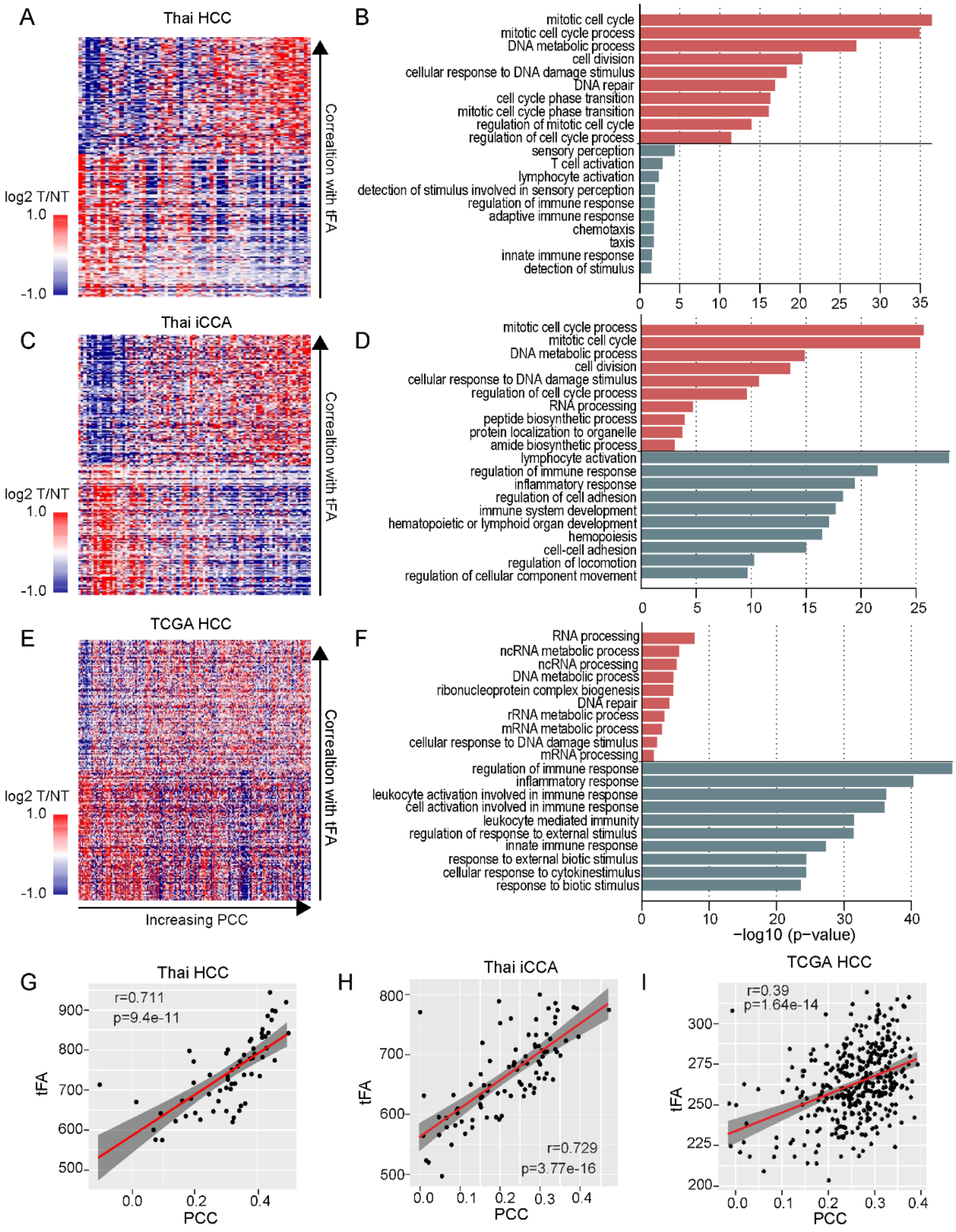


**Figure S5. Gene Ontology (GO) of tFA associated genes** (A-F) Functional relevance of PCC with tFA were examined among Thai HCC, Thai iCCA and TCGA HCC, respectively. Positively or negatively tFA associated genes were selected based on the correlation coefficient and p-value (more than 95% or less than 5% of estimate and p-value <0.01). Heatmap shows the expression level of selected genes in Thai HCC, iCCA, and TCGA HCC cohorts. Samples were represented in columns according to the FGC increasing order and selected genes were represented in the row according to the decreasing of correlation coefficient with tFA (A, C, and D, respectively). GO Enrichment Analysis of selected genes in Thai HCC, iCCA, and TCGA HCC cohorts was performed (B, D, and F, respectively). Top10 ranked process based on the precision rank were shown. Orange and green color indicate positively and negatively correlated gene sets, respectively. (G-I) PCC shows strong association with tFA in Thai HCC, Thai iCCA and TCGA HCC, respectively. Coefficient estimates and p-value based on Pearson's correlation were depicted.

Figure S6

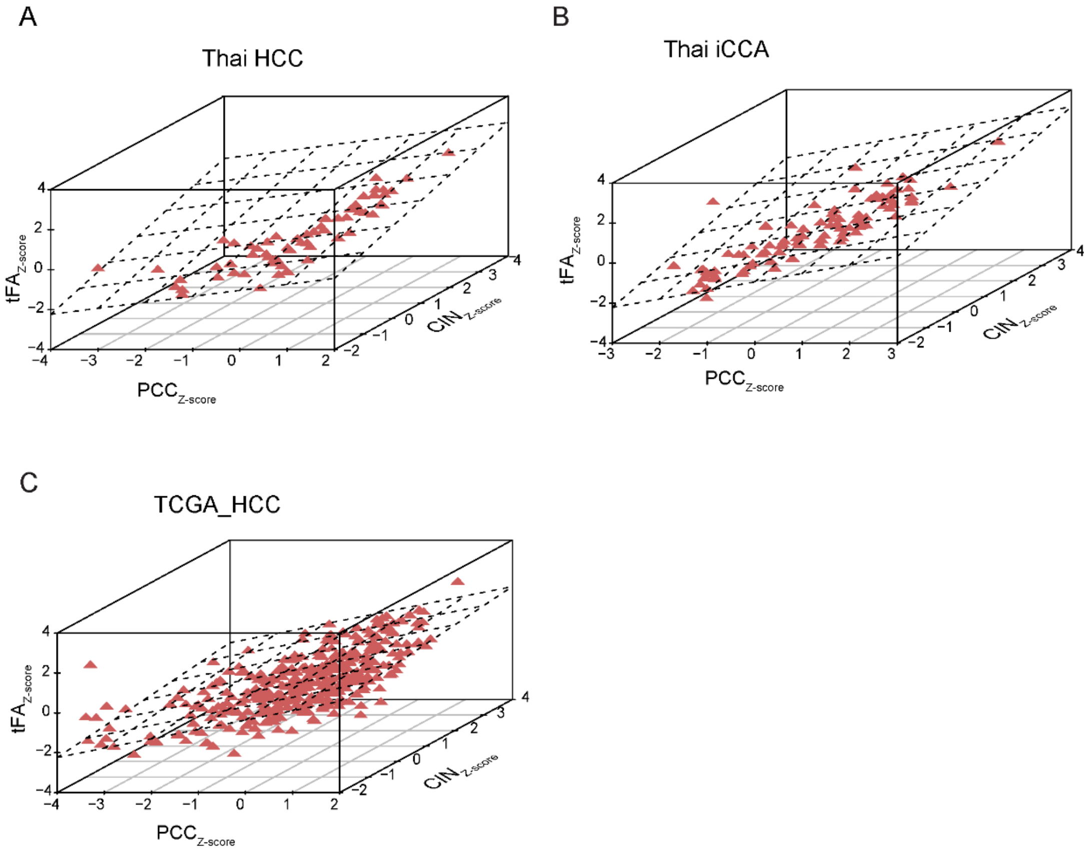


**Figure S6. Collective association among PCC, CIN, and tFA** (A-C) Collective association among the PCC (x axis), CIN (y axis), and tFA (z axis) are shown in Thai HCC, iCCA, and TCGA HCC, respectively.

Figure S7

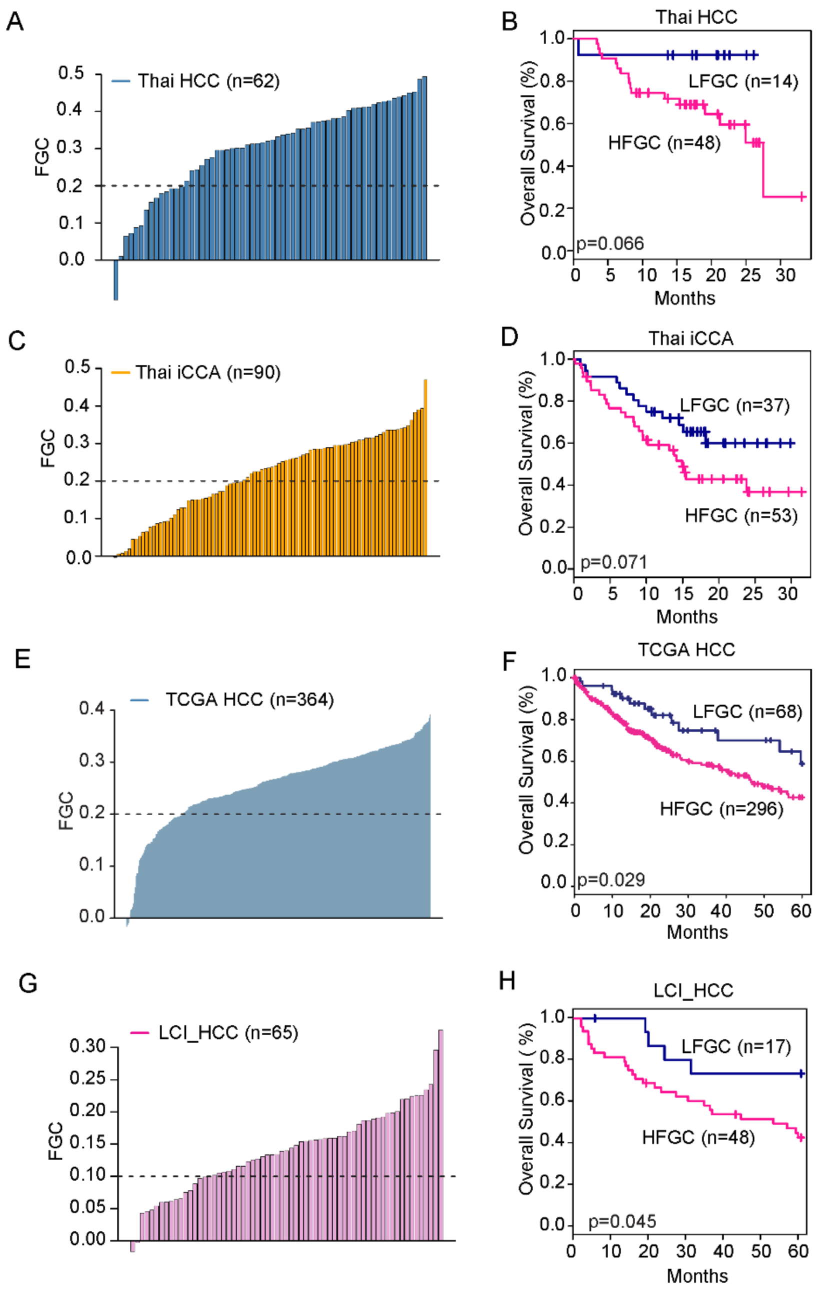


**Figure S7. Validation of FGC in independent cohorts** (A, C, E and G) (A) FGC values among the Thai HCC, Thai iCCA, TCGA HCC, and LCI HCC are plotted in a rank order, respectively. Dotted line indicates the cut-off FGC value, 0.2, applied to separate into FGC high (HFGC) and FGC low (LFGC) group in each tumor type, except for LCI HCC. (B, D, F and H) Kaplan-Meier survival analysis performed based on LFGC and HFGC among the Thai HCC, Thai iCCA, TCGA HCC, and LCI HCC shows significant difference on the overall survival, respectively. The statistical P value was generated by the Cox-Mantel log-rank test.

Figure S8

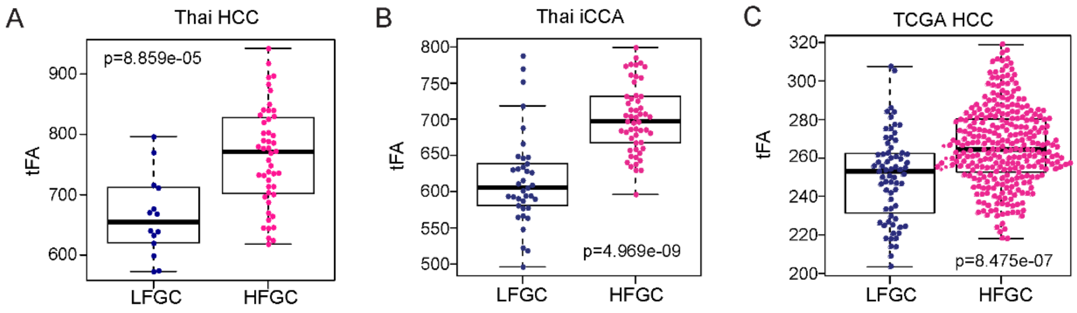


**Figure S8. Comparison of tFA between HFGC and LFGC** (A-C) HFGC show higher value of tFA in Thai HCC, Thai iCCA and TCGA HCC, respectively. P-value based on Welch two sample t-test was depicted.

Figure S9

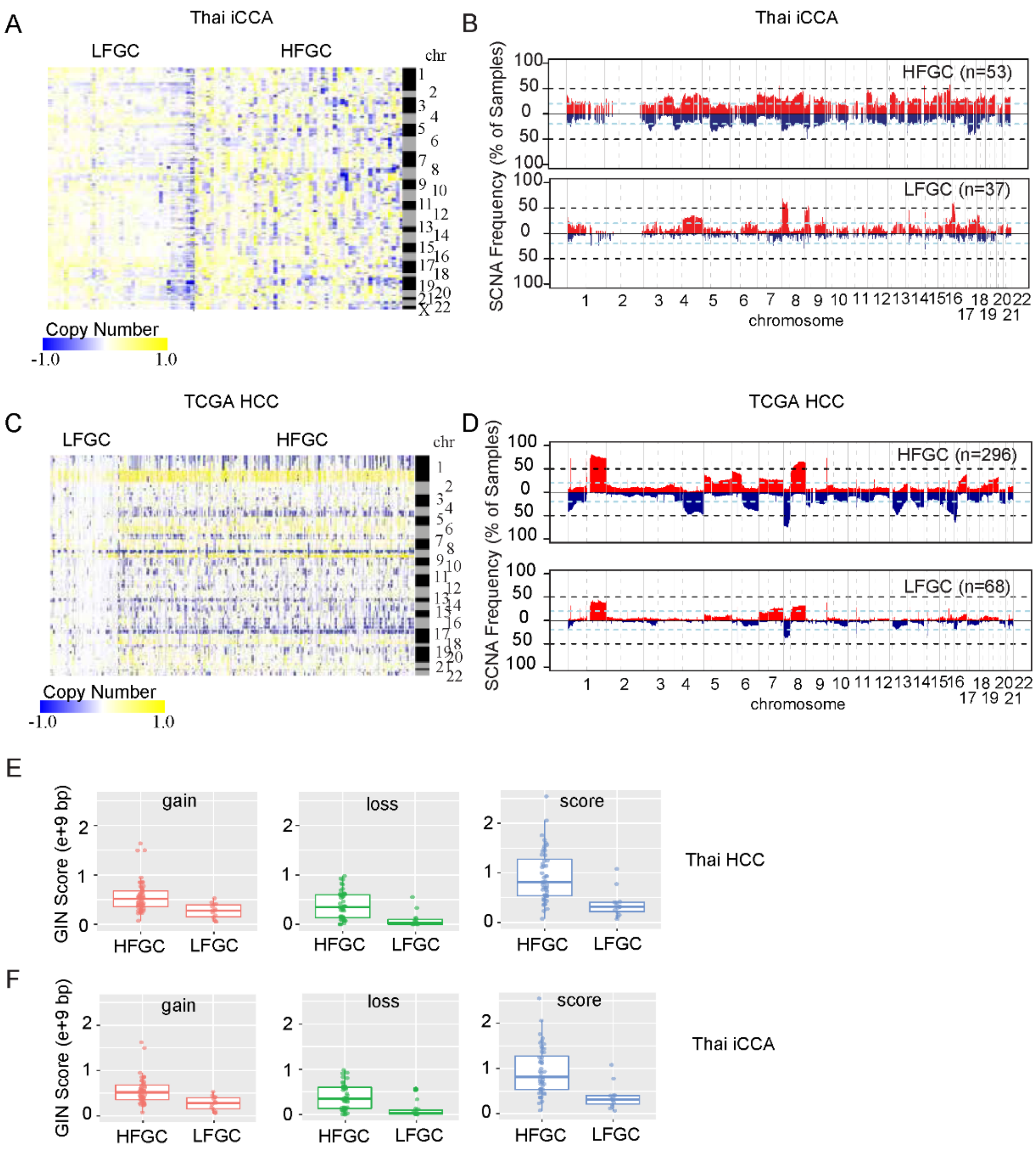


**Figure S9. Comparison of SCNA in HFGC and LFGC** (A and C) Heatmap shows copy number value of individual sample of Thai iCCA (A) and TCGA HCC (C) corresponding to the correlated segments regions, respectively. Samples are represented in columns, grouped by the HFGC and LFGC and segment regions are represented in rows according to the chromosomal location. (B and D) The frequency of SCNA among HFGC and LFGC subtype of Thai iCCA (B) and TCGA HCC (D) are plotted corresponding to the correlated segments region, respectively. The sample frequencies with copy number gain and loss (log2 (copy number) >0.2 or log2 (copy number) < -0.2) are shown in red and blue, respectively. Upper panel is the SCNA frequency plot for HFGC subtype and lower panel is the SCNA frequency plot for LFGC subtype. Chromosome boundaries and centromere position are indicated by vertical solid and dashed lines, respectively. Horizontal dashed blue lines indicate frequency of 50%. Horizontal dotted black lines indicate frequency of 20%. (E-F) Genomic instability (GIN) scores were compared between HFGC and LFGC. Boxplots for GIN length regarding to the gain (top), loss (middle), and score (bottom) for HFGC and LFGC subtype of Thai HCC (E) and Thai iCCA (F) are shown. GIN length regarding to the copy number gain or copy number loss (Methods) was calculated in the individual sample and the summation of the total SCNA length was calculated as score.

Figure S9

G

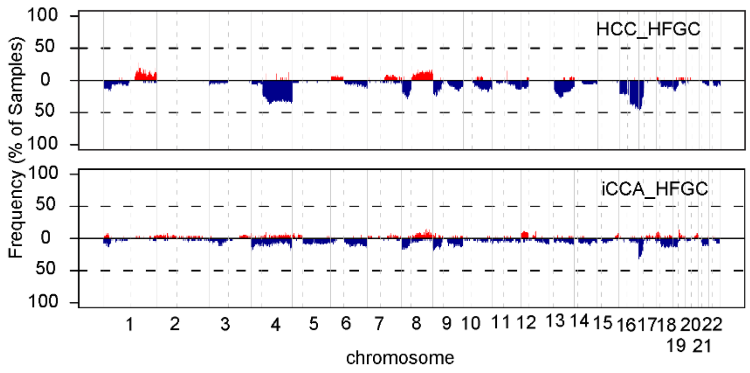


H

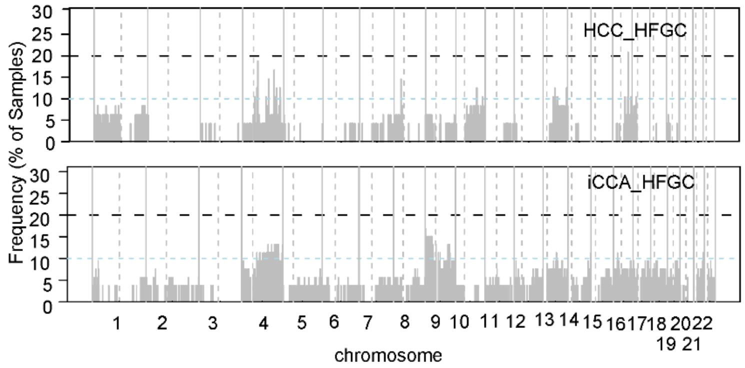


**Figure S9. Comparison of SCNA in HFGC and** (G-H) Allelic imbalance frequency between HFGC and LFGC was compared. (G) Frequencies of samples with amplified (AMP W/ LOH) or deletion region with LOH (DEL W/ LOH) among HCC_HFGC (upper panel) and iCCA_HFGC (lower panel) are plotted according to the chromosome location. AMP W/ LOH or DEL W/ LOH are shown in red or blue, respectively. Chromosome boundaries and centromere positions are indicated by vertical solid and dashed liens, respectively. Horizontal dashed blue lines indicate frequency of 20%. (H) Frequencies of samples with segment region with CN LOH among HCC_HFGC (upper panel) and iCCA_HFGC (lower panel) are plotted according to the chromosome location. Chromosome boundaries and centromere position are indicated by vertical solid and dashed liens, respectively. Horizontal dashed blue lines indicate frequency of 10%.

Figure S10

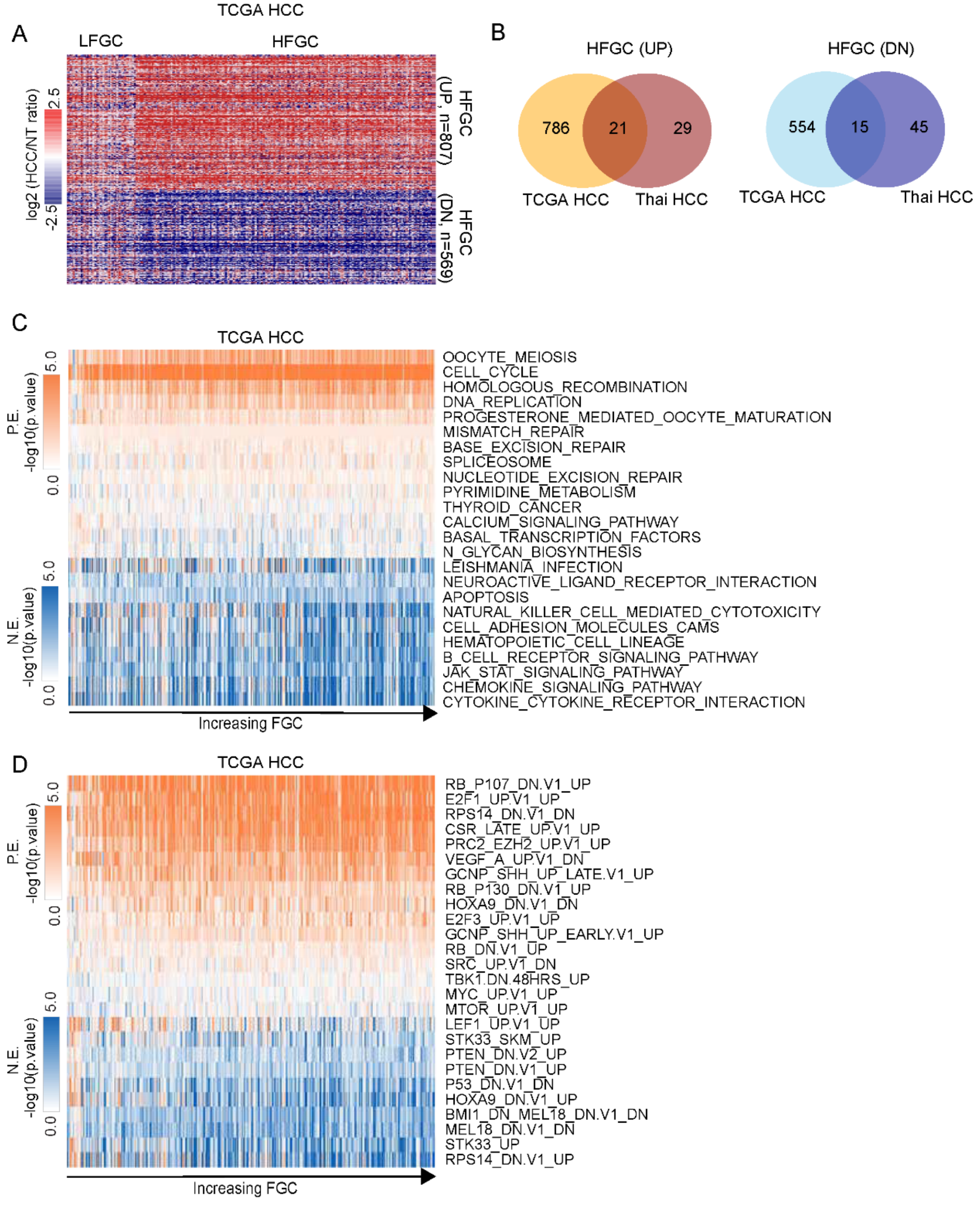


**Figure S10. Differentially expressed genes (DEG) between HFGC and LFGC of TCGA** (A) Heatmap shows the expression of DEG between HFGC and LFGC of TCGA HCC. 807 Up-regulated genes and 569 Down-regulated genes were selected based on the permutation t-test (p-value < 0.005 and log2 fold change >0.5 or <-0.5, respectively). Each gene expression value was normalized based on the mean of non-tumor tissue. Samples are represented in columns, grouped by the HFGC and LFGC and genes are represented in rows. (B) Venn diagrams shows the overlapped genes between DEG of HCC and of TCGA HCC. Up- and down-regulated genes are analyzed separately. (C and D) Gene Set Enrichment Analysis was performed with mRNA expression data from TCGA HCC based on the gene sets derived from the KEGG pathway gene sets (C) and oncogenic signature (D) in Molecular Signatures Database (MSigDB database v5.2). P-value from the Kolmogorov-Smirnov (ks) test was transformed in -log scaled and used in the plot. Samples were represented in columns according to the FGC increasing order and log transformed p-value for each gene set was represented in rows in the rank-order. Shown are the gene sets selected based on the significant difference between HFGC and LFGC subtype. P.E. p-value and N.E. p-value denote the p-value for positively and negatively enriched gene sets, respectively.

Figure S11

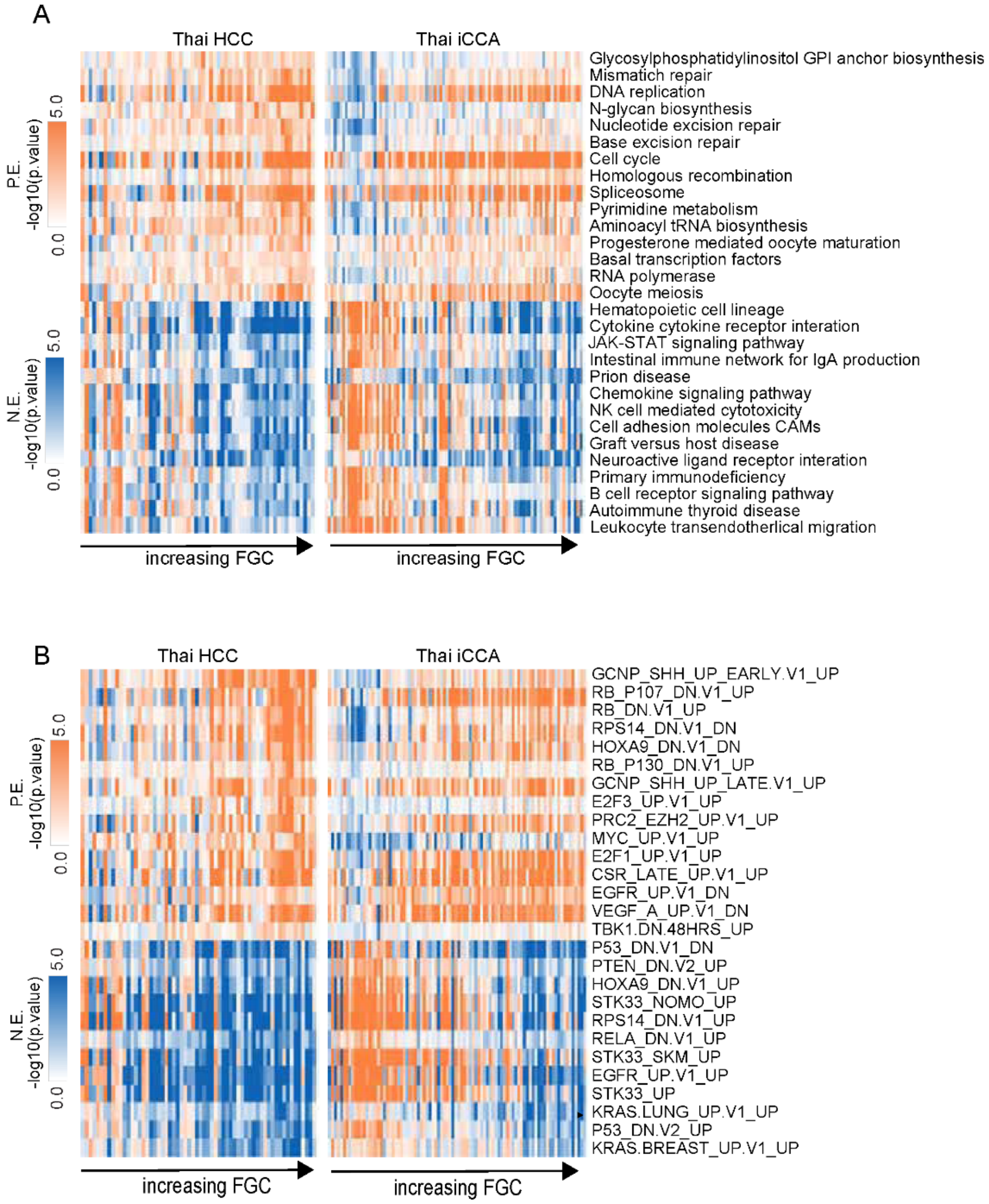


**Figure S11. Gene Set Enrichment Analysis of HFGC and LFGC of Thai PLC** (A and B) Single sample Gene Set Enrichment Analysis (ssGSEA) was performed with mRNA expression data from Thai HCC and Thai iCCA, respectively, based on the gene sets derived from the KEGG pathway gene sets (A) and oncogenic signature (B) (MSigDB database v5.2). P-value from the Kolmogorov-Smirnov (ks) test was transformed in -log scaled and used in the plot. Samples are represented in columns according to the rank order of FGC value and log transformed p-value for each gene set was represented in rows. Shown are the overlapped gene sets significantly enriched both Thai HCC and iCCA. P.E. p-value and N.E. p-value denote the p-value for positively and negatively enriched gene sets, respectively.

Figure S12

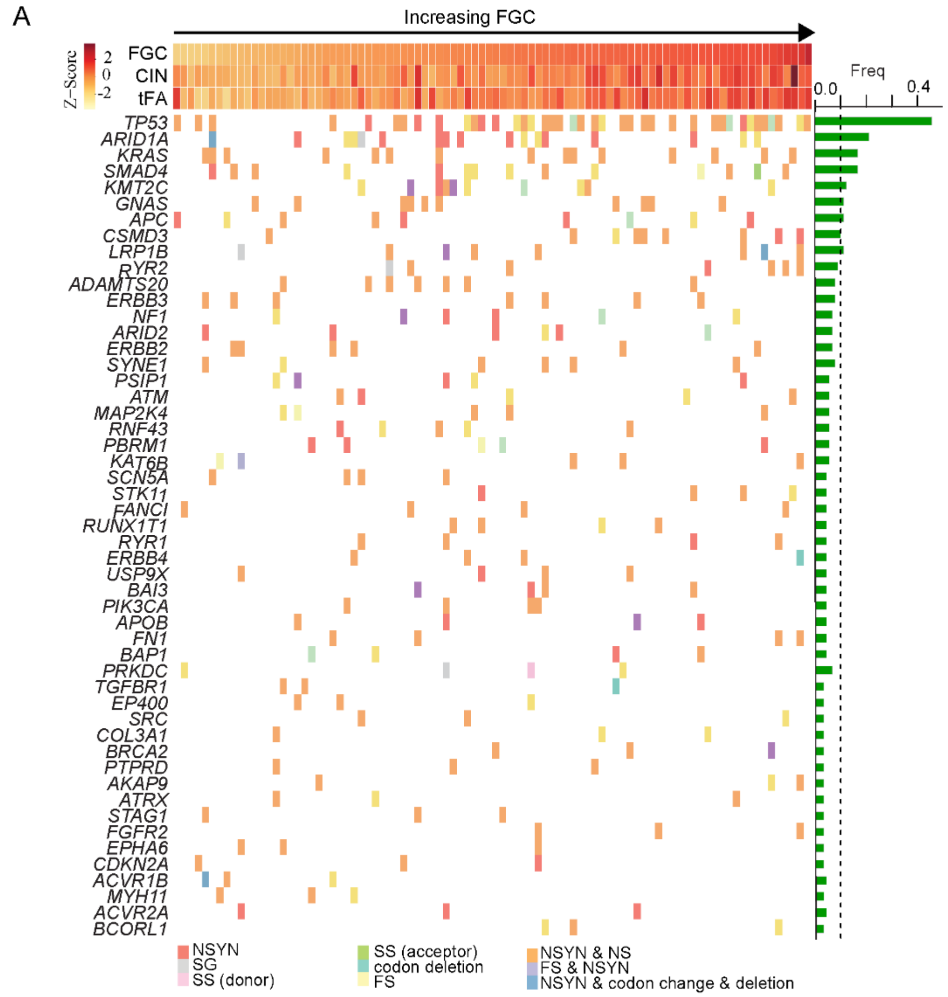


**Figure S12. Integrative analysis based on PCC showed *TP53* as a Cancer functional genomic complexity (FGCs) driver.** (A) (Top panel) Association between CIN, FGC, and tFA is shown in the barplot. Z-scores for FGC, CIN, and tFA in each Thai iCCA sample were plotted in each barplot in the FGC ranked order. (Bottom panel) Shown were 51 genes with mutations of more than 3 samples in Thai iCCA. Right plot shows the mutation frequency for each gene in the frequency order. Dotted line indicates the mutation frequency of 0.1. Left plot shows the occurrence of mutation of regarding gene in each sample. Each bar plot represents each gene. Different color indicates different mutation type. Thai iCCA samples were represented in columns in the same order of top panel. NSYN, non-synonymous mutation; FS, frame shift mutation; SS, splice site mutation; NS, non-sense mutation.

Figure S12

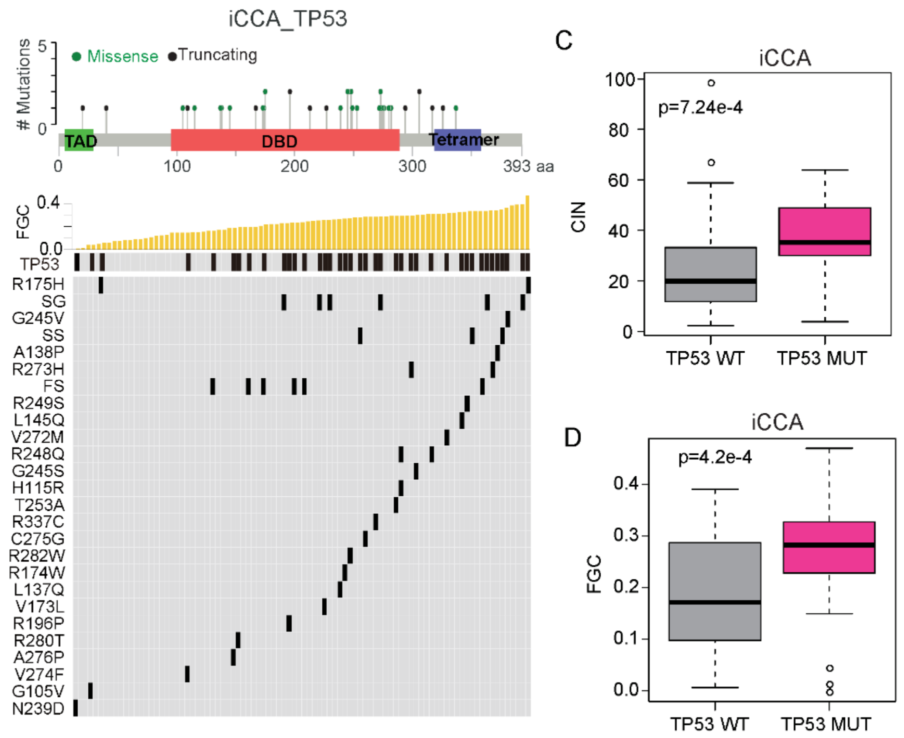


**Figure S12. Integrative analysis based on PCC showed *TP53* as a Cancer functional genomic complexity (FGCs) driver.** (B) (Top panel) Mutation mapper indicates the site where the *TP53* mutation occurred. Transactivation motif (TAD; 6-29), DNA binding motif (DBD; 95-288), and tetramerisation motif (Tetramer; 318-358) were depicted in different colored box; green, orange, and navy, respectively. Green or black dots indicates missense or truncating mutation, respectively. (Bottom panel) Top plot indicates the FGC score of each sample in the rank order. The incidence of TP53 mutation in each sample plotted in black in the bottom plot according to the mutation sites. (C-D) The CIN(C) and PCC (D) between *TP53* WT and *TP53* mutation among Thai iCCA. P-values based on the Welch two-sample t-test were depicted

### Figure S12

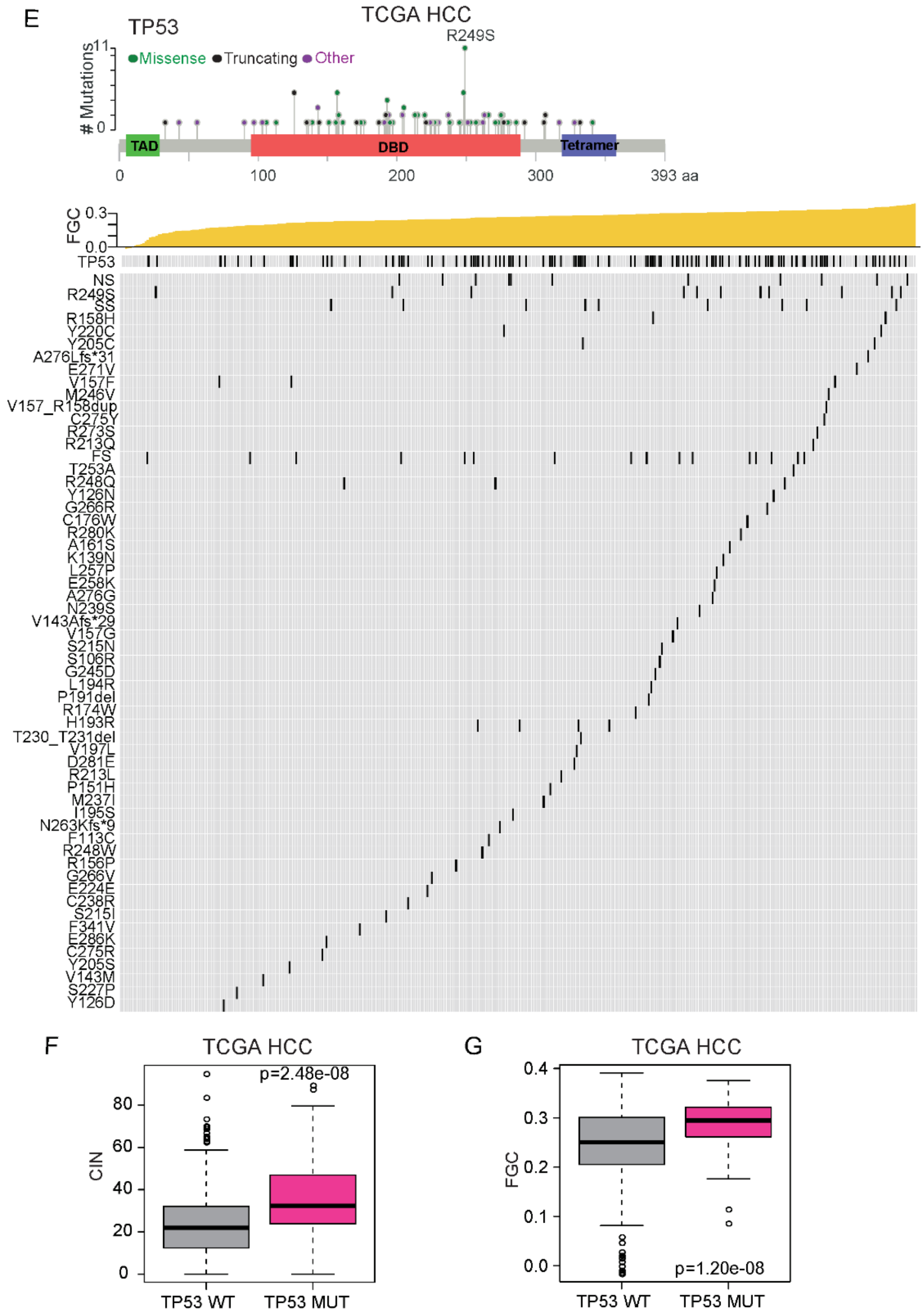


**Figure S12. Integrative analysis based on PCC showed *TP53* as a Cancer functional genomic complexity (FGCs) driver.** (E) (Top panel) Mutation mapper indicates the site where the *TP53* mutation occurred among TCGA HCC. Transactivation motif (TAD) (6-29), DNA binding motif (DBD) (95-288), and tetramerisation motif (Tetramer) (318-358) were depicted in different colored box; green, orange, and navy, respectively. Green or black dots indicates missense or truncating mutation, respectively. (Bottom panel) Top plot indicates the FGC score of each sample in the rank order. The incidence of *TP53* mutation in each sample plotted in black in the bottom plot according to the mutation sites. (F-G) The CIN (F) and FGC (G) level between *TP53* WT and *TP53* mutation among TCGA HCC. P-values based on the Welch two-sample t-test were depicted. NS, SS, and FS stand for non-sense mutation, splice site mutation, and frame shift mutation, respectively

Figure S13

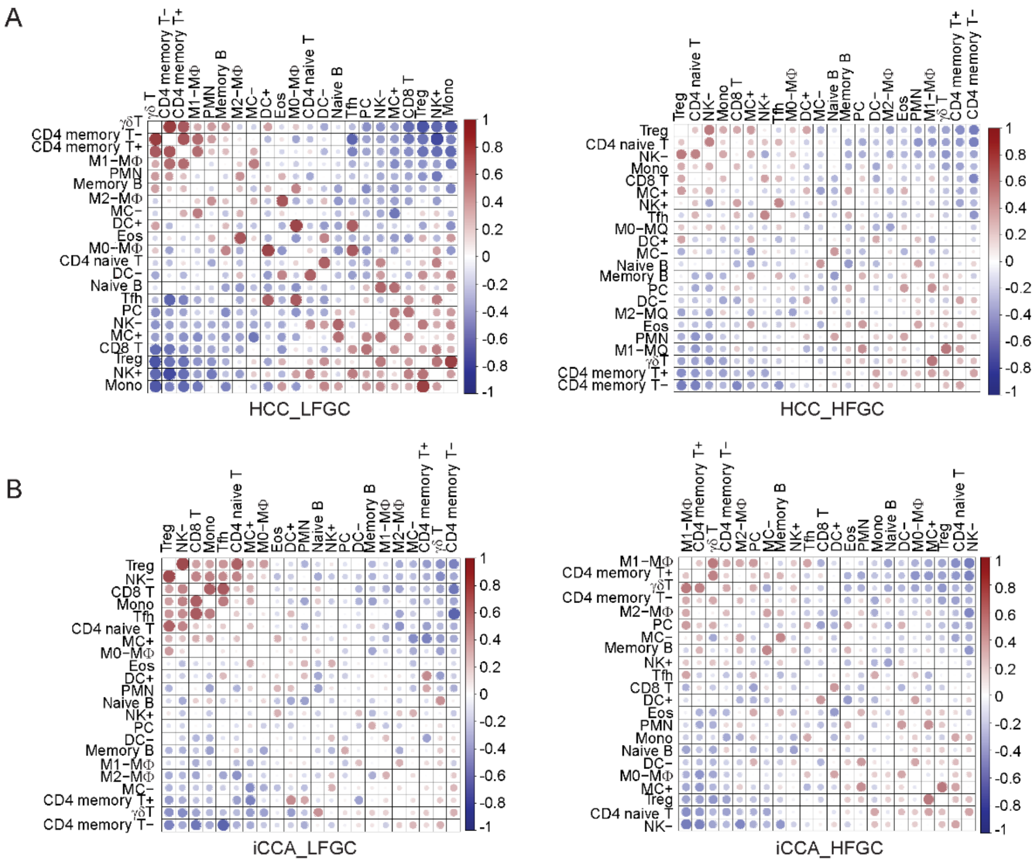


**Figure S13. Comparison of TIL subpopulations between HFGC and LFGC.** (A-B) Associations between 22 types of TIL subpopulations in the LFGC (left panel of each) and HFGC (right panel of each) of Thai HCC (A) and iCCA (B) are shown on a scale from red to blue (1 to -1). Color intensity and the size of the circle are proportional to the correlation coefficients. Proportion of 22 types of TILs based on the CIBERSORT analysis output in the LFGC and HFGC of HCC and iCCA are used.

.

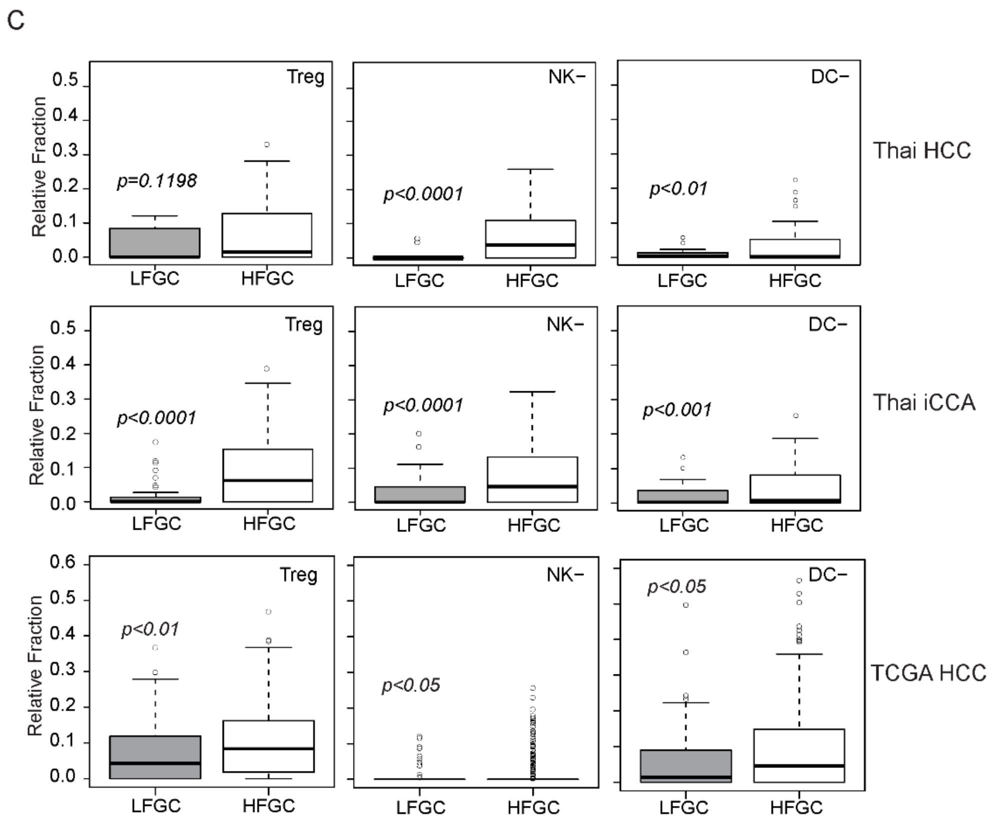


**Figure S13. Comparison of TIL subpopulations between HFGC and LFGC.** (C) Comparison of TIL subpopulations between HFGC and LFGC of Thai HCC, Thai iCCA, and TCGA HCC. Each boxplot shows the relative abundance of TIL subpopulation between HFGC and LFGC. From left to right, representative TILs, regulatory T cell (Treg), NK- cell, dendritic cells (DC) in LFGC and HFGC of Thai HCC (top), Thai iCCA (middle), and TCGA HCC (bottom) are compared. P-values by Welch two-sample t-test are depicted in the plot.

### Figure S14

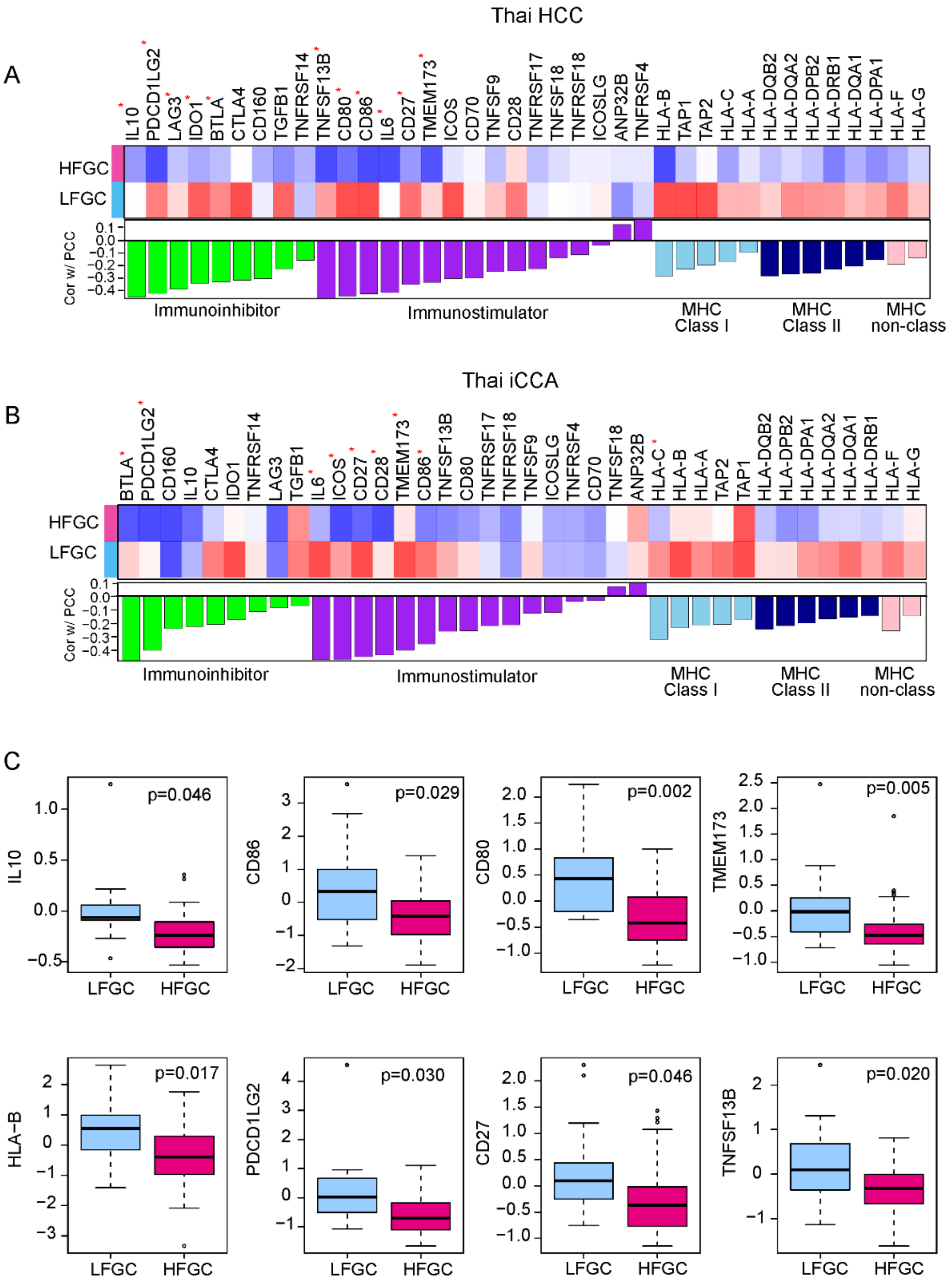


**Figure S14. Association of FGC with immunomodulators** (A-B) (Top panel) Heatmaps show expression level of genes regarding selected inhibitors, stimulators of immune response, MHC class I, II, and non-class among HFGC and LFGC of Thai HCC (A) and iCCA (B), respectively. (Bottom panel) Associations of FGC with selected genes were shown in bar among Thai HCC and iCCA, respectively. Coefficient estimates and p-value based on Pearson's correlation were estimated. Significantly FGC associated genes were marked with red star (p-value <0.01). (C-D) Comparison of selected genes between HFGC and LFGC of Thai HCC (C) Thai iCCA (D). P-values by Welch two-sample t-test are depicted in the plot.

### Figure S14

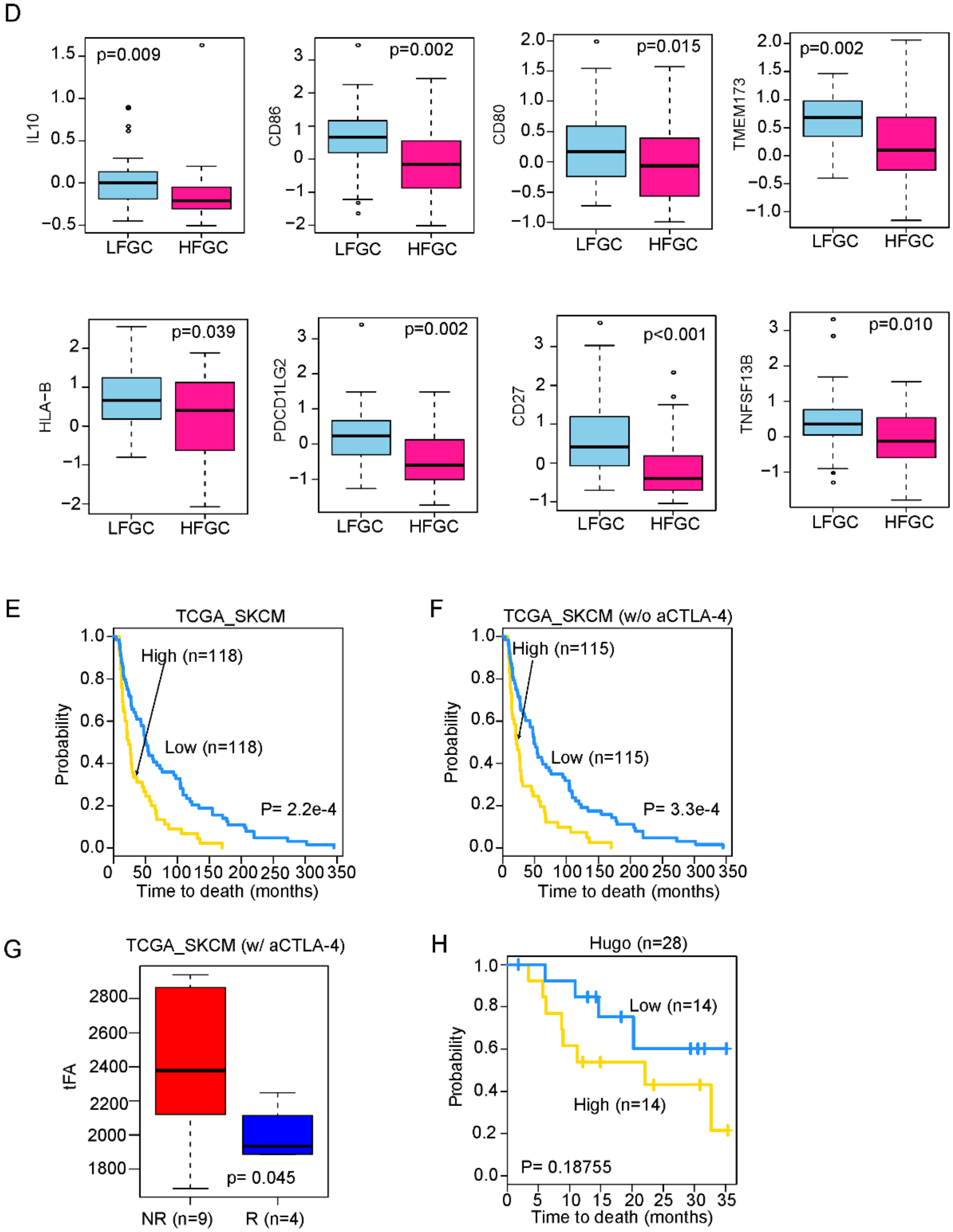


**Figure S14. Association of FGC with immunomodulators** (C-D) Comparison of selected genes between HFGC and LFGC of Thai HCC (C) Thai iCCA (D). P-values by Welch two-sample t-test are depicted in the plot. (E-G) Skin cutaneous melanoma data from TCGA (TCGA_SKCM, n=472) was used to examine the association between tFA and immunotherapy. KM survival analysis was performed including or excluding the patients groups who received pre-treatment of anti-CTLA-4 (E and F, respectively).

Patients were stratified into high and low group based on the tFA level. Patients with tFA levels above 3rd quartile or below 1st quartile were assigned into high and low group, respectively. (G) In the TCGA_SKCM anti-CTLA-4 pre-treatment subset, tFA levels between responders (R) and non-responders (NR) was compared. (H) Metastatic melanoma patients with pre-treatment of anti-PD-1 therapy was used. Patients were stratified into high and low group based on median value of tFA. KM survival analysis was performed between high and low tFA group (H). Numbers of patients in each group are shown.

### References


1. Scharpf RB, Irizarry RA, Ritchie ME, Carvalho B, & Ruczinski I (2011) Using the R Package crlmm for Genotyping and Copy Number Estimation. *J Stat Softw* 40(12):1-32.

2. Carvalho B, Bengtsson H, Speed TP, & Irizarry RA (2007) Exploration, normalization, and genotype calls of high-density oligonucleotide SNP array data. *Biostatistics* 8(2):485-499.

3. Bolstad BM, Irizarry RA, Astrand M, & Speed TP (2003) A comparison of normalization methods for high density oligonucleotide array data based on variance and bias. *Bioinformatics* 19(2):185-193.

4. Olshen AB, Venkatraman ES, Lucito R, & Wigler M (2004) Circular binary segmentation for the analysis of array-based DNA copy number data. *Biostatistics* 5(4):557-572.

5. Roessler S*, et al.* (2010) A unique metastasis gene signature enables prediction of tumor relapse in early-stage hepatocellular carcinoma patients. *Cancer Res.* 70(24):10202-10212.

6. Roessler S*, et al.* (2012) Integrative genomic identification of genes on 8p associated with hepatocellular carcinoma progression and patient survival. *Gastroenterology* 142:957-966.

7. Mermel CH*, et al.* (2011) GISTIC2.0 facilitates sensitive and confident localization of the targets of focal somatic copy-number alteration in human cancers. *Genome Biol.* 12(4):R41.

8. Subramanian A*, et al.* (2005) Gene set enrichment analysis: a knowledge-based approach for interpreting genome-wide expression profiles. *Proc. Natl. Acad. Sci. U. S. A.* 102(43):15545-15550.

9. Carter SL, Eklund AC, Kohane IS, Harris LN, & Szallasi Z (2006) A signature of chromosomal instability inferred from gene expression profiles predicts clinical outcome in multiple human cancers. *Nat. Genet.* 38(9):1043-1048.

10. Huang da W, Sherman BT, & Lempicki RA (2009) Systematic and integrative analysis of large gene lists using DAVID bioinformatics resources. *Nat. Protoc.* 4(1):44-57.

11. Newman AM*, et al.* (2015) Robust enumeration of cell subsets from tissue expression profiles. *Nat Methods* 12(5):453-457.

12. Gao J*, et al.* (2013) Integrative analysis of complex cancer genomics and clinical profiles using the cBioPortal. *Sci Signal* 6(269):pl1.

13. Cerami E, *et al.* (2012) The cBio cancer genomics portal: an open platform for exploring multidimensional cancer genomics data. *Cancer Discov.* 2(5):401-404.

14. Cancer Genome Atlas N (2015) Genomic Classification of Cutaneous Melanoma. *Cell* 161(7):1681-1696.

15. Hugo W, *et al.* (2017) Genomic and Transcriptomic Features of Response to Anti-PD-1 Therapy in Metastatic Melanoma. *Cell* 168(3):542.
