## Supplementary Tables for "Functional Genomic Complexity Defines Intratumor Heterogeneity and Tumor Aggressiveness in Liver Cancer"

**This PDF file includes:**

Tables S1 to S14
- Table S1a. Table for tFA, CIN, GIN and PCC scores among TIGER_LC cohort
- Table S1b. Table for tFA, CIN, and PCC scores for TCGA HCC cohort
- Table S1c. Table for PCC of LCI cohort

Table S2. Overlapped Differentially Expressed Genes among correlated genes between Thai HCC and iCCA
Table S3. Comparison table of overlapped differentially expressed genes among correlated genes.
Table S4. GSEA result compared HFGC with LFGC of Thai iCCA
Table S5. GSEA result compared HFGC with LFGC of Thai HCC
Table S6. GSEA result compared HFGC with LFGC of TCGA HCC
Table S7. Overlapped GSEA result HFGC and LFCG of Thai HCC and Thai iCCA
Table S8. Overlapped recurrent mutation genes between Thai HCC and Thai iCCA
Table S9. Recurrently mutated genes in Thai HCC
Table S10. Recurrently mutated genes in Thai iCCA
Table S11. Information of TP53 mutations mapping.
Table S12. Comparison table of TP53 mutation between HFGC and LFGC among PLC cohorts
Table S13a. Table for tFA of TCGA SKCM cohort
Table S13b. Table for tFA of Hugo Melanoma cohort

### Table S1a. Table for tFA, CIN and GIN and PCC scores among TIGER_LC cohort.

| Thai HCC | | | | | |
|---|---|---|---|---|---|
| **Sample ID** | **tFA** | **CIN** | **GIN** | **PCC** | **Group** |
| LCS_504A | 640.681 | 16.979 | 4.034E+08 | 0.197 | Low |
| LCS_505A | 778.102 | 52.261 | 1.451E+09 | 0.380 | High |
| LCS_506A | 802.326 | 22.908 | 5.356E+08 | 0.382 | High |
| LCS_501A | 825.739 | 31.992 | 8.112E+08 | 0.375 | High |
| LCS_502A | 658.129 | 10.109 | 2.299E+08 | 0.342 | High |
| LCS_503A | 713.620 | 24.450 | 5.497E+08 | 0.313 | High |
| LCS_557A | 769.821 | 18.055 | 3.994E+08 | 0.192 | Low |
| LCS_565A | 598.844 | 11.442 | 1.489E+08 | 0.065 | Low |
| LCS_541A | 785.926 | 53.129 | 1.356E+09 | 0.353 | High |
| LCS_515A | 820.240 | 33.127 | 7.586E+08 | 0.372 | High |
| LCS_530A | 918.184 | 41.706 | 1.135E+09 | 0.487 | High |
| LCS_550A | 757.557 | 62.330 | 1.758E+09 | 0.332 | High |
| LCS_551A | 676.774 | 17.951 | 3.449E+08 | 0.183 | Low |
| LCS_564A | 733.027 | 33.328 | 8.117E+08 | 0.403 | High |
| LCS_567A | 735.945 | 20.485 | 5.346E+08 | 0.243 | High |
| LCS_507A | 839.909 | 33.276 | 9.313E+08 | 0.494 | High |
| LCS_534A | 897.275 | 47.556 | 1.242E+09 | 0.443 | High |
| LCS_542A | 832.472 | 58.284 | 1.665E+09 | 0.420 | High |
| LCS_517A | 703.686 | 35.660 | 9.336E+08 | 0.296 | High |
| LCS_518A | 624.489 | 19.425 | 4.509E+08 | 0.213 | High |
| LCS_536A | 789.915 | 13.971 | 3.666E+08 | 0.297 | High |
| LCS_546A | 803.054 | 55.191 | 1.537E+09 | 0.424 | High |
| LCS_528A | 644.484 | 27.393 | 6.994E+08 | 0.255 | High |
| LCS_512A | 714.272 | 32.523 | 9.036E+08 | 0.324 | High |
| LCS_514A | 669.954 | 24.241 | 6.572E+08 | 0.272 | High |
| LCS_537A | 796.132 | 14.443 | 2.967E+08 | 0.179 | Low |
| LCS_524A | 942.299 | 49.457 | 1.491E+09 | 0.435 | High |
| LCS_540A | 840.796 | 60.515 | 1.603E+09 | 0.430 | High |
| LCS_511A | 724.569 | 13.700 | 2.667E+08 | 0.301 | High |
| LCS_543A | 572.680 | 9.152 | 1.168E+08 | 0.092 | Low |
| LCS_547A | 873.516 | 47.273 | 1.141E+09 | 0.440 | High |
| LCS_533A | 620.124 | 43.539 | 1.078E+09 | 0.135 | Low |
| LCS_549A | 798.784 | 18.315 | 4.313E+08 | 0.302 | High |
| LCS_510A | 664.320 | 25.297 | 5.498E+08 | 0.356 | High |
| LCS_519A | 772.474 | 29.342 | 7.468E+08 | 0.450 | High |
| LCS_529A | 646.271 | 15.923 | 3.524E+08 | 0.338 | High |
| LCS_568A | 776.757 | 46.241 | 1.113E+09 | 0.371 | High |
| LCS_508A | 895.199 | 91.113 | 2.546E+09 | 0.453 | High |

| | | | | | |
|---|---|---|---|---|---|
| LCS_513A | 716.319 | 28.509 | 7.753E+08 | 0.193 | Low |
| LCS_523A | 645.455 | 23.448 | 5.375E+08 | 0.296 | High |
| LCS_552A | 849.871 | 42.182 | 1.243E+09 | 0.412 | High |
| LCS_558A | 670.991 | 15.813 | 3.140E+08 | 0.155 | Low |
| LCS_538A | 574.235 | 14.355 | 2.481E+08 | 0.071 | Low |
| LCS_539A | 628.682 | 21.140 | 4.815E+08 | 0.320 | High |
| LCS_559A | 738.192 | 21.879 | 4.524E+08 | 0.377 | High |
| LCS_516A | 696.648 | 31.652 | 7.028E+08 | 0.277 | High |
| LCS_520A | 769.062 | 37.655 | 1.367E+09 | 0.340 | High |
| LCS_531A | 735.662 | 62.277 | 1.567E+09 | 0.353 | High |
| LCS_553A | 732.752 | 38.133 | 9.934E+08 | 0.300 | High |
| LCS_527A | 700.571 | 16.713 | 3.646E+08 | 0.310 | High |
| LCS_545A | 788.601 | 28.515 | 6.990E+08 | 0.386 | High |
| LCS_532A | 780.401 | 59.329 | 1.436E+09 | 0.411 | High |
| LCS_562A | 748.455 | 28.754 | 6.540E+08 | 0.314 | High |
| LCS_566A | 618.078 | 4.784 | 7.303E+07 | 0.316 | High |
| LCS_584A | 882.992 | 43.545 | 1.180E+09 | 0.427 | High |
| LCS_548A | 633.274 | 13.631 | 3.137E+08 | 0.167 | Low |
| LCS_544A | 839.814 | 75.377 | 2.057E+09 | 0.409 | High |
| LCS_560A | 639.351 | 9.042 | 2.032E+08 | 0.087 | Low |
| LCS_563A | 829.761 | 31.478 | 8.252E+08 | 0.410 | High |
| LCS_509A | 711.933 | 3.624 | 6.673E+07 | -0.107 | Low |
| LCS_535A | 668.365 | 13.556 | 4.155E+08 | 0.009 | Low |
| LCS_525A | 687.470 | 26.698 | 6.159E+08 | 0.241 | High |
| **Thai iCCA** | | | | | |
| **Sample ID** | **tFA** | **CIN** | **GIN** | **PCC** | **Group** |
| LCS_597A | 678.519 | 11.454 | 2.076E+08 | 0.220 | High |
| LCS_623A | 773.185 | 32.819 | 7.289E+08 | 0.342 | High |
| LCS_639A | 609.062 | 22.902 | 7.519E+08 | 0.092 | Low |
| LCS_657A | 713.145 | 58.278 | 1.549E+09 | 0.295 | High |
| LCS_658A | 703.025 | 54.643 | 1.672E+09 | 0.309 | High |
| LCS_580A | 696.278 | 35.215 | 9.142E+08 | 0.262 | High |
| LCS_600A | 684.673 | 33.155 | 9.429E+08 | 0.265 | High |
| LCS_616A | 667.434 | 37.352 | 1.117E+09 | 0.314 | High |
| LCS_637A | 605.441 | 5.480 | 5.683E+07 | 0.089 | Low |
| LCS_650A | 587.171 | 2.256 | 4.643E+07 | 0.102 | Low |
| LCS_651A | 630.054 | 46.268 | 1.212E+09 | 0.225 | High |
| LCS_586A | 647.809 | 34.884 | 1.089E+09 | 0.273 | High |
| LCS_587A | 761.435 | 58.814 | 1.590E+09 | 0.336 | High |
| LCS_595A | 773.857 | 46.443 | 1.341E+09 | 0.470 | High |
| LCS_645A | 697.373 | 56.373 | 1.546E+09 | 0.348 | High |

| | | | | | |
|---|---|---|---|---|---|
| LCS_570A | 628.549 | 54.049 | 1.447E+09 | 0.153 | Low |
| LCS_592A | 757.727 | 24.828 | 6.238E+08 | 0.288 | High |
| LCS_603A | 776.387 | 49.256 | 1.542E+09 | 0.319 | High |
| LCS_684A | 626.820 | 20.176 | 4.883E+08 | 0.150 | Low |
| LCS_643A | 577.480 | 6.113 | 1.374E+08 | 0.065 | Low |
| LCS_601A | 580.850 | 6.497 | 1.092E+08 | 0.086 | Low |
| LCS_612A | 635.219 | 29.895 | 7.444E+08 | 0.282 | High |
| LCS_653A | 615.784 | 30.508 | 9.623E+08 | 0.148 | Low |
| LCS_578A | 659.671 | 38.443 | 9.926E+08 | 0.294 | High |
| LCS_618A | 649.149 | 26.224 | 6.003E+08 | 0.251 | High |
| LCS_573A | 769.858 | 30.096 | 8.296E+08 | -0.002 | Low |
| LCS_681A | 682.591 | 41.728 | 1.101E+09 | 0.298 | High |
| LCS_575A | 590.652 | 13.258 | 2.288E+08 | 0.196 | Low |
| LCS_583A | 687.135 | 53.495 | 1.495E+09 | 0.295 | High |
| LCS_614A | 495.974 | 10.878 | 2.812E+08 | 0.053 | Low |
| LCS_634A | 674.090 | 15.298 | 4.095E+08 | 0.240 | High |
| LCS_664A | 732.907 | 47.914 | 1.410E+09 | 0.314 | High |
| LCS_596A | 705.811 | 67.105 | 1.824E+09 | 0.303 | High |
| LCS_598A | 722.202 | 52.126 | 1.418E+09 | 0.362 | High |
| LCS_644A | 703.386 | 44.953 | 1.104E+09 | 0.340 | High |
| LCS_646A | 784.935 | 54.046 | 1.530E+09 | 0.336 | High |
| LCS_674A | 594.768 | 44.826 | 1.064E+09 | 0.188 | Low |
| LCS_581A | 718.540 | 19.636 | 3.852E+08 | 0.154 | Low |
| LCS_626A | 751.591 | 50.648 | 1.327E+09 | 0.335 | High |
| LCS_668A | 566.749 | 17.829 | 4.078E+08 | 0.111 | Low |
| LCS_585A | 601.376 | 3.536 | 5.582E+07 | 0.091 | Low |
| LCS_602A | 631.518 | 16.423 | 3.794E+08 | 0.081 | Low |
| LCS_606A | 518.923 | 11.716 | 3.259E+08 | 0.019 | Low |
| LCS_621A | 649.324 | 19.102 | 4.349E+08 | 0.124 | Low |
| LCS_615A | 653.261 | 30.296 | 6.975E+08 | 0.246 | High |
| LCS_631A | 629.553 | 49.979 | 1.333E+09 | 0.257 | High |
| LCS_675A | 638.788 | 13.624 | 2.643E+08 | 0.166 | Low |
| LCS_683A | 594.087 | 19.823 | 6.047E+08 | 0.164 | Low |
| LCS_687A | 799.126 | 35.146 | 1.016E+09 | 0.297 | High |
| LCS_636A | 577.275 | 11.827 | 4.381E+08 | 0.128 | Low |
| LCS_574A | 728.821 | 43.278 | 1.199E+09 | 0.394 | High |
| LCS_624A | 666.230 | 3.808 | 8.968E+07 | 0.149 | Low |
| LCS_688A | 684.929 | 31.601 | 8.239E+08 | 0.255 | High |
| LCS_608A | 775.442 | 98.621 | 2.713E+09 | 0.390 | High |
| LCS_610A | 707.752 | 47.889 | 1.270E+09 | 0.336 | High |
| LCS_625A | 715.527 | 51.786 | 1.464E+09 | 0.285 | High |

| | | | | | |
|---|---|---|---|---|---|
| LCS_590A | 658.820 | 32.303 | 7.169E+08 | 0.309 | High |
| LCS_638A | 704.271 | 31.870 | 7.962E+08 | 0.315 | High |
| LCS_642A | 594.571 | 10.872 | 1.611E+08 | 0.045 | Low |
| LCS_670A | 522.376 | 17.844 | 5.368E+08 | 0.013 | Low |
| LCS_572A | 657.343 | 51.279 | 1.299E+09 | 0.330 | High |
| LCS_577A | 688.471 | 30.125 | 6.198E+08 | 0.172 | Low |
| LCS_671A | 685.355 | 16.495 | 3.433E+08 | 0.234 | High |
| LCS_571A | 686.204 | 38.836 | 1.159E+09 | 0.233 | High |
| LCS_599A | 640.682 | 23.899 | 6.156E+08 | 0.260 | High |
| LCS_604A | 707.234 | 19.668 | 5.608E+08 | 0.237 | High |
| LCS_613A | 548.419 | 4.191 | 8.803E+07 | 0.077 | Low |
| LCS_667A | 630.378 | 3.064 | 6.197E+07 | 0.007 | Low |
| LCS_678A | 564.741 | 11.787 | 2.242E+08 | 0.045 | Low |
| LCS_635A | 634.352 | 17.341 | 3.854E+08 | 0.192 | Low |
| LCS_641A | 593.070 | 7.808 | 9.613E+07 | 0.063 | Low |
| LCS_677A | 778.054 | 31.913 | 8.536E+08 | 0.382 | High |
| LCS_579A | 589.743 | 16.307 | 3.418E+08 | 0.129 | Low |
| LCS_620A | 712.323 | 55.379 | 1.723E+09 | 0.286 | High |
| LCS_628A | 751.489 | 7.155 | 2.156E+08 | 0.198 | Low |
| LCS_630A | 759.285 | 30.049 | 8.186E+08 | 0.225 | High |
| LCS_680A | 787.884 | 63.929 | 1.692E+09 | 0.195 | Low |
| LCS_633A | 611.311 | 12.287 | 1.708E+08 | 0.150 | Low |
| LCS_640A | 732.058 | 28.958 | 7.543E+08 | 0.269 | High |
| LCS_649A | 725.227 | 30.076 | 8.488E+08 | 0.323 | High |
| LCS_685A | 596.239 | 22.752 | 5.781E+08 | 0.202 | High |
| LCS_594A | 668.579 | 34.094 | 9.314E+08 | 0.288 | High |
| LCS_627A | 643.123 | 30.926 | 7.291E+08 | 0.283 | High |
| LCS_648A | 638.022 | 24.899 | 6.340E+08 | 0.209 | High |
| LCS_654A | 563.573 | 20.772 | 4.626E+08 | 0.006 | Low |
| LCS_679A | 681.193 | 20.499 | 5.922E+08 | 0.231 | High |
| LCS_609A | 646.746 | 26.012 | 6.495E+08 | 0.172 | Low |
| LCS_652A | 731.838 | 51.120 | 1.345E+09 | 0.288 | High |
| LCS_669A | 647.614 | 20.800 | 5.065E+08 | 0.158 | Low |
| LCS_682A | 695.584 | 34.073 | 9.812E+08 | 0.304 | High |

### Table S1b. Table for tFA, CIN and PCC for TCGA HCC cohort

| Sample ID | tFA | CIN | PCC | Group |
|---|---|---|---|---|
| TCGA-DD-AACO-01A-11R-A41C-07 | 260.416 | 31.593 | 0.188 | Low |
| TCGA-CC-5263-01A-01R-A131-07 | 231.347 | 32.143 | 0.278 | High |
| TCGA-CC-5261-01A-01R-A131-07 | 270.961 | 45.330 | 0.314 | High |
| TCGA-CC-A5UC-01A-11R-A28V-07 | 236.177 | 22.802 | 0.231 | High |
| TCGA-RC-A6M5-01A-11R-A32O-07 | 233.243 | 1.648 | 0.153 | Low |
| TCGA-DD-A1EH-01A-11R-A131-07 | 287.457 | 23.901 | 0.314 | High |
| TCGA-ED-A8O5-01A-11R-A36F-07 | 314.650 | 25.824 | 0.347 | High |
| TCGA-CC-5264-01A-01R-A131-07 | 266.629 | 69.780 | 0.335 | High |
| TCGA-DD-AACS-01A-11R-A41C-07 | 273.232 | 41.484 | 0.322 | High |
| TCGA-G3-A5SL-01A-11R-A27V-07 | 275.664 | 17.582 | 0.330 | High |
| TCGA-DD-AACE-01A-11R-A41C-07 | 291.702 | 56.044 | 0.327 | High |
| TCGA-BC-4072-01B-11R-A155-07 | 235.645 | 27.198 | 0.235 | High |
| TCGA-CC-A7II-01A-11R-A33J-07 | 298.547 | 44.780 | 0.259 | High |
| TCGA-DD-AAEK-01A-11R-A41C-07 | 256.899 | 37.363 | 0.322 | High |
| TCGA-DD-AACB-01A-11R-A41C-07 | 267.812 | 48.901 | 0.281 | High |
| TCGA-EP-A3JL-01A-11R-A213-07 | 257.045 | 13.462 | 0.273 | High |
| TCGA-EP-A2KB-01A-11R-A180-07 | 304.423 | 56.319 | 0.303 | High |
| TCGA-ED-A4XI-01A-11R-A266-07 | 226.076 | 5.769 | 0.117 | Low |
| TCGA-G3-A7M7-01A-12R-A352-07 | 246.379 | 3.297 | 0.175 | Low |
| TCGA-CC-A7IG-01A-11R-A33J-07 | 262.417 | 55.769 | 0.267 | High |
| TCGA-DD-AACT-01A-11R-A41C-07 | 289.843 | 28.022 | 0.331 | High |
| TCGA-UB-A7MF-01A-11R-A33J-07 | 273.117 | 22.802 | 0.237 | High |
| TCGA-HP-A5N0-01A-11R-A28V-07 | 261.943 | 17.582 | 0.341 | High |
| TCGA-CC-5260-01A-01R-A131-07 | 260.224 | 27.747 | 0.203 | High |
| TCGA-G3-A25X-01A-11R-A16W-07 | 266.378 | 37.912 | 0.279 | High |
| TCGA-DD-AAE9-01A-11R-A41C-07 | 264.033 | 34.615 | 0.270 | High |
| TCGA-RC-A7SF-01A-11R-A352-07 | 236.807 | 18.681 | 0.330 | High |
| TCGA-NI-A8LF-01A-11R-A36F-07 | 262.422 | 47.253 | 0.345 | High |
| TCGA-2Y-A9H6-01A-11R-A39D-07 | 267.611 | 21.429 | 0.280 | High |
| TCGA-DD-AAD6-01A-11R-A41C-07 | 272.579 | 27.198 | 0.326 | High |
| TCGA-ZP-A9D2-01A-11R-A38B-07 | 256.995 | 31.044 | 0.314 | High |
| TCGA-DD-A4NQ-01A-21R-A28V-07 | 232.242 | 32.143 | 0.304 | High |
| TCGA-NI-A4U2-01A-11R-A28V-07 | 253.876 | 16.484 | 0.194 | Low |
| TCGA-ED-A97K-01A-21R-A38B-07 | 255.413 | 34.066 | 0.224 | High |
| TCGA-DD-A4NB-01A-12R-A266-07 | 213.755 | 8.516 | 0.153 | Low |
| TCGA-2Y-A9H7-01A-11R-A39D-07 | 255.019 | 15.934 | 0.273 | High |
| TCGA-CC-A3MC-01A-11R-A22L-07 | 247.075 | 18.407 | 0.322 | High |
| TCGA-BC-A10Q-01A-11R-A131-07 | 268.860 | 70.055 | 0.193 | Low |
| TCGA-MI-A75E-01A-11R-A32O-07 | 224.774 | 13.736 | 0.177 | Low |

| | | | | |
|---|---|---|---|---|
| TCGA-WX-AA46-01A-11R-A39D-07 | 253.677 | 6.593 | 0.151 | Low |
| TCGA-FV-A3I1-01A-11R-A22L-07 | 263.288 | 25.824 | 0.287 | High |
| TCGA-DD-A4NJ-01A-11R-A27V-07 | 260.172 | 27.473 | 0.317 | High |
| TCGA-DD-AAVY-01A-11R-A41C-07 | 287.682 | 17.857 | 0.283 | High |
| TCGA-DD-AACK-01A-11R-A41C-07 | 228.015 | 3.022 | 0.122 | Low |
| TCGA-DD-AAE2-01A-11R-A41C-07 | 232.749 | 13.187 | 0.249 | High |
| TCGA-XR-A8TF-01A-11R-A36F-07 | 297.783 | 53.297 | 0.279 | High |
| TCGA-KR-A7K8-01A-11R-A33J-07 | 249.926 | 18.681 | 0.252 | High |
| TCGA-DD-A118-01A-11R-A131-07 | 255.236 | 47.253 | 0.246 | High |
| TCGA-BD-A3EP-01A-11R-A22L-07 | 231.000 | 34.341 | 0.278 | High |
| TCGA-KR-A7K2-01A-12R-A33R-07 | 276.767 | 58.791 | 0.267 | High |
| TCGA-ED-A7XO-01A-11R-A352-07 | 305.880 | 20.330 | 0.309 | High |
| TCGA-DD-A4NF-01A-11R-A27V-07 | 259.792 | 4.945 | 0.130 | Low |
| TCGA-ZS-A9CG-01A-11R-A37K-07 | 232.120 | 9.890 | 0.214 | High |
| TCGA-EP-A3RK-01A-11R-A22L-07 | 241.578 | 18.956 | 0.283 | High |
| TCGA-DD-A1EK-01A-11R-A213-07 | 248.430 | 11.538 | 0.244 | High |
| TCGA-WJ-A86L-01A-12R-A39D-07 | 255.149 | 20.879 | 0.241 | High |
| TCGA-DD-A1ED-01A-11R-A155-07 | 226.657 | 0.275 | 0.016 | Low |
| TCGA-G3-AAV3-01A-11R-A37K-07 | 236.328 | 9.615 | 0.226 | High |
| TCGA-2Y-A9H5-01A-11R-A38B-07 | 284.378 | 25.824 | 0.311 | High |
| TCGA-G3-A7M8-01A-11R-A33R-07 | 224.294 | 0.000 | -0.013 | Low |
| TCGA-DD-A4NK-01A-11R-A28V-07 | 282.209 | 20.604 | 0.283 | High |
| TCGA-DD-A39X-01A-11R-A213-07 | 256.587 | 12.637 | 0.321 | High |
| TCGA-DD-A39Z-01A-11R-A213-07 | 284.736 | 29.670 | 0.233 | High |
| TCGA-DD-AAD2-01A-11R-A41C-07 | 241.354 | 9.341 | 0.269 | High |
| TCGA-DD-AAE7-01A-11R-A41C-07 | 213.623 | 5.220 | 0.173 | Low |
| TCGA-BC-A216-01A-11R-A155-07 | 255.448 | 23.352 | 0.263 | High |
| TCGA-DD-A11A-01A-11R-A131-07 | 261.165 | 57.967 | 0.334 | High |
| TCGA-CC-A8HV-01A-11R-A36F-07 | 301.973 | 69.505 | 0.337 | High |
| TCGA-G3-AAV4-01A-11R-A38B-07 | 305.411 | 31.868 | 0.247 | High |
| TCGA-DD-A4NP-01A-11R-A28V-07 | 255.375 | 7.143 | 0.140 | Low |
| TCGA-ED-A459-01A-11R-A266-07 | 270.033 | 39.286 | 0.357 | High |
| TCGA-DD-AADV-01A-11R-A39D-07 | 287.640 | 31.319 | 0.300 | High |
| TCGA-DD-AADF-01A-11R-A41C-07 | 295.856 | 45.604 | 0.253 | High |
| TCGA-2Y-A9H8-01A-11R-A39D-07 | 275.156 | 55.769 | 0.298 | High |
| TCGA-DD-A113-01A-11R-A131-07 | 240.990 | 64.286 | 0.314 | High |
| TCGA-G3-AAV7-01A-11R-A38B-07 | 264.372 | 48.626 | 0.318 | High |
| TCGA-CC-A8HT-01A-11R-A36F-07 | 246.263 | 25.000 | 0.275 | High |
| TCGA-G3-A3CK-01A-11R-A213-07 | 255.166 | 10.989 | 0.180 | Low |
| TCGA-DD-A73D-01A-12R-A32O-07 | 286.004 | 35.440 | 0.301 | High |
| TCGA-RC-A7SK-01A-11R-A352-07 | 281.033 | 26.374 | 0.292 | High |

| | | | | |
|---|---|---|---|---|
| TCGA-DD-AAEB-01A-11R-A41C-07 | 270.490 | 15.110 | 0.147 | Low |
| TCGA-ZS-A9CD-01A-11R-A37K-07 | 245.998 | 10.989 | 0.213 | High |
| TCGA-DD-A4NN-01A-11R-A28V-07 | 301.580 | 39.011 | 0.347 | High |
| TCGA-T1-A6J8-01A-11R-A32O-07 | 246.107 | 23.901 | 0.330 | High |
| TCGA-FV-A4ZP-01A-12R-A266-07 | 269.774 | 50.549 | 0.226 | High |
| TCGA-DD-AADM-01A-11R-A41C-07 | 258.412 | 31.593 | 0.285 | High |
| TCGA-BC-A10Z-01A-11R-A131-07 | 276.224 | 25.549 | 0.238 | High |
| TCGA-BW-A5NO-01A-11R-A27V-07 | 282.413 | 54.396 | 0.321 | High |
| TCGA-DD-AADA-01A-11R-A41C-07 | 279.322 | 26.648 | 0.289 | High |
| TCGA-DD-A3A4-01A-11R-A22L-07 | 272.748 | 7.692 | 0.120 | Low |
| TCGA-DD-AAW2-01A-11R-A41C-07 | 280.589 | 23.901 | 0.318 | High |
| TCGA-DD-AAC8-01A-11R-A41C-07 | 278.117 | 45.330 | 0.291 | High |
| TCGA-DD-A119-01A-11R-A131-07 | 261.286 | 29.945 | 0.294 | High |
| TCGA-DD-A3A3-01A-11R-A22L-07 | 287.410 | 19.231 | 0.247 | High |
| TCGA-DD-AACQ-01A-11R-A41C-07 | 260.112 | 19.505 | 0.279 | High |
| TCGA-BC-A110-01A-11R-A131-07 | 238.159 | 0.000 | 0.021 | Low |
| TCGA-DD-AACU-01A-11R-A41C-07 | 262.727 | 77.473 | 0.334 | High |
| TCGA-PD-A5DF-01A-11R-A27V-07 | 218.277 | 18.407 | 0.264 | High |
| TCGA-DD-A3A8-01A-11R-A22L-07 | 267.811 | 13.187 | 0.237 | High |
| TCGA-K7-A6G5-01A-11R-A311-07 | 253.425 | 7.967 | 0.195 | Low |
| TCGA-LG-A9QD-01A-11R-A38B-07 | 246.293 | 9.890 | 0.200 | Low |
| TCGA-EP-A2KC-01A-11R-A213-07 | 267.252 | 39.011 | 0.330 | High |
| TCGA-DD-AADD-01A-11R-A41C-07 | 294.868 | 41.209 | 0.267 | High |
| TCGA-FV-A4ZQ-01A-11R-A266-07 | 244.964 | 26.374 | 0.248 | High |
| TCGA-BC-A5W4-01A-11R-A28V-07 | 278.064 | 79.670 | 0.308 | High |
| TCGA-ED-A82E-01A-11R-A352-07 | 266.591 | 62.637 | 0.215 | High |
| TCGA-BC-A10U-01A-11R-A131-07 | 264.661 | 28.022 | 0.332 | High |
| TCGA-BD-A3ER-01A-11R-A213-07 | 234.864 | 7.692 | 0.180 | Low |
| TCGA-DD-AAVV-01A-11R-A41C-07 | 249.730 | 35.440 | 0.335 | High |
| TCGA-BC-A3KG-01A-11R-A213-07 | 307.723 | 35.165 | 0.307 | High |
| TCGA-FV-A2QR-01A-11R-A213-07 | 235.086 | 26.099 | 0.293 | High |
| TCGA-DD-AAC9-01A-11R-A41C-07 | 247.083 | 9.615 | 0.197 | Low |
| TCGA-DD-A4NI-01A-11R-A27V-07 | 251.822 | 17.857 | 0.264 | High |
| TCGA-5C-AAPD-01A-21R-A39D-07 | 297.216 | 46.703 | 0.317 | High |
| TCGA-MI-A75I-01A-11R-A32O-07 | 282.767 | 27.747 | 0.359 | High |
| TCGA-FV-A3R2-01A-11R-A22L-07 | 284.317 | 37.912 | 0.376 | High |
| TCGA-DD-A1EL-01A-11R-A155-07 | 256.835 | 41.758 | 0.312 | High |
| TCGA-DD-A1EA-01A-11R-A131-07 | 255.273 | 15.385 | 0.220 | High |
| TCGA-DD-A73B-01A-12R-A32O-07 | 285.105 | 67.033 | 0.342 | High |
| TCGA-G3-A25S-01A-11R-A16W-07 | 271.510 | 53.297 | 0.360 | High |
| TCGA-UB-AA0U-01A-11R-A38B-07 | 256.133 | 35.165 | 0.332 | High |

| | | | | |
|---|---|---|---|---|
| TCGA-FV-A3I0-01A-11R-A22L-07 | 274.356 | 32.692 | 0.249 | High |
| TCGA-FV-A23B-01A-11R-A16W-07 | 290.434 | 26.099 | 0.315 | High |
| TCGA-2V-A95S-01A-11R-A37K-07 | 272.212 | 19.780 | 0.208 | High |
| TCGA-DD-AADL-01A-11R-A41C-07 | 291.173 | 38.462 | 0.351 | High |
| TCGA-CC-5258-01A-01R-A131-07 | 262.818 | 28.571 | 0.295 | High |
| TCGA-EP-A12J-01A-11R-A131-07 | 275.494 | 12.912 | 0.238 | High |
| TCGA-5R-AA1C-01A-11R-A41C-07 | 272.862 | 30.220 | 0.267 | High |
| TCGA-DD-A3A6-01A-11R-A22L-07 | 262.330 | 0.000 | 0.017 | Low |
| TCGA-DD-AADC-01A-11R-A41C-07 | 259.797 | 40.110 | 0.367 | High |
| TCGA-DD-A3A1-01A-11R-A213-07 | 247.331 | 27.473 | 0.284 | High |
| TCGA-DD-AACA-01A-11R-A41C-07 | 279.834 | 22.802 | 0.231 | High |
| TCGA-DD-A4NA-01A-11R-A266-07 | 277.137 | 25.275 | 0.170 | Low |
| TCGA-MI-A75H-01A-11R-A32O-07 | 249.304 | 10.989 | 0.257 | High |
| TCGA-CC-5262-01A-01R-A131-07 | 231.546 | 16.484 | 0.206 | High |
| TCGA-DD-AAE3-01A-11R-A41C-07 | 257.778 | 34.341 | 0.253 | High |
| TCGA-2Y-A9H4-01A-11R-A38B-07 | 252.345 | 20.604 | 0.261 | High |
| TCGA-DD-A3A7-01A-11R-A22L-07 | 271.264 | 50.000 | 0.267 | High |
| TCGA-DD-AAEH-01A-11R-A41C-07 | 247.020 | 17.033 | 0.288 | High |
| TCGA-FV-A496-01A-11R-A266-07 | 293.555 | 44.231 | 0.306 | High |
| TCGA-G3-AAV2-01A-11R-A37K-07 | 278.249 | 16.484 | 0.223 | High |
| TCGA-MI-A75G-01A-11R-A32O-07 | 298.360 | 38.462 | 0.269 | High |
| TCGA-DD-A1EF-01A-11R-A131-07 | 286.839 | 33.242 | 0.231 | High |
| TCGA-GJ-A3OU-01A-31R-A38B-07 | 217.999 | 12.912 | 0.225 | High |
| TCGA-GJ-A9DB-01A-11R-A37K-07 | 230.459 | 8.242 | 0.231 | High |
| TCGA-DD-A11D-01A-11R-A131-07 | 254.002 | 18.956 | 0.206 | High |
| TCGA-CC-A1HT-01A-11R-A131-07 | 256.159 | 19.231 | 0.229 | High |
| TCGA-DD-AADW-01A-11R-A39D-07 | 285.979 | 36.538 | 0.316 | High |
| TCGA-CC-A9FU-01A-11R-A37K-07 | 301.514 | 63.187 | 0.301 | High |
| TCGA-ED-A5KG-01A-11R-A27V-07 | 257.727 | 0.275 | 0.185 | Low |
| TCGA-DD-AAEE-01A-11R-A41C-07 | 263.540 | 31.868 | 0.340 | High |
| TCGA-WX-AA47-01A-11R-A39D-07 | 257.448 | 13.736 | 0.143 | Low |
| TCGA-BC-A10S-01A-22R-A131-07 | 272.285 | 9.890 | 0.229 | High |
| TCGA-G3-A3CJ-01A-11R-A213-07 | 261.481 | 16.484 | 0.198 | Low |
| TCGA-DD-A39Y-01A-11R-A213-07 | 257.197 | 45.879 | 0.298 | High |
| TCGA-FV-A495-01A-11R-A266-07 | 281.040 | 26.099 | 0.161 | Low |
| TCGA-DD-AACP-01A-11R-A41C-07 | 265.545 | 52.473 | 0.276 | High |
| TCGA-DD-A4NR-01A-11R-A311-07 | 229.735 | 4.670 | 0.221 | High |
| TCGA-DD-AADP-01A-11R-A39D-07 | 241.814 | 26.923 | 0.384 | High |
| TCGA-DD-AAD5-01A-11R-A41C-07 | 249.788 | 52.473 | 0.232 | High |
| TCGA-RC-A7SH-01A-11R-A38B-07 | 318.990 | 35.165 | 0.281 | High |
| TCGA-DD-AAD8-01A-11R-A41C-07 | 282.463 | 76.374 | 0.324 | High |

| | | | | |
|---|---|---|---|---|
| TCGA-DD-AAVP-01A-11R-A41C-07 | 268.066 | 18.132 | 0.250 | High |
| TCGA-DD-AACI-01A-11R-A41C-07 | 274.140 | 36.264 | 0.283 | High |
| TCGA-DD-A114-01A-11R-A131-07 | 268.878 | 9.341 | 0.238 | High |
| TCGA-2Y-A9GW-01A-11R-A38B-07 | 254.905 | 9.066 | 0.307 | High |
| TCGA-DD-A116-01A-11R-A131-07 | 278.716 | 21.978 | 0.262 | High |
| TCGA-DD-AACW-01A-11R-A41C-07 | 283.124 | 23.352 | 0.335 | High |
| TCGA-DD-A39W-01A-11R-A213-07 | 254.580 | 9.890 | 0.137 | Low |
| TCGA-2Y-A9GU-01A-11R-A38B-07 | 259.268 | 32.143 | 0.235 | High |
| TCGA-CC-A9FV-01A-11R-A37K-07 | 307.515 | 0.000 | -0.009 | Low |
| TCGA-DD-AAEG-01A-11R-A39D-07 | 260.499 | 22.527 | 0.340 | High |
| TCGA-DD-AAW0-01A-11R-A41C-07 | 254.335 | 29.121 | 0.376 | High |
| TCGA-2Y-A9H0-01A-11R-A38B-07 | 255.371 | 22.527 | 0.314 | High |
| TCGA-DD-AACG-01A-11R-A41C-07 | 269.025 | 39.560 | 0.333 | High |
| TCGA-RG-A7D4-01A-12R-A33R-07 | 279.351 | 47.253 | 0.333 | High |
| TCGA-ZP-A9D0-01A-11R-A37K-07 | 266.361 | 16.484 | 0.281 | High |
| TCGA-DD-AAVX-01A-11R-A41C-07 | 243.246 | 10.989 | 0.170 | Low |
| TCGA-2Y-A9GS-01A-12R-A38B-07 | 257.149 | 27.473 | 0.320 | High |
| TCGA-XR-A8TG-01A-11R-A36F-07 | 250.933 | 21.703 | 0.266 | High |
| TCGA-ZP-A9CZ-01A-11R-A38B-07 | 225.622 | 22.253 | 0.290 | High |
| TCGA-DD-A1EC-01A-21R-A131-07 | 241.721 | 21.429 | 0.140 | Low |
| TCGA-ED-A8O6-01A-11R-A36F-07 | 269.299 | 52.747 | 0.280 | High |
| TCGA-G3-A3CH-01A-11R-A22L-07 | 250.513 | 14.011 | 0.231 | High |
| TCGA-EP-A26S-01A-11R-A16W-07 | 277.035 | 11.813 | 0.162 | Low |
| TCGA-BC-A69I-01A-11R-A311-07 | 264.234 | 12.088 | 0.220 | High |
| TCGA-RC-A7S9-01A-11R-A33R-07 | 253.401 | 19.231 | 0.249 | High |
| TCGA-BW-A5NQ-01A-11R-A27V-07 | 256.292 | 35.165 | 0.249 | High |
| TCGA-CC-A8HU-01A-11R-A36F-07 | 310.050 | 48.077 | 0.362 | High |
| TCGA-BC-A3KF-01A-11R-A213-07 | 274.568 | 35.714 | 0.391 | High |
| TCGA-DD-A4NO-01A-11R-A28V-07 | 268.805 | 65.110 | 0.344 | High |
| TCGA-ED-A7PZ-01A-11R-A33R-07 | 298.805 | 60.989 | 0.330 | High |
| TCGA-DD-AAE0-01A-11R-A41C-07 | 294.683 | 53.297 | 0.327 | High |
| TCGA-LG-A9QC-01A-11R-A37K-07 | 273.416 | 30.495 | 0.280 | High |
| TCGA-DD-AACD-01A-11R-A41C-07 | 259.914 | 16.209 | 0.230 | High |
| TCGA-BC-A217-01A-11R-A155-07 | 293.566 | 51.923 | 0.340 | High |
| TCGA-5C-A9VH-01A-11R-A37K-07 | 247.693 | 18.681 | 0.289 | High |
| TCGA-G3-A25Z-01A-11R-A16W-07 | 239.946 | 38.462 | 0.282 | High |
| TCGA-CC-A3MA-01A-11R-A213-07 | 260.424 | 37.637 | 0.264 | High |
| TCGA-UB-A7MC-01A-11R-A33R-07 | 290.010 | 45.055 | 0.325 | High |
| TCGA-ZP-A9CV-01A-11R-A38B-07 | 279.904 | 10.165 | 0.248 | High |
| TCGA-BD-A2L6-01A-11R-A213-07 | 273.637 | 18.407 | 0.237 | High |
| TCGA-GJ-A6C0-01A-12R-A311-07 | 260.885 | 23.626 | 0.278 | High |

| | | | | |
|---|---|---|---|---|
| TCGA-DD-AAW1-01A-11R-A41C-07 | 246.586 | 10.714 | 0.185 | Low |
| TCGA-DD-AADB-01A-11R-A41C-07 | 259.629 | 31.319 | 0.333 | High |
| TCGA-UB-A7MD-01A-12R-A352-07 | 257.889 | 18.132 | 0.242 | High |
| TCGA-2Y-A9HA-01A-11R-A39D-07 | 260.797 | 21.154 | 0.322 | High |
| TCGA-ZP-A9D1-01A-11R-A38B-07 | 248.282 | 49.725 | 0.283 | High |
| TCGA-BC-4073-01B-02R-A131-07 | 224.987 | 8.516 | 0.146 | Low |
| TCGA-G3-A7M9-01A-23R-A352-07 | 253.649 | 48.077 | 0.242 | High |
| TCGA-RC-A6M6-01A-11R-A32O-07 | 235.672 | 23.077 | 0.261 | High |
| TCGA-CC-A7IH-01A-11R-A33J-07 | 278.554 | 17.582 | 0.251 | High |
| TCGA-DD-A4NH-01A-11R-A27V-07 | 260.406 | 35.440 | 0.356 | High |
| TCGA-DD-AAE6-01A-11R-A41C-07 | 300.530 | 57.418 | 0.303 | High |
| TCGA-WX-AA44-01A-11R-A39D-07 | 250.887 | 23.077 | 0.284 | High |
| TCGA-YA-A8S7-01A-11R-A37K-07 | 234.805 | 28.846 | 0.216 | High |
| TCGA-DD-AADG-01A-11R-A41C-07 | 290.507 | 24.451 | 0.238 | High |
| TCGA-DD-A4NL-01A-11R-A28V-07 | 218.028 | 0.000 | 0.028 | Low |
| TCGA-G3-A6UC-01A-21R-A33J-07 | 285.104 | 20.879 | 0.289 | High |
| TCGA-ED-A66Y-01A-11R-A311-07 | 283.584 | 17.857 | 0.166 | Low |
| TCGA-DD-A11B-01A-11R-A131-07 | 287.640 | 23.626 | 0.342 | High |
| TCGA-BC-A10X-01A-11R-A131-07 | 259.259 | 0.000 | 0.047 | Low |
| TCGA-DD-AAVW-01A-11R-A41C-07 | 223.770 | 30.769 | 0.342 | High |
| TCGA-ED-A7XP-01A-11R-A352-07 | 274.684 | 28.571 | 0.371 | High |
| TCGA-2Y-A9H9-01A-21R-A39D-07 | 243.947 | 25.549 | 0.248 | High |
| TCGA-2Y-A9H2-01A-12R-A38B-07 | 255.452 | 68.956 | 0.309 | High |
| TCGA-DD-A1EI-01A-11R-A131-07 | 238.839 | 25.549 | 0.356 | High |
| TCGA-CC-A7IJ-01A-11R-A33R-07 | 252.727 | 25.824 | 0.176 | Low |
| TCGA-DD-AADN-01A-11R-A41C-07 | 261.727 | 41.484 | 0.194 | Low |
| TCGA-CC-A3MB-01A-11R-A213-07 | 249.434 | 48.077 | 0.309 | High |
| TCGA-ED-A7PY-01A-11R-A33R-07 | 284.925 | 39.286 | 0.307 | High |
| TCGA-DD-AA3A-01A-11R-A37K-07 | 284.291 | 34.066 | 0.273 | High |
| TCGA-DD-AAD3-01A-11R-A41C-07 | 244.863 | 23.626 | 0.342 | High |
| TCGA-FV-A2QQ-01A-11R-A22L-07 | 244.218 | 11.538 | 0.251 | High |
| TCGA-5R-AAAM-01A-12R-A41C-07 | 203.303 | 7.967 | 0.199 | Low |
| TCGA-ED-A66X-01A-11R-A311-07 | 266.578 | 27.198 | 0.305 | High |
| TCGA-G3-A25U-01A-11R-A16W-07 | 267.606 | 22.527 | 0.306 | High |
| TCGA-DD-AACJ-01A-11R-A41C-07 | 277.233 | 26.374 | 0.288 | High |
| TCGA-UB-AA0V-01A-11R-A38B-07 | 220.563 | 0.000 | 0.008 | Low |
| TCGA-G3-A5SJ-01A-11R-A27V-07 | 255.690 | 23.901 | 0.311 | High |
| TCGA-UB-A7MB-01A-11R-A33R-07 | 263.841 | 19.231 | 0.282 | High |
| TCGA-DD-A3A5-01A-11R-A22L-07 | 281.575 | 22.253 | 0.234 | High |
| TCGA-UB-A7MA-01A-11R-A33R-07 | 280.343 | 53.022 | 0.342 | High |
| TCGA-WQ-A9G7-01A-11R-A37K-07 | 279.912 | 43.681 | 0.243 | High |

| | | | | |
|---|---|---|---|---|
| TCGA-WQ-AB4B-01A-11R-A41C-07 | 273.634 | 31.593 | 0.231 | High |
| TCGA-G3-A7M5-01A-11R-A33R-07 | 261.025 | 18.681 | 0.263 | High |
| TCGA-BC-A10T-01A-11R-A131-07 | 250.639 | 28.297 | 0.301 | High |
| TCGA-O8-A75V-01A-11R-A32O-07 | 249.770 | 31.868 | 0.344 | High |
| TCGA-DD-AADY-01A-11R-A41C-07 | 308.982 | 95.055 | 0.333 | High |
| TCGA-K7-A5RG-01A-11R-A28V-07 | 248.873 | 0.000 | 0.283 | High |
| TCGA-2Y-A9H3-01A-11R-A38B-07 | 274.932 | 20.330 | 0.193 | Low |
| TCGA-DD-AAEI-01A-11R-A41C-07 | 239.036 | 18.681 | 0.218 | High |
| TCGA-3K-AAZ8-01A-12R-A39D-07 | 303.124 | 11.813 | 0.287 | High |
| TCGA-DD-AAVZ-01A-11R-A41C-07 | 279.609 | 24.725 | 0.374 | High |
| TCGA-CC-A7IK-01A-12R-A33R-07 | 302.725 | 59.615 | 0.303 | High |
| TCGA-K7-A5RF-01A-11R-A28V-07 | 229.443 | 0.000 | 0.086 | Low |
| TCGA-BC-A10W-01A-11R-A131-07 | 261.296 | 29.945 | 0.358 | High |
| TCGA-DD-AADS-01A-11R-A41C-07 | 267.493 | 10.989 | 0.143 | Low |
| TCGA-G3-A3CI-01A-11R-A213-07 | 208.805 | 0.549 | 0.059 | Low |
| TCGA-G3-A7M6-01A-11R-A33R-07 | 281.204 | 27.198 | 0.301 | High |
| TCGA-G3-A3CG-01A-11R-A213-07 | 240.411 | 18.407 | 0.217 | High |
| TCGA-DD-A73A-01A-12R-A32O-07 | 229.537 | 8.791 | 0.230 | High |
| TCGA-G3-A25V-01A-11R-A16W-07 | 223.624 | 0.275 | 0.086 | Low |
| TCGA-DD-A4ND-01A-11R-A266-07 | 263.787 | 42.033 | 0.320 | High |
| TCGA-2Y-A9GZ-01A-11R-A39D-07 | 276.307 | 48.352 | 0.285 | High |
| TCGA-DD-A73G-01A-22R-A32O-07 | 305.272 | 19.231 | 0.193 | Low |
| TCGA-CC-A7IE-01A-21R-A38B-07 | 267.851 | 26.374 | 0.249 | High |
| TCGA-DD-AAVS-01A-11R-A41C-07 | 264.074 | 34.615 | 0.379 | High |
| TCGA-DD-A73C-01A-12R-A33J-07 | 265.055 | 7.143 | 0.111 | Low |
| TCGA-DD-A3A2-01A-11R-A213-07 | 257.524 | 7.143 | 0.100 | Low |
| TCGA-DD-AADK-01A-11R-A41C-07 | 267.315 | 22.802 | 0.363 | High |
| TCGA-DD-A1EE-01A-11R-A131-07 | 268.539 | 15.934 | 0.262 | High |
| TCGA-ZS-A9CF-01A-11R-A38B-07 | 263.806 | 26.099 | 0.306 | High |
| TCGA-G3-AAV1-01A-11R-A38B-07 | 258.116 | 46.703 | 0.307 | High |
| TCGA-XR-A8TE-01A-11R-A36F-07 | 270.509 | 21.429 | 0.143 | Low |
| TCGA-DD-AAE1-01A-11R-A41C-07 | 291.615 | 27.747 | 0.326 | High |
| TCGA-DD-A115-01A-11R-A131-07 | 257.596 | 26.099 | 0.301 | High |
| TCGA-DD-A1EB-01A-11R-A131-07 | 247.446 | 20.055 | 0.179 | Low |
| TCGA-ED-A627-01A-12R-A311-07 | 249.484 | 0.275 | 0.000 | Low |
| TCGA-DD-AACF-01A-11R-A41C-07 | 258.093 | 27.198 | 0.290 | High |
| TCGA-DD-AACX-01A-11R-A41C-07 | 254.264 | 17.033 | 0.218 | High |
| TCGA-DD-AACC-01A-11R-A41C-07 | 255.230 | 21.703 | 0.235 | High |
| TCGA-DD-AACZ-01A-11R-A41C-07 | 238.200 | 53.297 | 0.267 | High |
| TCGA-CC-A9FW-01A-11R-A37K-07 | 258.823 | 73.626 | 0.336 | High |
| TCGA-2Y-A9GV-01A-11R-A38B-07 | 244.556 | 10.165 | 0.234 | High |

| | | | | |
|---|---|---|---|---|
| TCGA-5C-A9VG-01A-11R-A37K-07 | 275.033 | 20.330 | 0.196 | Low |
| TCGA-DD-AAVR-01A-11R-A41C-07 | 230.023 | 19.231 | 0.226 | High |
| TCGA-BC-A8YO-01A-11R-A37K-07 | 244.285 | 16.484 | 0.282 | High |
| TCGA-DD-AAEA-01A-11R-A41C-07 | 271.162 | 28.846 | 0.268 | High |
| TCGA-QA-A7B7-01A-11R-A32O-07 | 280.574 | 27.747 | 0.305 | High |
| TCGA-DD-A4NS-01A-11R-A311-07 | 243.184 | 0.000 | 0.155 | Low |
| TCGA-HP-A5MZ-01A-21R-A27V-07 | 255.449 | 7.692 | 0.269 | High |
| TCGA-MR-A8JO-01A-12R-A36F-07 | 233.384 | 8.516 | 0.175 | Low |
| TCGA-2Y-A9GX-01A-11R-A38B-07 | 255.646 | 6.593 | 0.183 | Low |
| TCGA-DD-A73E-01A-12R-A32O-07 | 259.853 | 23.077 | 0.258 | High |
| TCGA-DD-AADQ-01A-11R-A41C-07 | 258.256 | 12.912 | 0.279 | High |
| TCGA-DD-A1EJ-01A-11R-A155-07 | 269.060 | 44.505 | 0.290 | High |
| TCGA-KR-A7K7-01A-11R-A33J-07 | 267.241 | 57.692 | 0.245 | High |
| TCGA-DD-AACY-01A-11R-A41C-07 | 275.482 | 19.780 | 0.308 | High |
| TCGA-UB-A7ME-01A-11R-A33J-07 | 283.236 | 36.264 | 0.336 | High |
| TCGA-G3-A25Y-01A-11R-A16W-07 | 266.442 | 29.670 | 0.312 | High |
| TCGA-DD-A4NV-01A-11R-A311-07 | 244.035 | 12.363 | 0.298 | High |
| TCGA-DD-A1EG-01A-11R-A213-07 | 236.043 | 32.418 | 0.301 | High |
| TCGA-ED-A7PX-01A-51R-A352-07 | 254.024 | 34.341 | 0.231 | High |
| TCGA-CC-A7IF-01A-11R-A33J-07 | 276.199 | 34.066 | 0.306 | High |
| TCGA-G3-AAV0-01A-11R-A37K-07 | 275.941 | 17.308 | 0.274 | High |
| TCGA-G3-A5SM-01A-12R-A28V-07 | 231.490 | 26.099 | 0.270 | High |
| TCGA-DD-AAED-01A-12R-A41C-07 | 283.887 | 40.385 | 0.308 | High |
| TCGA-5R-AA1D-01A-11R-A38B-07 | 217.686 | 5.769 | 0.081 | Low |
| TCGA-DD-AACV-01A-11R-A41C-07 | 311.248 | 34.341 | 0.305 | High |
| TCGA-CC-A5UD-01A-11R-A28V-07 | 254.674 | 89.560 | 0.216 | High |
| TCGA-CC-A9FS-01A-11R-A37K-07 | 290.146 | 25.549 | 0.369 | High |
| TCGA-DD-AAVQ-01A-11R-A41C-07 | 276.037 | 17.033 | 0.290 | High |
| TCGA-RC-A7SB-01A-21R-A352-07 | 257.526 | 20.604 | 0.308 | High |
| TCGA-BW-A5NP-01A-11R-A27V-07 | 310.660 | 54.396 | 0.360 | High |
| TCGA-DD-AADJ-01A-11R-A41C-07 | 250.854 | 18.132 | 0.245 | High |
| TCGA-DD-AACN-01A-11R-A41C-07 | 315.793 | 35.165 | 0.372 | High |
| TCGA-DD-AAD1-01A-11R-A41C-07 | 232.528 | 12.637 | 0.237 | High |
| TCGA-BC-A69H-01A-11R-A311-07 | 250.696 | 56.044 | 0.257 | High |
| TCGA-BC-A10R-01A-11R-A131-07 | 267.189 | 67.033 | 0.323 | High |
| TCGA-ZS-A9CE-01A-11R-A37K-07 | 286.428 | 29.396 | 0.237 | High |
| TCGA-DD-AAD0-01A-11R-A41C-07 | 241.656 | 17.033 | 0.232 | High |
| TCGA-ZP-A9D4-01A-11R-A37K-07 | 259.397 | 17.033 | 0.225 | High |
| TCGA-4R-AA8I-01A-11R-A38B-07 | 255.103 | 9.066 | 0.296 | High |
| TCGA-FV-A3R3-01A-11R-A22L-07 | 217.858 | 6.868 | 0.125 | Low |
| TCGA-LG-A6GG-01A-11R-A311-07 | 280.364 | 48.626 | 0.313 | High |

| | | | | |
|---|---|---|---|---|
| TCGA-DD-AADO-01A-11R-A41C-07 | 292.296 | 43.407 | 0.269 | High |
| TCGA-2Y-A9H1-01A-11R-A38B-07 | 273.908 | 18.132 | 0.218 | High |
| TCGA-BC-A10Y-01A-11R-A131-07 | 260.666 | 26.923 | 0.247 | High |
| TCGA-G3-A5SI-01A-31R-A27V-07 | 281.584 | 28.846 | 0.293 | High |
| TCGA-CC-A5UE-01A-11R-A28V-07 | 280.305 | 45.604 | 0.321 | High |
| TCGA-DD-AADU-01A-11R-A41C-07 | 267.753 | 83.791 | 0.244 | High |
| TCGA-DD-AAW3-01A-11R-A41C-07 | 298.783 | 21.154 | 0.327 | High |
| TCGA-G3-AAV5-01A-11R-A37K-07 | 269.221 | 31.593 | 0.233 | High |
| TCGA-CC-A7IL-01A-11R-A33R-07 | 304.318 | 50.000 | 0.320 | High |
| TCGA-DD-AAE4-01A-11R-A41C-07 | 262.484 | 15.934 | 0.167 | Low |
| TCGA-2Y-A9GT-01A-11R-A38B-07 | 220.526 | 1.923 | 0.172 | Low |
| TCGA-DD-AACH-01A-11R-A41C-07 | 243.067 | 34.341 | 0.326 | High |
| TCGA-EP-A2KA-01A-11R-A180-07 | 236.229 | 23.901 | 0.222 | High |
| TCGA-DD-AADI-01A-11R-A41C-07 | 271.844 | 26.099 | 0.273 | High |
| TCGA-RC-A6M4-01A-11R-A32O-07 | 265.721 | 39.560 | 0.273 | High |
| TCGA-RC-A6M3-01A-11R-A32O-07 | 270.870 | 48.626 | 0.203 | High |
| TCGA-DD-A3A9-01A-11R-A266-07 | 283.756 | 65.385 | 0.266 | High |
| TCGA-G3-A25T-01A-11R-A16W-07 | 285.980 | 14.286 | 0.195 | Low |
| TCGA-ES-A2HT-01A-12R-A180-07 | 261.613 | 11.538 | 0.186 | Low |
| TCGA-ES-A2HS-01A-11R-A180-07 | 311.565 | 32.143 | 0.251 | High |
| TCGA-KR-A7K0-01A-12R-A33R-07 | 251.401 | 5.220 | 0.190 | Low |
| TCGA-K7-AAU7-01A-11R-A38B-07 | 222.761 | 14.560 | 0.252 | High |
| TCGA-DD-A73F-01A-11R-A32O-07 | 240.798 | 10.165 | 0.208 | High |
| TCGA-G3-AAV6-01A-21R-A37K-07 | 287.045 | 70.330 | 0.293 | High |
| TCGA-DD-AACL-01A-11R-A41C-07 | 252.838 | 29.396 | 0.244 | High |
| TCGA-XR-A8TD-01A-12R-A39D-07 | 249.095 | 27.747 | 0.245 | High |
| TCGA-DD-A11C-01A-11R-A131-07 | 244.326 | 51.923 | 0.264 | High |
| TCGA-MR-A520-01A-11R-A266-07 | 250.506 | 0.000 | -0.016 | Low |
| TCGA-DD-AAVU-01A-11R-A41C-07 | 282.965 | 87.637 | 0.298 | High |
| TCGA-2Y-A9GY-01A-11R-A38B-07 | 262.723 | 42.308 | 0.279 | High |
| TCGA-DD-A39V-01A-11R-A213-07 | 241.350 | 25.549 | 0.233 | High |
| TCGA-BC-A112-01A-11R-A131-07 | 221.344 | 28.571 | 0.272 | High |
| TCGA-ZP-A9CY-01A-11R-A38B-07 | 249.760 | 3.022 | 0.142 | Low |
| TCGA-CC-A3M9-01A-11R-A213-07 | 284.039 | 28.022 | 0.115 | Low |
| TCGA-2Y-A9HB-01A-11R-A39D-07 | 254.293 | 17.308 | 0.232 | High |
| TCGA-DD-AADR-01A-11R-A41C-07 | 241.027 | 31.319 | 0.287 | High |
| TCGA-G3-AAUZ-01A-11R-A38B-07 | 271.209 | 23.626 | 0.221 | High |
| TCGA-MI-A75C-01A-11R-A32O-07 | 282.129 | 16.484 | 0.220 | High |

**Table S1c. Table for PCC of LCI cohort**

| Sample ID | PCC | Group |
|---|---|---|
| LCS_004A | -0.0168 | Low |
| LCS_007A | 0.0477 | Low |
| LCS_023A | 0.1302 | High |
| LCS_067A | 0.1584 | High |
| LCS_065A | 0.1331 | High |
| LCS_073A | 0.0779 | Low |
| LCS_024A | 0.1620 | High |
| LCS_042A | 0.2200 | High |
| LCS_062A | 0.1259 | High |
| LCS_015A | 0.1077 | High |
| LCS_069A | 0.1808 | High |
| LCS_033A | 0.0652 | Low |
| LCS_061A | 0.1397 | High |
| LCS_020A | 0.0613 | Low |
| LCS_045A | 0.1535 | High |
| LCS_057A | 0.1688 | High |
| LCS_009A | 0.2959 | High |
| LCS_049A | 0.0640 | Low |
| LCS_047A | 0.2006 | High |
| LCS_075A | 0.0594 | Low |
| LCS_034A | 0.0456 | Low |
| LCS_041A | 0.2258 | High |
| LCS_044A | -0.0030 | Low |
| LCS_071A | 0.1441 | High |
| LCS_036A | 0.1864 | High |
| LCS_064A | 0.2237 | High |
| LCS_038A | 0.1564 | High |
| LCS_002A | 0.0885 | Low |
| LCS_054A | 0.1344 | High |
| LCS_063A | 0.1109 | High |
| LCS_022A | 0.1863 | High |
| LCS_025A | 0.0539 | Low |
| LCS_066A | 0.1591 | High |
| LCS_027A | 0.1703 | High |
| LCS_048A | 0.0992 | Low |
| LCS_012A | 0.1333 | High |
| LCS_051A | 0.1051 | High |

| | | |
|---|---|---|
| LCS_056A | 0.1909 | High |
| LCS_028A | 0.1985 | High |
| LCS_016A | 0.1540 | High |
| LCS_196A | 0.1571 | High |
| LCS_072A | 0.1593 | High |
| LCS_357A | 0.1484 | High |
| LCS_005A | 0.1554 | High |
| LCS_019A | 0.1922 | High |
| LCS_029A | 0.1017 | High |
| LCS_039A | 0.0971 | Low |
| LCS_254A | 0.0431 | Low |
| LCS_068A | 0.0990 | Low |
| LCS_076A | 0.3269 | High |
| LCS_333A | 0.2193 | High |
| LCS_339A | 0.0606 | Low |
| LCS_341A | 0.0750 | Low |
| LCS_343A | 0.1621 | High |
| LCS_344A | 0.1981 | High |
| LCS_346A | 0.1155 | High |
| LCS_347A | 0.2424 | High |
| LCS_393A | 0.1158 | High |
| LCS_400A | 0.2249 | High |
| LCS_401A | 0.1066 | High |
| LCS_403A | 0.1890 | High |
| LCS_406A | 0.1228 | High |
| LCS_415A | 0.1272 | High |
| LCS_424A | 0.2341 | High |
| LCS_426A | 0.1391 | High |

**Table S2. Overlapped Differentially Expressed Genes among correlated genes between Thai HCC and Thai iCCA**

| | Probe ID | Official Symbol | HCC | | | | | | iCCA | | | | | |
|---|---|---|---|---|---|---|---|---|---|---|---|---|---|---|
| | | | LFGC (Mean) | HFGC (Mean) | perm.p | Fold Change | cor.r | cor.p | LFGC (Mean) | HFGC (Mean) | perm.p | Fold Change | cor.r | cor.p |
| HFGC_UP | TC17001769.hg | BRIP1 | 0.0142 | 1.1001 | 0.0017 | 1.0860 | 0.5851 | 0.0000 | 0.2369 | 0.9013 | 0.0024 | 0.6645 | 0.4350 | 0.0000 |
| | TC05000134.hg | C5orf22 | 0.0416 | 0.6175 | 0.0018 | 0.5759 | 0.5762 | 0.0000 | -0.0638 | 0.6110 | 0.0000 | 0.6748 | 0.5907 | 0.0000 |
| | TC03001882.hg | HLTF | 0.1843 | 1.0937 | 0.0028 | 0.9094 | 0.5510 | 0.0000 | -0.2542 | 0.5102 | 0.0015 | 0.7644 | 0.4663 | 0.0000 |
| | TC08000701.hg | MTBP | 0.0354 | 0.8016 | 0.0004 | 0.7662 | 0.5097 | 0.0000 | 0.1454 | 0.8641 | 0.0001 | 0.7187 | 0.6017 | 0.0000 |
| | TC01001781.hg | FLVCR1 | 0.4805 | 1.4848 | 0.0008 | 1.0043 | 0.5002 | 0.0000 | 0.3281 | 1.0638 | 0.0002 | 0.7357 | 0.4285 | 0.0000 |
| | TC10001914.hg | - | 0.1514 | 0.7716 | 0.0039 | 0.6202 | 0.4810 | 0.0001 | -0.0738 | 0.5058 | 0.0001 | 0.5796 | 0.4807 | 0.0000 |
| | TC19001630.hg | - | -0.2373 | 0.6659 | 0.0039 | 0.9032 | 0.4580 | 0.0002 | -0.1117 | 0.6015 | 0.0003 | 0.7133 | 0.3103 | 0.0029 |
| | TC01005322.hg | - | -0.2565 | 0.3482 | 0.0031 | 0.6046 | 0.4102 | 0.0009 | -0.1403 | 0.5285 | 0.0014 | 0.6688 | 0.6045 | 0.0000 |
| | TC20000063.hg | MCM8 | 0.0263 | 0.9355 | 0.0043 | 0.9093 | 0.3707 | 0.0030 | 0.4325 | 1.5114 | 0.0000 | 1.0789 | 0.5561 | 0.0000 |
| | TC07001654.hg | MCM7 | -0.0801 | 0.6606 | 0.0006 | 0.7408 | 0.3695 | 0.0031 | 0.1244 | 0.9751 | 0.0000 | 0.8507 | 0.5547 | 0.0000 |
| | TC05000161.hg | SKP2 | -0.0199 | 1.0414 | 0.0009 | 1.0614 | 0.3537 | 0.0048 | -0.8532 | 0.1413 | 0.0000 | 0.9945 | 0.5339 | 0.0000 |
| | TC09002185.hg | - | -0.1209 | 0.7151 | 0.0039 | 0.8360 | 0.3309 | 0.0086 | -0.5622 | -0.0146 | 0.0003 | 0.5477 | 0.3671 | 0.0004 |
| | TC01002916.hg | SASS6 | -0.0896 | 0.6298 | 0.0030 | 0.7194 | 0.3192 | 0.0114 | 0.3593 | 1.0462 | 0.0001 | 0.6868 | 0.3587 | 0.0005 |
| | TC20000784.hg | PIGU | 0.2316 | 1.0044 | 0.0015 | 0.7728 | 0.2949 | 0.0200 | 0.1652 | 0.8425 | 0.0000 | 0.6773 | 0.6154 | 0.0000 |
| | TC08000695.hg | MAL2 | -0.9544 | 0.2122 | 0.0025 | 1.1665 | 0.2941 | 0.0203 | 0.6231 | 1.6283 | 0.0001 | 1.0052 | 0.5509 | 0.0000 |
| | TC17002618.hg | - | 0.1968 | 1.9895 | 0.0049 | 1.7927 | 0.2895 | 0.0225 | 1.4700 | 3.0272 | 0.0000 | 1.5572 | 0.3490 | 0.0007 |
| | TC12001605.hg | TIMELESS | 0.0507 | 0.8189 | 0.0017 | 0.7682 | 0.2763 | 0.0297 | -0.3542 | 0.3319 | 0.0002 | 0.6860 | 0.5740 | 0.0000 |
| HFGC_DN | TC16000443.hg | CYLD | 0.1088 | -0.4176 | 0.0008 | -0.5264 | 0.6379 | 0.0000 | 0.1743 | -0.5150 | 0.0000 | -0.6893 | 0.3223 | 0.0019 |
| | TC04001323.hg | CNOT6L | 0.3452 | -0.3297 | 0.0003 | -0.6749 | 0.6173 | 0.0000 | 0.1911 | -0.3290 | 0.0002 | -0.5201 | 0.4735 | 0.0000 |
| | TC05001570.hg | CCNH | 0.0704 | -0.5432 | 0.0003 | -0.6136 | 0.4298 | 0.0005 | 0.0399 | -0.5382 | 0.0000 | -0.5782 | 0.6222 | 0.0000 |
| | TC09000035.hg | JAK2 | 0.1345 | -0.5793 | 0.0010 | -0.7138 | 0.4244 | 0.0006 | 0.9534 | 0.2215 | 0.0012 | -0.7319 | 0.3768 | 0.0003 |
| | TC05001367.hg | LOC100652914 | 0.2876 | -0.5078 | 0.0045 | -0.7954 | 0.3987 | 0.0013 | -0.8985 | -1.5924 | 0.0001 | -0.6939 | 0.2842 | 0.0066 |
| | TC09000037.hg | CD274 | 0.3362 | -0.3846 | 0.0012 | -0.7208 | 0.3534 | 0.0048 | 0.3626 | -0.3030 | 0.0011 | -0.6655 | 0.3748 | 0.0003 |
| | TC09001325.hg | NFIL3 | -0.0454 | -0.8257 | 0.0034 | -0.7803 | 0.3426 | 0.0064 | -0.6668 | -1.3746 | 0.0003 | -0.7078 | 0.2388 | 0.0234 |
| | TC11002246.hg | CASP4 | 0.1596 | -0.5252 | 0.0007 | -0.6848 | 0.3013 | 0.0173 | 0.3340 | -0.1859 | 0.0008 | -0.5199 | 0.4825 | 0.0000 |
| | TC17002849.hg | PER1 | 0.0407 | -0.6408 | 0.0010 | -0.6814 | 0.2963 | 0.0194 | -0.3776 | -0.9433 | 0.0001 | -0.5657 | 0.2691 | 0.0103 |
| | TC17001053.hg | SCIMP | 0.2858 | -0.6687 | 0.0002 | -0.9545 | 0.2812 | 0.0269 | -0.0821 | -0.6832 | 0.0009 | -0.6011 | 0.3166 | 0.0024 |
| | TC08000310.hg | C8orf4 | -0.4859 | -1.8605 | 0.0033 | -1.3745 | 0.2598 | 0.0414 | -0.0326 | -1.2503 | 0.0001 | -1.2176 | 0.2699 | 0.0101 |
| | TC07001295.hg | TRGV10 | 0.3831 | -0.3456 | 0.0012 | -0.7288 | -0.2387 | 0.0617 | 0.2736 | -0.5858 | 0.0000 | -0.8594 | -0.2417 | 0.0217 |

**Table S3. Comparison table of overlapped differentially expressed genes among correlated genes.**

| Comparison | Overlapped gene | |
|---|---|---|
| | **HFGC_UP** | **HFGC_DN** |
| **Thai HCC vs. Thai iCCA** | FLVCR1, SASS6, HLTF, C5orf22, SKP2, MCM7, MAL2, MTBP, TIMELESS, BRIP1, PIGU | CNOT6L, C8orf4, JAK2, NFIL3, CASP4, CYLD, SCIMP |
| **Thai HCC vs. TCGA_HCC** | ANXA9, ASPM, BRIP1, BRPF3, CAP2, CENPQ, FDPS, FLVCR1, INTS7, MAL2, MCM3, MCM7, MOGAT3, MTBP, PRIM2, SASS6, SQLE, STXBP6, TIMELESS, VPS72, ZNF572 | ALPK1, AOAH, SCIMP, CHD9, COTL1, EPSTI1, FYB, HLA-DOA, IRF8, NDEL1, NFIL3, PPM1K, RGS2, SLAMF6, ZC3H13 |
| **Thai iCCA vs. TCGA_HCC** | ACTL6A, ANKRD27, ATAD2, ATAD5, AURKA, BRCA1, BRIP1, PARPBP, BORA, CASC5, CEP85, CCT3, CDC20, CDC45, CDC6, CDCA8, CENPA, CENPO, CKS2, DNA2, DSCC1, DSN1, E2F7, ECT2, EFNA4, EZH2, FAM83D, FANCD2, FANCI, FLVCR1, FOXM1, GDPD1, GINS1, GPSM2, IGF2BP3, INTS8, IQGAP3, KIF15, KIF20B, KIF2C, KIFC1, KNTC1, KPNA2, LAPTM4B, LLGL2, LSR, MAL2, MCM4, MCM7, MELK, MSH2, MTBP, MYBL2, NCAPG2, NDC80, NEK2, NUP155, PBK, PCNA, POLQ, PRIM1, PSPH, RACGAP1, RAD51AP1, RBL1, RFC3, RFC4, RRM2, SASS6, SHCBP1, SKA1, TIMELESS, TKT, TOP2A, TRIP13, TUBA1B, TUBG1, TYMS, UBE2C, UBE2T, XPO5, XRCC2, ZNF623 | AIF1, SCIMP, CDC37L1, CLEC10A, CSGALNACT1, CTSO, DOCK8, ETS1, GAS7, IL7R, JUNB, NAT2, NFIL3, PDCD1LG2, PECAM1, PMP22, RCAN1 |
| **Thai HCC vs. Thai iCCA vs. TCGA HCC** | BRIP1, FLVCR1, MAL2, MCM7, MTBP, SASS6, TIMELESS | SCIMP, NFIL3 |

### Table S4. GSEA result compared HFGC with LFGC of Thai iCCA

| Biological Process (HFGC) | | | | | | | | |
|---|---|---|---|---|---|---|---|---|
| NAME | SIZE | ES | NES | NOM p-val | FDR q-val | FWER p-val | RANK AT MAX | LEADING EDGE |
| DNA_DEPENDENT_DNA_REPLICATION | 43 | -0.674 | -1.907 | 0.002 | 0.2221 | 0.122 | 2601 | tags=60%, list=19%, signal=75% |
| MITOSIS | 50 | -0.689 | -1.882 | 0.002 | 0.1576 | 0.166 | 1564 | tags=52%, list=12%, signal=59% |
| M_PHASE_OF_MITOTIC_CELL_CYCLE | 53 | -0.685 | -1.861 | 0.002 | 0.1475 | 0.204 | 1564 | tags=51%, list=12%, signal=57% |
| M_PHASE | 70 | -0.664 | -1.855 | 0.004 | 0.1176 | 0.21 | 1855 | tags=53%, list=14%, signal=61% |
| CELL_CYCLE_PHASE | 107 | -0.593 | -1.837 | 0.0079 | 0.1166 | 0.242 | 1564 | tags=42%, list=12%, signal=47% |
| CELL_CYCLE_PROCESS | 124 | -0.601 | -1.834 | 0.0041 | 0.1008 | 0.249 | 1957 | tags=47%, list=15%, signal=54% |
| MITOTIC_CELL_CYCLE | 99 | -0.604 | -1.82 | 0.0081 | 0.1015 | 0.277 | 1766 | tags=43%, list=13%, signal=50% |
| DNA_REPLICATION | 75 | -0.599 | -1.804 | 0.008 | 0.1073 | 0.307 | 2620 | tags=52%, list=19%, signal=64% |
| CHROMOSOME_SEGREGATION | 21 | -0.702 | -1.801 | 0.0039 | 0.0981 | 0.309 | 1024 | tags=48%, list=8%, signal=51% |
| CELL_CYCLE_GO_0007049 | 215 | -0.505 | -1.799 | 0.0101 | 0.0907 | 0.313 | 2042 | tags=38%, list=15%, signal=44% |
| DNA_INTEGRITY_CHECKPOINT | 16 | -0.719 | -1.785 | 0.012 | 0.0955 | 0.34 | 2579 | tags=63%, list=19%, signal=77% |
| REGULATION_OF_MITOSIS | 26 | -0.682 | -1.784 | 0.008 | 0.0885 | 0.342 | 1564 | tags=54%, list=12%, signal=61% |
| MICROTUBULE_CYTOSKELETON_ORGANIZATION_AND_BIOGENESIS | 28 | -0.632 | -1.747 | 0.0251 | 0.1188 | 0.431 | 3152 | tags=61%, list=23%, signal=79% |
| MICROTUBULE_BASED_PROCESS | 60 | -0.518 | -1.744 | 0.008 | 0.1132 | 0.437 | 3152 | tags=47%, list=23%, signal=61% |
| CHROMATIN_MODIFICATION | 39 | -0.571 | -1.731 | 0.0078 | 0.1195 | 0.469 | 3230 | tags=56%, list=24%, signal=74% |
| CHROMOSOME_ORGANIZATION_AND_BIOGENESIS | 82 | -0.54 | -1.73 | 0.0138 | 0.1138 | 0.471 | 3230 | tags=54%, list=24%, signal=70% |
| CELL_CYCLE_CHECKPOINT_GO_0000075 | 34 | -0.614 | -1.705 | 0.0356 | 0.1347 | 0.524 | 3112 | tags=59%, list=23%, signal=76% |
| MEIOTIC_CELL_CYCLE | 21 | -0.619 | -1.702 | 0.0149 | 0.1315 | 0.535 | 1855 | tags=52%, list=14%, signal=61% |

| | | | | | | | | |
|---|---|---|---|---|---|---|---|---|
| INTERPHASE_OF_MITOTIC_CELL_CYCLE | 37 | -0.564 | -1.686 | 0.0163 | 0.1442 | 0.573 | 4206 | tags=62%, list=31%, signal=90% |
| PHOSPHOINOSITIDE_BIOSYNTHETIC_PROCESS | 20 | -0.642 | -1.685 | 0.0187 | 0.1384 | 0.579 | 2275 | tags=50%, list=17%, signal=60% |
| **Biological Process (LFGC)** | | | | | | | | |
| NAME | SIZE | ES | NES | NOM p-val | FDR q-val | FWER p-val | RANK AT MAX | LEADING EDGE |
| DEFENSE_RESPONSE | 189 | 0.6299 | 1.8836 | 0.0162 | 0.4864 | 0.174 | 2508 | tags=59%, list=19%, signal=71% |
| RESPONSE_TO_WOUNDING | 142 | 0.6262 | 1.8659 | 0.0081 | 0.2967 | 0.204 | 2839 | tags=61%, list=21%, signal=77% |
| INFLAMMATORY_RESPONSE | 100 | 0.6446 | 1.8588 | 0.0142 | 0.2127 | 0.211 | 2508 | tags=61%, list=19%, signal=74% |
| IMMUNE_RESPONSE | 152 | 0.6352 | 1.8528 | 0.0233 | 0.1687 | 0.221 | 2832 | tags=64%, list=21%, signal=80% |
| IMMUNE_SYSTEM_PROCESS | 219 | 0.5803 | 1.8454 | 0.0307 | 0.147 | 0.231 | 2848 | tags=58%, list=21%, signal=72% |
| POSITIVE_REGULATION_OF_MULTICELLULAR_ORGANISMAL_PROCESS | 42 | 0.6503 | 1.8285 | 0.0101 | 0.1507 | 0.263 | 2949 | tags=60%, list=22%, signal=76% |
| RESPONSE_TO_OTHER_ORGANISM | 48 | 0.6462 | 1.8282 | 0.0161 | 0.1298 | 0.263 | 2547 | tags=63%, list=19%, signal=77% |
| CYTOKINE_PRODUCTION | 41 | 0.6209 | 1.7841 | 0.0219 | 0.1792 | 0.362 | 2949 | tags=61%, list=22%, signal=78% |
| POSITIVE_REGULATION_OF_TRANSLATION | 21 | 0.6338 | 1.7774 | 0.014 | 0.1697 | 0.376 | 2190 | tags=62%, list=16%, signal=74% |
| LOCOMOTORY_BEHAVIOR | 74 | 0.6033 | 1.7728 | 0.0164 | 0.1596 | 0.386 | 2820 | tags=64%, list=21%, signal=80% |
| RESPONSE_TO_EXTERNAL_STIMULUS | 213 | 0.5415 | 1.7694 | 0.0164 | 0.1505 | 0.394 | 2848 | tags=55%, list=21%, signal=69% |
| MULTI_ORGANISM_PROCESS | 97 | 0.5187 | 1.7675 | 0.0182 | 0.1408 | 0.394 | 3121 | tags=55%, list=23%, signal=71% |
| POSITIVE_REGULATION_OF_IMMUNE_SYSTEM_PROCESS | 32 | 0.652 | 1.7663 | 0.0217 | 0.1321 | 0.397 | 1006 | tags=41%, list=7%, signal=44% |
| POSITIVE_REGULATION_OF_CYTOKINE_BIOSYNTHETIC_PROCESS | 15 | 0.7064 | 1.7575 | 0.0164 | 0.1328 | 0.423 | 2190 | tags=67%, list=16%, signal=80% |
| CELLULAR_DEFENSE_RESPONSE | 40 | 0.6893 | 1.7565 | 0.0291 | 0.1252 | 0.427 | 1874 | tags=63%, list=14%, signal=72% |
| RESPONSE_TO_BACTERIUM | 15 | 0.7077 | 1.7227 | 0.0163 | 0.1574 | 0.505 | 2488 | tags=67%, list=18%, signal=82% |

| | | | | | | | | |
|---|---|---|---|---|---|---|---|---|
| REGULATION_OF_IMMUNE_SYSTEM_PROCESS | 39 | 0.6153 | 1.7226 | 0.0339 | 0.1484 | 0.505 | 2848 | tags=56%, list=21%, signal=71% |
| POSITIVE_REGULATION_OF_IMMUNE_RESPONSE | 18 | 0.7265 | 1.7208 | 0.0142 | 0.1422 | 0.511 | 1987 | tags=61%, list=15%, signal=72% |
| REGULATION_OF_CYTOKINE_BIOSYNTHETIC_PROCESS | 21 | 0.6296 | 1.7202 | 0.0198 | 0.1351 | 0.513 | 2190 | tags=57%, list=16%, signal=68% |
| RESPONSE_TO_VIRUS | 31 | 0.6363 | 1.7147 | 0.0257 | 0.134 | 0.525 | 3695 | tags=81%, list=27%, signal=111% |
| **KEGG Pathway (HFGC)** | | | | | | | | |
| NAME | SIZE | ES | NES | NOM p-val | FDR q-val | FWER p-val | RANK AT MAX | LEADING EDGE |
| KEGG_DNA_REPLICATION | 29 | -0.812 | -1.86 | 0.0019 | 0.1886 | 0.125 | 969 | tags=72%, list=7%, signal=78% |
| KEGG_MISMATCH_REPAIR | 19 | -0.742 | -1.776 | 0.0059 | 0.2655 | 0.267 | 1688 | tags=63%, list=13%, signal=72% |
| KEGG_BASE_EXCISION_REPAIR | 21 | -0.676 | -1.739 | 0.0077 | 0.2569 | 0.338 | 3359 | tags=71%, list=25%, signal=95% |
| KEGG_CELL_CYCLE | 90 | -0.61 | -1.721 | 0.0211 | 0.2324 | 0.388 | 2306 | tags=52%, list=17%, signal=63% |
| KEGG_HOMOLOGOUS_RECOMBINATION | 16 | -0.71 | -1.689 | 0.0134 | 0.2545 | 0.464 | 1855 | tags=63%, list=14%, signal=72% |
| KEGG_RNA_POLYMERASE | 23 | -0.654 | -1.645 | 0.0059 | 0.3047 | 0.555 | 2754 | tags=65%, list=20%, signal=82% |
| KEGG_OOCYTE_MEIOSIS | 77 | -0.488 | -1.616 | 0.0206 | 0.3265 | 0.618 | 2892 | tags=44%, list=21%, signal=56% |
| KEGG_AMINOACYL_TRNA_BIOSYNTHESIS | 30 | -0.625 | -1.601 | 0.0329 | 0.3195 | 0.648 | 3080 | tags=67%, list=23%, signal=86% |
| KEGG_PYRIMIDINE_METABOLISM | 62 | -0.506 | -1.591 | 0.0476 | 0.308 | 0.675 | 3502 | tags=56%, list=26%, signal=76% |
| KEGG_NUCLEOTIDE_EXCISION_REPAIR | 34 | -0.571 | -1.588 | 0.0355 | 0.2814 | 0.678 | 3395 | tags=50%, list=25%, signal=67% |
| KEGG_N_GLYCAN_BIOSYNTHESIS | 37 | -0.502 | -1.562 | 0.0462 | 0.3011 | 0.729 | 3637 | tags=62%, list=27%, signal=85% |
| KEGG_SPLICEOSOME | 96 | -0.531 | -1.552 | 0.0825 | 0.2975 | 0.747 | 4748 | tags=68%, list=35%, signal=104% |
| KEGG_BASAL_TRANSCRIPTION_FACTORS | 24 | -0.564 | -1.548 | 0.0511 | 0.2811 | 0.754 | 933 | tags=33%, list=7%, signal=36% |
| KEGG_UBIQUITIN_MEDIATED_PROTEOLYSIS | 105 | -0.455 | -1.542 | 0.0405 | 0.2726 | 0.765 | 3469 | tags=46%, list=26%, signal=61% |
| KEGG_RNA_DEGRADATION | 44 | -0.524 | -1.508 | 0.0807 | 0.312 | 0.815 | 3259 | tags=48%, list=24%, |

| | | | | | | | | |
|---|---|---|---|---|---|---|---|---|
| | | | | | | | | signal=63% |
| KEGG_OXIDATIVE_PHOSPHORYLATION | 89 | -0.523 | -1.498 | 0.1 | 0.3092 | 0.826 | 4764 | tags=67%, list=35%, signal=104% |
| KEGG_GLYCOSYLPHOSPHATIDYLINOSITOL_GPI_ANCHOR_BIOSYNTHESIS | 18 | -0.582 | -1.495 | 0.0538 | 0.2962 | 0.829 | 3900 | tags=67%, list=29%, signal=94% |
| KEGG_HUNTINGTONS_DISEASE | 120 | -0.462 | -1.469 | 0.0904 | 0.3207 | 0.854 | 4145 | tags=52%, list=31%, signal=75% |
| KEGG_PROGESTERONE_MEDIATED_OOCYTE_MATURATION | 54 | -0.441 | -1.447 | 0.0696 | 0.3396 | 0.876 | 1134 | tags=28%, list=8%, signal=30% |
| **KEGG Pathway (LFGC)** | | | | | | | | |
| NAME | SIZE | ES | NES | NOM p-val | FDR q-val | FWER p-val | RANK AT MAX | LEADING EDGE |
| KEGG_HEMATOPOIETIC_CELL_LINEAGE | 49 | 0.7264 | 1.8663 | 0.002 | 0.2115 | 0.134 | 1945 | tags=63%, list=14%, signal=74% |
| KEGG_CYTOKINE_CYTOKINE_RECEPTOR_INTERACTION | 156 | 0.5901 | 1.8433 | 0.002 | 0.1416 | 0.167 | 3003 | tags=60%, list=22%, signal=77% |
| KEGG_NATURAL_KILLER_CELL_MEDIATED_CYTOTOXICITY | 80 | 0.5364 | 1.6603 | 0.0455 | 0.5888 | 0.516 | 1202 | tags=40%, list=9%, signal=44% |
| KEGG_NEUROACTIVE_LIGAND_RECEPTOR_INTERACTION | 108 | 0.4393 | 1.6404 | 0.0165 | 0.5135 | 0.557 | 2646 | tags=40%, list=20%, signal=49% |
| KEGG_CELL_ADHESION_MOLECULES_CAMS | 93 | 0.5438 | 1.6371 | 0.052 | 0.4226 | 0.56 | 2500 | tags=49%, list=19%, signal=60% |
| KEGG_CHEMOKINE_SIGNALING_PATHWAY | 142 | 0.458 | 1.5851 | 0.0538 | 0.5215 | 0.668 | 1088 | tags=31%, list=8%, signal=33% |
| KEGG_B_CELL_RECEPTOR_SIGNALING_PATHWAY | 58 | 0.4919 | 1.5708 | 0.0593 | 0.4922 | 0.693 | 1362 | tags=40%, list=10%, signal=44% |
| KEGG_LEUKOCYTE_TRANSENDOTHELIAL_MIGRATION | 76 | 0.4963 | 1.5683 | 0.0641 | 0.4378 | 0.695 | 1611 | tags=38%, list=12%, signal=43% |
| KEGG_T_CELL_RECEPTOR_SIGNALING_PATHWAY | 77 | 0.4747 | 1.567 | 0.0559 | 0.3925 | 0.7 | 1517 | tags=38%, list=11%, signal=42% |
| KEGG_COMPLEMENT_AND_COAGULATION_CASCADES | 59 | 0.734 | 1.5668 | 0.0539 | 0.3536 | 0.701 | 2577 | tags=80%, list=19%, signal=98% |
| KEGG_JAK_STAT_SIGNALING_PATHWAY | 84 | 0.4794 | 1.5541 | 0.0713 | 0.3496 | 0.724 | 2208 | tags=44%, list=16%, signal=52% |
| KEGG_NOD_LIKE_RECEPTOR_SIGNALING_PATHWAY | 48 | 0.5046 | 1.5262 | 0.082 | 0.3787 | 0.756 | 1717 | tags=42%, list=13%, signal=48% |
| KEGG_INTESTINAL_IMMUNE_NETWORK_FOR_IG | 29 | 0.6359 | 1.4996 | 0.1064 | 0.4074 | 0.795 | 4060 | tags=86%, list=30%, |

| | | | | | | | | |
|---|---|---|---|---|---|---|---|---|
| A_PRODUCTION | | | | | | | | signal=123% |
| KEGG_GRAFT_VERSUS_HOST_DISEASE | 25 | 0.7296 | 1.4963 | 0.0682 | 0.3859 | 0.799 | 2435 | tags=76%, list=18%, signal=93% |
| KEGG_PRIMARY_IMMUNODEFICIENCY | 26 | 0.6269 | 1.4957 | 0.1086 | 0.361 | 0.8 | 2588 | tags=73%, list=19%, signal=90% |
| KEGG_AUTOIMMUNE_THYROID_DISEASE | 23 | 0.6945 | 1.4832 | 0.0969 | 0.3616 | 0.817 | 2877 | tags=74%, list=21%, signal=94% |
| KEGG_ARACHIDONIC_ACID_METABOLISM | 32 | 0.5242 | 1.4747 | 0.1006 | 0.3558 | 0.829 | 3686 | tags=69%, list=27%, signal=94% |
| KEGG_TYPE_II_DIABETES_MELLITUS | 31 | 0.4034 | 1.4685 | 0.0393 | 0.3455 | 0.833 | 585 | tags=23%, list=4%, signal=24% |
| KEGG_PRION_DISEASES | 29 | 0.5504 | 1.4596 | 0.092 | 0.3433 | 0.842 | 2577 | tags=59%, list=19%, signal=72% |
| KEGG_NICOTINATE_AND_NICOTINAMIDE_METABOLISM | 17 | 0.4906 | 1.4434 | 0.0658 | 0.3543 | 0.856 | 2290 | tags=35%, list=17%, signal=42% |
| **Oncogenic (HFGC)** | | | | | | | | |
| NAME | SIZE | ES | NES | NOM p-val | FDR q-val | FWER p-val | RANK AT MAX | LEADING EDGE |
| RB_DN.V1_UP | 89 | -0.564 | -1.931 | 0.0039 | 0.0198 | 0.038 | 2798 | tags=52%, list=21%, signal=65% |
| RB_P107_DN.V1_UP | 88 | -0.606 | -1.923 | 0.0059 | 0.0108 | 0.041 | 1688 | tags=51%, list=13%, signal=58% |
| PRC2_EZH2_UP.V1_UP | 117 | -0.475 | -1.778 | 0.0115 | 0.0513 | 0.181 | 1547 | tags=35%, list=11%, signal=39% |
| RPS14_DN.V1_DN | 132 | -0.515 | -1.764 | 0.0213 | 0.0462 | 0.205 | 4011 | tags=59%, list=30%, signal=83% |
| E2F1_UP.V1_UP | 123 | -0.517 | -1.757 | 0.0179 | 0.04 | 0.219 | 3182 | tags=51%, list=24%, signal=66% |
| GCNP_SHH_UP_EARLY.V1_UP | 115 | -0.447 | -1.653 | 0.0189 | 0.0852 | 0.406 | 2257 | tags=37%, list=17%, signal=45% |
| CSR_LATE_UP.V1_UP | 125 | -0.503 | -1.616 | 0.0247 | 0.1003 | 0.478 | 2489 | tags=46%, list=18%, signal=56% |
| RB_P130_DN.V1_UP | 78 | -0.418 | -1.616 | 0.0203 | 0.0879 | 0.478 | 2950 | tags=38%, list=22%, signal=49% |
| PRC2_EDD_UP.V1_UP | 110 | -0.443 | -1.564 | 0.0506 | 0.1202 | 0.581 | 2160 | tags=34%, list=16%, signal=40% |
| HOXA9_DN.V1_DN | 135 | -0.406 | -1.535 | 0.0503 | 0.1341 | 0.629 | 2943 | tags=38%, list=22%, signal=48% |
| VEGF_A_UP.V1_DN | 147 | -0.399 | -1.447 | 0.1016 | 0.2237 | 0.764 | 2360 | tags=33%, list=18%, signal=39% |

| | | | | | | | | |
|---|---|---|---|---|---|---|---|---|
| E2F3_UP.V1_UP | 105 | -0.355 | -1.407 | 0.0911 | 0.258 | 0.81 | 1986 | tags=28%, list=15%, signal=32% |
| SIRNA_EIF4GI_UP | 73 | -0.403 | -1.405 | 0.1023 | 0.2412 | 0.812 | 3063 | tags=45%, list=23%, signal=58% |
| GCNP_SHH_UP_LATE.V1_UP | 128 | -0.371 | -1.398 | 0.1212 | 0.234 | 0.822 | 3118 | tags=39%, list=23%, signal=50% |
| EGFR_UP.V1_DN | 135 | -0.323 | -1.391 | 0.0588 | 0.2267 | 0.826 | 3102 | tags=33%, list=23%, signal=43% |
| TBK1.DN.48HRS_UP | 39 | -0.403 | -1.341 | 0.1044 | 0.2821 | 0.891 | 2892 | tags=38%, list=21%, signal=49% |
| TBK1.DF_DN | 228 | -0.337 | -1.34 | 0.1445 | 0.2661 | 0.891 | 4018 | tags=41%, list=30%, signal=57% |
| MYC_UP.V1_UP | 105 | -0.403 | -1.321 | 0.1899 | 0.2775 | 0.91 | 2375 | tags=33%, list=18%, signal=40% |
| EIF4E_UP | 54 | -0.391 | -1.288 | 0.1908 | 0.3111 | 0.933 | 3506 | tags=46%, list=26%, signal=62% |
| RB_P107_DN.V1_DN | 94 | -0.339 | -1.243 | 0.1879 | 0.3658 | 0.955 | 3218 | tags=36%, list=24%, signal=47% |
| **Oncogenic (LFGC)** | | | | | | | | |
| NAME | SIZE | ES | NES | NOM p-val | FDR q-val | FWER p-val | RANK AT MAX | LEADING EDGE |
| P53_DN.V2_UP | 79 | 0.5704 | 1.9195 | 0 | 0.102 | 0.048 | 2412 | tags=44%, list=18%, signal=54% |
| RPS14_DN.V1_UP | 142 | 0.6025 | 1.8577 | 0.0081 | 0.1009 | 0.09 | 2491 | tags=54%, list=18%, signal=66% |
| EGFR_UP.V1_UP | 152 | 0.5307 | 1.8076 | 0.014 | 0.1238 | 0.143 | 2685 | tags=45%, list=20%, signal=55% |
| P53_DN.V1_DN | 133 | 0.5355 | 1.7705 | 0.0101 | 0.1346 | 0.192 | 2926 | tags=52%, list=22%, signal=66% |
| RELA_DN.V1_UP | 91 | 0.4621 | 1.7161 | 0.0141 | 0.1838 | 0.29 | 2402 | tags=38%, list=18%, signal=46% |
| KRAS.PROSTATE_UP.V1_UP | 71 | 0.5174 | 1.6938 | 0.0145 | 0.1842 | 0.334 | 3189 | tags=58%, list=24%, signal=75% |
| HOXA9_DN.V1_UP | 147 | 0.5021 | 1.6755 | 0.039 | 0.1848 | 0.369 | 1942 | tags=37%, list=14%, signal=42% |
| KRAS.LUNG.BREAST_UP.V1_UP | 88 | 0.4688 | 1.6688 | 0.012 | 0.1709 | 0.379 | 2890 | tags=43%, list=21%, signal=55% |
| PTEN_DN.V2_UP | 94 | 0.4766 | 1.6472 | 0.0261 | 0.1827 | 0.416 | 2132 | tags=40%, list=16%, signal=48% |
| PTEN_DN.V1_DN | 96 | 0.4537 | 1.6429 | 0.020 | 0.171 | 0.424 | 3254 | tags=49%, |

| | | | | | | | | |
|---|---|---|---|---|---|---|---|---|
| | | | | 4 | 4 | | | list=24%, signal=64% |
| ATM_DN.V1_UP | 79 | 0.4708 | 1.6424 | 0.020 4 | 0.156 3 | 0.425 | 3473 | tags=51%, list=26%, signal=68% |
| STK33_NOMO_UP | 227 | 0.4533 | 1.6374 | 0.045 6 | 0.148 7 | 0.435 | 2294 | tags=39%, list=17%, signal=46% |
| STK33_UP | 225 | 0.4324 | 1.6319 | 0.050 4 | 0.144 | 0.445 | 2417 | tags=41%, list=18%, signal=49% |
| KRAS.BREAST_UP.V1_UP | 71 | 0.4614 | 1.6316 | 0.032 8 | 0.134 3 | 0.447 | 3579 | tags=48%, list=27%, signal=65% |
| KRAS.LUNG_UP.V1_UP | 81 | 0.4791 | 1.6239 | 0.020 7 | 0.131 8 | 0.459 | 2890 | tags=47%, list=21%, signal=59% |
| CRX_DN.V1_DN | 86 | 0.4552 | 1.623 | 0.023 5 | 0.124 | 0.46 | 1958 | tags=34%, list=15%, signal=39% |
| HINATA_NFKB_IMMU_IN F | 16 | 0.7641 | 1.6149 | 0.029 6 | 0.124 9 | 0.475 | 2133 | tags=81%, list=16%, signal=96% |
| IL2_UP.V1_DN | 114 | 0.3958 | 1.6109 | 0.006 5 | 0.122 | 0.486 | 2989 | tags=38%, list=22%, signal=48% |
| STK33_SKM_UP | 214 | 0.4085 | 1.6005 | 0.059 3 | 0.125 6 | 0.507 | 2417 | tags=40%, list=18%, signal=48% |
| KRAS.BREAST_UP.V1_DN | 74 | 0.452 | 1.5994 | 0.016 6 | 0.120 3 | 0.508 | 2493 | tags=38%, list=18%, signal=46% |

### Table S5. GSEA result compared HFGC with LFGC of Thai HCC

| **Biological Process (HFGC)** | | | | | | | | |
|---|---|---|---|---|---|---|---|---|
| NAME | SIZE | ES | NES | NOM p-val | FDR q-val | FWER p-val | RANK AT MAX | LEADING EDGE |
| DNA_DEPENDENT_DNA_REPLICATION | 40 | -0.62918 | -1.76602 | 0.022133 | 1 | 0.379 | 2617 | tags=52%, list=19%, signal=65% |
| GOLGI_VESICLE_TRANSPORT | 33 | -0.56013 | -1.71775 | 0.008621 | 0.912368 | 0.504 | 2368 | tags=39%, list=17%, signal=48% |
| CHROMOSOME_SEGREGATION | 21 | -0.63975 | -1.63112 | 0.059305 | 1 | 0.709 | 2362 | tags=62%, list=17%, signal=75% |
| DNA_REPLICATION | 70 | -0.53649 | -1.61436 | 0.056112 | 1 | 0.744 | 2617 | tags=46%, list=19%, signal=56% |
| CHROMATIN_REMODELING | 18 | -0.63872 | -1.61236 | 0.034205 | 0.932113 | 0.745 | 2705 | tags=56%, list=20%, signal=69% |
| REGULATION_OF_DNA_REPLICATION | 15 | -0.61605 | -1.54532 | 0.056112 | 1 | 0.839 | 1571 | tags=47%, list=12%, signal=53% |
| DNA_PACKAGING | 23 | -0.55972 | -1.52803 | 0.079051 | 1 | 0.86 | 2852 | tags=52%, list=21%, signal=66% |
| LIPOPROTEIN_BIOSYNTHETIC_PROCESS | 20 | -0.54358 | -1.52769 | 0.065764 | 1 | 0.861 | 2499 | tags=40%, list=18%, signal=49% |
| PHOSPHOINOSITIDE_METABOLIC_PROCESS | 25 | -0.49369 | -1.47285 | 0.089796 | 1 | 0.905 | 2499 | tags=40%, list=18%, signal=49% |
| SECRETORY_PATHWAY | 53 | -0.42492 | -1.4727 | 0.048583 | 1 | 0.905 | 2368 | tags=36%, list=17%, signal=43% |
| LIPID_BIOSYNTHETIC_PROCESS | 68 | -0.38891 | -1.45206 | 0.069364 | 1 | 0.919 | 2499 | tags=32%, list=18%, signal=39% |
| GLYCEROPHOSPHOLIPID_METABOLIC_PROCESS | 34 | -0.45017 | -1.44896 | 0.086242 | 1 | 0.921 | 2499 | tags=38%, list=18%, signal=47% |
| DNA_METABOLIC_PROCESS | 169 | -0.42659 | -1.44562 | 0.125514 | 1 | 0.924 | 3189 | tags=44%, list=23%, signal=56% |
| DNA_REPAIR | 83 | -0.44888 | -1.43731 | 0.122407 | 1 | 0.93 | 3085 | tags=46%, list=23%, signal=59% |
| PROTEIN_DNA_COMPLEX_ASSEMBLY | 35 | -0.47914 | -1.42807 | 0.124744 | 1 | 0.938 | 2625 | tags=43%, list=19%, signal=53% |
| REGULATION_OF_MITOSIS | 26 | -0.53891 | -1.42042 | 0.149901 | 1 | 0.943 | 2596 | tags=54%, list=19%, signal=66% |
| M_PHASE | 71 | -0.49874 | -1.41633 | 0.174699 | 0.979201 | 0.947 | 3072 | tags=54%, list=23%, signal=69% |
| TRANSCRIPTION_INITIATION | 23 | -0.5176 | -1.41132 | 0.108932 | 0.94973 | 0.95 | 1757 | tags=39%, list=13%, signal=45% |
| DOUBLE_STRAND_BREAK_REPAIR | 18 | -0.50342 | -1.39604 | 0.114345 | 0.978372 | 0.956 | 1558 | tags=39%, list=11%, signal=44% |
| MRNA_PROCESSING_GO_0006397 | 50 | -0.47793 | -1.37971 | 0.153684 | 1 | 0.966 | 2536 | tags=48%, list=19%, signal=59% |
| **Biological Process (LFGC)** | | | | | | | | |
| NAME | SIZE | ES | NES | NOM p-val | FDR q-val | FWER p-val | RANK AT MAX | LEADING EDGE |
| IMMUNE_RESPONSE | 145 | 0.61873 | 1.97102 | 0.004 | 0.09 | 0.063 | 2401 | tags=56%, list=18%, |

| | | | | | | | | |
|---|---|---|---|---|---|---|---|---|
| | | | 4 | 283 | 9877 | | | signal=67% |
| DEFENSE_RESPONSE | 173 | 0.571424 | 1.936295 | 0.008869 | 0.082701 | 0.088 | 2461 | tags=51%, list=18%, signal=61% |
| IMMUNE_SYSTEM_PROCESS | 207 | 0.576359 | 1.8863 | 0.008811 | 0.11421 | 0.152 | 2401 | tags=52%, list=18%, signal=62% |
| CYCLIC_NUCLEOTIDE_MEDIATED_SIGNALING | 43 | 0.538298 | 1.886154 | 0 | 0.08635 | 0.154 | 1277 | tags=40%, list=9%, signal=43% |
| G_PROTEIN_SIGNALING_COUPLED_TO_CYCLIC_NUCLEOTIDE_SECOND_MESSENGER | 42 | 0.540901 | 1.846746 | 0.007444 | 0.112183 | 0.217 | 1277 | tags=40%, list=9%, signal=45% |
| REGULATION_OF_PROTEIN_AMINO_ACID_PHOSPHORYLATION | 18 | 0.65502 | 1.839378 | 0 | 0.10229 | 0.23 | 1422 | tags=44%, list=10%, signal=50% |
| RESPONSE_TO_OTHER_ORGANISM | 37 | 0.633649 | 1.831636 | 0.006637 | 0.09651 | 0.243 | 2458 | tags=57%, list=18%, signal=69% |
| MULTI_ORGANISM_PROCESS | 77 | 0.505827 | 1.830873 | 0.006494 | 0.085133 | 0.246 | 2699 | tags=47%, list=20%, signal=58% |
| CELLULAR_DEFENSE_RESPONSE | 38 | 0.667433 | 1.82391 | 0.013667 | 0.083166 | 0.265 | 1947 | tags=55%, list=14%, signal=64% |
| CAMP_MEDIATED_SIGNALING | 29 | 0.567475 | 1.821902 | 0 | 0.076653 | 0.268 | 2709 | tags=55%, list=20%, signal=69% |
| BEHAVIOR | 84 | 0.527808 | 1.820751 | 0.013699 | 0.070419 | 0.268 | 2395 | tags=46%, list=18%, signal=56% |
| LOCOMOTORY_BEHAVIOR | 64 | 0.595132 | 1.808114 | 0.013393 | 0.073622 | 0.298 | 2395 | tags=56%, list=18%, signal=68% |
| G_PROTEIN_SIGNALING_COUPLED_TO_CAMP_NUCLEOTIDE_SECOND_MESSENGER | 28 | 0.572484 | 1.791727 | 0.002353 | 0.081384 | 0.33 | 2709 | tags=57%, list=20%, signal=71% |
| JAK_STAT_CASCADE | 22 | 0.614424 | 1.790257 | 0.007737 | 0.077138 | 0.332 | 2370 | tags=55%, list=17%, signal=66% |
| POSITIVE_REGULATION_OF_MULTICELLULAR_ORGANISMAL_PROCESS | 39 | 0.55981 | 1.78999 | 0.01073 | 0.072088 | 0.332 | 2401 | tags=49%, list=18%, signal=59% |
| IMMUNE_EFFECTOR_PROCESS | 23 | 0.648251 | 1.78198 | 0.008 | 0.073971 | 0.348 | 1376 | tags=57%, list=10%, signal=63% |
| SECOND_MESSENGER_MEDIATED_SIGNALING | 74 | 0.478572 | 1.759456 | 0.007177 | 0.088508 | 0.403 | 1277 | tags=31%, list=9%, signal=34% |
| INFLAMMATORY_RESPONSE | 90 | 0.536287 | 1.749392 | 0.027837 | 0.09205 | 0.418 | 1608 | tags=39%, list=12%, signal=44% |
| PEPTIDYL_TYROSINE_PHOSPHORYLATION | 19 | 0.662132 | 1.74779 | 0.01227 | 0.088396 | 0.42 | 2370 | tags=47%, list=17%, signal=57% |
| REGULATION_OF_IMMUNE_SYSTEM_PROCESS | 39 | 0.561961 | 1.72683 | 0.02766 | 0.10236 | 0.472 | 2401 | tags=51%, list=18%, signal=62% |
| **KEGG Pathway (HFGC)** | | | | | | | | |
| NAME | SIZE | ES | NES | NOM p-val | FDR q- | FWER p- | RANK | LEADING EDGE |

| | | | | | val | val | AT MA X | |
|---|---|---|---|---|---|---|---|---|
| KEGG_GLYCOSYLPHOS PHATIDYLINOSITOL_G PI_ANCHOR_BIOSYNTH ESIS | 17 | -0.6601 | - 1.68827 | 0.013 699 | 0.91 326 8 | 0.488 | 2499 | tags=53%, list=18%, signal=65% |
| KEGG_MISMATCH_REP AIR | 20 | - 0.67839 | -1.6523 | 0.030 675 | 0.62 350 5 | 0.557 | 3127 | tags=65%, list=23%, signal=84% |
| KEGG_DNA_REPLICATI ON | 30 | - 0.69259 | - 1.58928 | 0.061 475 | 0.68 348 2 | 0.697 | 1140 | tags=53%, list=8%, signal=58% |
| KEGG_N_GLYCAN_BIO SYNTHESIS | 35 | -0.478 | - 1.51036 | 0.062 5 | 0.85 499 5 | 0.814 | 3252 | tags=51%, list=24%, signal=67% |
| KEGG_PROTEIN_EXPO RT | 21 | - 0.54417 | -1.4696 | 0.092 | 0.86 004 1 | 0.86 | 3078 | tags=52%, list=23%, signal=68% |
| KEGG_NUCLEOTIDE_E XCISION_REPAIR | 36 | - 0.49862 | - 1.43636 | 0.128 846 | 0.86 175 9 | 0.895 | 3127 | tags=47%, list=23%, signal=61% |
| KEGG_BASE_EXCISION _REPAIR | 23 | - 0.53646 | - 1.41653 | 0.124 498 | 0.81 724 5 | 0.904 | 3127 | tags=61%, list=23%, signal=79% |
| KEGG_THYROID_CANC ER | 21 | - 0.44684 | - 1.35522 | 0.112 936 | 0.94 311 | 0.941 | 4477 | tags=67%, list=33%, signal=99% |
| KEGG_CELL_CYCLE | 92 | - 0.46083 | - 1.35385 | 0.196 787 | 0.84 353 5 | 0.941 | 1726 | tags=41%, list=13%, signal=47% |
| KEGG_HOMOLOGOUS_ RECOMBINATION | 17 | - 0.56311 | - 1.32721 | 0.182 | 0.84 840 9 | 0.96 | 3127 | tags=47%, list=23%, signal=61% |
| KEGG_SPLICEOSOME | 95 | - 0.45813 | - 1.31074 | 0.216 216 | 0.82 594 9 | 0.968 | 3377 | tags=45%, list=25%, signal=60% |
| KEGG_PYRIMIDINE_ME TABOLISM | 65 | -0.3794 | - 1.22892 | 0.249 012 | 1 | 0.994 | 3208 | tags=38%, list=24%, signal=50% |
| KEGG_AMINOACYL_TR NA_BIOSYNTHESIS | 32 | - 0.44189 | -1.2235 | 0.281 8 | 0.95 994 3 | 0.995 | 4371 | tags=56%, list=32%, signal=83% |
| KEGG_PROGESTERONE _MEDIATED_OOCYTE_ MATURATION | 54 | -0.3868 | -1.2147 | 0.276 423 | 0.91 679 7 | 0.995 | 2871 | tags=46%, list=21%, signal=58% |
| KEGG_BIOSYNTHESIS_ OF_UNSATURATED_FA TTY_ACIDS | 17 | -0.4515 | - 1.20649 | 0.278 21 | 0.87 862 9 | 0.995 | 4881 | tags=71%, list=36%, signal=110% |
| KEGG_BASAL_TRANSC RIPTION_FACTORS | 25 | - 0.40496 | - 1.17575 | 0.268 39 | 0.91 246 6 | 0.998 | 2431 | tags=44%, list=18%, signal=53% |
| KEGG_LYSINE_DEGRA DATION | 30 | - 0.40296 | -1.1726 | 0.278 | 0.86 696 3 | 0.998 | 3032 | tags=40%, list=22%, signal=51% |
| KEGG_RNA_POLYMER ASE | 23 | - 0.44177 | - 1.16567 | 0.305 785 | 0.83 607 6 | 0.998 | 3208 | tags=43%, list=24%, signal=57% |
| KEGG_ADHERENS_JUN CTION | 55 | - 0.33143 | - 1.10704 | 0.324 268 | 0.93 999 8 | 1 | 1869 | tags=27%, list=14%, signal=31% |
| KEGG_OOCYTE_MEIOS IS | 75 | - 0.33657 | - 1.09607 | 0.376 426 | 0.91 905 | 1 | 2886 | tags=35%, list=21%, signal=44% |

| | | | | | | | | |
|---|---|---|---|---|---|---|---|---|
| | | | | | 3 | | | |
| **KEGG Pathway (LFGC)** | | | | | | | | |
| NAME | SIZE | ES | NES | NOM p-val | FDR q-val | FWER p-val | RANK AT MAX | LEADING EDGE |
| KEGG_HEMATOPOIETIC_CELL_LINEAGE | 47 | 0.704545 | 2.007128 | 0.00655 | 0.054984 | 0.027 | 1584 | tags=53%, list=12%, signal=60% |
| KEGG_CYTOKINE_CYTOKINE_RECEPTOR_INTERACTION | 139 | 0.561605 | 1.909341 | 0.009302 | 0.103436 | 0.104 | 2512 | tags=50%, list=18%, signal=60% |
| KEGG_JAK_STAT_SIGNALING_PATHWAY | 81 | 0.548258 | 1.882923 | 0.010438 | 0.094065 | 0.136 | 1689 | tags=42%, list=12%, signal=48% |
| KEGG_INTESTINAL_IMMUNE_NETWORK_FOR_IGA_PRODUCTION | 27 | 0.734203 | 1.830915 | 0.00655 | 0.132395 | 0.201 | 2957 | tags=78%, list=22%, signal=99% |
| KEGG_PRION_DISEASES | 30 | 0.602044 | 1.797722 | 0.008421 | 0.145171 | 0.259 | 2395 | tags=50%, list=18%, signal=61% |
| KEGG_CHEMOKINE_SIGNALING_PATHWAY | 132 | 0.509207 | 1.776465 | 0.029787 | 0.148752 | 0.3 | 1422 | tags=36%, list=10%, signal=39% |
| KEGG_LEISHMANIA_INFECTION | 47 | 0.652137 | 1.775539 | 0.02079 | 0.129828 | 0.305 | 2981 | tags=68%, list=22%, signal=87% |
| KEGG_NATURAL_KILLER_CELL_MEDIATED_CYTOTOXICITY | 75 | 0.567174 | 1.768625 | 0.039014 | 0.12248 | 0.318 | 2461 | tags=49%, list=18%, signal=60% |
| KEGG_CALCIUM_SIGNALING_PATHWAY | 93 | 0.457141 | 1.748602 | 0.009153 | 0.132013 | 0.367 | 3143 | tags=44%, list=23%, signal=57% |
| KEGG_TOLL_LIKE_RECEPTOR_SIGNALING_PATHWAY | 63 | 0.525927 | 1.713769 | 0.030675 | 0.161496 | 0.423 | 1631 | tags=40%, list=12%, signal=45% |
| KEGG_CELL_ADHESION_MOLECULES_CAMS | 94 | 0.515669 | 1.71064 | 0.037946 | 0.151088 | 0.435 | 2493 | tags=48%, list=18%, signal=58% |
| KEGG_TYPE_I_DIABETES_MELLITUS | 25 | 0.675559 | 1.680333 | 0.036797 | 0.176451 | 0.488 | 3079 | tags=72%, list=23%, signal=93% |
| KEGG_GRAFT_VERSUS_HOST_DISEASE | 24 | 0.746227 | 1.672133 | 0.044681 | 0.173156 | 0.501 | 2493 | tags=79%, list=18%, signal=97% |
| KEGG_NEUROACTIVE_LIGAND_RECEPTOR_INTERACTION | 107 | 0.434694 | 1.658422 | 0.005115 | 0.177866 | 0.53 | 2111 | tags=36%, list=15%, signal=43% |
| KEGG_PRIMARY_IMMUNODEFICIENCY | 24 | 0.675547 | 1.627136 | 0.048523 | 0.211331 | 0.59 | 1938 | tags=63%, list=14%, signal=73% |
| KEGG_B_CELL_RECEPTOR_SIGNALING_PATHWAY | 55 | 0.519313 | 1.609815 | 0.067762 | 0.225019 | 0.629 | 1631 | tags=40%, list=12%, signal=45% |
| KEGG_ALLOGRAFT_REJECTION | 20 | 0.745882 | 1.606325 | 0.046512 | 0.217333 | 0.633 | 3079 | tags=90%, list=23%, signal=116% |
| KEGG_AUTOIMMUNE_THYROID_DISEASE | 22 | 0.70187 | 1.59605 | 0.060606 | 0.220174 | 0.654 | 3079 | tags=82%, list=23%, signal=106% |

| | | | | | | | | |
|---|---|---|---|---|---|---|---|---|
| KEGG_LEUKOCYTE_TRANSENDOTHELIAL_MIGRATION | 77 | 0.45788 | 1.586102 | 0.072581 | 0.223359 | 0.68 | 2254 | tags=40%, list=17%, signal=48% |
| KEGG_APOPTOSIS | 62 | 0.467115 | 1.554045 | 0.05157 | 0.261476 | 0.728 | 3053 | tags=45%, list=22%, signal=58% |
| **Oncogenic (HFGC)** | | | | | | | | |
| NAME | SIZE | ES | NES | NOM p-val | FDR q-val | FWER p-val | RANK AT MAX | LEADING EDGE |
| GCNP_SHH_UP_EARLY.V1_UP | 112 | -0.44558 | -1.61302 | 0.022945 | 0.570273 | 0.455 | 2893 | tags=47%, list=21%, signal=60% |
| RB_P107_DN.V1_UP | 94 | -0.52063 | -1.57264 | 0.081439 | 0.395183 | 0.518 | 2187 | tags=47%, list=16%, signal=55% |
| RB_DN.V1_UP | 91 | -0.46223 | -1.53291 | 0.055985 | 0.351861 | 0.583 | 3121 | tags=45%, list=23%, signal=58% |
| RPS14_DN.V1_DN | 128 | -0.43877 | -1.51859 | 0.066288 | 0.2904 | 0.609 | 2817 | tags=41%, list=21%, signal=52% |
| HOXA9_DN.V1_DN | 131 | -0.39457 | -1.49063 | 0.052632 | 0.280516 | 0.67 | 1376 | tags=24%, list=10%, signal=27% |
| RB_P130_DN.V1_UP | 73 | -0.40246 | -1.44796 | 0.06203 | 0.305194 | 0.726 | 3345 | tags=42%, list=25%, signal=56% |
| GCNP_SHH_UP_LATE.V1_UP | 128 | -0.38965 | -1.42891 | 0.093458 | 0.294296 | 0.747 | 2842 | tags=43%, list=21%, signal=54% |
| YAP1_DN | 24 | -0.43614 | -1.38845 | 0.108987 | 0.325464 | 0.791 | 1761 | tags=33%, list=13%, signal=38% |
| SRC_UP.V1_DN | 105 | -0.34118 | -1.31503 | 0.109185 | 0.428184 | 0.869 | 3483 | tags=41%, list=26%, signal=55% |
| MTOR_UP.V1_UP | 107 | -0.33723 | -1.30871 | 0.155009 | 0.399462 | 0.885 | 2733 | tags=34%, list=20%, signal=42% |
| E2F3_UP.V1_UP | 102 | -0.3626 | -1.30344 | 0.166963 | 0.373698 | 0.888 | 3872 | tags=45%, list=28%, signal=63% |
| PRC2_EZH2_UP.V1_UP | 109 | -0.36104 | -1.29241 | 0.179487 | 0.362678 | 0.9 | 2825 | tags=38%, list=21%, signal=47% |
| MYC_UP.V1_UP | 106 | -0.34623 | -1.28213 | 0.174147 | 0.352583 | 0.908 | 3044 | tags=35%, list=22%, signal=45% |
| E2F1_UP.V1_UP | 119 | -0.37075 | -1.21113 | 0.30916 | 0.458367 | 0.953 | 3075 | tags=41%, list=23%, signal=53% |
| CSR_LATE_UP.V1_UP | 125 | -0.37744 | -1.20792 | 0.306569 | 0.433931 | 0.954 | 3417 | tags=42%, list=25%, signal=56% |
| EGFR_UP.V1_DN | 136 | -0.29014 | -1.19597 | 0.196 | 0.430325 | 0.959 | 2466 | tags=26%, list=18%, signal=31% |
| AKT_UP_MTOR_DN.V1_DN | 117 | -0.2773 | -1.11006 | 0.307692 | 0.57449 | 0.979 | 2660 | tags=27%, list=20%, signal=34% |

| | | | | | | | | |
|---|---|---|---|---|---|---|---|---|
| | | | | | 9 | | | |
| GLI1_UP.V1_DN | 23 | -0.38021 | -1.07906 | 0.381481 | 0.613293 | 0.983 | 2637 | tags=35%, list=19%, signal=43% |
| VEGF_A_UP.V1_DN | 145 | -0.29105 | -1.05405 | 0.385338 | 0.637762 | 0.987 | 3072 | tags=34%, list=23%, signal=43% |
| TBK1.DN.48HRS_UP | 37 | -0.32568 | -1.04908 | 0.412863 | 0.617005 | 0.988 | 4923 | tags=57%, list=36%, signal=89% |
| **Oncogenic (LFGC)** | | | | | | | | |
| NAME | SIZE | ES | NES | NOM p-val | FDR q-val | FWER p-val | RANK AT MAX | LEADING EDGE |
| P53_DN.V1_DN | 132 | 0.498719 | 1.865869 | 0 | 0.146619 | 0.072 | 2879 | tags=48%, list=21%, signal=60% |
| PTEN_DN.V2_UP | 94 | 0.478087 | 1.825921 | 0 | 0.122728 | 0.104 | 2384 | tags=40%, list=17%, signal=49% |
| HOXA9_DN.V1_UP | 145 | 0.532434 | 1.818311 | 0.004107 | 0.086655 | 0.11 | 3358 | tags=52%, list=25%, signal=69% |
| STK33_NOMO_UP | 225 | 0.491031 | 1.809651 | 0.010395 | 0.073094 | 0.118 | 3463 | tags=52%, list=25%, signal=68% |
| RPS14_DN.V1_UP | 136 | 0.581069 | 1.798718 | 0.014675 | 0.06671 | 0.13 | 2856 | tags=57%, list=21%, signal=71% |
| RELA_DN.V1_UP | 86 | 0.457427 | 1.758811 | 0.002141 | 0.088525 | 0.176 | 2304 | tags=41%, list=17%, signal=49% |
| STK33_SKM_UP | 211 | 0.475827 | 1.739224 | 0.012793 | 0.091894 | 0.205 | 2843 | tags=47%, list=21%, signal=58% |
| EGFR_UP.V1_UP | 149 | 0.473234 | 1.737659 | 0.010417 | 0.082116 | 0.208 | 2460 | tags=41%, list=18%, signal=49% |
| STK33_UP | 221 | 0.473561 | 1.731673 | 0.02079 | 0.0773 | 0.216 | 2577 | tags=42%, list=19%, signal=51% |
| KRAS.LUNG_UP.V1_UP | 74 | 0.461387 | 1.71326 | 0.006682 | 0.083486 | 0.241 | 3118 | tags=42%, list=23%, signal=54% |
| VEGF_A_UP.V1_UP | 99 | 0.445195 | 1.704662 | 0.008403 | 0.082356 | 0.26 | 2205 | tags=34%, list=16%, signal=41% |
| P53_DN.V2_UP | 73 | 0.501725 | 1.699962 | 0.025229 | 0.079479 | 0.272 | 2736 | tags=45%, list=20%, signal=56% |
| LEF1_UP.V1_UP | 109 | 0.475131 | 1.698026 | 0.008949 | 0.075302 | 0.275 | 2520 | tags=41%, list=18%, signal=50% |
| MEL18_DN.V1_DN | 99 | 0.454059 | 1.687313 | 0.008333 | 0.078173 | 0.289 | 2586 | tags=41%, list=19%, signal=51% |
| IL2_UP.V1_UP | 118 | 0.43754 | 1.622774 | 0.025114 | 0.130928 | 0.403 | 2605 | tags=37%, list=19%, signal=46% |
| BCAT_GDS748_UP | 35 | 0.46845 | 1.60452 | 0.020 | 0.14 | 0.442 | 1610 | tags=29%, list=12%, |

| | | | | | | | | |
|---|---|---|---|---|---|---|---|---|
| | | | | 325 | 197<br>4 | | | signal=32% |
| KRAS.BREAST_UP.V1_UP | 68 | 0.45164 | 1.60253<br>1 | 0.043<br>373 | 0.13<br>565<br>1 | 0.443 | 2676 | tags=41%, list=20%, signal=51% |
| PTEN_DN.V1_UP | 103 | 0.39776<br>4 | 1.60025<br>6 | 0.019<br>093 | 0.12<br>977<br>9 | 0.445 | 2812 | tags=38%, list=21%, signal=47% |
| BMI1_DN_MEL18_DN.V1_DN | 98 | 0.41994<br>3 | 1.58354<br>6 | 0.024<br>59 | 0.13<br>773<br>4 | 0.471 | 2432 | tags=37%, list=18%, signal=44% |
| BMI1_DN.V1_DN | 87 | 0.38760<br>4 | 1.55729 | 0.018<br>908 | 0.15<br>855<br>6 | 0.526 | 2472 | tags=33%, list=18%, signal=40% |

### Table S6. GSEA result compared HFGC with LFGC of TCGA HCC

| Biological Process (HFGC) | | | | | | | | |
|---|---|---|---|---|---|---|---|---|
| NAME | SIZE | ES | NES | NOM p-val | FDR q-val | FWER p-val | RANK AT MAX | LEADING EDGE |
| CHROMOSOME_SEGREGATION | 29 | -0.7256807 | -2.1769378 | 0.0021142 | 0.0108505 | 0.005 | 509 | tags=45%, list=3%, signal=46% |
| M_PHASE | 101 | -0.6577637 | -2.1104443 | 0.0039841 | 0.0159817 | 0.015 | 2517 | tags=49%, list=15%, signal=57% |
| DNA_DEPENDENT_DNA_REPLICATION | 51 | -0.6878951 | -2.1082685 | 0 | 0.011113 | 0.016 | 1911 | tags=47%, list=12%, signal=53% |
| REGULATION_OF_MITOSIS | 36 | -0.7227784 | -2.103995 | 0.0020325 | 0.0094778 | 0.018 | 441 | tags=39%, list=3%, signal=40% |
| CELL_CYCLE_PROCESS | 175 | -0.6020439 | -2.098091 | 0.0060606 | 0.0081045 | 0.021 | 1896 | tags=39%, list=11%, signal=44% |
| DNA_REPAIR | 110 | -0.5683734 | -2.078699 | 0 | 0.0090214 | 0.03 | 1911 | tags=35%, list=12%, signal=39% |
| CELL_CYCLE_PHASE | 153 | -0.5939316 | -2.0732765 | 0.0080645 | 0.0082688 | 0.033 | 2517 | tags=42%, list=15%, signal=49% |
| DNA_METABOLIC_PROCESS | 229 | -0.5257379 | -2.0633385 | 0.0040404 | 0.0084337 | 0.037 | 1911 | tags=32%, list=12%, signal=36% |
| DNA_REPLICATION | 93 | -0.5967884 | -2.0442684 | 0.0041494 | 0.0088635 | 0.046 | 1911 | tags=38%, list=12%, signal=42% |
| M_PHASE_OF_MITOTIC_CELL_CYCLE | 76 | -0.6681321 | -2.0280175 | 0.0082988 | 0.0103759 | 0.057 | 589 | tags=36%, list=4%, signal=37% |
| MITOSIS | 73 | -0.6653343 | -2.0132692 | 0.008316 | 0.0105645 | 0.066 | 589 | tags=36%, list=4%, signal=37% |
| RESPONSE_TO_DNA_DAMAGE_STIMULUS | 143 | -0.5180857 | -2.0058506 | 0.0020492 | 0.0105438 | 0.072 | 2706 | tags=37%, list=16%, signal=44% |
| REGULATION_OF_GENE_EXPRESSION_EPIGENETIC | 28 | -0.620373 | -1.9912269 | 0 | 0.0115952 | 0.083 | 1359 | tags=36%, list=8%, signal=39% |
| MEIOTIC_CELL_CYCLE | 30 | -0.6747697 | -1.977847 | 0 | 0.01243 | 0.089 | 2517 | tags=53%, list=15%, |

| | | | | | | | | |
|---|---|---|---|---|---|---|---|---|
| | | | | | 48 | | | signal=63% |
| RESPONSE_TO_ENDOGENOUS_STIMULUS | 178 | -0.4732107 | -1.9758451 | 0 | 0.011817 | 0.089 | 3235 | tags=38%, list=20%, signal=46% |
| ORGANELLE_LOCALIZATION | 23 | -0.6114302 | -1.9455433 | 0 | 0.0161809 | 0.132 | 441 | tags=30%, list=3%, signal=31% |
| CELL_CYCLE_CHECKPOINT_GO_0000075 | 42 | -0.6341423 | -1.9398936 | 0.01 | 0.0165203 | 0.144 | 2706 | tags=52%, list=16%, signal=62% |
| MITOTIC_CELL_CYCLE | 138 | -0.5599372 | -1.93874 | 0.0199203 | 0.0157299 | 0.145 | 1849 | tags=34%, list=11%, signal=38% |
| CELL_CYCLE_GO_0007049 | 284 | -0.4878363 | -1.9308627 | 0.0099404 | 0.0165395 | 0.16 | 1956 | tags=30%, list=12%, signal=33% |
| MEIOSIS_I | 17 | -0.7186291 | -1.9257898 | 0.001996 | 0.0167939 | 0.167 | 1504 | tags=53%, list=9%, signal=58% |
| **Biological Process (LFGC)** | | | | | | | | |
| NAME | SIZE | ES | NES | NOM p-val | FDR q-val | FWER p-val | RANK AT MAX | LEADING EDGE |
| BONE_REMODELING | 28 | 0.5837959 | 1.8827065 | 0.0022676 | 0.8077714 | 0.242 | 2183 | tags=39%, list=13%, signal=45% |
| G_PROTEIN_SIGNALING_COUPLED_TO_CYCLIC_NUCLEOTIDE_SECOND_MESSENGER | 94 | 0.4412195 | 1.8440132 | 0 | 0.6043246 | 0.326 | 3773 | tags=40%, list=23%, signal=52% |
| CYCLIC_NUCLEOTIDE_MEDIATED_SIGNALING | 96 | 0.4327054 | 1.8185772 | 0 | 0.5129932 | 0.403 | 3773 | tags=40%, list=23%, signal=51% |
| SECOND_MESSENGER_MEDIATED_SIGNALING | 142 | 0.4236116 | 1.7985157 | 0 | 0.4633081 | 0.457 | 3773 | tags=39%, list=23%, signal=51% |
| TISSUE_REMODELING | 29 | 0.547134 | 1.7777083 | 0.0022676 | 0.4474539 | 0.509 | 2183 | tags=38%, list=13%, signal=44% |
| G_PROTEIN_SIGNALING_COUPLED_TO_CAMP_NUCLEOTIDE_SECOND_MESSENGER | 62 | 0.4338301 | 1.7289587 | 0.0043764 | 0.5601189 | 0.64 | 3773 | tags=40%, list=23%, signal=52% |
| CAMP_MEDIATED_SIGNALING | 63 | 0.429736 | 1.7278168 | 0.0044346 | 0.4838306 | 0.643 | 3773 | tags=40%, list=23%, signal=51% |

| | | | | | | | | |
|---|---|---|---|---|---|---|---|---|
| REGULATION_OF_HEART_CONTRACTION | 24 | 0.5629414 | 1.7137011 | 0.02412 28 | 0.47892 64 | 0.68 | 3060 | tags=46%, list=18%, signal=56% |
| RESPONSE_TO_VIRUS | 37 | 0.6080874 | 1.7080084 | 0.03082 85 | 0.44412 79 | 0.695 | 2834 | tags=54%, list=17%, signal=65% |
| ENERGY_DERIVATION_BY_OXIDATION_OF_ORGANIC_COMPOUNDS | 35 | 0.5616012 | 1.6965986 | 0.02191 24 | 0.43500 27 | 0.726 | 3566 | tags=51%, list=22%, signal=65% |
| G_PROTEIN_SIGNALING_COUPLED_TO_IP3_SECOND_MESSENGERPHOSPHOLIPASE_C_ACTIVATING | 43 | 0.4625911 | 1.6809778 | 0.00451 47 | 0.44715 5 | 0.765 | 3406 | tags=40%, list=21%, signal=50% |
| PHOSPHOINOSITIDE_MEDIATED_SIGNALING | 46 | 0.4494167 | 1.6508359 | 0.00674 16 | 0.51648 34 | 0.818 | 3406 | tags=39%, list=21%, signal=49% |
| CELL_SUBSTRATE_ADHESION | 38 | 0.4938582 | 1.6485391 | 0.02169 63 | 0.48533 69 | 0.823 | 3722 | tags=47%, list=22%, signal=61% |
| RESPONSE_TO_OTHER_ORGANISM | 65 | 0.5113334 | 1.607881 | 0.04259 63 | 0.59871 35 | 0.888 | 4209 | tags=54%, list=25%, signal=72% |
| AXON_GUIDANCE | 20 | 0.546113 | 1.5928898 | 0.03671 71 | 0.61644 1 | 0.914 | 1687 | tags=40%, list=10%, signal=44% |
| CELL_MATRIX_ADHESION | 37 | 0.4754125 | 1.5913014 | 0.03571 43 | 0.58457 06 | 0.915 | 3722 | tags=46%, list=22%, signal=59% |
| INSULIN_RECEPTOR_SIGNALING_PATHWAY | 17 | 0.5940896 | 1.5898261 | 0.04713 11 | 0.55672 32 | 0.916 | 4604 | tags=65%, list=28%, signal=89% |
| RESPONSE_TO_BIOTIC_STIMULUS | 101 | 0.4569852 | 1.5837369 | 0.03594 08 | 0.54674 75 | 0.922 | 3423 | tags=43%, list=21%, signal=53% |
| SKELETAL_DEVELOPMENT | 95 | 0.4345817 | 1.5740607 | 0.05694 76 | 0.55145 54 | 0.931 | 3553 | tags=44%, list=21%, signal=56% |
| CALCIUM_ION_TRANSPORT | 25 | 0.5010728 | 1.5728935 | 0.02422 91 | 0.52764 92 | 0.932 | 3870 | tags=44%, list=23%, signal=57% |
| **KEGG Pathway (HFGC)** | | | | | | | | |
| NAME | SIZE | ES | NES | NOM p-val | FDR q-val | FWER p-val | RANK AT MAX | LEADING EDGE |
| KEGG_HOMOLOGOUS_RECOMBINATION | 27 | -0.7821631 | -2.0921094 | 0 | 0.00834 | 0.013 | 1419 | tags=52%, list=9%, |

| | | | | | | | | |
|---|---|---|---|---|---|---|---|---|
| | | | | | 65 | | | signal=57% |
| KEGG_MISMATCH_REPAIR | 22 | -0.7855697 | -1.9719963 | 0.0020833 | 0.0225981 | 0.059 | 2093 | tags=55%, list=13%, signal=62% |
| KEGG_CELL_CYCLE | 115 | -0.5960533 | -1.9206722 | 0.006237 | 0.0272204 | 0.09 | 2339 | tags=47%, list=14%, signal=54% |
| KEGG_DNA_REPLICATION | 33 | -0.8120576 | -1.9124084 | 0 | 0.0227869 | 0.096 | 570 | tags=58%, list=3%, signal=60% |
| KEGG_BASE_EXCISION_REPAIR | 30 | -0.6350449 | -1.8123506 | 0.0081301 | 0.0599964 | 0.239 | 2604 | tags=53%, list=16%, signal=63% |
| KEGG_PYRIMIDINE_METABOLISM | 84 | -0.457888 | -1.648735 | 0.0167364 | 0.2059161 | 0.615 | 1291 | tags=25%, list=8%, signal=27% |
| KEGG_SPLICEOSOME | 116 | -0.5450181 | -1.6371747 | 0.0433925 | 0.1944765 | 0.642 | 3614 | tags=52%, list=22%, signal=66% |
| KEGG_OOCYTE_MEIOSIS | 101 | -0.4245597 | -1.6295837 | 0.0168224 | 0.1786417 | 0.66 | 976 | tags=22%, list=6%, signal=23% |
| KEGG_NUCLEOTIDE_EXCISION_REPAIR | 39 | -0.5257825 | -1.6200255 | 0.0351563 | 0.1698185 | 0.684 | 2143 | tags=33%, list=13%, signal=38% |
| KEGG_N_GLYCAN_BIOSYNTHESIS | 42 | -0.4592321 | -1.5965395 | 0.0368664 | 0.181347 | 0.733 | 2905 | tags=38%, list=18%, signal=46% |
| KEGG_MATURITY_ONSET_DIABETES_OF_THE_YOUNG | 25 | -0.5502435 | -1.5653161 | 0.0581162 | 0.2020812 | 0.794 | 3270 | tags=44%, list=20%, signal=55% |
| KEGG_BASAL_TRANSCRIPTION_FACTORS | 31 | -0.4453705 | -1.4513489 | 0.082996 | 0.3563504 | 0.934 | 2762 | tags=32%, list=17%, signal=39% |
| KEGG_UBIQUITIN_MEDIATED_PROTEOLYSIS | 119 | -0.3430879 | -1.4071141 | 0.0550847 | 0.4076045 | 0.969 | 3056 | tags=29%, list=18%, signal=35% |
| KEGG_PROGESTERONE_MEDIATED_OOCYTE_MATURATION | 77 | -0.3630972 | -1.3535997 | 0.1175337 | 0.4821322 | 0.987 | 2171 | tags=27%, list=13%, signal=31% |
| KEGG_P53_SIGNALING_PATHWAY | 60 | -0.3579201 | -1.3314016 | 0.1078838 | 0.4952371 | 0.99 | 2250 | tags=28%, list=14%, signal=33% |
| KEGG_PENTOSE_AND_GLUCURONATE_INTERCONVERSIONS | 26 | -0.520304 | -1.2967172 | 0.2302632 | 0.5338726 | 0.993 | 4224 | tags=46%, list=25%, signal=62 |

| | | | | | | | | |
|---|---|---|---|---|---|---|---|---|
| | | | | | | | | % |
| KEGG_BLADDER_CANCER | 37 | -0.3534546 | -1.1797932 | 0.244898 | 0.7698757 | 1 | 4124 | tags=43%, list=25%, signal=57% |
| KEGG_THYROID_CANCER | 28 | -0.351849 | -1.1598387 | 0.2665289 | 0.7766116 | 1 | 2043 | tags=21%, list=12%, signal=24% |
| KEGG_TERPENOID_BACKBONE_BIOSYNTHESIS | 15 | -0.4792478 | -1.1281612 | 0.3621053 | 0.8139191 | 1 | 1813 | tags=40%, list=11%, signal=45% |
| KEGG_RIBOSOME | 80 | -0.5202168 | -1.118066 | 0.4309278 | 0.7972894 | 1 | 5320 | tags=66%, list=32%, signal=97% |
| **KEGG Pathway (LFGC)** | | | | | | | | |
| NAME | SIZE | ES | NES | NOM p-val | FDR q-val | FWER p-val | RANK AT MAX | LEADING EDGE |
| KEGG_CALCIUM_SIGNALING_PATHWAY | 155 | 0.4612967 | 1.9231007 | 0 | 0.2147519 | 0.101 | 3474 | tags=39%, list=21%, signal=49% |
| KEGG_REGULATION_OF_AUTOPHAGY | 19 | 0.5696816 | 1.750542 | 0.0102041 | 0.7194228 | 0.397 | 1729 | tags=32%, list=10%, signal=35% |
| KEGG_VASCULAR_SMOOTH_MUSCLE_CONTRACTION | 106 | 0.4589802 | 1.7177393 | 0.0108225 | 0.6369793 | 0.471 | 2705 | tags=37%, list=16%, signal=44% |
| KEGG_VALINE_LEUCINE_AND_ISOLEUCINE_DEGRADATION | 42 | 0.6580745 | 1.6309192 | 0.0534653 | 0.9256749 | 0.673 | 2805 | tags=60%, list=17%, signal=71% |
| KEGG_HEMATOPOIETIC_CELL_LINEAGE | 80 | 0.4910937 | 1.6131573 | 0.0688259 | 0.8442543 | 0.715 | 4248 | tags=52%, list=26%, signal=70% |
| KEGG_GLYCOSAMINOGLYCAN_BIOSYNTHESIS_CHONDROITIN_SULFATE | 21 | 0.5366713 | 1.5693448 | 0.0434783 | 0.927301 | 0.803 | 1176 | tags=38%, list=7%, signal=41% |
| KEGG_CYTOKINE_CYTOKINE_RECEPTOR_INTERACTION | 226 | 0.4284237 | 1.5615697 | 0.0597015 | 0.8342742 | 0.815 | 4104 | tags=43%, list=25%, signal=56% |
| KEGG_APOPTOSIS | 78 | 0.4165952 | 1.5483484 | 0.0372549 | 0.790282 | 0.836 | 3311 | tags=37%, list=20%, signal=46% |
| KEGG_DILATED_CARDIOMYOPATHY | 82 | 0.4527537 | 1.5069128 | 0.0651163 | 0.8841164 | 0.891 | 3267 | tags=41%, list=20%, signal=51% |

| | | | | | | | | |
|---|---|---|---|---|---|---|---|---|
| KEGG_NICOTINATE_AND_NICOTINAMIDE_METABOLISM | 22 | 0.4871063 | 1.4966484 | 0.0565657 | 0.8410843 | 0.896 | 5949 | tags=64%, list=36%, signal=99% |
| KEGG_NEUROACTIVE_LIGAND_RECEPTOR_INTERACTION | 229 | 0.3238937 | 1.4924084 | 0.0221607 | 0.7820019 | 0.901 | 3755 | tags=31%, list=23%, signal=40% |
| KEGG_B_CELL_RECEPTOR_SIGNALING_PATHWAY | 71 | 0.4492866 | 1.4923524 | 0.0848485 | 0.7168351 | 0.901 | 2989 | tags=41%, list=18%, signal=50% |
| KEGG_CITRATE_CYCLE_TCA_CYCLE | 28 | 0.573049 | 1.485822 | 0.0989899 | 0.6881833 | 0.909 | 3566 | tags=54%, list=22%, signal=68% |
| KEGG_CHEMOKINE_SIGNALING_PATHWAY | 179 | 0.403439 | 1.4697447 | 0.0907173 | 0.6943442 | 0.925 | 4240 | tags=45%, list=26%, signal=60% |
| KEGG_ETHER_LIPID_METABOLISM | 30 | 0.4054771 | 1.4601877 | 0.0378947 | 0.6810028 | 0.938 | 1951 | tags=30%, list=12%, signal=34% |
| KEGG_NATURAL_KILLER_CELL_MEDIATED_CYTOTOXICITY | 111 | 0.4265771 | 1.4512093 | 0.1234568 | 0.6678835 | 0.944 | 3561 | tags=43%, list=21%, signal=55% |
| KEGG_JAK_STAT_SIGNALING_PATHWAY | 122 | 0.3924675 | 1.4459618 | 0.0817204 | 0.6446661 | 0.949 | 3064 | tags=32%, list=18%, signal=39% |
| KEGG_CELL_ADHESION_MOLECULES_CAMS | 115 | 0.4258497 | 1.4085304 | 0.1283644 | 0.7238066 | 0.968 | 3303 | tags=41%, list=20%, signal=51% |
| KEGG_ECM_RECEPTOR_INTERACTION | 79 | 0.4428824 | 1.4052496 | 0.169697 | 0.6949843 | 0.97 | 3400 | tags=46%, list=21%, signal=57% |
| KEGG_LEISHMANIA_INFECTION | 63 | 0.4759485 | 1.3842937 | 0.1816367 | 0.7285357 | 0.976 | 3303 | tags=43%, list=20%, signal=53% |
| **Oncogenic (HFGC)** | | | | | | | | |
| NAME | SIZE | ES | NES | NOM p-val | FDR q-val | FWER p-val | RANK AT MAX | LEADING EDGE |
| RPS14_DN.V1_DN | 163 | -0.5448914 | -2.1645298 | 0 | 0 | 0 | 1825 | tags=33%, list=11%, signal=37% |
| PRC2_EZH2_UP.V1_UP | 165 | -0.4864141 | -2.0560021 | 0 | 8.97E-04 | 0.005 | 886 | tags=27%, list=5%, signal=28% |
| E2F1_UP.V1_UP | 173 | -0.4767531 | -1.8829107 | 0.0037951 | 0.0119927 | 0.072 | 2545 | tags=35%, list=15%, signal=41 |

| | | | | | | | | |
|---|---|---|---|---|---|---|---|---|
| | | | | | | | | % |
| RB_P107_DN.V1_UP | 125 | -0.5648047 | -1.833929 | 0.0201613 | 0.0170637 | 0.114 | 1595 | tags=42%, list=10%, signal=47% |
| RB_DN.V1_UP | 119 | -0.4355741 | -1.7348772 | 0.0061856 | 0.0374932 | 0.227 | 2465 | tags=33%, list=15%, signal=38% |
| GCNP_SHH_UP_EARLY.V1_UP | 152 | -0.4522155 | -1.7185619 | 0.0174757 | 0.0363483 | 0.251 | 3025 | tags=36%, list=18%, signal=44% |
| CSR_LATE_UP.V1_UP | 152 | -0.4781577 | -1.688554 | 0.0246212 | 0.0403553 | 0.31 | 2151 | tags=38%, list=13%, signal=43% |
| E2F3_UP.V1_UP | 162 | -0.3926694 | -1.6751021 | 0.0058366 | 0.0394561 | 0.326 | 2796 | tags=31%, list=17%, signal=37% |
| HOXA9_DN.V1_DN | 164 | -0.4006853 | -1.6392138 | 0.0080321 | 0.0488003 | 0.412 | 3375 | tags=37%, list=20%, signal=46% |
| GCNP_SHH_UP_LATE.V1_UP | 160 | -0.4417787 | -1.6136187 | 0.0329457 | 0.0539783 | 0.465 | 1563 | tags=25%, list=9%, signal=27% |
| PRC2_EDD_UP.V1_UP | 171 | -0.3744294 | -1.5758784 | 0.0288066 | 0.0666854 | 0.558 | 2143 | tags=26%, list=13%, signal=30% |
| MTOR_UP.V1_UP | 153 | -0.344041 | -1.5182328 | 0.0316832 | 0.0948124 | 0.684 | 2817 | tags=28%, list=17%, signal=34% |
| RB_P130_DN.V1_UP | 116 | -0.3776458 | -1.5029774 | 0.0246407 | 0.0959503 | 0.711 | 2850 | tags=32%, list=17%, signal=38% |
| SRC_UP.V1_DN | 145 | -0.3309808 | -1.4914079 | 0.0149533 | 0.0968481 | 0.73 | 1739 | tags=21%, list=10%, signal=24% |
| MYC_UP.V1_UP | 146 | -0.3925988 | -1.4314686 | 0.1064718 | 0.1335323 | 0.838 | 4985 | tags=49%, list=30%, signal=69% |
| VEGF_A_UP.V1_DN | 176 | -0.3885405 | -1.419865 | 0.0994264 | 0.1346864 | 0.855 | 1489 | tags=21%, list=9%, signal=23% |
| TBK1.DN.48HRS_UP | 46 | -0.397412 | -1.419133 | 0.0611814 | 0.1273293 | 0.857 | 3288 | tags=35%, list=20%, signal=43% |
| BCAT_GDS748_DN | 40 | -0.3437548 | -1.3097826 | 0.1162325 | 0.2279908 | 0.96 | 3751 | tags=45%, list=23%, signal=58 |

| | | | | | | | | |
|---|---|---|---|---|---|---|---|---|
| | | | | | | | | % |
| NFE2L2.V2 | 381 | - 0.2793905 | - 1.2796575 | 0.12 375 25 | 0.25 516 36 | 0.976 | 306 6 | tags=24%, list=18%, signal=28 % |
| DCA_UP.V1_DN | 150 | - 0.2611077 | - 1.2338156 | 0.09 638 55 | 0.30 858 01 | 0.991 | 272 4 | tags=23%, list=16%, signal=27 % |
| **Oncogenic (LFGC)** | | | | | | | | |
| NAME | SIZ E | ES | NES | NO M p- val | FDR q- val | FWE R p- val | RA NK AT MA X | LEADING EDGE |
| HOXA9_DN.V1_UP | 171 | 0.519038 | 1.9048997 | 0.00 394 48 | 0.10 100 86 | 0.054 | 441 4 | tags=56%, list=27%, signal=75 % |
| PTEN_DN.V1_UP | 160 | 0.4476807 | 1.8342359 | 0.00 222 72 | 0.12 340 86 | 0.112 | 347 9 | tags=41%, list=21%, signal=52 % |
| LEF1_UP.V1_UP | 163 | 0.511918 | 1.8233509 | 0.01 470 59 | 0.09 209 85 | 0.12 | 352 6 | tags=47%, list=21%, signal=59 % |
| RPS14_DN.V1_UP | 179 | 0.5407417 | 1.773334 | 0.03 080 08 | 0.12 119 03 | 0.176 | 352 5 | tags=48%, list=21%, signal=60 % |
| BMI1_DN_MEL18_DN.V1_DN | 131 | 0.4469608 | 1.7732651 | 0 | 0.09 695 23 | 0.176 | 368 2 | tags=37%, list=22%, signal=48 % |
| P53_DN.V1_DN | 175 | 0.4529273 | 1.7417918 | 0.01 006 04 | 0.11 072 58 | 0.224 | 386 7 | tags=45%, list=23%, signal=58 % |
| CAHOY_ASTROGLIAL | 86 | 0.5048821 | 1.7266909 | 0.02 298 85 | 0.11 169 5 | 0.252 | 297 6 | tags=44%, list=18%, signal=54 % |
| ATF2_UP.V1_DN | 171 | 0.4762719 | 1.7226397 | 0.01 4 | 0.10 003 24 | 0.254 | 377 0 | tags=49%, list=23%, signal=62 % |
| KRAS.600_UP.V1_UP | 236 | 0.3785581 | 1.7125484 | 0.00 423 73 | 0.09 801 76 | 0.268 | 325 7 | tags=33%, list=20%, signal=41 % |
| MEL18_DN.V1_DN | 132 | 0.4435261 | 1.708167 | 0.00 398 41 | 0.09 133 19 | 0.273 | 400 1 | tags=43%, list=24%, signal=56 % |
| PRC2_EZH2_UP.V1_DN | 158 | 0.4094486 | 1.6979431 | 0.00 210 97 | 0.09 284 | 0.295 | 282 0 | tags=30%, list=17%, signal=36 % |

| | | | | | | | | |
|---|---|---|---|---|---|---|---|---|
| ATF2_S_UP.V1_DN | 165 | 0.4447916 | 1.678042 | 0.0218254 | 0.1017227 | 0.328 | 3682 | tags=44%, list=22%, signal=56% |
| SNF5_DN.V1_DN | 144 | 0.398627 | 1.6590326 | 0.001992 | 0.1127894 | 0.372 | 3607 | tags=38%, list=22%, signal=48% |
| PTEN_DN.V2_UP | 121 | 0.4029441 | 1.6550689 | 0.008016 | 0.1077028 | 0.382 | 3786 | tags=40%, list=23%, signal=51% |
| CYCLIN_D1_KE_.V1_DN | 171 | 0.3794514 | 1.6550121 | 0.0105485 | 0.1005226 | 0.382 | 4093 | tags=40%, list=25%, signal=53% |
| STK33_SKM_UP | 245 | 0.4252608 | 1.6459278 | 0.0202429 | 0.1004487 | 0.398 | 2845 | tags=35%, list=17%, signal=41% |
| BMI1_DN.V1_UP | 131 | 0.4983971 | 1.6403983 | 0.0392562 | 0.0990974 | 0.409 | 2817 | tags=40%, list=17%, signal=48% |
| AKT_UP.V1_DN | 171 | 0.4042722 | 1.6097466 | 0.0150094 | 0.1165851 | 0.46 | 3838 | tags=43%, list=23%, signal=56% |
| STK33_UP | 256 | 0.4384923 | 1.6074026 | 0.0281125 | 0.1116833 | 0.464 | 4335 | tags=45%, list=26%, signal=59% |
| KRAS.KIDNEY_UP.V1_UP | 130 | 0.4079987 | 1.6069605 | 0.0246914 | 0.1065567 | 0.465 | 5084 | tags=53%, list=31%, signal=76% |

### Table S7. Overlapped GSEA result HFGC and LFGC of Thai HCC and Thai iCCA

| Name | Size | ES | NES | NOM.p.val | Size | ES | NES | NOM.p.val |
|---|---|---|---|---|---|---|---|---|
| | **HCC_HFGC** | | | | **iCCA_HFGC** | | | |
| **Biological Process** | | | | | | | | |
| DNA_DEPENDENT_DNA_REPLICATION | 40 | -0.6292 | -1.76602 | 0.0221 | 43 | -0.6745 | -1.90739 | 0.002 |
| CHROMOSOME_SEGREGATION | 21 | -0.6397 | -1.63112 | 0.0593 | 21 | -0.7025 | -1.8006 | 0.0039 |
| DNA_REPLICATION | 70 | -0.5365 | -1.61436 | 0.0561 | 75 | -0.5991 | -1.80371 | 0.008 |
| REGULATION_OF_MITOSIS | 26 | -0.5389 | -1.42042 | 0.1499 | 26 | -0.6822 | -1.78362 | 0.008 |
| M_PHASE | 71 | -0.4987 | -1.41633 | 0.1747 | 70 | -0.6641 | -1.85491 | 0.004 |
| **KEGG pathway** | | | | | | | | |
| KEGG_GLYCOSYLPHOSPHATIDYLINOSITOL_GPI_ANCHOR_BIOSYNTHESIS | 17 | -0.6601 | -1.68827 | 0.0137 | 18 | -0.5825 | -1.49526 | 0.0538 |
| KEGG_MISMATCH_REPAIR | 20 | -0.6784 | -1.6523 | 0.0307 | 19 | -0.7417 | -1.77585 | 0.0059 |
| KEGG_DNA_REPLICATION | 30 | -0.6926 | -1.58928 | 0.0615 | 29 | -0.8124 | -1.86008 | 0.0019 |
| KEGG_N_GLYCAN_BIOSYNTHESIS | 35 | -0.478 | -1.51036 | 0.0625 | 37 | -0.5023 | -1.56237 | 0.0462 |
| KEGG_NUCLEOTIDE_EXCISION_REPAIR | 36 | -0.4986 | -1.43636 | 0.1288 | 34 | -0.5713 | -1.58788 | 0.0355 |
| KEGG_BASE_EXCISION_REPAIR | 23 | -0.5365 | -1.41653 | 0.1245 | 21 | -0.6762 | -1.7393 | 0.0077 |
| KEGG_CELL_CYCLE | 92 | -0.4608 | -1.35385 | 0.1968 | 90 | -0.6096 | -1.72089 | 0.0211 |
| KEGG_HOMOLOGOUS_RECOMBINATION | 17 | -0.5631 | -1.32721 | 0.182 | 16 | -0.7101 | -1.68874 | 0.0134 |
| KEGG_SPLICEOSOME | 95 | -0.4581 | -1.31074 | 0.2162 | 96 | -0.5311 | -1.552 | 0.0825 |
| KEGG_PYRIMIDINE_METABOLISM | 65 | -0.3794 | -1.22892 | 0.249 | 62 | -0.5061 | -1.59065 | 0.0476 |
| KEGG_AMINOACYL_TRNA_BIOSYNTHESIS | 32 | -0.4419 | -1.2235 | 0.2818 | 30 | -0.6253 | -1.60137 | 0.0329 |
| KEGG_PROGESTERONE_MEDIATED_OOCYTE_MATURATION | 54 | -0.3868 | -1.2147 | 0.2764 | 54 | -0.4414 | -1.44729 | 0.0696 |

| | | | | | | | | |
|---|---|---|---|---|---|---|---|---|
| KEGG_BASAL_TRANSCRIPTION_FACTORS | 25 | -0.405 | -1.17575 | 0.2684 | 24 | -0.5638 | -1.54818 | 0.0511 |
| KEGG_RNA_POLYMERASE | 23 | -0.4418 | -1.16567 | 0.3058 | 23 | -0.6545 | -1.64476 | 0.0059 |
| KEGG_OOCYTE_MEIOSIS | 75 | -0.3366 | -1.09607 | 0.3764 | 77 | -0.4877 | -1.61648 | 0.0206 |
| **Oncogenic pathway** | | | | | | | | |
| GCNP_SHH_UP_EARLY.V1_UP | 112 | -0.4456 | -1.61302 | 0.0229 | 115 | -0.4474 | -1.65347 | 0.0189 |
| RB_P107_DN.V1_UP | 94 | -0.5206 | -1.57264 | 0.0814 | 88 | -0.6059 | -1.923 | 0.0059 |
| RB_DN.V1_UP | 91 | -0.4622 | -1.53291 | 0.056 | 89 | -0.5635 | -1.93142 | 0.0039 |
| RPS14_DN.V1_DN | 128 | -0.4388 | -1.51859 | 0.0663 | 132 | -0.5152 | -1.76369 | 0.0213 |
| HOXA9_DN.V1_DN | 131 | -0.3946 | -1.49063 | 0.0526 | 135 | -0.4055 | -1.53515 | 0.0503 |
| RB_P130_DN.V1_UP | 73 | -0.4025 | -1.44796 | 0.062 | 78 | -0.4179 | -1.61577 | 0.0203 |
| GCNP_SHH_UP_LATE.V1_UP | 128 | -0.3897 | -1.42891 | 0.0935 | 128 | -0.3713 | -1.39788 | 0.1212 |
| E2F3_UP.V1_UP | 102 | -0.3626 | -1.30344 | 0.167 | 105 | -0.3547 | -1.40734 | 0.0911 |
| PRC2_EZH2_UP.V1_UP | 109 | -0.361 | -1.29241 | 0.1795 | 117 | -0.4752 | -1.77755 | 0.0115 |
| MYC_UP.V1_UP | 106 | -0.3462 | -1.28213 | 0.1741 | 105 | -0.4026 | -1.32091 | 0.1899 |
| E2F1_UP.V1_UP | 119 | -0.3708 | -1.21113 | 0.3092 | 123 | -0.5175 | -1.75653 | 0.0179 |
| CSR_LATE_UP.V1_UP | 125 | -0.3774 | -1.20792 | 0.3066 | 125 | -0.5026 | -1.61629 | 0.0247 |
| EGFR_UP.V1_DN | 136 | -0.2901 | -1.19597 | 0.196 | 135 | -0.3226 | -1.39144 | 0.0588 |
| VEGF_A_UP.V1_DN | 145 | -0.2911 | -1.05405 | 0.3853 | 147 | -0.3992 | -1.44666 | 0.1016 |
| TBK1.DN.48HRS_UP | 37 | -0.3257 | -1.04908 | 0.4129 | 39 | -0.4025 | -1.34081 | 0.1044 |
| | **HCC_LFGC** | | | | **iCCA_LFGC** | | | |
| **Biological Process** | | | | | | | | |

| | | | | | | | | |
|---|---|---|---|---|---|---|---|---|
| IMMUNE_RESPONSE | 145 | 0.6187 | 1.971024 | 0.0043 | 152 | 0.6352 | 1.852819 | 0.0233 |
| DEFENSE_RESPONSE | 173 | 0.5714 | 1.936295 | 0.0089 | 189 | 0.6299 | 1.883574 | 0.0162 |
| IMMUNE_SYSTEM_PROCESS | 207 | 0.5764 | 1.8863 | 0.0088 | 219 | 0.5803 | 1.845352 | 0.0307 |
| RESPONSE_TO_OTHER_ORGANISM | 37 | 0.6336 | 1.831636 | 0.0066 | 48 | 0.6462 | 1.828193 | 0.0161 |
| MULTI_ORGANISM_PROCESS | 77 | 0.5058 | 1.830873 | 0.0065 | 97 | 0.5187 | 1.767454 | 0.0182 |
| CELLULAR_DEFENSE_RESPONSE | 38 | 0.6674 | 1.82391 | 0.0137 | 40 | 0.6893 | 1.756464 | 0.0291 |
| LOCOMOTORY_BEHAVIOR | 64 | 0.5951 | 1.808114 | 0.0134 | 74 | 0.6033 | 1.772803 | 0.0164 |
| POSITIVE_REGULATION_OF_MULTICELLULAR_ORGANISMAL_PROCESS | 39 | 0.5598 | 1.78999 | 0.0107 | 42 | 0.6503 | 1.828494 | 0.0101 |
| INFLAMMATORY_RESPONSE | 90 | 0.5363 | 1.749392 | 0.0278 | 100 | 0.6446 | 1.858764 | 0.0142 |
| REGULATION_OF_IMMUNE_SYSTEM_PROCESS | 39 | 0.562 | 1.72683 | 0.0277 | 39 | 0.6153 | 1.72256 | 0.0339 |
| **KEGG pathway** | | | | | | | | |
| KEGG_HEMATOPOIETIC_CELL_LINEAGE | 47 | 0.7045 | 2.007128 | 0.0066 | 49 | 0.7264 | 1.866346 | 0.002 |
| KEGG_CYTOKINE_CYTOKINE_RECEPTOR_INTERACTION | 139 | 0.5616 | 1.909341 | 0.0093 | 156 | 0.5901 | 1.843299 | 0.002 |
| KEGG_JAK_STAT_SIGNALING_PATHWAY | 81 | 0.5483 | 1.882923 | 0.0104 | 84 | 0.4794 | 1.554089 | 0.0713 |
| KEGG_INTESTINAL_IMMUNE_NETWORK_FOR_IGA_PRODUCTION | 27 | 0.7342 | 1.830915 | 0.0066 | 29 | 0.6359 | 1.499596 | 0.1064 |
| KEGG_PRION_DISEASES | 30 | 0.602 | 1.797722 | 0.0084 | 29 | 0.5504 | 1.459598 | 0.092 |
| KEGG_CHEMOKINE_SIGNALING_PATHWAY | 132 | 0.5092 | 1.776465 | 0.0298 | 142 | 0.458 | 1.585065 | 0.0538 |
| KEGG_NATURAL_KILLER_CELL_MEDIATED_CYTOTOXICITY | 75 | 0.5672 | 1.768625 | 0.039 | 80 | 0.5364 | 1.660275 | 0.0455 |
| KEGG_CELL_ADHESION_MOLECULES_CAMS | 94 | 0.5157 | 1.71064 | 0.0379 | 93 | 0.5438 | 1.637101 | 0.052 |
| KEGG_GRAFT_VERSUS_HOST_DISEASE | 24 | 0.7462 | 1.672133 | 0.0447 | 25 | 0.7296 | 1.496257 | 0.0682 |
| KEGG_NEUROACTIVE_LIGAND_RECEPTOR_INTERACTION | 107 | 0.4347 | 1.658422 | 0.0051 | 108 | 0.4393 | 1.640409 | 0.0165 |
| KEGG_PRIMARY_IMMUNODEFICIENCY | 24 | 0.6755 | 1.627136 | 0.0485 | 26 | 0.6269 | 1.495657 | 0.1086 |
| KEGG_B_CELL_RECEPTOR_SIGNALING_PATHWAY | 55 | 0.5193 | 1.609815 | 0.0678 | 58 | 0.4919 | 1.570837 | 0.0593 |
| KEGG_AUTOIMMUNE_THYROID_DISEASE | 22 | 0.7019 | 1.59605 | 0.0606 | 23 | 0.6945 | 1.483169 | 0.0969 |
| KEGG_LEUKOCYTE_TRANSENDOTHELIAL_MIGRATION | 77 | 0.4579 | 1.586102 | 0.0726 | 76 | 0.4963 | 1.568311 | 0.0641 |

| **Oncogenic pathway** | | | | | | | | |
|---|---|---|---|---|---|---|---|---|
| P53_DN.V1_DN | 132 | 0.4987 | 1.865869 | 0 | 133 | 0.5355 | 1.770526 | 0.0101 |
| PTEN_DN.V2_UP | 94 | 0.4781 | 1.825921 | 0 | 94 | 0.4766 | 1.647227 | 0.0261 |
| HOXA9_DN.V1_UP | 145 | 0.5324 | 1.818311 | 0.0041 | 147 | 0.5021 | 1.675491 | 0.039 |
| STK33_NOMO_UP | 225 | 0.491 | 1.809651 | 0.0104 | 227 | 0.4533 | 1.637368 | 0.0456 |
| RPS14_DN.V1_UP | 136 | 0.5811 | 1.798718 | 0.0147 | 142 | 0.6025 | 1.857724 | 0.0081 |
| RELA_DN.V1_UP | 86 | 0.4574 | 1.758811 | 0.0021 | 91 | 0.4621 | 1.716071 | 0.0141 |
| STK33_SKM_UP | 211 | 0.4758 | 1.739224 | 0.0128 | 214 | 0.4085 | 1.600525 | 0.0593 |
| EGFR_UP.V1_UP | 149 | 0.4732 | 1.737659 | 0.0104 | 152 | 0.5307 | 1.807555 | 0.014 |
| STK33_UP | 221 | 0.4736 | 1.731673 | 0.0208 | 225 | 0.4324 | 1.631893 | 0.0504 |
| KRAS.LUNG_UP.V1_UP | 74 | 0.4614 | 1.71326 | 0.0067 | 81 | 0.4791 | 1.6239 | 0.0207 |
| P53_DN.V2_UP | 73 | 0.5017 | 1.699962 | 0.0252 | 79 | 0.5704 | 1.919475 | 0 |
| KRAS.BREAST_UP.V1_UP | 68 | 0.4516 | 1.602531 | 0.0434 | 71 | 0.4614 | 1.631583 | 0.0328 |

**Table S8. Overlapped recurrent mutation genes between Thai HCC and Thai iCCA.**

| Gene | mut.No.HCC | mut.freq.HCC | mut.No.iCCA | mut.freq.iCCA |
|---|---|---|---|---|
| TP53 | 22 | 0.355 | 37 | 0.411 |
| ARID2 | 9 | 0.145 | 6 | 0.067 |
| ARID1A | 9 | 0.145 | 17 | 0.189 |
| CTNNB1 | 8 | 0.129 | 2 | 0.022 |
| APOB | 7 | 0.113 | 4 | 0.044 |
| CSMD3 | 6 | 0.097 | 9 | 0.100 |
| AXIN1 | 5 | 0.081 | 1 | 0.011 |
| RYR2 | 4 | 0.065 | 7 | 0.078 |
| RB1 | 3 | 0.048 | 1 | 0.011 |
| ASXL1 | 3 | 0.048 | 1 | 0.011 |
| PTPRD | 3 | 0.048 | 3 | 0.033 |
| PKHD1 | 3 | 0.048 | 1 | 0.011 |
| PIK3CA | 3 | 0.048 | 4 | 0.044 |
| PDE4DIP | 3 | 0.048 | 2 | 0.022 |
| SMARCA4 | 3 | 0.048 | 2 | 0.022 |
| PRKDC | 3 | 0.048 | 6 | 0.067 |
| SCN5A | 3 | 0.048 | 4 | 0.044 |
| NF1 | 3 | 0.048 | 6 | 0.067 |
| PBRM1 | 3 | 0.048 | 4 | 0.044 |
| RYR1 | 3 | 0.048 | 4 | 0.044 |
| KEAP1 | 3 | 0.048 | 2 | 0.022 |
| MBD1 | 2 | 0.032 | 1 | 0.011 |
| EPHA5 | 2 | 0.032 | 1 | 0.011 |
| NCOA3 | 2 | 0.032 | 1 | 0.011 |
| GRM8 | 2 | 0.032 | 1 | 0.011 |
| IGF2R | 2 | 0.032 | 1 | 0.011 |
| KIT | 2 | 0.032 | 1 | 0.011 |
| COL1A1 | 2 | 0.032 | 2 | 0.022 |
| LRP1B | 2 | 0.032 | 10 | 0.111 |
| EPHA7 | 2 | 0.032 | 2 | 0.022 |
| ERBB4 | 2 | 0.032 | 4 | 0.044 |
| KMT2C | 2 | 0.032 | 11 | 0.122 |
| EML4 | 2 | 0.032 | 1 | 0.011 |
| SAMD9 | 2 | 0.032 | 1 | 0.011 |
| EPHB1 | 2 | 0.032 | 1 | 0.011 |
| PSIP1 | 2 | 0.032 | 5 | 0.056 |
| UBR5 | 2 | 0.032 | 2 | 0.022 |
| WHSC1 | 2 | 0.032 | 1 | 0.011 |
| ACVR2A | 2 | 0.032 | 3 | 0.033 |
| SF3B1 | 2 | 0.032 | 1 | 0.011 |
| DCC | 2 | 0.032 | 1 | 0.011 |
| ADAMTS20 | 2 | 0.032 | 7 | 0.078 |
| NF2 | 2 | 0.032 | 1 | 0.011 |
| DNMT3A | 1 | 0.016 | 2 | 0.022 |
| PTEN | 1 | 0.016 | 1 | 0.011 |
| MYH11 | 1 | 0.016 | 3 | 0.033 |

| | | | | |
|---|---|---|---|---|
| JAK1 | 1 | 0.016 | 2 | 0.022 |
| RICTOR | 1 | 0.016 | 1 | 0.011 |
| TSHR | 1 | 0.016 | 1 | 0.011 |
| NLRP1 | 1 | 0.016 | 1 | 0.011 |
| MED12 | 1 | 0.016 | 1 | 0.011 |
| FBXO11 | 1 | 0.016 | 2 | 0.022 |
| RNF213 | 1 | 0.016 | 1 | 0.011 |
| POT1 | 1 | 0.016 | 1 | 0.011 |
| ACVR1 | 1 | 0.016 | 1 | 0.011 |
| CHEK2 | 1 | 0.016 | 1 | 0.011 |
| MYBPC3 | 1 | 0.016 | 1 | 0.011 |
| BRCA2 | 1 | 0.016 | 3 | 0.033 |
| TSC2 | 1 | 0.016 | 2 | 0.022 |
| KMT2A | 1 | 0.016 | 1 | 0.011 |
| NSD1 | 1 | 0.016 | 1 | 0.011 |
| BCORL1 | 1 | 0.016 | 3 | 0.033 |
| ABL1 | 1 | 0.016 | 2 | 0.022 |
| EPHA3 | 1 | 0.016 | 1 | 0.011 |
| NTRK3 | 1 | 0.016 | 1 | 0.011 |
| AKAP9 | 1 | 0.016 | 3 | 0.033 |
| FN1 | 1 | 0.016 | 4 | 0.044 |
| KDM6A | 1 | 0.016 | 1 | 0.011 |
| ITGA9 | 1 | 0.016 | 1 | 0.011 |
| TGFBR2 | 1 | 0.016 | 1 | 0.011 |
| SYNE1 | 1 | 0.016 | 7 | 0.078 |
| RNASEL | 1 | 0.016 | 1 | 0.011 |
| PTPRT | 1 | 0.016 | 2 | 0.022 |
| POLH | 1 | 0.016 | 2 | 0.022 |
| CDH5 | 1 | 0.016 | 1 | 0.011 |
| GUCY1A2 | 1 | 0.016 | 1 | 0.011 |
| BAI3 | 1 | 0.016 | 4 | 0.044 |
| LPHN3 | 1 | 0.016 | 1 | 0.011 |
| COL3A1 | 1 | 0.016 | 3 | 0.033 |
| AFF3 | 1 | 0.016 | 1 | 0.011 |
| CACNA1S | 1 | 0.016 | 1 | 0.011 |
| ARID1B | 1 | 0.016 | 1 | 0.011 |
| HGF | 1 | 0.016 | 2 | 0.022 |
| TBX22 | 1 | 0.016 | 1 | 0.011 |
| IGF1R | 1 | 0.016 | 1 | 0.011 |
| EP400 | 1 | 0.016 | 3 | 0.033 |
| DST | 1 | 0.016 | 2 | 0.022 |
| GNAS | 1 | 0.016 | 3 | 0.033 |
| ATM | 1 | 0.016 | 5 | 0.056 |
| ERBB3 | 1 | 0.016 | 6 | 0.067 |
| NBN | 1 | 0.016 | 1 | 0.011 |
| BAP1 | 1 | 0.016 | 4 | 0.044 |
| TRRAP | 1 | 0.016 | 1 | 0.011 |
| PIK3R1 | 1 | 0.016 | 1 | 0.011 |

| | | | | |
|---|---|---|---|---|
| KRAS | 1 | 0.016 | 9 | 0.100 |
| FGFR2 | 1 | 0.016 | 3 | 0.033 |
| CTNNA1 | 1 | 0.016 | 2 | 0.022 |
| CDH11 | 1 | 0.016 | 2 | 0.022 |
| MYLK | 1 | 0.016 | 1 | 0.011 |
| MAP3K1 | 1 | 0.016 | 1 | 0.011 |

**Table S9. Recurrently mutated genes in Thai HCC**

| Gene | Chr. | POS | REF | ALT | AA Change | Mutation. Type | Effect | var.frq | snp.alt |
|---|---|---|---|---|---|---|---|---|---|
| TP53 | chr17 | 7578257 | C | A | E198* | SNV | STOP_GAINED | 1 | C>A |
| TP53 | chr17 | 7577509 | C | T | E258K | SNV | NON_SYNONYMOUS_CODING | 1 | C>T |
| TP53 | chr17 | 7577535 | C | A | R249M | SNV | NON_SYNONYMOUS_CODING | 1 | C>A |
| TP53 | chr17 | 7577534 | C | A | R249S | SNV | NON_SYNONYMOUS_CODING | 7 | C>A |
| TP53 | chr17 | 7578397 | TG | T | -178 | Indel | FRAME_SHIFT | 1 | Indel |
| TP53 | chr17 | 7577114 | C | T | C275Y | SNV | NON_SYNONYMOUS_CODING | 1 | C>T |
| TP53 | chr17 | 7577132 | CT | C | -269 | Indel | FRAME_SHIFT | 1 | Indel |
| TP53 | chr17 | 7578196 | A | T | V218E | SNV | NON_SYNONYMOUS_CODING | 1 | A>T |
| TP53 | chr17 | 7577610 | T | A | | Indel | SPLICE_SITE_ACCEPTOR | 1 | Indel |
| TP53 | chr17 | 7577120 | C | T | R273H | SNV | NON_SYNONYMOUS_CODING | 3 | C>T |
| TP53 | chr17 | 7578515 | T | C | K139E | SNV | NON_SYNONYMOUS_CODING | 1 | T>C |
| TP53 | chr17 | 7578526 | C | G | C135S | SNV | NON_SYNONYMOUS_CODING | 1 | C>G |
| TP53 | chr17 | 7577550 | C | A | G244V | SNV | NON_SYNONYMOUS_CODING | 1 | C>A |
| TP53 | chr17 | 7578271 | T | A | H193L | SNV | NON_SYNONYMOUS_CODING | 1 | T>A |
| TP53 | chr17 | 7577559 | G | T | S241Y | SNV | NON_SYNONYMOUS_CODING | 1 | G>T |
| TP53 | chr17 | 7579350 | A | C | F113V | SNV | NON_SYNONYMOUS_CODING | 1 | A>C |
| TP53 | chr17 | 7578554 | A | G | Y126H | SNV | NON_SYNONYMOUS_CODING | 1 | A>G |
| TP53 | chr17 | 7579528 | C | CCAT | -53Q? | Indel | FRAME_SHIFT | 1 | Indel |

| | | | | | | | | | |
|---|---|---|---|---|---|---|---|---|---|
| | | | | TG | | | | | |
| TP53 | chr17 | 7577085 | C | CT | -284? | Indel | FRAME_SHIFT | 1 | Indel |
| TP53 | chr17 | 7578535 | T | A | K132M | SNV | NON_SYNONYMOUS_CODING | 1 | T>A |
| TP53 | chr17 | 7577501 | GGA | G | -260 | Indel | FRAME_SHIFT | 1 | Indel |
| TP53 | chr17 | 7577550 | C | T | G244D | SNV | NON_SYNONYMOUS_CODING | 1 | C>T |
| SETD2 | chr3 | 47162374 | G | A | S1251L | SNV | NON_SYNONYMOUS_CODING | 1 | G>A |
| SETD2 | chr3 | 47125383 | T | A | K1963* | SNV | STOP_GAINED | 1 | T>A |
| SETD2 | chr3 | 47147605 | G | T | T1574N | SNV | NON_SYNONYMOUS_CODING | 1 | G>T |
| RYR2 | chr1 | 237880547 | G | A | R3456Q | SNV | NON_SYNONYMOUS_CODING | 1 | G>A |
| RYR2 | chr1 | 237494269 | A | T | R85* | SNV | STOP_GAINED | 1 | A>T |
| RYR2 | chr1 | 237777775 | C | T | P1781S | SNV | NON_SYNONYMOUS_CODING | 1 | C>T |
| RYR2 | chr1 | 237841410 | A | G | K2963E | SNV | NON_SYNONYMOUS_CODING | 1 | A>G |
| RYR1 | chr19 | 39001168 | A | C | K2988T | SNV | NON_SYNONYMOUS_CODING | 1 | A>C |
| RYR1 | chr19 | 39034492 | C | T | H3997Y | SNV | NON_SYNONYMOUS_CODING | 1 | C>T |
| RYR1 | chr19 | 39052033 | G | T | R4188L | SNV | NON_SYNONYMOUS_CODING | 1 | G>T |
| RPTOR | chr17 | 78796030 | C | T | T307M | SNV | NON_SYNONYMOUS_CODING | 1 | C>T |
| RPTOR | chr17 | 78897465 | C | A | P934T | SNV | NON_SYNONYMOUS_CODING | 1 | C>A |
| RPTOR | chr17 | 78921076 | A | T | N1064Y | SNV | NON_SYNONYMOUS_CODING | 1 | A>T |
| RB1 | chr13 | 48947539 | A | G | | Indel | SPLICE_SITE_ACCEPTOR | 1 | Indel |
| RB1 | chr13 | 4894169 | AT | A | -336 | Indel | FRAME_SHIFT | 1 | Indel |

| | | | | | | | | | |
|---|---|---|---|---|---|---|---|---|---|
| | | 5 | | | | | | | |
| RB1 | chr13 | 48955572 | G | C | W563S | SNV | NON_SYNONYMOUS_CODING | 1 | G>C |
| PSIP1 | chr9 | 15490113 | T | TA | -53? | Indel | FRAME_SHIFT | 2 | Indel |
| PSIP1 | chr9 | 15490045 | CCTTTT | C | -74 | Indel | FRAME_SHIFT | 3 | Indel |
| PRKDC | chr8 | 48765266 | G | C | L2324V | SNV | NON_SYNONYMOUS_CODING | 1 | G>C |
| PRKDC | chr8 | 48766699 | C | G | V2279L | SNV | NON_SYNONYMOUS_CODING | 1 | C>G |
| PRKDC | chr8 | 48805763 | C | T | A1261T | SNV | NON_SYNONYMOUS_CODING | 1 | C>T |
| PML | chr15 | 74315195 | C | A | P210Q | SNV | NON_SYNONYMOUS_CODING | 1 | C>A |
| PML | chr15 | 74337311 | G | T | G871C | SNV | NON_SYNONYMOUS_CODING | 1 | G>T |
| PML | chr15 | 74315374 | C | T | R270C | SNV | NON_SYNONYMOUS_CODING | 1 | C>T |
| PKHD1 | chr6 | 51732731 | T | A | S2555C | SNV | NON_SYNONYMOUS_CODING | 1 | T>A |
| PKHD1 | chr6 | 51771140 | T | C | | Indel | SPLICE_SITE_ACCEPTOR | 1 | Indel |
| PKHD1 | chr6 | 51930833 | C | A | R274I | SNV | NON_SYNONYMOUS_CODING | 1 | C>A |
| PIK3CA | chr3 | 178916936 | G | A | R108H | SNV | NON_SYNONYMOUS_CODING | 1 | G>A |
| PIK3CA | chr3 | 178952150 | TGAAAA | T | -1069 | Indel | FRAME_SHIFT | 1 | Indel |
| PIK3CA | chr3 | 178916713 | T | G | L34V | SNV | NON_SYNONYMOUS_CODING | 1 | T>G |
| PDE4DIP | chr1 | 144866678 | C | A | R1991L | SNV | NON_SYNONYMOUS_CODING | 1 | C>A |
| PDE4DIP | chr1 | 144916577 | T | A | E730V | SNV | NON_SYNONYMOUS_CODING | 2 | T>A |

| | | | | | | | | | |
|---|---|---|---|---|---|---|---|---|---|
| PDE4DIP | chr1 | 145021150 | T | C | D84G | SNV | NON_SYNONYMOUS_CODING | 2 | T>C |
| NFE2L2 | chr2 | 178098816 | C | G | D77H | SNV | NON_SYNONYMOUS_CODING | 1 | C>G |
| NFE2L2 | chr2 | 178098954 | C | T | G31R | SNV | NON_SYNONYMOUS_CODING | 1 | C>T |
| NFE2L2 | chr2 | 178098953 | C | A | G31V | SNV | NON_SYNONYMOUS_CODING | 1 | C>A |
| NFE2L2 | chr2 | 178098799 | T | A | E82D | SNV | NON_SYNONYMOUS_CODING | 1 | T>A |
| NFE2L2 | chr2 | 178098959 | T | A | D29V | SNV | NON_SYNONYMOUS_CODING | 1 | T>A |
| KIF1B | chr1 | 10397592 | G | A | | Indel | SPLICE_SITE_DONOR | 1 | Indel |
| KIF1B | chr1 | 10431274 | A | G | T1634A | SNV | NON_SYNONYMOUS_CODING | 1 | A>G |
| KIF1B | chr1 | 10292411 | G | A | A9T | SNV | NON_SYNONYMOUS_CODING | 1 | G>A |
| KEAP1 | chr19 | 10602866 | T | G | N238H | SNV | NON_SYNONYMOUS_CODING | 1 | T>G |
| KEAP1 | chr19 | 10602391 | T | TA | -396? | Indel | FRAME_SHIFT | 1 | Indel |
| KEAP1 | chr19 | 10600446 | C | T | R470H | SNV | NON_SYNONYMOUS_CODING | 1 | C>T |
| KDR | chr4 | 55984852 | C | A | V93L | SNV | NON_SYNONYMOUS_CODING | 1 | C>A |
| KDR | chr4 | 55979551 | CG | C | -299 | Indel | FRAME_SHIFT | 1 | Indel |
| KDR | chr4 | 55971152 | C | T | | Indel | SPLICE_SITE_ACCEPTOR | 1 | Indel |
| GRM8 | chr7 | 126883137 | C | T | R41Q | SNV | NON_SYNONYMOUS_CODING | 1 | C>T |
| GRM8 | chr7 | 126173088 | A | G | I783T | SNV | NON_SYNONYMOUS_CODING | 3 | A>G |
| CUL2 | chr10 | 35300859 | C | T | D694N | SNV | NON_SYNONYMOUS_CODING | 1 | C>T |

| | | | | | | | | | |
|---|---|---|---|---|---|---|---|---|---|
| CUL2 | chr10 | 35314123 | C | A | E560* | SNV | STOP_GAINED | 4 | C>A |
| CTNNB1 | chr3 | 41266124 | A | G | T41A | SNV | NON_SYNONYMOUS_CODING | 3 | A>G |
| CTNNB1 | chr3 | 41266100 | T | C | S33P | SNV | NON_SYNONYMOUS_CODING | 2 | T>C |
| CTNNB1 | chr3 | 41266113 | C | G | S37C | SNV | NON_SYNONYMOUS_CODING | 1 | C>G |
| CTNNB1 | chr3 | 41266104 | GAATCCATTCTGGTGCCACTACCAC | G | GIHSGATTT34G | Indel | CODON_DELETION | 1 | Indel |
| CTNNB1 | chr3 | 41266137 | C | T | S45F | SNV | NON_SYNONYMOUS_CODING | 2 | C>T |
| CTNNB1 | chr3 | 41266097 | G | C | D32H | SNV | NON_SYNONYMOUS_CODING | 3 | G>C |
| CTNNB1 | chr3 | 41266097 | G | T | D32Y | SNV | NON_SYNONYMOUS_CODING | 1 | G>T |
| CTNNB1 | chr3 | 41266098 | A | G | D32G | SNV | NON_SYNONYMOUS_CODING | 1 | A>G |
| CSMD3 | chr8 | 113504810 | G | T | A1729D | SNV | NON_SYNONYMOUS_CODING | 1 | G>T |
| CSMD3 | chr8 | 114448950 | G | C | T45R | SNV | NON_SYNONYMOUS_CODING | 1 | G>C |
| CSMD3 | chr8 | 113702205 | G | A | R683C | SNV | NON_SYNONYMOUS_CODING | 1 | G>A |
| CSMD3 | chr8 | 113933964 | G | A | Q509* | SNV | STOP_GAINED | 1 | G>A |
| CSMD3 | chr8 | 113871409 | A | G | C574R | SNV | NON_SYNONYMOUS_CODING | 2 | A>G |

| | | | | | | | | | |
|---|---|---|---|---|---|---|---|---|---|
| CSMD3 | chr8 | 113395877 | A | T | F1984I | SNV | NON_SYNONYMOUS_CODING | 1 | A>T |
| AXIN1 | chr16 | 396446 | C | A | E194* | SNV | STOP_GAINED | 1 | C>A |
| AXIN1 | chr16 | 343603 | G | A | Q691* | SNV | STOP_GAINED | 1 | G>A |
| AXIN1 | chr16 | 347107 | C | T | W635* | SNV | STOP_GAINED | 1 | C>T |
| AXIN1 | chr16 | 359972 | C | T | | Indel | SPLICE_SITE_DONOR | 1 | Indel |
| AXIN1 | chr16 | 354428 | AC | A | -377 | Indel | FRAME_SHIFT | 1 | Indel |
| ASXL1 | chr20 | 31020700 | A | T | M333L | SNV | NON_SYNONYMOUS_CODING | 1 | A>T |
| ASXL1 | chr20 | 31024764 | C | A | P1417T | SNV | NON_SYNONYMOUS_CODING | 1 | C>A |
| ASXL1 | chr20 | 31021502 | T | A | S501T | SNV | NON_SYNONYMOUS_CODING | 1 | T>A |
| ARID2 | chr12 | 46230748 | G | T | D333Y | SNV | NON_SYNONYMOUS_CODING | 1 | G>T |
| ARID2 | chr12 | 46245837 | A | T | K1311* | SNV | STOP_GAINED | 1 | A>T |
| ARID2 | chr12 | 46244453 | A | AC | -850? | Indel | FRAME_SHIFT | 1 | Indel |
| ARID2 | chr12 | 46233207 | C | CT | -476? | Indel | FRAME_SHIFT | 1 | Indel |
| ARID2 | chr12 | 46244262 | G | T | G786C | SNV | NON_SYNONYMOUS_CODING | 1 | G>T |
| ARID2 | chr12 | 46245288 | C | T | Q1128* | SNV | STOP_GAINED | 1 | C>T |
| ARID2 | chr12 | 46231135 | C | CTCAA | -352S? | Indel | FRAME_SHIFT | 1 | Indel |
| ARID2 | chr12 | 46246299 | CCAAGTGTA | C | -1465 | Indel | FRAME_SHIFT | 1 | Indel |
| ARID2 | chr12 | 46233219 | G | T | E480* | SNV | STOP_GAINED | 1 | G>T |
| ARID1A | chr1 | 27097658 | G | T | G1083C | SNV | NON_SYNONYMOUS_CODING | 1 | G>T |
| ARID1A | chr1 | 27107059 | G | T | E2224* | SNV | STOP_GAINED | 1 | G>T |
| ARID1A | chr1 | 27094279 | A | T | | Indel | SPLICE_SITE_ACCEPTOR | 1 | Indel |
| ARID1A | chr1 | 27107023 | C | T | Q2212* | SNV | STOP_GAINED | 1 | C>T |
| ARID1A | chr1 | 2709994 | C | T | R1276* | SNV | STOP_GAINED | 2 | C>T |

| | | | | | | | | | |
|---|---|---|---|---|---|---|---|---|---|
| | | 7 | | | | | | | |
| ARID1A | chr1 | 27106803 | AC | A | -2139 | Indel | FRAME_SHIFT | 2 | Indel |
| ARID1A | chr1 | 27105841 | C | T | Q1818* | SNV | STOP_GAINED | 1 | C>T |
| ARID1A | chr1 | 27105756 | C | CT | -1790? | Indel | FRAME_SHIFT | 1 | Indel |
| ARID1A | chr1 | 27105774 | GC | G | -1796 | Indel | FRAME_SHIFT | 1 | Indel |
| APOB | chr2 | 21232728 | T | A | R2338* | SNV | STOP_GAINED | 1 | T>A |
| APOB | chr2 | 21258549 | C | A | S242I | SNV | NON_SYNONYMOUS_CODING | 1 | C>A |
| APOB | chr2 | 21233884 | ATG | A | -1952 | Indel | FRAME_SHIFT | 1 | Indel |
| APOB | chr2 | 21224995 | AGTAAAGTTAGAGGCACTGAC | A | -4427 | Indel | FRAME_SHIFT | 1 | Indel |
| APOB | chr2 | 21234944 | C | T | R1599H | SNV | NON_SYNONYMOUS_CODING | 1 | C>T |
| APOB | chr2 | 21235067 | G | C | T1558S | SNV | NON_SYNONYMOUS_CODING | 1 | G>C |
| APOB | chr2 | 21232800 | GAATGTCATTTATTCTTTCAAATGAAAT | G | ISFERINDI2305- | Indel | CODON_DELETION | 1 | Indel |
| ACVR2A | chr2 | 148683685 | TA | T | -435 | Indel | FRAME_SHIFT | 3 | Indel |
| ACVR2A | chr2 | 148680560 | A | G | M366V | SNV | NON_SYNONYMOUS_CODING | 1 | A>G |

**Table S10. Recurrently mutated genes in Thai iCCA**

| Gene | Chr. | POS | REF | ALT | AA Change | Mutation.Type | Effect | var.frq | snp.alt |
|---|---|---|---|---|---|---|---|---|---|
| USP9X | chrX | 41075689 | G | A | D1957N | SNV | NON_SYNONYMOUS_CODING | 1 | G>A |
| USP9X | chrX | 41010177 | C | T | R544C | SNV | NON_SYNONYMOUS_CODING | 1 | C>T |
| USP9X | chrX | 41057909 | ATTAC | A | -1504 | Indel | FRAME_SHIFT | 1 | Indel |
| USP9X | chrX | 41073850 | T | G | L1740R | SNV | NON_SYNONYMOUS_CODING | 1 | T>G |
| TP53 | chr17 | 7590693 | A | T | | Indel | SPLICE_SITE_DONOR | 1 | Indel |
| TP53 | chr17 | 7577524 | T | C | T253A | SNV | NON_SYNONYMOUS_CODING | 1 | T>C |
| TP53 | chr17 | 7577022 | G | A | R306* | SNV | STOP_GAINED | 2 | G>A |
| TP53 | chr17 | 7577534 | C | A | R249S | SNV | NON_SYNONYMOUS_CODING | 7 | C>A |
| TP53 | chr17 | 7577566 | T | C | N239D | SNV | NON_SYNONYMOUS_CODING | 1 | T>C |
| TP53 | chr17 | 7577548 | C | T | G245S | SNV | NON_SYNONYMOUS_CODING | 1 | C>T |
| TP53 | chr17 | 7578406 | C | T | R175H | SNV | NON_SYNONYMOUS_CODING | 2 | C>T |
| TP53 | chr17 | 7590692 | T | TA | | Indel | SPLICE_SITE_DONOR | 1 | Indel |
| TP53 | chr17 | 7577118 | C | A | V274F | SNV | NON_SYNONYMOUS_CODING | 1 | C>A |
| TP53 | chr17 | 7577094 | G | A | R282W | SNV | NON_SYNONYMOUS_CODING | 2 | G>A |
| TP53 | chr17 | 7577112 | C | G | A276P | SNV | NON_SYNONYMOUS_CODING | 1 | C>G |
| TP53 | chr17 | 7577120 | C | T | R273H | SNV | NON_SYNONYMOUS_CODING | 3 | C>T |
| TP53 | chr17 | 7579554 | GCATCAAATCATCCA | G | -40 | Indel | FRAME_SHIFT | 1 | Indel |
| TP53 | chr17 | 7578496 | A | T | L145Q | SNV | NON_SYNONYMOUS_CODING | 1 | A>T |
| TP53 | chr17 | 7578518 | C | G | A138P | SNV | NON_SYNONYMOUS_CODING | 1 | C>G |
| TP53 | chr17 | 7578520 | A | T | L137Q | SNV | NON_SYNONYMOUS_CODING | 1 | A>T |

| | | | | | | | | | |
|---|---|---|---|---|---|---|---|---|---|
| TP53 | chr17 | 7577547 | C | A | G245V | SNV | NON_SYNONYMOUS_CODING | 1 | C>A |
| TP53 | chr17 | 7578431 | G | A | Q167* | SNV | STOP_GAINED | 1 | G>A |
| TP53 | chr17 | 7577538 | C | T | R248Q | SNV | NON_SYNONYMOUS_CODING | 3 | C>T |
| TP53 | chr17 | 7577599 | C | CA | -227? | Indel | FRAME_SHIFT | 1 | Indel |
| TP53 | chr17 | 7577115 | A | C | C275G | SNV | NON_SYNONYMOUS_CODING | 1 | A>C |
| TP53 | chr17 | 7577057 | TC | T | -294 | Indel | FRAME_SHIFT | 1 | Indel |
| TP53 | chr17 | 7578176 | C | T | | Indel | SPLICE_SITE_DONOR | 1 | Indel |
| TP53 | chr17 | 7578413 | C | A | V173L | SNV | NON_SYNONYMOUS_CODING | 1 | C>A |
| TP53 | chr17 | 7579359 | G | GGAAACCGT | -109TV? | Indel | FRAME_SHIFT | 1 | Indel |
| TP53 | chr17 | 7576870 | C | A | E326* | SNV | STOP_GAINED | 1 | C>A |
| TP53 | chr17 | 7578212 | G | A | R213* | SNV | STOP_GAINED | 1 | G>A |
| TP53 | chr17 | 7578410 | T | A | R174W | SNV | NON_SYNONYMOUS_CODING | 1 | T>A |
| TP53 | chr17 | 7577124 | C | T | V272M | SNV | NON_SYNONYMOUS_CODING | 1 | C>T |
| TP53 | chr17 | 7578263 | G | A | R196* | SNV | STOP_GAINED | 1 | G>A |
| TP53 | chr17 | 7576896 | TG | T | -317 | Indel | FRAME_SHIFT | 1 | Indel |
| TP53 | chr17 | 7579343 | T | C | H115R | SNV | NON_SYNONYMOUS_CODING | 1 | T>C |
| TP53 | chr17 | 7579373 | C | A | G105V | SNV | NON_SYNONYMOUS_CODING | 1 | C>A |
| TP53 | chr17 | 7579854 | GA | G | -20 | Indel | FRAME_SHIFT | 1 | Indel |
| TP53 | chr17 | 7574018 | G | A | R337C | SNV | NON_SYNONYMOUS_CODING | 1 | G>A |
| TP53 | chr17 | 7578262 | C | G | R196P | SNV | NON_SYNONYMOUS_CODING | 1 | C>G |
| TP53 | chr17 | 7577099 | C | G | R280T | SNV | NON_SYNONYMOUS_CODING | 1 | C>G |
| TGFBR1 | chr9 | 101904946 | G | A | G316S | SNV | NON_SYNONYMOUS_CODING | 1 | G>A |
| TGFBR1 | chr9 | 101891287 | C | T | P83L | SNV | NON_SYNONYMOUS_CODING | 1 | C>T |
| TGFBR1 | chr9 | 101891136 | GCGTTAC | G | ALQ33E | Indel | CODON_CHANGE_PLUS_CODON_DELETION | 1 | Indel |

| | | | | | | | | | |
|---|---|---|---|---|---|---|---|---|---|
| SYN E1 | chr6 | 152651967 | A | C | L4618R | SNV | NON_SYNONYMOUS _CODING | 1 | A> C |
| SYN E1 | chr6 | 152651058 | T | C | Q4921R | SNV | NON_SYNONYMOUS _CODING | 1 | T> C |
| SYN E1 | chr6 | 152527476 | G | A | R7616 W | SNV | NON_SYNONYMOUS _CODING | 1 | G> A |
| SYN E1 | chr6 | 152485406 | C | T | M7894I | SNV | NON_SYNONYMOUS _CODING | 1 | C> T |
| SYN E1 | chr6 | 152555825 | C | T | R6836H | SNV | NON_SYNONYMOUS _CODING | 1 | C> T |
| SYN E1 | chr6 | 152599315 | T | G | E6161A | SNV | NON_SYNONYMOUS _CODING | 1 | T> G |
| SYN E1 | chr6 | 152765644 | C | A | E1247* | SNV | STOP_GAINED | 1 | C> A |
| STK1 1 | chr19 | 1207033 | AA GC G | A | -41 | Indel | FRAME_SHIFT | 1 | In del |
| STK1 1 | chr19 | 1221995 | C | T | R304W | SNV | NON_SYNONYMOUS _CODING | 1 | C> T |
| STK1 1 | chr19 | 1220487 | G | A | D194N | SNV | NON_SYNONYMOUS _CODING | 1 | G> A |
| STK1 1 | chr19 | 1207054 | A | T | K48* | SNV | STOP_GAINED | 1 | A> T |
| STA G1 | chr3 | 136191310 | C | G | D384H | SNV | NON_SYNONYMOUS _CODING | 1 | C> G |
| STA G1 | chr3 | 136221548 | T | A | Q250H | SNV | NON_SYNONYMOUS _CODING | 1 | T> A |
| STA G1 | chr3 | 136342005 | G | A | R39C | SNV | NON_SYNONYMOUS _CODING | 1 | G> A |
| SRC | chr20 | 36031729 | G | T | E526* | SNV | STOP_GAINED | 1 | G> T |
| SRC | chr20 | 36014513 | G | A | E96K | SNV | NON_SYNONYMOUS _CODING | 1 | G> A |
| SRC | chr20 | 36031750 | G | A | E533K | SNV | NON_SYNONYMOUS _CODING | 1 | G> A |
| SMA D4 | chr18 | 48593406 | G | A | G386D | SNV | NON_SYNONYMOUS _CODING | 1 | G> A |
| SMA D4 | chr18 | 48591880 | TT AC TG TT GA | T | VTVD3 48V | Indel | CODON_DELETION | 1 | In del |
| SMA D4 | chr18 | 48591898 | T | G | V354G | SNV | NON_SYNONYMOUS _CODING | 1 | T> G |
| SMA D4 | chr18 | 48591918 | C | T | R361C | SNV | NON_SYNONYMOUS _CODING | 4 | C> T |
| SMA D4 | chr18 | 48604764 | T | TA | -529? | Indel | FRAME_SHIFT | 1 | In del |
| SMA D4 | chr18 | 48591910 | GA GG AG | G | GGD35 8G | Indel | CODON_CHANGE_PL US_CODON_DELETIO N | 1 | In del |

| | | | | | | | | | |
|---|---|---|---|---|---|---|---|---|---|
| | | | A | | | | | | |
| SMA D4 | chr18 | 48604750 | G | T | W524C | SNV | NON_SYNONYMOUS _CODING | 1 | G> T |
| SMA D4 | chr18 | 48584497 | C | T | Q224* | SNV | STOP_GAINED | 1 | C> T |
| SMA D4 | chr18 | 48575219 | C | G | S138* | SNV | STOP_GAINED | 1 | C> G |
| SMA D4 | chr18 | 48593531 | A | T | K428* | SNV | STOP_GAINED | 1 | A> T |
| SMA D4 | chr18 | 48584593 | C | T | Q256* | SNV | STOP_GAINED | 1 | C> T |
| SMA D4 | chr18 | 48573528 | A | T | R38* | SNV | STOP_GAINED | 1 | A> T |
| SMA D4 | chr18 | 48584513 | TG | T | -229 | Indel | FRAME_SHIFT | 1 | In del |
| SCN5 A | chr3 | 38608045 | C | T | R1232Q | SNV | NON_SYNONYMOUS _CODING | 1 | C> T |
| SCN5 A | chr3 | 38640457 | G | A | R659W | SNV | NON_SYNONYMOUS _CODING | 1 | G> A |
| SCN5 A | chr3 | 38603943 | C | T | R1309H | SNV | NON_SYNONYMOUS _CODING | 1 | C> T |
| SCN5 A | chr3 | 38647567 | C | T | V405M | SNV | NON_SYNONYMOUS _CODING | 1 | C> T |
| RYR2 | chr1 | 237969498 | T | C | F4744S | SNV | NON_SYNONYMOUS _CODING | 1 | T> C |
| RYR2 | chr1 | 237804248 | G | A | M2387I | SNV | NON_SYNONYMOUS _CODING | 1 | G> A |
| RYR2 | chr1 | 237850754 | GC | G | -3004 | Indel | FRAME_SHIFT | 1 | In del |
| RYR2 | chr1 | 237868562 | C | T | P3165S | SNV | NON_SYNONYMOUS _CODING | 3 | C> T |
| RYR2 | chr1 | 237670041 | G | C | R880T | SNV | NON_SYNONYMOUS _CODING | 1 | G> C |
| RYR2 | chr1 | 237580409 | G | C | E276D | SNV | NON_SYNONYMOUS _CODING | 1 | G> C |
| RYR2 | chr1 | 237755097 | G | T | E1405* | SNV | STOP_GAINED | 1 | G> T |
| RYR1 | chr19 | 38934854 | G | T | V164F | SNV | NON_SYNONYMOUS _CODING | 1 | G> T |
| RYR1 | chr19 | 38979850 | CA | C | -1861 | Indel | FRAME_SHIFT | 1 | In del |
| RYR1 | chr19 | 38949797 | C | T | R727C | SNV | NON_SYNONYMOUS _CODING | 1 | C> T |
| RYR1 | chr19 | 38998374 | G | A | D2947N | SNV | NON_SYNONYMOUS _CODING | 1 | G> A |
| RUN X1T1 | chr8 | 92998513 | C | T | R384Q | SNV | NON_SYNONYMOUS _CODING | 1 | C> T |
| RUN X1T1 | chr8 | 93029565 | G | A | P50S | SNV | NON_SYNONYMOUS _CODING | 1 | G> A |
| RUN | chr8 | 93004062 | G | A | R277* | SNV | STOP_GAINED | 1 | G> |

| | | | | | | | | | |
|---|---|---|---|---|---|---|---|---|---|
| X1T1 | | | | | | | | | A |
| RUN X1T1 | chr8 | 92982983 | G | A | T492I | SNV | NON_SYNONYMOUS _CODING | 1 | G> A |
| RNF4 3 | chr17 | 56436140 | G | A | Q333* | SNV | STOP_GAINED | 1 | G> A |
| RNF4 3 | chr17 | 56492739 | C | T | G67D | SNV | NON_SYNONYMOUS _CODING | 1 | C> T |
| RNF4 3 | chr17 | 56440741 | C | T | W159* | SNV | STOP_GAINED | 1 | C> T |
| RNF4 3 | chr17 | 56438222 | AC CC CG | A | -256 | Indel | FRAME_SHIFT | 1 | In del |
| RNF4 3 | chr17 | 56437534 | G | T | P310T | SNV | NON_SYNONYMOUS _CODING | 1 | G> T |
| PTPR D | chr9 | 8636761 | C | G | D50H | SNV | NON_SYNONYMOUS _CODING | 1 | C> G |
| PTPR D | chr9 | 8633407 | G | A | R88W | SNV | NON_SYNONYMOUS _CODING | 1 | G> A |
| PTPR D | chr9 | 8460493 | C | T | E1265K | SNV | NON_SYNONYMOUS _CODING | 1 | C> T |
| PSIP1 | chr9 | 15490045 | CC TTT T | C | -74 | Indel | FRAME_SHIFT | 3 | In del |
| PSIP1 | chr9 | 15490113 | TA | T | -53 | Indel | FRAME_SHIFT | 1 | In del |
| PSIP1 | chr9 | 15469290 | T | A | K360* | SNV | STOP_GAINED | 1 | T> A |
| PSIP1 | chr9 | 15506572 | C | A | G46* | SNV | STOP_GAINED | 1 | C> A |
| PSIP1 | chr9 | 15472749 | C | A | | Indel | SPLICE_SITE_ACCEP TOR | 1 | In del |
| PRK DC | chr8 | 48830924 | C | T | W813* | SNV | STOP_GAINED | 1 | C> T |
| PRK DC | chr8 | 48749959 | C | T | W2524* | SNV | STOP_GAINED | 1 | C> T |
| PRK DC | chr8 | 48713478 | C | T | G3330D | SNV | NON_SYNONYMOUS _CODING | 1 | C> T |
| PRK DC | chr8 | 48817428 | C | G | | Indel | SPLICE_SITE_DONOR | 1 | In del |
| PRK DC | chr8 | 48848395 | C | A | Q448H | SNV | NON_SYNONYMOUS _CODING | 1 | C> A |
| PRK DC | chr8 | 48811031 | G | A | R1155* | SNV | STOP_GAINED | 1 | G> A |
| PIK3 CA | chr3 | 178936091 | G | A | E545K | SNV | NON_SYNONYMOUS _CODING | 1 | G> A |
| PIK3 CA | chr3 | 178921553 | T | A | N345K | SNV | NON_SYNONYMOUS _CODING | 2 | T> A |
| PIK3 CA | chr3 | 178943780 | T | G | I816S | SNV | NON_SYNONYMOUS _CODING | 1 | T> G |
| PIK3 CA | chr3 | 178952085 | A | G | H1047R | SNV | NON_SYNONYMOUS _CODING | 1 | A> G |

| | | | | | | | | | |
|---|---|---|---|---|---|---|---|---|---|
| PBR M1 | chr3 | 52584801 | CT AC CA TA | C | -1545 | Indel | FRAME_SHIFT | 1 | In del |
| PBR M1 | chr3 | 52682399 | A | AT | -258? | Indel | FRAME_SHIFT | 2 | In del |
| PBR M1 | chr3 | 52643328 | C | A | | Indel | SPLICE_SITE_DONOR | 1 | In del |
| PBR M1 | chr3 | 52682407 | CG AG | C | L255- | Indel | CODON_DELETION | 1 | In del |
| NF1 | chr17 | 29684326 | C | T | R2637* | SNV | STOP_GAINED | 1 | C> T |
| NF1 | chr17 | 29554310 | G | C | | Indel | SPLICE_SITE_DONOR | 1 | In del |
| NF1 | chr17 | 29486094 | G | GA | -91? | Indel | FRAME_SHIFT | 1 | In del |
| NF1 | chr17 | 29652920 | G | T | E1640* | SNV | STOP_GAINED | 1 | G> T |
| NF1 | chr17 | 29657433 | C | CT | -1910? | Indel | FRAME_SHIFT | 1 | In del |
| NF1 | chr17 | 29663350 | G | A | | Indel | SPLICE_SITE_ACCEP TOR | 1 | In del |
| MYH 11 | chr16 | 15815482 | C | T | E1466K | SNV | NON_SYNONYMOUS _CODING | 1 | C> T |
| MYH 11 | chr16 | 15932094 | G | A | Q6* | SNV | STOP_GAINED | 1 | G> A |
| MYH 11 | chr16 | 15844174 | C | T | G634S | SNV | NON_SYNONYMOUS _CODING | 1 | C> T |
| MAP 2K4 | chr17 | 12028657 | G | A | R298H | SNV | NON_SYNONYMOUS _CODING | 1 | G> A |
| MAP 2K4 | chr17 | 12028638 | C | T | R292* | SNV | STOP_GAINED | 1 | C> T |
| MAP 2K4 | chr17 | 12028660 | C | T | S299F | SNV | NON_SYNONYMOUS _CODING | 1 | C> T |
| MAP 2K4 | chr17 | 12016634 | C | G | S268C | SNV | NON_SYNONYMOUS _CODING | 1 | C> G |
| MAP 2K4 | chr17 | 11984802 | CA AC AA AA TG | C | NKM12 8- | Indel | CODON_DELETION | 1 | In del |
| LRP1 B | chr2 | 141143498 | T | A | N3499Y | SNV | NON_SYNONYMOUS _CODING | 1 | T> A |
| LRP1 B | chr2 | 141122280 | TC A | T | -3693 | Indel | FRAME_SHIFT | 1 | In del |
| LRP1 B | chr2 | 141214083 | T | C | T3302A | SNV | NON_SYNONYMOUS _CODING | 1 | T> C |
| LRP1 B | chr2 | 140995804 | G | T | P4493T | SNV | NON_SYNONYMOUS _CODING | 1 | G> T |
| LRP1 B | chr2 | 141253256 | T | C | E2971G | SNV | NON_SYNONYMOUS _CODING | 1 | T> C |
| LRP1 B | chr2 | 141264371 | G | A | Q2839* | SNV | STOP_GAINED | 1 | G> A |

| | | | | | | | | | |
|---|---|---|---|---|---|---|---|---|---|
| LRP1B | chr2 | 141643848 | G | C | L1275V | SNV | NON_SYNONYMOUS_CODING | 1 | G>C |
| LRP1B | chr2 | 141243094 | C | A | | Indel | SPLICE_SITE_ACCEPTOR | 1 | Indel |
| LRP1B | chr2 | 141473551 | G | C | P2005R | SNV | NON_SYNONYMOUS_CODING | 1 | G>C |
| LRP1B | chr2 | 141299411 | G | T | L2442I | SNV | NON_SYNONYMOUS_CODING | 1 | G>T |
| KRAS | chr12 | 25398284 | C | T | G12D | SNV | NON_SYNONYMOUS_CODING | 6 | C>T |
| KRAS | chr12 | 25398284 | C | A | G12V | SNV | NON_SYNONYMOUS_CODING | 3 | C>A |
| KRAS | chr12 | 25398285 | C | T | G12S | SNV | NON_SYNONYMOUS_CODING | 2 | C>T |
| KRAS | chr12 | 25378562 | C | T | A146T | SNV | NON_SYNONYMOUS_CODING | 1 | C>T |
| KRAS | chr12 | 25380275 | T | G | Q61H | SNV | NON_SYNONYMOUS_CODING | 1 | T>G |
| KRAS | chr12 | 25398282 | C | A | G13C | SNV | NON_SYNONYMOUS_CODING | 1 | C>A |
| KRAS | chr12 | 25398285 | C | A | G12C | SNV | NON_SYNONYMOUS_CODING | 2 | C>A |
| KRAS | chr12 | 25398262 | C | A | L19F | SNV | NON_SYNONYMOUS_CODING | 1 | C>A |
| KRAS | chr12 | 25398281 | C | T | G13D | SNV | NON_SYNONYMOUS_CODING | 1 | C>T |
| KMT2C | chr7 | 151874120 | GAGAACCA | G | -2804 | Indel | FRAME_SHIFT | 1 | Indel |
| KMT2C | chr7 | 152012245 | G | A | R190* | SNV | STOP_GAINED | 1 | G>A |
| KMT2C | chr7 | 151949801 | C | G | | Indel | SPLICE_SITE_ACCEPTOR | 1 | Indel |
| KMT2C | chr7 | 151856055 | G | A | Q3855* | SNV | STOP_GAINED | 1 | G>A |
| KMT2C | chr7 | 151859201 | C | A | | Indel | SPLICE_SITE_DONOR | 1 | Indel |
| KMT2C | chr7 | 151851531 | C | T | | Indel | SPLICE_SITE_ACCEPTOR | 2 | Indel |
| KMT2C | chr7 | 151947040 | T | C | | Indel | SPLICE_SITE_ACCEPTOR | 1 | Indel |
| KMT2C | chr7 | 151879136 | G | A | Q1937* | SNV | STOP_GAINED | 1 | G>A |
| KMT2C | chr7 | 151845261 | C | T | R4641Q | SNV | NON_SYNONYMOUS_CODING | 1 | C>T |
| KMT2C | chr7 | 151884352 | A | C | F1668C | SNV | NON_SYNONYMOUS_CODING | 1 | A>C |
| KMT2C | chr7 | 151949138 | C | A | E503* | SNV | STOP_GAINED | 1 | C>A |

| | | | | | | | | | |
|---|---|---|---|---|---|---|---|---|---|
| KAT6 B | chr10 | 76739051 | G | C | E729Q | SNV | NON_SYNONYMOUS _CODING | 1 | G> C |
| KAT6 B | chr10 | 76788859 | C | T | A1426V | SNV | NON_SYNONYMOUS _CODING | 1 | C> T |
| KAT6 B | chr10 | 76789762 | A | T | N1727I | SNV | NON_SYNONYMOUS _CODING | 1 | A> T |
| KAT6 B | chr10 | 76781905 | G | GGA A | -1097E | Indel | CODON_INSERTION | 1 | In del |
| KAT6 B | chr10 | 76781905 | GG AA | G | E1097- | Indel | CODON_DELETION | 1 | In del |
| GNA S | chr20 | 57484421 | G | A | R844H | SNV | NON_SYNONYMOUS _CODING | 6 | G> A |
| GNA S | chr20 | 57484420 | C | T | R844C | SNV | NON_SYNONYMOUS _CODING | 9 | C> T |
| GNA S | chr20 | 57428925 | C | T | P202L | SNV | NON_SYNONYMOUS _CODING | 1 | C> T |
| FN1 | chr2 | 216271877 | G | C | Q896E | SNV | NON_SYNONYMOUS _CODING | 1 | G> C |
| FN1 | chr2 | 216284088 | T | C | T566A | SNV | NON_SYNONYMOUS _CODING | 1 | T> C |
| FN1 | chr2 | 216286852 | C | G | R503P | SNV | NON_SYNONYMOUS _CODING | 1 | C> G |
| FN1 | chr2 | 216274397 | C | T | E730K | SNV | NON_SYNONYMOUS _CODING | 1 | C> T |
| FGFR 2 | chr10 | 123247514 | C | A | K660N | SNV | NON_SYNONYMOUS _CODING | 1 | C> A |
| FGFR 2 | chr10 | 123247516 | T | C | K660E | SNV | NON_SYNONYMOUS _CODING | 1 | T> C |
| FGFR 2 | chr10 | 123258034 | A | C | N550K | SNV | NON_SYNONYMOUS _CODING | 1 | A> C |
| FAN CI | chr15 | 89816675 | C | A | T317K | SNV | NON_SYNONYMOUS _CODING | 4 | C> A |
| FAN CI | chr15 | 89816698 | C | A | Q325K | SNV | NON_SYNONYMOUS _CODING | 4 | C> A |
| ERB B4 | chr2 | 212566818 | C | T | A455T | SNV | NON_SYNONYMOUS _CODING | 1 | C> T |
| ERB B4 | chr2 | 212251768 | AG CA CC CT GT | A | AQGA1 094A | Indel | CODON_CHANGE_PL US_CODON_DELETIO N | 1 | In del |
| ERB B4 | chr2 | 212426794 | C | A | S774I | SNV | NON_SYNONYMOUS _CODING | 1 | C> A |
| ERB B4 | chr2 | 212570064 | G | A | R393W | SNV | NON_SYNONYMOUS _CODING | 1 | G> A |
| ERB B3 | chr12 | 56481696 | T | C | F244S | SNV | NON_SYNONYMOUS _CODING | 1 | T> C |
| ERB B3 | chr12 | 56481922 | G | A | G284R | SNV | NON_SYNONYMOUS _CODING | 3 | G> A |

| | | | | | | | | | |
|---|---|---|---|---|---|---|---|---|---|
| ERB B3 | chr12 | 56478854 | G | A | V104M | SNV | NON_SYNONYMOUS _CODING | 1 | G> A |
| ERB B3 | chr12 | 56482537 | G | A | E332K | SNV | NON_SYNONYMOUS _CODING | 1 | G> A |
| ERB B3 | chr12 | 56482341 | G | T | D297Y | SNV | NON_SYNONYMOUS _CODING | 1 | G> T |
| ERB B3 | chr12 | 56488249 | C | T | P590S | SNV | NON_SYNONYMOUS _CODING | 1 | C> T |
| ERB B2 | chr17 | 37868208 | C | T | S310F | SNV | NON_SYNONYMOUS _CODING | 6 | C> T |
| ERB B2 | chr17 | 37880220 | T | C | L755S | SNV | NON_SYNONYMOUS _CODING | 1 | T> C |
| ERB B2 | chr17 | 37868208 | C | A | S310Y | SNV | NON_SYNONYMOUS _CODING | 3 | C> A |
| EPH A6 | chr3 | 97124093 | A | G | D569G | SNV | NON_SYNONYMOUS _CODING | 1 | A> G |
| EPH A6 | chr3 | 97202892 | T | C | V730A | SNV | NON_SYNONYMOUS _CODING | 1 | T> C |
| EPH A6 | chr3 | 97251304 | G | A | R768K | SNV | NON_SYNONYMOUS _CODING | 1 | G> A |
| EP40 0 | chr12 | 132530375 | G | A | R2413Q | SNV | NON_SYNONYMOUS _CODING | 1 | G> A |
| EP40 0 | chr12 | 132522288 | C | T | R2041* | SNV | STOP_GAINED | 1 | C> T |
| EP40 0 | chr12 | 132512637 | C | A | P1765T | SNV | NON_SYNONYMOUS _CODING | 1 | C> A |
| CSM D3 | chr8 | 114326810 | A | T | F131I | SNV | NON_SYNONYMOUS _CODING | 1 | A> T |
| CSM D3 | chr8 | 113304844 | C | T | G2904R | SNV | NON_SYNONYMOUS _CODING | 1 | C> T |
| CSM D3 | chr8 | 113266510 | C | G | G3361A | SNV | NON_SYNONYMOUS _CODING | 1 | C> G |
| CSM D3 | chr8 | 113348949 | AG | A | -2317 | Indel | FRAME_SHIFT | 1 | In del |
| CSM D3 | chr8 | 113702263 | AC TTT | A | -662 | Indel | FRAME_SHIFT | 1 | In del |
| CSM D3 | chr8 | 113871376 | T | G | K585Q | SNV | NON_SYNONYMOUS _CODING | 1 | T> G |
| CSM D3 | chr8 | 113871409 | A | G | C574R | SNV | NON_SYNONYMOUS _CODING | 2 | A> G |
| CSM D3 | chr8 | 113308152 | G | A | Q2842* | SNV | STOP_GAINED | 1 | G> A |
| CSM D3 | chr8 | 113267612 | C | T | E3303K | SNV | NON_SYNONYMOUS _CODING | 1 | C> T |
| COL3 A1 | chr2 | 189861915 | C | T | R596* | SNV | STOP_GAINED | 1 | C> T |
| COL3 A1 | chr2 | 189854843 | C | T | R238* | SNV | STOP_GAINED | 1 | C> T |
| COL3 A1 | chr2 | 189870103 | G | A | G987S | SNV | NON_SYNONYMOUS _CODING | 1 | G> A |

| | | | | | | | | | |
|---|---|---|---|---|---|---|---|---|---|
| CDK N2A | chr9 | 21971203 | AT GA C | A | -106 | Indel | FRAME_SHIFT | 1 | In del |
| CDK N2A | chr9 | 21971111 | G | A | A138V | SNV | NON_SYNONYMOUS _CODING | 2 | G> A |
| CDK N2A | chr9 | 21971099 | G | A | P142L | SNV | NON_SYNONYMOUS _CODING | 1 | G> A |
| BRC A2 | chr13 | 32905055 | G | T | | Indel | SPLICE_SITE_ACCEP TOR | 1 | In del |
| BRC A2 | chr13 | 32929036 | T | G | F2349C | SNV | NON_SYNONYMOUS _CODING | 1 | T> G |
| BRC A2 | chr13 | 32912621 | A | C | N1377H | SNV | NON_SYNONYMOUS _CODING | 1 | A> C |
| BCO RL1 | chrX | 129159070 | G | A | R1265H | SNV | NON_SYNONYMOUS _CODING | 1 | G> A |
| BCO RL1 | chrX | 129155104 | C | T | R1196* | SNV | STOP_GAINED | 1 | C> T |
| BCO RL1 | chrX | 129149383 | G | T | E879* | SNV | STOP_GAINED | 1 | G> T |
| BAP1 | chr3 | 52441413 | A | T | | Indel | SPLICE_SITE_DONOR | 1 | In del |
| BAP1 | chr3 | 52441235 | G | A | R179W | SNV | NON_SYNONYMOUS _CODING | 1 | G> A |
| BAP1 | chr3 | 52440906 | C | A | E200* | SNV | STOP_GAINED | 1 | C> A |
| BAP1 | chr3 | 52439796 | C | CAA | -305? | Indel | FRAME_SHIFT | 1 | In del |
| BAI3 | chr6 | 70037710 | A | C | | Indel | SPLICE_SITE_ACCEP TOR | 1 | In del |
| BAI3 | chr6 | 70049284 | C | T | A1116V | SNV | NON_SYNONYMOUS _CODING | 1 | C> T |
| BAI3 | chr6 | 70071276 | G | C | E1371Q | SNV | NON_SYNONYMOUS _CODING | 1 | G> C |
| BAI3 | chr6 | 69348853 | G | GA | -96? | Indel | FRAME_SHIFT | 1 | In del |
| ATR X | chrX | 76939697 | C | A | E351* | SNV | STOP_GAINED | 1 | C> A |
| ATR X | chrX | 76849277 | T | C | D2000G | SNV | NON_SYNONYMOUS _CODING | 1 | T> C |
| ATR X | chrX | 76972716 | T | G | S9R | SNV | NON_SYNONYMOUS _CODING | 1 | T> G |
| ATM | chr11 | 108160464 | G | A | G1458R | SNV | NON_SYNONYMOUS _CODING | 1 | G> A |
| ATM | chr11 | 108198387 | A | T | K2331* | SNV | STOP_GAINED | 1 | A> T |
| ATM | chr11 | 108186766 | T | C | W2042 R | SNV | NON_SYNONYMOUS _CODING | 1 | T> C |
| ATM | chr11 | 108121684 | G | T | E498* | SNV | STOP_GAINED | 1 | G> T |
| ATM | chr11 | 108216476 | C | CA | -2809? | Indel | FRAME_SHIFT | 1 | In del |
| ARID | chr12 | 46244611 | CT | C | -902 | Indel | FRAME_SHIFT | 1 | In |

| | | | | | | | | | | |
|---|---|---|---|---|---|---|---|---|---|---|
| 2 | | | | | | | | | | del |
| ARID2 | chr12 | 46245865 | C | A | S1320* | SNV | STOP_GAINED | | 1 | C>A |
| ARID2 | chr12 | 46231491 | G | A | | Indel | SPLICE_SITE_DONOR | | 1 | Indel |
| ARID2 | chr12 | 46231104 | CT | C | -342 | Indel | FRAME_SHIFT | | 1 | Indel |
| ARID2 | chr12 | 46215255 | T | TA | -231? | Indel | FRAME_SHIFT | | 1 | Indel |
| ARID2 | chr12 | 46244803 | CTAACA | C | -966 | Indel | FRAME_SHIFT | | 1 | Indel |
| ARID1A | chr1 | 27105579 | T | TC | -1731? | Indel | FRAME_SHIFT | | 1 | Indel |
| ARID1A | chr1 | 27101300 | C | T | R1528* | SNV | STOP_GAINED | | 1 | C>T |
| ARID1A | chr1 | 27105930 | T | TG | -1848? | Indel | FRAME_SHIFT | | 2 | Indel |
| ARID1A | chr1 | 27105736 | G | T | E1783* | SNV | STOP_GAINED | | 1 | G>T |
| ARID1A | chr1 | 27106335 | CTGTG | C | -1983 | Indel | FRAME_SHIFT | | 1 | Indel |
| ARID1A | chr1 | 27106549 | G | T | E2054* | SNV | STOP_GAINED | | 1 | G>T |
| ARID1A | chr1 | 27057835 | C | T | Q515* | SNV | STOP_GAINED | | 2 | C>T |
| ARID1A | chr1 | 27057967 | CCT | C | -559 | Indel | FRAME_SHIFT | | 1 | Indel |
| ARID1A | chr1 | 27099947 | C | T | R1276* | SNV | STOP_GAINED | | 2 | C>T |
| ARID1A | chr1 | 27106803 | AC | A | -2139 | Indel | FRAME_SHIFT | | 2 | Indel |
| ARID1A | chr1 | 27106005 | AC | A | -1873 | Indel | FRAME_SHIFT | | 1 | Indel |
| ARID1A | chr1 | 27100284 | CTCCTATAGACATGAT | C | -1335 | Indel | FRAME_SHIFT | | 1 | Indel |
| ARID1A | chr1 | 27056282 | CCCGCAGCG | C | -427 | Indel | FRAME_SHIFT | | 1 | Indel |
| ARID1A | chr1 | 27056292 | T | C | Y430H | SNV | NON_SYNONYMOUS_CODING | | 1 | T>C |
| ARID1A | chr1 | 27099878 | AACGGCGGGA | A | -1253 | Indel | FRAME_SHIFT | | 1 | Indel |

| | | | | | | | | | |
|---|---|---|---|---|---|---|---|---|---|
| | | | TG GG | | | | | | |
| ARID 1A | chr1 | 27099023 | C | A | P1147T | SNV | NON_SYNONYMOUS _CODING | 1 | C> A |
| ARID 1A | chr1 | 27056211 | C | T | Q403* | SNV | STOP_GAINED | 1 | C> T |
| APO B | chr2 | 21233296 | C | CT | -2148? | Indel | FRAME_SHIFT | 1 | In del |
| APO B | chr2 | 21230961 | CT | C | -2926 | Indel | FRAME_SHIFT | 1 | In del |
| APO B | chr2 | 21238133 | C | T | | Indel | SPLICE_SITE_ACCEP TOR | 1 | In del |
| APO B | chr2 | 21246458 | A | C | I848R | SNV | NON_SYNONYMOUS _CODING | 1 | A> C |
| APC | chr5 | 112175951 | G | GA | -1554? | Indel | FRAME_SHIFT | 1 | In del |
| APC | chr5 | 112175925 | CA | C | -1545 | Indel | FRAME_SHIFT | 1 | In del |
| APC | chr5 | 112177529 | T | A | L2080 M | SNV | NON_SYNONYMOUS _CODING | 1 | T> A |
| APC | chr5 | 112154822 | A | T | K365* | SNV | STOP_GAINED | 1 | A> T |
| APC | chr5 | 112175108 | A | T | R1273* | SNV | STOP_GAINED | 1 | A> T |
| APC | chr5 | 112175879 | G | T | E1530* | SNV | STOP_GAINED | 1 | G> T |
| APC | chr5 | 112175675 | A | AAG | -1462? | Indel | FRAME_SHIFT | 1 | In del |
| APC | chr5 | 112177217 | G | A | D1976N | SNV | NON_SYNONYMOUS _CODING | 1 | G> A |
| APC | chr5 | 112164670 | G | T | | Indel | SPLICE_SITE_DONOR | 1 | In del |
| APC | chr5 | 112175897 | G | GA | -1536? | Indel | FRAME_SHIFT | 1 | In del |
| AKA P9 | chr7 | 91714999 | A | C | E3012A | SNV | NON_SYNONYMOUS _CODING | 1 | A> C |
| AKA P9 | chr7 | 91631813 | A | G | Y873C | SNV | NON_SYNONYMOUS _CODING | 1 | A> G |
| AKA P9 | chr7 | 91691747 | T | A | L1987* | SNV | STOP_GAINED | 1 | T> A |
| ADA MTS2 0 | chr12 | 43944731 | A | C | V145G | SNV | NON_SYNONYMOUS _CODING | 1 | A> C |
| ADA MTS2 0 | chr12 | 43833726 | G | C | R813G | SNV | NON_SYNONYMOUS _CODING | 1 | G> C |
| ADA MTS2 0 | chr12 | 43858516 | G | T | L463I | SNV | NON_SYNONYMOUS _CODING | 1 | G> T |
| ADA MTS2 0 | chr12 | 43828120 | G | A | S883F | SNV | NON_SYNONYMOUS _CODING | 1 | G> A |

| | | | | | | | | | |
|---|---|---|---|---|---|---|---|---|---|
| ADA MTS20 | chr12 | 43777734 | C | T | C1500Y | SNV | NON_SYNONYMOUS_CODING | 1 | C>T |
| ADA MTS20 | chr12 | 43792997 | C | A | V1442L | SNV | NON_SYNONYMOUS_CODING | 1 | C>A |
| ADA MTS20 | chr12 | 43840436 | C | T | C720Y | SNV | NON_SYNONYMOUS_CODING | 1 | C>T |
| ACV R2A | chr2 | 148683685 | TA | T | -435 | Indel | FRAME_SHIFT | 3 | Indel |
| ACV R2A | chr2 | 148657040 | G | GA | -93? | Indel | FRAME_SHIFT | 1 | Indel |
| ACV R2A | chr2 | 148683685 | T | TA | -435? | Indel | FRAME_SHIFT | 2 | Indel |
| ACV R1B | chr12 | 52379131 | C | T | R420* | SNV | STOP_GAINED | 1 | C>T |
| ACV R1B | chr12 | 52379123 | G | A | G417E | SNV | NON_SYNONYMOUS_CODING | 1 | G>A |
| ACV R1B | chr12 | 52369210 | CTGAGCTCGGAG | C | -85 | Indel | FRAME_SHIFT | 1 | Indel |
| ACV R1B | chr12 | 52377825 | A | G | H326R | SNV | NON_SYNONYMOUS_CODING | 1 | A>G |

**Table S11. Information of TP53 mutations mapping.**

| Sample ID | AA change | Type | Chr | Start Pos | End Pos | Ref | Var |
|---|---|---|---|---|---|---|---|
| **Thai HCC** | | | | | | | |
| LCS_501A | E198* | Nonsense | chr17 | 7578257 | 7578257 | C | A |
| LCS_505A | E258K | Missense | chr17 | 7577509 | 7577509 | C | T |
| LCS_528A | R249M | Missense | chr17 | 7577535 | 7577535 | C | A |
| LCS_512A | R249S | Missense | chr17 | 7577534 | 7577534 | C | A |
| LCS_508A | R249S | Missense | chr17 | 7577534 | 7577534 | C | A |
| LCS_513A | -178 | FS | chr17 | 7578397 | 7578397 | TG | T |
| LCS_541A | C275Y | Missense | chr17 | 7577114 | 7577114 | C | T |
| LCS_542A | -269 | FS | chr17 | 7577132 | 7577132 | CT | C |
| LCS_540A | V218E | Missense | chr17 | 7578196 | 7578196 | A | T |
| LCS_538A | | Splice | chr17 | 7577610 | 7577610 | T | A |
| LCS_562A | R273H | Missense | chr17 | 7577120 | 7577120 | C | T |
| LCS_524A | K139E | Missense | chr17 | 7578515 | 7578515 | T | C |
| LCS_533A | C135S | Missense | chr17 | 7578526 | 7578526 | C | G |
| LCS_568A | G244V | Missense | chr17 | 7577550 | 7577550 | C | A |
| LCS_584A | H193L | Missense | chr17 | 7578271 | 7578271 | T | A |
| LCS_530A | R249S | Missense | chr17 | 7577534 | 7577534 | C | A |
| LCS_511A | S241Y | Missense | chr17 | 7577559 | 7577559 | G | T |
| LCS_510A | R249S | Missense | chr17 | 7577534 | 7577534 | C | A |
| LCS_559A | F113V | Missense | chr17 | 7579350 | 7579350 | A | C |
| LCS_566A | Y126H | Missense | chr17 | 7578554 | 7578554 | A | G |
| LCS_536A | -53Q? | FS | chr17 | 7579528 | 7579528 | C | CCATT G |
| LCS_532A | -284? | FS | chr17 | 7577085 | 7577085 | C | CT |
| LCS_563A | K132M | Missense | chr17 | 7578535 | 7578535 | T | A |
| LCS_509A | -260 | FS | chr17 | 7577501 | 7577501 | GGA | G |
| LCS_507A | G244D | Missense | chr17 | 7577550 | 7577550 | C | T |
| LCS_519A | R249S | Missense | chr17 | 7577534 | 7577534 | C | A |
| LCS_550A | R249S | Missense | chr17 | 7577534 | 7577534 | C | A |
| **Thai iCCA** | | | | | | | |
| LCS_612A | | Splice | chr17 | 7590693 | 7590693 | A | T |
| LCS_578A | T253A | Missense | chr17 | 7577524 | 7577524 | T | C |
| LCS_618A | R306* | Nonsense | chr17 | 7577022 | 7577022 | G | A |
| LCS_573A | N239D | Missense | chr17 | 7577566 | 7577566 | T | C |
| LCS_681A | G245S | Missense | chr17 | 7577548 | 7577548 | C | T |
| LCS_595A | R175H | Missense | chr17 | 7578406 | 7578406 | C | T |
| LCS_626A | | Splice | chr17 | 7590692 | 7590692 | T | TA |
| LCS_624A | V274F | Missense | chr17 | 7577118 | 7577118 | C | A |
| LCS_577A | A276P | Missense | chr17 | 7577112 | 7577112 | C | G |
| LCS_680A | -40 | FS | chr17 | 7579554 | 7579554 | GCA TCA AAT CAT CCA | G |
| LCS_649A | L145Q | Missense | chr17 | 7578496 | 7578496 | A | T |
| LCS_623A | A138P | Missense | chr17 | 7578518 | 7578518 | C | G |

| | | | | | | | |
|---|---|---|---|---|---|---|---|
| LCS_580A | L137Q | Missense | chr17 | 7578520 | 7578520 | A | T |
| LCS_598A | G245V | Missense | chr17 | 7577547 | 7577547 | C | A |
| LCS_574A | Q167* | Nonsense | chr17 | 7578431 | 7578431 | G | A |
| LCS_685A | -227? | FS | chr17 | 7577599 | 7577599 | C | CA |
| LCS_627A | C275G | Missense | chr17 | 7577115 | 7577115 | A | C |
| LCS_669A | -294 | FS | chr17 | 7577057 | 7577057 | TC | T |
| LCS_645A | | Splice | chr17 | 7578176 | 7578176 | C | T |
| LCS_644A | R273H | Missense | chr17 | 7577120 | 7577120 | C | T |
| LCS_687A | R273H | Missense | chr17 | 7577120 | 7577120 | C | T |
| LCS_688A | V173L | Missense | chr17 | 7578413 | 7578413 | C | A |
| LCS_572A | R249S | Missense | chr17 | 7577534 | 7577534 | C | A |
| LCS_571A | -109TV? | FS | chr17 | 7579359 | 7579359 | G | GGAAA CCGT |
| LCS_630A | E326* | Nonsense | chr17 | 7576870 | 7576870 | C | A |
| LCS_640A | R282W | Missense | chr17 | 7577094 | 7577094 | G | A |
| LCS_652A | R213* | Nonsense | chr17 | 7578212 | 7578212 | G | A |
| LCS_658A | R248Q | Missense | chr17 | 7577538 | 7577538 | C | T |
| LCS_600A | R174W | Missense | chr17 | 7578410 | 7578410 | T | A |
| LCS_616A | V272M | Missense | chr17 | 7577124 | 7577124 | C | T |
| LCS_646A | R196* | Nonsense | chr17 | 7578263 | 7578263 | G | A |
| LCS_631A | R306* | Nonsense | chr17 | 7577022 | 7577022 | G | A |
| LCS_610A | -317 | FS | chr17 | 7576896 | 7576896 | TG | T |
| LCS_583A | R248Q | Missense | chr17 | 7577538 | 7577538 | C | T |
| LCS_583A | H115R | Missense | chr17 | 7579343 | 7579343 | T | C |
| LCS_670A | G105V | Missense | chr17 | 7579373 | 7579373 | C | A |
| LCS_604A | -20 | FS | chr17 | 7579854 | 7579854 | GA | G |
| LCS_620A | R337C | Missense | chr17 | 7574018 | 7574018 | G | A |
| LCS_679A | R196P | Missense | chr17 | 7578262 | 7578262 | C | G |
| LCS_642A | R175H | Missense | chr17 | 7578406 | 7578406 | C | T |
| LCS_674A | R280T | Missense | chr17 | 7577099 | 7577099 | C | G |
| **TCGA HCC** | | | | | | | |
| TCGA-UB-A7MC-01 | R248Q | Missense | chr17 | 7577538 | 7577538 | C | T |
| TCGA-XR-A8TG-01 | R248Q | Missense | chr17 | 7577538 | 7577538 | C | T |
| TCGA-WQ-AB4B-01 | R248Q | Missense | chr17 | 7577538 | 7577538 | C | T |
| TCGA-ED-A459-01 | Y220C | Missense | chr17 | 7578190 | 7578190 | T | C |
| TCGA-DD-AADD-01 | Y220C | Missense | chr17 | 7578190 | 7578190 | T | C |
| TCGA-DD-AACL-01 | C238R | Missense | chr17 | 7577569 | 7577569 | A | G |
| TCGA-ZP-A9D2-01 | R280K | Missense | chr17 | 7577099 | 7577099 | C | T |
| TCGA-BC-A10U-01 | R213Q | Missense | chr17 | 7578211 | 7578211 | C | T |
| TCGA-BC-A216-01 | R248W | Missense | chr17 | 7577539 | 7577539 | G | A |
| TCGA-DD-AAVU-01 | L194R | Missense | chr17 | 7578268 | 7578268 | A | C |

| | | | | | | | |
|---|---|---|---|---|---|---|---|
| TCGA-2Y-A9H8-01 | G245D | Missense | chr17 | 7577547 | 7577547 | C | T |
| TCGA-UB-A7MB-01 | Y205C | Missense | chr17 | 7578235 | 7578235 | T | C |
| TCGA-DD-AADL-01 | Y205C | Missense | chr17 | 7578235 | 7578235 | T | C |
| TCGA-FV-A4ZP-01 | E286K | Missense | chr17 | 7577082 | 7577082 | C | T |
| TCGA-BW-A5NO-01 | G266R | Missense | chr17 | 7577142 | 7577142 | C | T |
| TCGA-RG-A7D4-01 | C275Y | Missense | chr17 | 7577114 | 7577114 | C | T |
| TCGA-DD-A1EE-01 | H193R | Missense | chr17 | 7578271 | 7578271 | T | C |
| TCGA-G3-A25Z-01 | H193R | Missense | chr17 | 7578271 | 7578271 | T | C |
| TCGA-FV-A3I1-01 | H193R | Missense | chr17 | 7578271 | 7578271 | T | C |
| TCGA-MI-A75G-01 | H193R | Missense | chr17 | 7578271 | 7578271 | T | C |
| TCGA-UB-A7MF-01 | S215I | Missense | chr17 | 7578205 | 7578205 | C | A |
| TCGA-DD-A114-01 | R249S | Missense | chr17 | 7577534 | 7577534 | C | A |
| TCGA-G3-A25U-01 | R249S | Missense | chr17 | 7577534 | 7577534 | C | A |
| TCGA-CC-A3M9-01 | R249S | Missense | chr17 | 7577534 | 7577534 | C | A |
| TCGA-CC-A3MB-01 | R249S | Missense | chr17 | 7577534 | 7577534 | C | A |
| TCGA-CC-A5UE-01 | R249S | Missense | chr17 | 7577534 | 7577534 | C | A |
| TCGA-QA-A7B7-01 | R249S | Missense | chr17 | 7577534 | 7577534 | C | A |
| TCGA-CC-A7II-01 | R249S | Missense | chr17 | 7577534 | 7577534 | C | A |
| TCGA-ED-A7XP-01 | R249S | Missense | chr17 | 7577534 | 7577534 | C | A |
| TCGA-CC-A8HU-01 | R249S | Missense | chr17 | 7577534 | 7577534 | C | A |
| TCGA-CC-A8HV-01 | R249S | Missense | chr17 | 7577534 | 7577534 | C | A |
| TCGA-2Y-A9GS-01 | R249S | Missense | chr17 | 7577534 | 7577534 | C | A |
| TCGA-BC-A3KG-01 | N239S | Missense | chr17 | 7577565 | 7577565 | T | C |
| TCGA-2Y-A9GY-01 | R213L | Missense | chr17 | 7578211 | 7578211 | C | A |
| TCGA-FV-A4ZQ-01 | G266V | Missense | chr17 | 7577141 | 7577141 | C | A |
| TCGA-CC-A8HT-01 | M237I | Missense | chr17 | 7577570 | 7577570 | C | A |
| TCGA-BD-A3EP-01 | P151H | Missense | chr17 | 7578478 | 7578478 | G | T |
| TCGA-DD- | C176W | Missense | chr17 | 7578402 | 7578402 | G | C |

| | | | | | | | |
|---|---|---|---|---|---|---|---|
| AADW-01 | | | | | | | |
| TCGA-GJ-A3OU-01 | C275R | Missense | chr17 | 7577115 | 7577115 | A | G |
| TCGA-YA-A8S7-01 | Y205S | Missense | chr17 | 7578235 | 7578235 | T | G |
| TCGA-DD-AAEA-01 | I195S | Missense | chr17 | 7578265 | 7578265 | A | C |
| TCGA-G3-A7M6-01 | S215N | Missense | chr17 | 7578205 | 7578205 | C | T |
| TCGA-DD-AADB-01 | R273S | Missense | chr17 | 7577121 | 7577121 | G | T |
| TCGA-DD-AACB-01 | D281E | Missense | chr17 | 7577095 | 7577095 | G | C |
| TCGA-BC-A10R-01 | Q136* | Nonsense | chr17 | 7578524 | 7578524 | G | A |
| TCGA-G3-A25V-01 | I251Sfs*94 | fs del | chr17 | 7577531 | 7577531 | G | - |
| TCGA-DD-AACU-01 | M246V | Missense | chr17 | 7577545 | 7577545 | T | C |
| TCGA-DD-AAD3-01 | E271V | Missense | chr17 | 7577126 | 7577126 | T | A |
| TCGA-RC-A7SK-01 | R174W | Missense | chr17 | 7578410 | 7578410 | T | A |
| TCGA-CC-5263-01 | C277* | Nonsense | chr17 | 7577107 | 7577107 | A | T |
| TCGA-2Y-A9HA-01 | M237Cfs*10 | fs del | chr17 | 7577572 | 7577572 | T | - |
| TCGA-BC-A5W4-01 | A276G | Missense | chr17 | 7577111 | 7577111 | G | C |
| TCGA-DD-A3A7-01 | C135* | Nonsense | chr17 | 7578525 | 7578525 | G | T |
| TCGA-DD-AAD5-01 | F341V | Missense | chr17 | 7574006 | 7574006 | A | C |
| TCGA-G3-A5SJ-01 | K139N | Missense | chr17 | 7578513 | 7578513 | C | A |
| TCGA-DD-A39Y-01 | R158H | Missense | chr17 | 7578457 | 7578457 | C | T |
| TCGA-MI-A75I-01 | R158H | Missense | chr17 | 7578457 | 7578457 | C | T |
| TCGA-FV-A3R2-01 | R306* | Nonsense | chr17 | 7577022 | 7577022 | G | A |
| TCGA-4R-AA8I-01 | P191del | if del | chr17 | 7578275 | 7578277 | GAG | - |
| TCGA-T1-A6J8-01 | X187_splice | Splice | chr17 | 7578290 | 7578290 | C | T |
| TCGA-CC-A5UD-01 | V157F | Missense | chr17 | 7578461 | 7578461 | C | A |
| TCGA-CC-A7IJ-01 | V157F | Missense | chr17 | 7578461 | 7578461 | C | A |
| TCGA-DD-AAVV-01 | V157F | Missense | chr17 | 7578461 | 7578461 | C | A |
| TCGA-BW-A5NQ-01 | Q192* | Nonsense | chr17 | 7578275 | 7578275 | G | A |
| TCGA-DD- | Q192* | Nonsense | chr17 | 7578275 | 7578275 | G | A |

| | | | | | | | |
|---|---|---|---|---|---|---|---|
| AACG-01 | | | | | | | |
| TCGA-K7-AAU7-01 | R156P | Missense | chr17 | 7578463 | 7578463 | C | G |
| TCGA-WJ-A86L-01 | E204* | Nonsense | chr17 | 7578239 | 7578239 | C | A |
| TCGA-CC-A7IG-01 | E221* | Nonsense | chr17 | 7578188 | 7578188 | C | A |
| TCGA-CC-A3MA-01 | F113C | Missense | chr17 | 7579349 | 7579349 | A | C |
| TCGA-CC-A1HT-01 | X126_splice | Splice | chr17 | 7578555 | 7578555 | C | T |
| TCGA-DD-AADC-01 | X126_splice | Splice | chr17 | 7578555 | 7578555 | C | T |
| TCGA-DD-AAED-01 | X126_splice | Splice | chr17 | 7578555 | 7578555 | C | T |
| TCGA-RC-A7SH-01 | V197L | Missense | chr17 | 7578260 | 7578260 | C | A |
| TCGA-DD-AADV-01 | S106R | Missense | chr17 | 7579369 | 7579369 | G | C |
| TCGA-DD-AACS-01 | Y126N | Missense | chr17 | 7578554 | 7578554 | A | T |
| TCGA-UB-A7MD-01 | X33_splice | Splice | chr17 | 7579591 | 7579591 | C | T |
| TCGA-DD-A1EB-01 | Y126D | Missense | chr17 | 7578554 | 7578554 | A | C |
| TCGA-DD-AACE-01 | F328Sfs*17 | fs del | chr17 | 7576863 | 7576863 | A | - |
| TCGA-DD-AACV-01 | Q144* | Nonsense | chr17 | 7578500 | 7578500 | G | A |
| TCGA-DD-A1EG-01 | V157G | Missense | chr17 | 7578460 | 7578460 | A | C |
| TCGA-G3-A3CJ-01 | V143M | Missense | chr17 | 7578503 | 7578503 | C | T |
| TCGA-RC-A6M6-01 | K292* | Nonsense | chr17 | 7577064 | 7577064 | T | A |
| TCGA-2Y-A9H4-01 | S90Pfs*33 | fs del | chr17 | 7579420 | 7579420 | G | - |
| TCGA-5C-AAPD-01 | L43* | fs del | chr17 | 7579559 | 7579559 | A | - |
| TCGA-K7-A5RG-01 | X307_splice | Splice | chr17 | 7576928 | 7576928 | T | C |
| TCGA-WX-AA44-01 | X307_splice | Splice | chr17 | 7576928 | 7576928 | T | C |
| TCGA-G3-AAV4-01 | E224E | splice_region | chr17 | 7578177 | 7578177 | C | T |
| TCGA-G3-A5SM-01 | X332_splice | Splice | chr17 | 7574035 | 7574035 | T | C |
| TCGA-DD-A1EI-01 | E171* | Nonsense | chr17 | 7578419 | 7578419 | C | A |
| TCGA-DD-AAD8-01 | X225_splice | Splice | chr17 | 7577610 | 7577610 | T | A |
| TCGA-RC-A7SB-01 | E258K | Missense | chr17 | 7577509 | 7577509 | C | T |
| TCGA-DD-A1EL- | A161S | Missense | chr17 | 7578449 | 7578449 | C | A |

| | | | | | | | |
|---|---|---|---|---|---|---|---|
| 01 | | | | | | | |
| TCGA-DD-A11A-01 | V157_R158dup | if ins | chr17 | 7578454 | 7578455 | NA | CGCGGA |
| TCGA-DD-AAE6-01 | V143Afs*29 | fs ins | chr17 | 7578502 | 7578503 | NA | CAGGG |
| TCGA-NI-A8LF-01 | A276Lfs*31 | fs ins | chr17 | 7577112 | 7577113 | NA | ACAA |
| TCGA-ED-A7XO-01 | L257P | Missense | chr17 | 7577511 | 7577511 | A | G |
| TCGA-BC-A69H-01 | V97Efs*48 | fs del | chr17 | 7579389 | 7579399 | GGGAAGGGACA | - |
| TCGA-BC-A8YO-01 | T230_T231del | if del | chr17 | 7577589 | 7577594 | GTGGTA | - |
| TCGA-CC-5258-01 | N263Ifs*82 | fs del | chr17 | 7577150 | 7577150 | T | - |
| TCGA-CC-A7IK-01 | Q317Ffs*9 | fs del | chr17 | 7576866 | 7576897 | TATTCTCCATCCAGTGGTTTCTTCTTTGGCTG | - |
| TCGA-DD-AACF-01 | V143Rfs*18 | fs del | chr17 | 7578477 | 7578504 | GGGTGTGGAATCAACCCACAGCTGCACA | - |
| TCGA-DD-AACQ-01 | D228Vfs*18 | fs del | chr17 | 7577595 | 7577598 | CAGT | - |
| TCGA-DD-AACZ-01 | N263Kfs*9 | fs ins | chr17 | 7577149 | 7577150 | - | C |
| TCGA-DD-AADN-01 | E204Vfs*4 | fs del | chr17 | 7578237 | 7578238 | CT | - |
| TCGA-DD-AAE0-01 | T253A | Missense | chr17 | 7577524 | 7577524 | T | C |
| TCGA-DD-AAEI-01 | E56Kfs*67 | fs del | chr17 | 7579521 | 7579521 | C | - |
| TCGA-ED-A7PZ-01 | R248Hfs*13 | fs del | chr17 | 7577531 | 7577538 | GGGCCTCC | - |
| TCGA-ES-A2HT-01 | S227P | Missense | chr17 | 7577602 | 7577602 | A | G |
| TCGA-FV-A496-01 | Y103Tfs*20 | fs del | chr17 | 7579380 | 7579380 | A | - |

| | | | | | | | |
|---|---|---|---|---|---|---|---|
| TCGA-G3-A7M9-01 | L194Efs *51 | fs del | chr17 | 7578264 | 7578270 | GAT AAG A | - |
| TCGA-G3-AAV7-01 | G262Vfs *83 | fs del | chr17 | 7577153 | 7577153 | C | - |

**Table S12. Comparison table of TP53 mutation between HFGC and LFGC among PLC cohorts**

| Cohort | Class | TP53 MUT | TP53 *DBD MUT | TP53 WT | Fisher's exact test (p) | |
|---|---|---|---|---|---|---|
| | | | | | MUT vs. WT | DBD MUT vs. WT |
| TIGER_LC (n=152) | HFGC | 55 | 38 | 46 | 2.85E-04 | 5.21E-04 |
| | LFGC | 12 | 7 | 39 | | |
| Thai_HCC (n=62) | HFGC | 23 | 19 | 25 | 0.24 | 4.20E-02 |
| | LFGC | 4 | 1 | 10 | | |
| Thai_iCCA (n=90) | HFGC | 32 | 19 | 21 | 4.78E-04 | 7.00E-03 |
| | LFGC | 8 | 6 | 29 | | |
| TCGA_HCC (n=364) | HFGC | 104 | 59 | 192 | 3.14E-05 | 3.16E-03 |
| | LFGC | 7 | 5 | 61 | | |

* DBD MUT : mutation occurred within DNA binding domain (DBD)

**Table S13a. Table for tFA of TCGA SKCM cohort**

| Sample ID | tFA | Group | subset |
|---|---|---|---|
| TCGA-ER-A1A1-06A-11R-A18U-07 | 1833.533 | Low | subset1 |
| TCGA-RP-A690-06A-11R-A311-07 | 1975.275 | NA | subset1 |
| TCGA-EE-A2GU-06A-11R-A18T-07 | 1861.785 | Low | subset1 |
| TCGA-EE-A3JE-06A-11R-A20F-07 | 1861.778 | Low | subset1 |
| TCGA-BF-AAOX-01A-11R-A39D-07 | 2439.111 | High | subset1 |
| TCGA-ER-A19P-06A-11R-A18S-07 | 2019.652 | NA | subset1 |
| TCGA-D3-A2J6-06A-11R-A18T-07 | 2308.362 | NA | subset1 |
| TCGA-D9-A6EG-06A-12R-A32P-07 | 1846.107 | Low | subset1 |
| TCGA-WE-A8ZR-06A-11R-A37K-07 | 1766.808 | Low | subset1 |
| TCGA-EB-A82C-01A-11R-A352-07 | 2376.790 | High | subset1 |
| TCGA-FS-A1YY-06A-11R-A18T-07 | 2048.012 | NA | subset1 |
| TCGA-ER-A19T-06A-11R-A18U-07 | 2747.214 | High | subset1 |
| TCGA-D3-A51T-06A-11R-A266-07 | 2083.324 | NA | subset1 |
| TCGA-EE-A3AF-06A-11R-A18S-07 | 2319.922 | NA | subset1 |
| TCGA-QB-AA9O-06A-11R-A39D-07 | 2551.261 | High | subset2 |
| TCGA-EE-A29R-06A-11R-A18T-07 | 2071.334 | NA | subset1 |
| TCGA-D3-A2J9-06A-11R-A18T-07 | 1963.680 | NA | subset1 |
| TCGA-FS-A4FC-06A-11R-A24X-07 | 1776.473 | Low | subset1 |
| TCGA-D3-A3C8-06A-12R-A18S-07 | 2114.548 | NA | subset1 |
| TCGA-EE-A2GJ-06A-11R-A18U-07 | 1769.805 | Low | subset1 |
| TCGA-EB-A5UL-06A-11R-A311-07 | 2132.368 | NA | subset1 |
| TCGA-D9-A1X3-06A-11R-A18S-07 | 1800.016 | Low | subset1 |
| TCGA-Z2-AA3V-06A-11R-A39D-07 | 2945.241 | High | subset1 |
| TCGA-BF-A5EP-01A-12R-A27Q-07 | 2282.277 | NA | subset1 |
| TCGA-FS-A1Z4-06A-11R-A18T-07 | 1702.189 | Low | subset1 |
| TCGA-EB-A5SG-06A-11R-A311-07 | 2307.041 | NA | subset1 |
| TCGA-D3-A3ML-06A-11R-A21D-07 | 1974.338 | NA | subset1 |
| TCGA-EE-A2M6-06A-12R-A18S-07 | 2023.653 | NA | subset1 |
| TCGA-D3-A2JE-06A-11R-A37K-07 | 2037.189 | NA | subset1 |
| TCGA-D3-A8GD-06A-11R-A37K-07 | 2183.112 | NA | subset1 |
| TCGA-ER-A42K-06A-11R-A24X-07 | 2062.055 | NA | subset1 |
| TCGA-EE-A2MQ-06A-11R-A18S-07 | 2026.100 | NA | subset1 |
| TCGA-DA-A3F8-06A-11R-A20F-07 | 1998.634 | NA | subset1 |
| TCGA-GN-A265-06A-21R-A18T-07 | 2176.448 | NA | subset1 |
| TCGA-EE-A3AA-06A-11R-A18S-07 | 1735.311 | Low | subset1 |
| TCGA-EE-A3J8-06A-11R-A20F-07 | 1711.218 | Low | subset1 |
| TCGA-FR-A7U9-06A-11R-A352-07 | 2339.388 | High | subset1 |
| TCGA-EB-A5SE-01A-11R-A311-07 | 2178.593 | NA | subset1 |
| TCGA-DA-A1IC-06A-11R-A18S-07 | 2023.989 | NA | subset1 |

| | | | |
|---|---|---|---|
| TCGA-ER-A3EV-06A-11R-A20F-07 | 2619.586 | High | subset1 |
| TCGA-EB-A551-01A-21R-A27Q-07 | 1833.502 | Low | subset1 |
| TCGA-EE-A2ME-06A-11R-A18T-07 | 1648.314 | Low | subset1 |
| TCGA-EE-A29S-06A-11R-A18T-07 | 2153.671 | NA | subset1 |
| TCGA-D3-A1QA-06A-11R-A18T-07 | 1841.090 | Low | subset1 |
| TCGA-EE-A2A0-06A-11R-A18T-07 | 1616.215 | Low | subset1 |
| TCGA-FW-A5DY-06A-11R-A311-07 | 1924.269 | NA | subset1 |
| TCGA-D3-A5GS-06A-11R-A27Q-07 | 2074.785 | NA | subset1 |
| TCGA-BF-A3DN-01A-11R-A20F-07 | 1723.537 | Low | subset1 |
| TCGA-FS-A1Z7-06A-11R-A18T-07 | 1973.111 | NA | subset1 |
| TCGA-EE-A29L-06A-12R-A18S-07 | 1920.932 | NA | subset1 |
| TCGA-WE-A8ZM-06A-11R-A37K-07 | 2074.598 | NA | subset1 |
| TCGA-BF-A5EO-01A-12R-A27Q-07 | 3161.755 | High | subset1 |
| TCGA-FS-A1ZU-06A-12R-A18T-07 | 2372.705 | High | subset1 |
| TCGA-WE-AA9Y-06A-12R-A38C-07 | 2221.827 | NA | subset1 |
| TCGA-D3-A3CF-06A-11R-A18T-07 | 1960.643 | NA | subset1 |
| TCGA-D3-A1Q1-06A-21R-A18T-07 | 2314.370 | NA | subset1 |
| TCGA-D3-A3C3-06A-12R-A18S-07 | 2487.458 | High | subset1 |
| TCGA-D9-A3Z4-01A-11R-A239-07 | 2181.147 | NA | subset1 |
| TCGA-ER-A2NF-01A-11R-A18T-07 | 2351.492 | High | subset1 |
| TCGA-FS-A1ZT-06A-11R-A18U-07 | 1802.765 | Low | subset1 |
| TCGA-D3-A8GB-06A-11R-A37K-07 | 2413.124 | High | subset1 |
| TCGA-D3-A8GP-06A-11R-A37K-07 | 2128.930 | NA | subset1 |
| TCGA-WE-A8ZY-06A-11R-A37K-07 | 2165.085 | NA | subset1 |
| TCGA-FS-A1YX-06A-11R-A18T-07 | 1874.948 | Low | subset1 |
| TCGA-EB-A3XB-01A-11R-A239-07 | 2013.839 | NA | subset1 |
| TCGA-ER-A19S-06A-11R-A18U-07 | 2074.735 | NA | subset1 |
| TCGA-EE-A2MR-06A-11R-A18S-07 | 1864.706 | Low | subset1 |
| TCGA-D3-A8GE-06A-11R-A37K-07 | 1713.446 | Low | subset1 |
| TCGA-FS-A1YW-06A-11R-A18T-07 | 1753.762 | Low | subset1 |
| TCGA-GN-A4U5-01A-11R-A32P-07 | 2380.036 | High | subset1 |
| TCGA-HR-A2OH-06A-11R-A18U-07 | 2112.931 | NA | subset1 |
| TCGA-D3-A8GS-06A-12R-A37K-07 | 2234.734 | NA | subset1 |
| TCGA-D3-A8GN-06A-11R-A37K-07 | 2336.416 | High | subset1 |
| TCGA-EE-A2MP-06A-11R-A18S-07 | 2230.703 | NA | subset1 |
| TCGA-EB-A5VU-01A-21R-A32P-07 | 1984.819 | NA | subset1 |
| TCGA-EE-A2GN-06A-11R-A18S-07 | 2178.228 | NA | subset1 |
| TCGA-GN-A8LN-01A-11R-A37K-07 | 1894.746 | Low | subset2 |
| TCGA-FR-A8YE-06A-11R-A37K-07 | 1907.599 | Low | subset1 |
| TCGA-EE-A185-06A-11R-A18S-07 | 2238.907 | NA | subset1 |
| TCGA-ER-A195-06A-11R-A18U-07 | 1850.852 | Low | subset1 |

| | | | |
|---|---|---|---|
| TCGA-GF-A769-01A-32R-A32P-07 | 2011.733 | NA | subset1 |
| TCGA-D3-A2JH-06A-11R-A18T-07 | 1969.564 | NA | subset1 |
| TCGA-EE-A29V-06A-12R-A18S-07 | 2721.253 | High | subset1 |
| TCGA-EB-A5SH-06A-11R-A311-07 | 3147.432 | High | subset1 |
| TCGA-D3-A1QA-07A-11R-A37K-07 | 1887.884 | Low | subset1 |
| TCGA-EB-A3HV-01A-11R-A21D-07 | 2765.014 | High | subset1 |
| TCGA-GF-A2C7-01A-11R-A18T-07 | 2015.868 | NA | subset1 |
| TCGA-DA-A1I4-06A-11R-A18U-07 | 1947.043 | NA | subset1 |
| TCGA-D3-A2JF-06A-11R-A18S-07 | 2355.263 | High | subset1 |
| TCGA-FR-A726-01A-11R-A32P-07 | 1931.381 | NA | subset1 |
| TCGA-D3-A3BZ-06A-12R-A18S-07 | 2145.162 | NA | subset1 |
| TCGA-DA-A1IB-06A-11R-A18S-07 | 1985.401 | NA | subset1 |
| TCGA-GN-A266-06A-11R-A18T-07 | 1911.179 | Low | subset1 |
| TCGA-EE-A2M8-06A-12R-A18S-07 | 1986.732 | NA | subset1 |
| TCGA-EE-A2A6-06A-11R-A18T-07 | 1945.138 | NA | subset1 |
| TCGA-ER-A2NF-06A-11R-A18T-07 | 2286.245 | NA | subset1 |
| TCGA-ER-A3ET-06A-11R-A20F-07 | 1875.620 | Low | subset1 |
| TCGA-FR-A3YN-06A-11R-A239-07 | 1987.026 | NA | subset2 |
| TCGA-EE-A3AH-06A-11R-A18S-07 | 2073.417 | NA | subset1 |
| TCGA-FR-A8YC-06A-11R-A37K-07 | 2146.297 | NA | subset1 |
| TCGA-FS-A1Z3-06A-11R-A18T-07 | 1643.659 | Low | subset1 |
| TCGA-D3-A2JP-06A-11R-A18S-07 | 2377.688 | High | subset1 |
| TCGA-RP-A695-06A-11R-A311-07 | 2045.651 | NA | subset1 |
| TCGA-WE-A8ZX-06A-11R-A37K-07 | 2318.953 | NA | subset1 |
| TCGA-DA-A95Y-06A-11R-A37K-07 | 2549.316 | High | subset1 |
| TCGA-ER-A19W-06A-41R-A239-07 | 1931.592 | NA | subset1 |
| TCGA-FS-A1ZQ-06A-11R-A18U-07 | 2542.477 | High | subset1 |
| TCGA-D3-A51G-06A-11R-A266-07 | 1840.934 | Low | subset1 |
| TCGA-GF-A4EO-06A-12R-A24X-07 | 1970.318 | NA | subset1 |
| TCGA-D9-A1JW-06A-11R-A18S-07 | 2448.345 | High | subset1 |
| TCGA-EE-A20C-06A-11R-A18S-07 | 2139.490 | NA | subset1 |
| TCGA-EE-A29X-06A-11R-A18T-07 | 2541.010 | High | subset1 |
| TCGA-EE-A2MM-06A-11R-A18S-07 | 2028.837 | NA | subset1 |
| TCGA-EE-A3JB-06A-11R-A21D-07 | 1986.424 | NA | subset1 |
| TCGA-EE-A2MK-06A-11R-A18S-07 | 2095.734 | NA | subset1 |
| TCGA-DA-A1HW-06A-11R-A18U-07 | 1973.726 | NA | subset1 |
| TCGA-FS-A1ZD-06A-11R-A18T-07 | 1624.258 | Low | subset1 |
| TCGA-FS-A1ZP-06A-11R-A18T-07 | 1777.354 | Low | subset1 |
| TCGA-D3-A2JL-06A-11R-A18S-07 | 2534.391 | High | subset1 |
| TCGA-BF-AAP2-01A-11R-A40A-07 | 2303.308 | NA | subset1 |
| TCGA-D3-A2JB-06A-11R-A18T-07 | 2070.220 | NA | subset1 |

| | | | |
|---|---|---|---|
| TCGA-D3-A3CE-06A-11R-A18S-07 | 1758.624 | Low | subset1 |
| TCGA-GF-A6C9-06A-11R-A311-07 | 2416.296 | High | subset1 |
| TCGA-ER-A19K-01A-21R-A18T-07 | 2027.554 | NA | subset1 |
| TCGA-W3-AA1V-06B-11R-A40A-07 | 1847.777 | Low | subset1 |
| TCGA-EB-A97M-01A-11R-A38C-07 | 2843.162 | High | subset1 |
| TCGA-EE-A29N-06A-12R-A18S-07 | 1758.215 | Low | subset1 |
| TCGA-EB-A4OZ-01A-12R-A266-07 | 3544.320 | High | subset1 |
| TCGA-W3-AA21-06A-11R-A38C-07 | 2925.908 | High | subset1 |
| TCGA-ER-A19F-06A-11R-A18S-07 | 2015.959 | NA | subset1 |
| TCGA-EE-A3AB-06A-11R-A18S-07 | 2371.241 | High | subset1 |
| TCGA-EB-A431-01A-11R-A266-07 | 1962.443 | NA | subset1 |
| TCGA-XV-A9W2-01A-11R-A39D-07 | 2268.226 | NA | subset1 |
| TCGA-BF-AAOU-01A-12R-A39D-07 | 2333.175 | High | subset1 |
| TCGA-EE-A2M5-06A-12R-A18S-07 | 1914.413 | NA | subset1 |
| TCGA-YG-AA3O-06A-11R-A38C-07 | 1926.656 | NA | subset1 |
| TCGA-D3-A8GJ-06A-11R-A37K-07 | 2290.560 | NA | subset1 |
| TCGA-EB-A550-01A-61R-A27Q-07 | 2053.782 | NA | subset1 |
| TCGA-FR-A729-06A-11R-A352-07 | 2166.960 | NA | subset1 |
| TCGA-ER-A42H-01A-11R-A24X-07 | 2329.346 | High | subset1 |
| TCGA-DA-A1IA-06A-11R-A18S-07 | 3272.298 | High | subset1 |
| TCGA-EB-A42Y-01A-12R-A24X-07 | 2126.079 | NA | subset1 |
| TCGA-LH-A9QB-06A-11R-A38C-07 | 1597.084 | Low | subset1 |
| TCGA-BF-A1Q0-01A-21R-A18S-07 | 2191.620 | NA | subset1 |
| TCGA-FS-A1ZB-06A-12R-A18S-07 | 2298.871 | NA | subset1 |
| TCGA-D3-A51R-06A-11R-A266-07 | 1856.005 | Low | subset1 |
| TCGA-FS-A1ZC-06A-11R-A18T-07 | 1928.130 | NA | subset1 |
| TCGA-EB-A41B-01A-11R-A24X-07 | 2602.208 | High | subset1 |
| TCGA-DA-A1HV-06A-21R-A18S-07 | 2240.725 | NA | subset1 |
| TCGA-FR-A44A-06A-11R-A24X-07 | 2060.373 | NA | subset1 |
| TCGA-EE-A2GL-06A-11R-A18S-07 | 2030.395 | NA | subset1 |
| TCGA-EE-A3J7-06A-11R-A20F-07 | 1923.413 | NA | subset1 |
| TCGA-EB-A553-01A-12R-A27Q-07 | 2742.727 | High | subset1 |
| TCGA-D3-A1Q4-06A-11R-A18T-07 | 2087.720 | NA | subset1 |
| TCGA-EE-A2MC-06A-12R-A18S-07 | 1912.422 | Low | subset1 |
| TCGA-EE-A2MN-06A-11R-A18S-07 | 2253.975 | NA | subset1 |
| TCGA-BF-A1PV-01A-11R-A18U-07 | 1816.889 | Low | subset1 |
| TCGA-3N-A9WD-06A-11R-A38C-07 | 2125.827 | NA | subset1 |
| TCGA-WE-A8ZN-06A-11R-A37K-07 | 3033.920 | High | subset1 |
| TCGA-BF-AAP7-01A-11R-A40A-07 | 2843.033 | High | subset1 |
| TCGA-EE-A2A1-06A-11R-A18T-07 | 2106.479 | NA | subset1 |
| TCGA-EE-A3JA-06A-11R-A20F-07 | 1734.669 | Low | subset1 |

| | | | |
|---|---|---|---|
| TCGA-EE-A29C-06A-21R-A18S-07 | 2411.624 | High | subset2 |
| TCGA-D3-A3MO-06A-11R-A21D-07 | 1982.460 | NA | subset1 |
| TCGA-ER-A19G-06A-11R-A18U-07 | 1930.281 | NA | subset1 |
| TCGA-FS-A1ZA-06A-11R-A18T-07 | 2030.301 | NA | subset1 |
| TCGA-EE-A181-06A-11R-A18S-07 | 1996.543 | NA | subset1 |
| TCGA-EE-A2GB-06A-11R-A18T-07 | 1719.227 | Low | subset1 |
| TCGA-EE-A2GI-06A-11R-A18T-07 | 2339.467 | High | subset1 |
| TCGA-D9-A6E9-06A-12R-A311-07 | 2542.533 | High | subset1 |
| TCGA-D3-A2JN-06A-11R-A18S-07 | 2213.550 | NA | subset1 |
| TCGA-DA-A1I2-06A-21R-A18U-07 | 1859.553 | Low | subset1 |
| TCGA-FW-A3R5-06A-11R-A239-07 | 1734.250 | Low | subset1 |
| TCGA-BF-A5EQ-01A-21R-A27Q-07 | 2156.379 | NA | subset1 |
| TCGA-EB-A3Y7-01A-11R-A239-07 | 2995.436 | High | subset1 |
| TCGA-WE-AAA3-06A-11R-A38C-07 | 2409.066 | High | subset1 |
| TCGA-BF-A3DM-01A-11R-A20F-07 | 1867.547 | Low | subset1 |
| TCGA-GF-A6C8-06A-12R-A311-07 | 2117.417 | NA | subset1 |
| TCGA-EB-A3Y6-01A-21R-A239-07 | 2084.139 | NA | subset1 |
| TCGA-FS-A1ZS-06A-12R-A18T-07 | 1727.620 | Low | subset1 |
| TCGA-EB-A5VV-06A-11R-A32P-07 | 2086.798 | NA | subset1 |
| TCGA-FS-A1ZM-06A-12R-A18S-07 | 2005.679 | NA | subset1 |
| TCGA-D3-A5GU-06A-11R-A27Q-07 | 2030.325 | NA | subset1 |
| TCGA-GN-A267-06A-21R-A18T-07 | 2433.771 | High | subset1 |
| TCGA-XV-A9VZ-01A-11R-A38C-07 | 2080.786 | NA | subset1 |
| TCGA-FW-A5DX-01A-11R-A27Q-07 | 1873.973 | Low | subset1 |
| TCGA-YD-A9TA-06A-11R-A39D-07 | 1855.413 | Low | subset1 |
| TCGA-GN-A262-06A-11R-A18T-07 | 2511.234 | High | subset1 |
| TCGA-WE-A8K6-06A-11R-A37K-07 | 2211.806 | NA | subset1 |
| TCGA-D3-A3CC-06A-11R-A18S-07 | 1986.728 | NA | subset1 |
| TCGA-FS-A4F5-06A-11R-A266-07 | 1949.295 | NA | subset1 |
| TCGA-FS-A1ZK-06A-11R-A18T-07 | 1966.155 | NA | subset1 |
| TCGA-EB-A85J-01A-12R-A352-07 | 2465.963 | High | subset1 |
| TCGA-EB-A6QZ-01A-12R-A32P-07 | 2285.247 | NA | subset1 |
| TCGA-DA-A3F2-06A-11R-A20F-07 | 1916.093 | NA | subset1 |
| TCGA-D9-A149-06A-11R-A18S-07 | 2179.247 | NA | subset1 |
| TCGA-EE-A180-06A-11R-A21D-07 | 1962.510 | NA | subset1 |
| TCGA-D9-A3Z3-06A-11R-A239-07 | 2329.837 | High | subset1 |
| TCGA-GN-A9SD-06A-11R-A40A-07 | 2121.817 | NA | subset1 |
| TCGA-EE-A2MI-06A-11R-A18U-07 | 1770.823 | Low | subset1 |
| TCGA-DA-A1I7-06A-22R-A18S-07 | 2155.671 | NA | subset1 |
| TCGA-W3-AA1Q-06A-11R-A38C-07 | 2080.683 | NA | subset1 |
| TCGA-BF-A9VF-01A-11R-A37K-07 | 2950.453 | High | subset1 |

| | | | |
|---|---|---|---|
| TCGA-W3-AA1R-06A-11R-A39D-07 | 2231.668 | NA | subset1 |
| TCGA-GN-A4U3-06A-11R-A32P-07 | 2117.681 | NA | subset1 |
| TCGA-RP-A693-06A-13R-A311-07 | 1956.734 | NA | subset1 |
| TCGA-D3-A5GO-06A-12R-A27Q-07 | 2364.763 | High | subset1 |
| TCGA-XV-AAZV-01A-11R-A40A-07 | 2662.743 | High | subset1 |
| TCGA-ER-A193-06A-12R-A18S-07 | 1985.362 | NA | subset1 |
| TCGA-D3-A5GL-06A-11R-A27Q-07 | 2007.073 | NA | subset1 |
| TCGA-D3-A3MU-06A-11R-A21D-07 | 2541.853 | High | subset1 |
| TCGA-EE-A29P-06A-11R-A18T-07 | 2028.607 | NA | subset1 |
| TCGA-BF-AAP6-01A-11R-A40A-07 | 2012.012 | NA | subset1 |
| TCGA-EB-A3XD-01A-22R-A239-07 | 2593.070 | High | subset1 |
| TCGA-XV-AAZY-01A-12R-A40A-07 | 3290.632 | High | subset1 |
| TCGA-EE-A2GE-06A-11R-A18T-07 | 2048.518 | NA | subset1 |
| TCGA-EB-A5KH-06A-11R-A27Q-07 | 2802.093 | High | subset1 |
| TCGA-D3-A8GQ-06A-11R-A37K-07 | 2308.990 | NA | subset1 |
| TCGA-D3-A8GL-06A-11R-A37K-07 | 1959.640 | NA | subset1 |
| TCGA-GN-A268-06A-11R-A18T-07 | 1933.873 | NA | subset1 |
| TCGA-FS-A1ZR-06A-21R-A18U-07 | 1835.580 | Low | subset1 |
| TCGA-D3-A1Q6-06A-11R-A18T-07 | 1544.720 | Low | subset1 |
| TCGA-EE-A3AC-06A-11R-A18S-07 | 1901.392 | Low | subset1 |
| TCGA-EE-A29H-06A-12R-A18S-07 | 2223.850 | NA | subset1 |
| TCGA-D3-A1Q3-06A-11R-A18T-07 | 1764.291 | Low | subset1 |
| TCGA-GN-A26C-01A-11R-A18T-07 | 2226.242 | NA | subset1 |
| TCGA-EE-A2GM-06B-11R-A18S-07 | 1931.751 | NA | subset1 |
| TCGA-D3-A1Q8-06A-11R-A18T-07 | 2000.435 | NA | subset1 |
| TCGA-BF-AAP0-06A-11R-A39D-07 | 2329.012 | NA | subset1 |
| TCGA-EB-A3XE-01A-12R-A239-07 | 2935.296 | High | subset1 |
| TCGA-ER-A2NB-01A-12R-A18S-07 | 2398.038 | High | subset1 |
| TCGA-HR-A2OG-06A-21R-A18U-07 | 1885.062 | Low | subset1 |
| TCGA-GN-A26A-06A-11R-A18T-07 | 2213.402 | NA | subset1 |
| TCGA-D9-A4Z2-01A-11R-A24X-07 | 1956.883 | NA | subset1 |
| TCGA-ER-A2NG-06A-11R-A18T-07 | 2086.277 | NA | subset1 |
| TCGA-XV-AAZW-01A-12R-A40A-07 | 2405.637 | High | subset1 |
| TCGA-D3-A51N-06A-11R-A266-07 | 2375.036 | High | subset1 |
| TCGA-ER-A42L-06A-11R-A24X-07 | 1884.028 | Low | subset1 |
| TCGA-W3-A828-06A-11R-A352-07 | 1728.455 | Low | subset1 |
| TCGA-D3-A51H-06A-12R-A266-07 | 2163.153 | NA | subset1 |
| TCGA-D3-A8GK-06A-11R-A37K-07 | 2323.482 | NA | subset1 |
| TCGA-YD-A89C-06A-11R-A37K-07 | 2257.450 | NA | subset1 |
| TCGA-EB-A3XF-01A-31R-A239-07 | 2003.544 | NA | subset1 |
| TCGA-WE-A8K4-01A-12R-A37K-07 | 2094.691 | NA | subset1 |

| | | | |
|---|---|---|---|
| TCGA-EE-A29E-06A-11R-A18T-07 | 1956.071 | NA | subset1 |
| TCGA-FS-A1ZJ-06A-12R-A18S-07 | 2172.719 | NA | subset1 |
| TCGA-ER-A19A-06A-21R-A18U-07 | 1892.173 | Low | subset1 |
| TCGA-DA-A1I0-06A-11R-A20F-07 | 3273.873 | High | subset1 |
| TCGA-EE-A182-06A-11R-A18T-07 | 3470.002 | High | subset1 |
| TCGA-DA-A95V-06A-11R-A37K-07 | 3114.335 | High | subset1 |
| TCGA-EE-A2MD-06A-11R-A18T-07 | 1836.676 | Low | subset1 |
| TCGA-FS-A1ZZ-06A-11R-A18S-07 | 1858.423 | Low | subset1 |
| TCGA-FS-A4F4-06A-12R-A266-07 | 2834.993 | High | subset1 |
| TCGA-GN-A263-01A-11R-A18T-07 | 1761.940 | Low | subset1 |
| TCGA-RP-A694-06A-11R-A311-07 | 1899.378 | Low | subset1 |
| TCGA-ER-A194-01A-11R-A18U-07 | 3309.356 | High | subset1 |
| TCGA-EE-A29B-06A-11R-A18U-07 | 2639.422 | High | subset1 |
| TCGA-GN-A4U7-06A-21R-A32P-07 | 2605.683 | High | subset1 |
| TCGA-EE-A3J3-06A-11R-A20F-07 | 2069.494 | NA | subset1 |
| TCGA-D3-A3CB-06A-11R-A18S-07 | 1964.008 | NA | subset1 |
| TCGA-FR-A2OS-01A-11R-A21D-07 | 1640.405 | Low | subset1 |
| TCGA-GN-A264-06A-11R-A18U-07 | 1909.148 | Low | subset1 |
| TCGA-EB-A85I-01A-11R-A352-07 | 2645.651 | High | subset1 |
| TCGA-D9-A148-06A-11R-A18S-07 | 2325.850 | NA | subset1 |
| TCGA-ER-A19T-01A-11R-A18T-07 | 2445.725 | High | subset1 |
| TCGA-D3-A51F-06A-11R-A266-07 | 2176.655 | NA | subset1 |
| TCGA-D9-A4Z6-06A-12R-A266-07 | 2008.184 | NA | subset1 |
| TCGA-D3-A8GI-06A-11R-A37K-07 | 3280.004 | High | subset1 |
| TCGA-WE-A8ZQ-06A-41R-A37K-07 | 2016.299 | NA | subset1 |
| TCGA-EE-A29A-06A-12R-A18U-07 | 1970.783 | NA | subset1 |
| TCGA-GN-A4U4-06A-11R-A32P-07 | 1906.754 | Low | subset2 |
| TCGA-EB-A41A-01A-11R-A24X-07 | 2100.956 | NA | subset1 |
| TCGA-FW-A3TV-06A-11R-A239-07 | 1771.361 | Low | subset1 |
| TCGA-BF-A5ES-01A-11R-A27Q-07 | 2474.821 | High | subset1 |
| TCGA-EB-A24D-01A-11R-A18T-07 | 2042.970 | NA | subset1 |
| TCGA-D3-A1Q7-06A-11R-A18T-07 | 1796.908 | Low | subset1 |
| TCGA-EE-A184-06A-11R-A18S-07 | 2126.913 | NA | subset1 |
| TCGA-D3-A1Q5-06A-11R-A18T-07 | 1751.546 | Low | subset1 |
| TCGA-D9-A4Z3-01A-11R-A266-07 | 2142.430 | NA | subset1 |
| TCGA-FS-A4F2-06A-11R-A24X-07 | 2296.874 | NA | subset1 |
| TCGA-ER-A2NH-06A-11R-A18S-07 | 2074.206 | NA | subset1 |
| TCGA-D3-A8GR-06A-11R-A37K-07 | 2804.760 | High | subset1 |
| TCGA-W3-AA1O-06A-11R-A38C-07 | 1843.408 | Low | subset1 |
| TCGA-WE-A8ZO-06A-11R-A37K-07 | 1859.589 | Low | subset1 |
| TCGA-EE-A2GR-06A-11R-A18S-07 | 1757.063 | Low | subset1 |

| | | | |
|---|---|---|---|
| TCGA-ER-A19O-06A-11R-A18S-07 | 2365.430 | High | subset1 |
| TCGA-D3-A3C7-06A-11R-A18U-07 | 2233.500 | NA | subset1 |
| TCGA-EB-A82B-01A-11R-A352-07 | 2657.561 | High | subset1 |
| TCGA-EB-A6L9-06A-11R-A32P-07 | 2948.606 | High | subset1 |
| TCGA-DA-A1I8-06A-11R-A18T-07 | 2050.480 | NA | subset1 |
| TCGA-ER-A2NC-06A-11R-A18T-07 | 2002.037 | NA | subset1 |
| TCGA-EB-A3XC-01A-11R-A239-07 | 2406.407 | High | subset1 |
| TCGA-EE-A29G-06A-12R-A18T-07 | 1986.555 | NA | subset1 |
| TCGA-BF-A3DL-01A-11R-A20F-07 | 2798.171 | High | subset1 |
| TCGA-EE-A2GD-06A-11R-A18T-07 | 1819.240 | Low | subset1 |
| TCGA-EE-A2MU-06A-21R-A18S-07 | 2107.591 | NA | subset1 |
| TCGA-EE-A2GT-06A-12R-A18S-07 | 2370.242 | High | subset1 |
| TCGA-ER-A19J-06A-11R-A18S-07 | 2056.954 | NA | subset1 |
| TCGA-FS-A4F8-06A-11R-A266-07 | 1910.427 | Low | subset1 |
| TCGA-EB-A44R-06A-41R-A266-07 | 2303.148 | NA | subset1 |
| TCGA-EB-A4IQ-01A-12R-A266-07 | 2373.122 | High | subset1 |
| TCGA-Z2-A8RT-06A-11R-A37K-07 | 2126.651 | NA | subset1 |
| TCGA-GN-A4U9-06A-11R-A32P-07 | 2266.429 | NA | subset2 |
| TCGA-GN-A8LK-06A-11R-A37K-07 | 2929.237 | High | subset2 |
| TCGA-D3-A5GT-01A-12R-A311-07 | 2591.782 | High | subset1 |
| TCGA-D3-A2JG-06A-11R-A18T-07 | 2215.628 | NA | subset1 |
| TCGA-YG-AA3N-01A-11R-A38C-07 | 2798.735 | High | subset1 |
| TCGA-EE-A2A2-06A-11R-A18T-07 | 1915.004 | NA | subset1 |
| TCGA-EE-A29Q-06A-11R-A18T-07 | 1741.164 | Low | subset1 |
| TCGA-EB-A51B-01A-11R-A27Q-07 | 2039.143 | NA | subset1 |
| TCGA-Z2-AA3S-06A-11R-A39D-07 | 1943.714 | NA | subset1 |
| TCGA-D3-A1Q9-06A-11R-A18T-07 | 1840.123 | Low | subset1 |
| TCGA-FS-A1ZH-06A-11R-A18T-07 | 1976.220 | NA | subset1 |
| TCGA-EB-A5UN-06A-11R-A311-07 | 1735.959 | Low | subset1 |
| TCGA-D3-A51E-06A-11R-A266-07 | 2178.013 | NA | subset1 |
| TCGA-D3-A8GM-06A-11R-A37K-07 | 1750.134 | Low | subset1 |
| TCGA-D3-A8GO-06A-11R-A37K-07 | 2687.796 | High | subset1 |
| TCGA-WE-A8JZ-06A-11R-A37K-07 | 2113.723 | NA | subset1 |
| TCGA-EE-A183-06A-11R-A18S-07 | 2910.901 | High | subset1 |
| TCGA-EE-A17Z-06A-11R-A18S-07 | 2333.957 | High | subset1 |
| TCGA-EE-A20F-06A-21R-A18S-07 | 2113.717 | NA | subset1 |
| TCGA-EB-A44P-01A-11R-A266-07 | 1889.396 | Low | subset1 |
| TCGA-DA-A95W-06A-11R-A37K-07 | 2365.247 | High | subset1 |
| TCGA-EB-A42Z-01A-12R-A24X-07 | 2199.235 | NA | subset1 |
| TCGA-DA-A95Z-06A-11R-A37K-07 | 2345.368 | High | subset1 |
| TCGA-WE-A8K5-06A-11R-A37K-07 | 2233.435 | NA | subset1 |

| | | | |
|---|---|---|---|
| TCGA-ER-A197-06A-32R-A18S-07 | 2352.582 | High | subset1 |
| TCGA-FS-A1ZE-06A-11R-A18T-07 | 1833.423 | Low | subset1 |
| TCGA-EB-A4XL-01A-11R-A27Q-07 | 1943.764 | NA | subset1 |
| TCGA-XV-A9W5-01A-11R-A38C-07 | 2341.378 | High | subset1 |
| TCGA-EE-A3J4-06A-11R-A20F-07 | 2020.619 | NA | subset1 |
| TCGA-WE-A8K1-06A-21R-A37K-07 | 2234.531 | NA | subset1 |
| TCGA-EE-A2MJ-06A-11R-A18S-07 | 1899.382 | Low | subset1 |
| TCGA-DA-A1I5-06A-11R-A18T-07 | 2159.440 | NA | subset1 |
| TCGA-D3-A8GV-06A-11R-A37K-07 | 2415.200 | High | subset1 |
| TCGA-EB-A1NK-01A-11R-A18T-07 | 2610.533 | High | subset1 |
| TCGA-EE-A17Y-06A-11R-A18T-07 | 2010.815 | NA | subset1 |
| TCGA-D3-A2J7-06A-11R-A18T-07 | 1852.938 | Low | subset1 |
| TCGA-EE-A2GH-06A-11R-A18T-07 | 2353.014 | High | subset1 |
| TCGA-RP-A6K9-06A-41R-A352-07 | 2191.958 | NA | subset1 |
| TCGA-EE-A2GS-06A-12R-A18S-07 | 2147.036 | NA | subset2 |
| TCGA-EE-A2MH-06A-11R-A18S-07 | 2276.324 | NA | subset1 |
| TCGA-EB-A430-01A-11R-A24X-07 | 2618.281 | High | subset1 |
| TCGA-EE-A3AG-06A-31R-A18S-07 | 2084.292 | NA | subset1 |
| TCGA-D3-A51K-06A-11R-A266-07 | 3450.853 | High | subset1 |
| TCGA-EE-A17X-06A-11R-A18S-07 | 2312.433 | NA | subset1 |
| TCGA-ER-A196-01A-11R-A18T-07 | 2102.420 | NA | subset1 |
| TCGA-BF-A1PX-01A-12R-A18T-07 | 2431.812 | High | subset1 |
| TCGA-FR-A728-01A-11R-A32P-07 | 2239.871 | NA | subset1 |
| TCGA-FR-A7U8-06A-21R-A352-07 | 2662.407 | High | subset1 |
| TCGA-EE-A3J5-06A-11R-A20F-07 | 2054.339 | NA | subset1 |
| TCGA-HR-A5NC-01A-11R-A27Q-07 | 1966.957 | NA | subset1 |
| TCGA-3N-A9WC-06A-11R-A38C-07 | 2032.608 | NA | subset1 |
| TCGA-FS-A1ZN-01A-11R-A18T-07 | 1860.979 | Low | subset1 |
| TCGA-EE-A29W-06A-11R-A18U-07 | 1871.479 | Low | subset1 |
| TCGA-D9-A6EA-06A-11R-A311-07 | 2393.943 | High | subset1 |
| TCGA-W3-A824-06A-21R-A352-07 | 2300.339 | NA | subset1 |
| TCGA-EE-A2MF-06A-11R-A21D-07 | 1771.066 | Low | subset1 |
| TCGA-YG-AA3P-06A-11R-A38C-07 | 2343.788 | High | subset1 |
| TCGA-DA-A3F3-06A-11R-A20F-07 | 1871.956 | Low | subset1 |
| TCGA-FR-A8YD-06A-11R-A37K-07 | 2969.640 | High | subset2 |
| TCGA-QB-A6FS-06A-11R-A311-07 | 3457.167 | High | subset1 |
| TCGA-FS-A1ZY-06A-11R-A18S-07 | 2479.247 | High | subset1 |
| TCGA-EB-A44Q-06A-11R-A266-07 | 1920.895 | NA | subset1 |
| TCGA-WE-AAA0-06A-11R-A38C-07 | 2132.450 | NA | subset2 |
| TCGA-D3-A2JD-06A-11R-A18T-07 | 2929.576 | High | subset1 |
| TCGA-FR-A3R1-01A-11R-A239-07 | 1710.592 | Low | subset1 |

| | | | |
|---|---|---|---|
| TCGA-BF-A3DJ-01A-11R-A20F-07 | 2116.079 | NA | subset1 |
| TCGA-EE-A2GK-06A-11R-A18S-07 | 2087.017 | NA | subset1 |
| TCGA-EE-A29M-06A-11R-A18T-07 | 1895.099 | Low | subset1 |
| TCGA-FR-A3YO-06A-11R-A239-07 | 2886.871 | High | subset2 |
| TCGA-BF-A1PU-01A-11R-A18S-07 | 2413.662 | High | subset1 |
| TCGA-D3-A2JC-06A-11R-A18T-07 | 2294.340 | NA | subset1 |
| TCGA-D3-A1QB-06A-11R-A18T-07 | 2234.045 | NA | subset1 |
| TCGA-D3-A3C6-06A-12R-A18U-07 | 1970.091 | NA | subset1 |
| TCGA-ER-A199-06A-11R-A18T-07 | 2123.379 | NA | subset1 |
| TCGA-GN-A8LL-06A-21R-A37K-07 | 3005.469 | High | subset1 |
| TCGA-XV-AB01-06A-12R-A40A-07 | 3383.119 | High | subset1 |
| TCGA-EE-A2GP-06A-11R-A18S-07 | 1634.481 | Low | subset1 |
| TCGA-EE-A2MG-06A-11R-A18T-07 | 1850.515 | Low | subset1 |
| TCGA-BF-A5ER-01A-12R-A27Q-07 | 3188.303 | High | subset1 |
| TCGA-D3-A2J8-06A-11R-A18T-07 | 2195.415 | NA | subset1 |
| TCGA-DA-A1HY-06A-11R-A18T-07 | 1835.734 | Low | subset1 |
| TCGA-DA-A95X-06A-11R-A37K-07 | 1876.848 | Low | subset1 |
| TCGA-D3-A3C1-06A-12R-A18S-07 | 2675.555 | High | subset1 |
| TCGA-EB-A24C-01A-11R-A18T-07 | 1955.662 | NA | subset1 |
| TCGA-FS-A4F0-06A-11R-A24X-07 | 1692.776 | Low | subset1 |
| TCGA-EE-A3JD-06A-11R-A20F-07 | 1698.645 | Low | subset1 |
| TCGA-D9-A6EC-06A-11R-A311-07 | 2210.169 | NA | subset1 |
| TCGA-EE-A2MS-06A-11R-A18S-07 | 2148.495 | NA | subset1 |
| TCGA-FS-A4F9-06A-11R-A24X-07 | 1902.278 | Low | subset1 |
| TCGA-BF-A1PZ-01A-11R-A18S-07 | 2393.962 | High | subset1 |
| TCGA-EE-A20H-06A-11R-A18S-07 | 1814.914 | Low | subset1 |
| TCGA-FS-A4FD-06A-11R-A266-07 | 2136.244 | NA | subset1 |
| TCGA-D3-A5GR-06A-11R-A27Q-07 | 1829.309 | Low | subset1 |
| TCGA-EB-A57M-01A-51R-A311-07 | 2890.017 | High | subset1 |
| TCGA-EB-A299-01A-21R-A18U-07 | 2382.678 | High | subset1 |
| TCGA-EE-A20B-06A-11R-A18U-07 | 2121.893 | NA | subset1 |
| TCGA-EE-A2A5-06A-11R-A18T-07 | 2049.528 | NA | subset1 |
| TCGA-EB-A4IS-01A-21R-A266-07 | 2130.696 | NA | subset1 |
| TCGA-D3-A3MV-06A-11R-A21D-07 | 1986.410 | NA | subset1 |
| TCGA-EB-A6QY-01A-12R-A32P-07 | 1909.892 | Low | subset1 |
| TCGA-DA-A1I1-06A-12R-A18U-07 | 1948.713 | NA | subset1 |
| TCGA-ER-A19D-06A-11R-A18S-07 | 1773.560 | Low | subset1 |
| TCGA-3N-A9WB-06A-11R-A38C-07 | 2184.664 | NA | subset1 |
| TCGA-ER-A19B-06A-11R-A18S-07 | 2173.003 | NA | subset1 |
| TCGA-D3-A3MR-06A-11R-A21D-07 | 2390.621 | High | subset1 |
| TCGA-FW-A3I3-06A-11R-A21D-07 | 1884.355 | Low | subset1 |

| | | | |
|---|---|---|---|
| TCGA-EE-A2GC-06A-11R-A18T-07 | 1644.236 | Low | subset1 |
| TCGA-EB-A4OY-01A-11R-A266-07 | 3620.139 | High | subset1 |
| TCGA-ER-A3PL-06A-11R-A239-07 | 1991.540 | NA | subset1 |
| TCGA-EE-A2M7-06A-11R-A18U-07 | 1919.662 | NA | subset1 |
| TCGA-W3-A825-06A-11R-A352-07 | 2208.690 | NA | subset1 |
| TCGA-EB-A44O-01A-11R-A266-07 | 1785.798 | Low | subset1 |
| TCGA-D3-A8GC-06A-11R-A37K-07 | 1959.354 | NA | subset1 |
| TCGA-EB-A44N-01A-11R-A266-07 | 1988.082 | NA | subset1 |
| TCGA-DA-A960-01A-11R-A37K-07 | 3006.129 | High | subset1 |
| TCGA-FR-A7UA-06A-32R-A352-07 | 1883.542 | Low | subset1 |
| TCGA-ER-A19Q-06A-11R-A18U-07 | 2150.867 | NA | subset1 |
| TCGA-BF-AAP8-01A-11R-A40A-07 | 2284.551 | NA | subset1 |
| TCGA-D9-A4Z5-01A-11R-A266-07 | 2118.556 | NA | subset1 |
| TCGA-BF-AAP1-01A-11R-A39D-07 | 2271.943 | NA | subset1 |
| TCGA-OD-A75X-06A-12R-A32P-07 | 1690.400 | Low | subset1 |
| TCGA-EB-A6R0-01A-12R-A32P-07 | 1875.735 | Low | subset1 |
| TCGA-ER-A198-06A-11R-A18T-07 | 1774.436 | Low | subset1 |
| TCGA-WE-A8ZT-06A-11R-A37K-07 | 1534.970 | Low | subset1 |
| TCGA-EE-A20I-06A-11R-A18U-07 | 2230.602 | NA | subset1 |
| TCGA-FS-A1Z0-06A-11R-A18T-07 | 1778.477 | Low | subset1 |
| TCGA-ER-A19L-06A-12R-A18S-07 | 1909.123 | Low | subset1 |
| TCGA-FS-A1ZF-06A-12R-A18S-07 | 2206.751 | NA | subset1 |
| TCGA-DA-A3F5-06A-11R-A20F-07 | 2173.165 | NA | subset1 |
| TCGA-BF-AAP4-01A-11R-A40A-07 | 2253.955 | NA | subset1 |
| TCGA-FR-A69P-06A-21R-A311-07 | 1825.268 | Low | subset1 |
| TCGA-W3-AA1W-06A-11R-A38C-07 | 2113.790 | NA | subset1 |
| TCGA-D3-A51J-06A-11R-A266-07 | 2105.771 | NA | subset1 |
| TCGA-EB-A4P0-01A-41R-A266-07 | 2607.393 | High | subset1 |
| TCGA-D3-A5GN-06A-11R-A27Q-07 | 1867.903 | Low | subset1 |
| TCGA-ER-A2NE-06A-21R-A18T-07 | 2045.589 | NA | subset1 |
| TCGA-EE-A29T-06A-11R-A18T-07 | 1897.981 | Low | subset1 |
| TCGA-EE-A2GO-06A-11R-A18S-07 | 2001.053 | NA | subset1 |
| TCGA-EE-A3AD-06A-11R-A18S-07 | 1939.183 | NA | subset1 |
| TCGA-D9-A1JX-06A-11R-A18S-07 | 2034.000 | NA | subset1 |
| TCGA-EE-A3JH-06A-11R-A21D-07 | 1983.903 | NA | subset1 |
| TCGA-EE-A3JI-06A-11R-A21D-07 | 1692.108 | Low | subset2 |
| TCGA-EE-A29D-06A-11R-A18T-07 | 1953.450 | NA | subset1 |
| TCGA-GN-A26D-06A-11R-A18T-07 | 2011.655 | NA | subset1 |
| TCGA-GN-A4U8-06A-11R-A32P-07 | 2123.931 | NA | subset1 |
| TCGA-ER-A19H-06A-12R-A18S-07 | 1714.553 | Low | subset1 |
| TCGA-EE-A2ML-06A-11R-A18S-07 | 1785.597 | Low | subset1 |

| | | | |
|---|---|---|---|
| TCGA-ER-A3ES-06A-11R-A20F-07 | 1965.942 | NA | subset1 |
| TCGA-IH-A3EA-01A-11R-A20F-07 | 3069.875 | High | subset1 |
| TCGA-FS-A1ZG-06A-11R-A18T-07 | 1905.587 | Low | subset1 |
| TCGA-D9-A3Z1-06A-11R-A239-07 | 1895.923 | Low | subset1 |
| TCGA-YD-A9TB-06A-12R-A40A-07 | 1846.249 | Low | subset1 |
| TCGA-ER-A2ND-06A-11R-A18T-07 | 2667.364 | High | subset1 |
| TCGA-EB-A5SF-01A-11R-A311-07 | 2844.826 | High | subset1 |
| TCGA-D3-A2JK-06A-11R-A18S-07 | 2894.867 | High | subset1 |
| TCGA-FW-A3TU-06A-11R-A239-07 | 3110.687 | High | subset1 |
| TCGA-EB-A5FP-01A-11R-A27Q-07 | 1781.826 | Low | subset1 |
| TCGA-ER-A19E-06A-11R-A18S-07 | 2002.127 | NA | subset1 |
| TCGA-ER-A19M-06A-61R-A239-07 | 1969.996 | NA | subset1 |
| TCGA-D3-A2JO-06A-11R-A18S-07 | 1919.488 | NA | subset1 |
| TCGA-D3-A2JA-06A-11R-A18T-07 | 1929.326 | NA | subset1 |
| TCGA-GF-A3OT-06A-23R-A239-07 | 1986.806 | NA | subset2 |
| TCGA-ER-A19N-06A-11R-A18S-07 | 1962.540 | NA | subset1 |
| TCGA-EE-A2MT-06A-11R-A18S-07 | 1947.686 | NA | subset1 |
| TCGA-FS-A4FB-06A-11R-A266-07 | 1895.969 | Low | subset1 |
| TCGA-GN-A4U8-11A-11R-A32P-07 | 2167.931 | NA | subset1 |
| TCGA-EB-A5UM-01A-11R-A311-07 | 2956.515 | High | subset1 |
| TCGA-FS-A1ZW-06A-12R-A18T-07 | 1665.910 | Low | subset1 |
| TCGA-WE-AAA4-06A-12R-A38C-07 | 2905.925 | High | subset1 |
| TCGA-ER-A19C-06A-11R-A18S-07 | 1911.208 | Low | subset1 |

**Table S13b. Table for tFA of Hugo Melanoma cohort**

| Sample ID | tFA | Group |
|---|---|---|
| Pt1 | 2232.637 | Low |
| Pt2 | 2731.533 | High |
| Pt4 | 2598.993 | Low |
| Pt5 | 2313.247 | Low |
| Pt6 | 2283.931 | Low |
| Pt7 | 2854.153 | High |
| Pt8 | 3014.902 | High |
| Pt9 | 2445.262 | Low |
| Pt10 | 2452.072 | Low |
| Pt12 | 2362.878 | Low |
| Pt13 | 2301.223 | Low |
| Pt14 | 2384.946 | Low |
| Pt15 | 3063.404 | High |
| Pt16 | 3016.530 | High |
| Pt19 | 3672.679 | High |
| Pt20 | 2736.222 | High |
| Pt22 | 2258.911 | Low |
| Pt23 | 2706.164 | High |
| Pt25 | 3924.079 | High |
| Pt27A | 2668.609 | Low |
| Pt27B | 2637.624 | Low |
| Pt28 | 2037.138 | Low |
| Pt29 | 2697.004 | High |
| Pt31 | 3363.477 | High |
| Pt32 | 2847.857 | High |
| Pt35 | 2508.802 | Low |
| Pt37 | 3509.842 | High |
| Pt38 | 2764.535 | High |
